# A Bivalent Molecular Glue Linking Lysine Acetyltransferases to Oncogene-induced Cell Death

**DOI:** 10.1101/2025.03.14.643404

**Authors:** Meredith N. Nix, Sai Gourisankar, Roman C. Sarott, Brendan G. Dwyer, Sabin. A. Nettles, Michael M. Martinez, Hind Abuzaid, Haopeng Yang, Yanlan Wang, Juste M. Simanauskaite, Bryan A. Romero, Hannah M. Jones, Andrey Krokhotin, Tara N. Lowensohn, Lei Chen, Cara Low, Mark M. Davis, Daniel Fernandez, Tinghu Zhang, Michael R. Green, Stephen M. Hinshaw, Nathanael S. Gray, Gerald R. Crabtree

**Affiliations:** Department of Chemical and Systems Biology, Stanford University, Stanford, CA, USA; Department of Chemistry, Stanford University, Stanford, CA, USA; Department of Pathology, Stanford University, Stanford, CA, USA; Department of Developmental Biology, Stanford University, Stanford, CA, USA; Department of Lymphoma-& Myeloma, University of Texas MD Anderson Cancer Center, Houston, TX, USA; Department of Microbiology and Immunology, Stanford University, Stanford, CA, USA; Macromolecular Structure, Nucleus at Sarafan ChEM-H, Stanford University, Stanford, CA, USA

**Keywords:** lysine acetyltransferases, induced proximity, lymphoma, transcription

## Abstract

Developing cancer therapies that induce robust death of the malignant cell is critical to prevent relapse. Highly effective strategies, such as immunotherapy, exemplify this observation. Here we provide the structural and molecular underpinnings for an approach that leverages chemical induced proximity to produce specific cell killing of diffuse large B cell lymphoma, the most common non-Hodgkin’s lymphoma. We develop KAT-TCIPs (lysine acetyltransferase transcriptional/epigenetic chemical inducers of proximity) that redirect p300 and CBP to activate programmed cell death genes normally repressed by the oncogenic driver, BCL6. Acute treatment rapidly reprograms the epigenome to initiate apoptosis and repress c-MYC. The crystal structure of the chemically induced p300-BCL6 complex reveals how chance interactions between the two proteins can be systematically exploited to produce the exquisite potency and selectivity of KAT-TCIPs. Thus, the malignant function of an oncogenic driver can be co-opted to activate robust cell death, with implications for precision epigenetic therapies.

## INTRODUCTION

The paralogous lysine acetyltransferases (KATs) p300 (<u>E</u>1A-associated protein <u>p300</u>, encoded by *EP300*) and CBP (<u>C</u>REB-<u>b</u>inding <u>p</u>rotein, encoded by *CREBBP*), are essential regulators of gene expression in healthy and malignant cells^1^. p300 and CBP (hereafter: p300/CBP) bind and catalyze histone acetylation, interact with transcription factors (TFs), and activate transcription at promoters and enhancers^2,3^. A catalytic KAT domain (86% identity between p300/CBP)^4–6^, an acetyl-lysine-binding bromodomain (97% identity), and several other conserved protein-interaction domains facilitate these functions^7,8^. In diverse cancers, p300/CBP support the expression of oncogenic networks by activating clusters of gene-regulatory regions called super-enhancers^8–13^. No p300/CBP inhibitor has received regulatory approval for oncology^14–18^. Given the physiological role of p300/CBP in activating transcription, we sought to develop small molecules that redirect them to induce the expression of proapoptotic pathways in cancer.

We pursued this objective by developing chemical inducers of proximity (CIPs). CIPs are small molecules that rewire protein-protein interactions. The realization that induced proximity governs diverse cellular processes such as signal transduction, post-translational-modification, and epigenetic regulation^19^ has led to the development of CIPs for therapeutic purposes. Proteolysis-targeting chimeras (PROTACs), which rely on inducing the proximity of a ubiquitin ligase to degrade a target protein, exemplify this concept^20^. Cyclosporin A^21^, FK506^22^ (tacrolimus) and rapamycin^23^ (sirolimus) belong to a related class of clinical proximity-inducing small molecules. Structural studies by Schreiber and Clardy showed that these compounds facilitate the formation of protein complexes by creating a composite binding surface containing the small molecule and the two binding proteins^24^. A high degree of cooperative binding enables the potency and selectivity of the CIP^25^. Small molecules that facilitate the cooperative assembly of complexes between unrelated proteins have come to be known as molecular glues^26^.

While CIPs are emerging as preclinical tools to degrade p300/CBP in hematological malignancies and solid cancers^27–30^, the development of p300/CBP-redirecting CIPs remains limited. Thus far, efforts have largely relied on transgene overexpression^31^, protein tags^32–34^, or large, DNA-binding pyrrole-imidazole polyamide moieties^35^; these technologies are not easily translatable to the clinic. Two notable recent exceptions are CIPs^36,37^ that harness KAT activity to inactivate a relatively rare^38^, mutant form of p53 via targeted p53 acetylation in genetically unmodified cells.

Our laboratories recently introduced a class of CIPs termed transcriptional/epigenetic chemical inducers of proximity (TCIPs) that relocalize transcriptional activators to chromatin bound by DNA sequence-specific TFs^39,40^. We previously synthesized TCIPs that redirect the RNA Polymerase II elongation-associated factors BRD4 and CDK9 to BCL6, a TF that represses pro-apoptotic and growth arrest genes^41–43^ and is overexpressed in 40 to 60% of diffuse large B cell lymphomas (DLBCLs)^44^. These molecules rewire BCL6 to activate transcriptionally silent proapoptotic pathways and potently kill cells. They offer an alternative therapeutic strategy to kill DLBCL cells when compared to inhibitors or degraders of BCL6^45–49^ and have been chemically optimized for *in vivo* administration^40^. The broad applicability of this approach hinges on (i) identifying a diverse set of activators that can be rewired and (ii) developing molecular glues that induce favorable protein-protein interactions. The latter requires structural studies of critical ternary complexes to drive chemical optimization of cooperativity.

Here, we developed lysine acetyltransferase TCIPs (KAT-TCIPs) that redirect p300/CBP to BCL6. We used a systematic chemical design cascade to discover molecules that are exquisitely selective to both protein partners and potent in DLBCL cells (cell proliferation IC_50_ ∼0.8 nM). KAT-TCIPs redistributed p300/CBP-catalyzed chromatin acetylation and activated BCL6-regulated gene expression. These molecular effects led to potent induction of pro-apoptotic proteins and durable repression of the oncogene c-MYC (IC_50_ ∼ 0.8 nM). A co-crystal structure of the KAT-TCIP in complex with p300 and BCL6 and biophysical analyses demonstrated that selectivity and potency arise from the chance formation of complementary protein-protein interactions on the composite drug-protein interface. Thus, CIPs that redirect p300/CBP can be systematically designed to be precise and powerful killers of cancer cells.

## RESULTS

### Design and activity of KAT-TCIPs

To explore the potential of harnessing co-activating lysine acetyltransferase activity for transcriptional activation, we synthesized a library of bivalent compounds designed to recruit p300/CBP to BCL6-bound genes (Fig. 1A). These bivalent compounds link the BCL6 BTB-domain-binding ligand BI-3812^48^ to p300/CBP-specific bromodomain (BD) inhibitors, including GNE-781^50^ and Cmpd33^51^, via conjugatable handles identified from X-ray co-crystal structures (Supplemental Fig. 1A, B). A diverse library of alkyl, PEG, and rigid linkers was employed to increase the probability of generating cell-permeable TCIPs that induce productive ternary complexes (Supplemental Fig. 1A, 1B).

**Figure 1.**
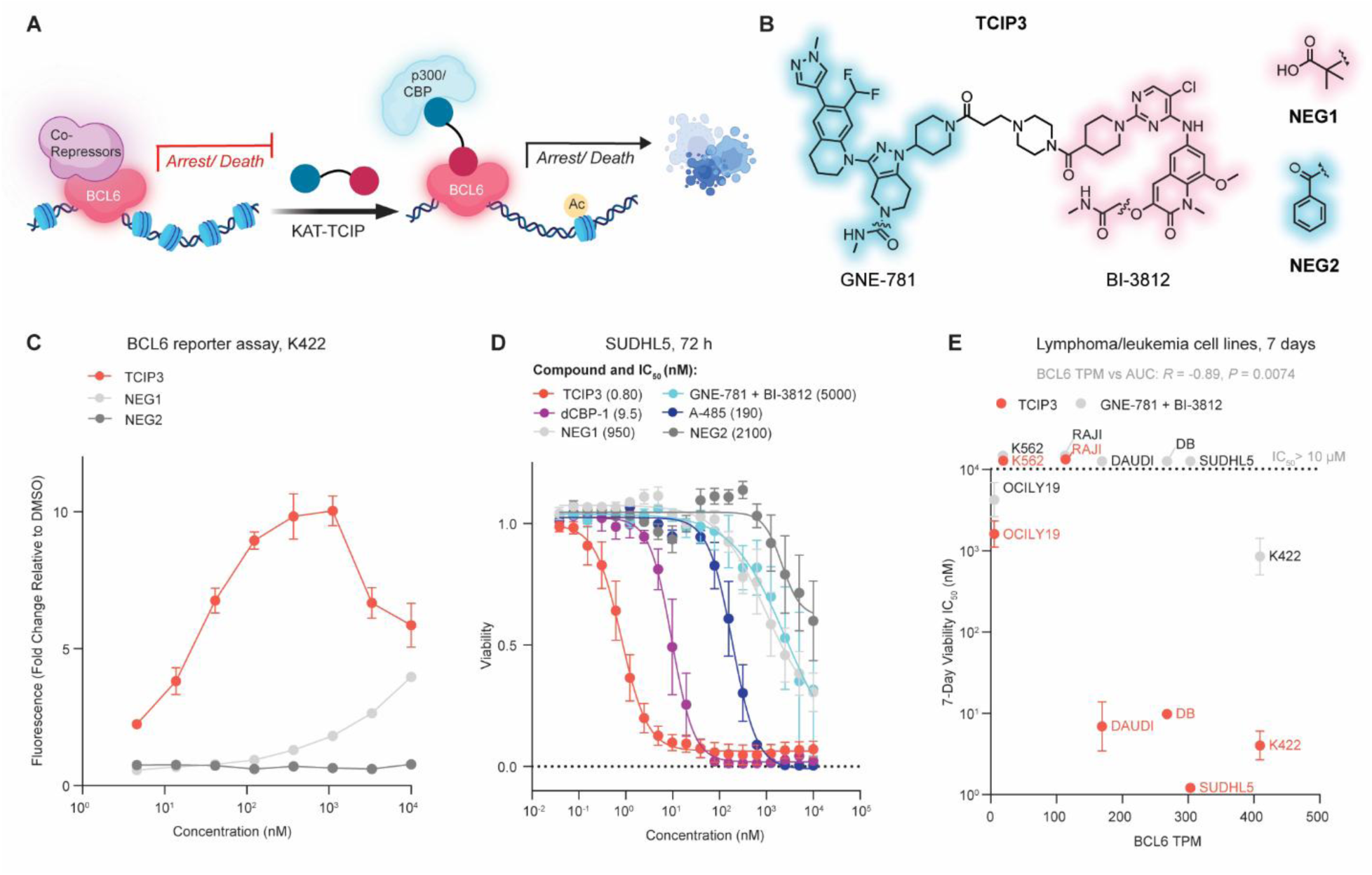
Design and Activity of KAT-TCIPs. **(A)** KAT-TCIPs targeting p300/CBP are designed to de-repress cell death and cell cycle arrest pathways controlled by the transcription factor BCL6 in DLCBL cells. **(B)** Structure of **TCIP3, NEG1**, which retains binding to p300/CBP, and **NEG2**, which retains binding to BCL6. **(C)** Activation of a BCL6-repressed GFP reporter construct integrated into K422 cells after compound treatment for 24 h; 3 biological replicates, mean ± s.e.m. **(D)** Cell-killing potencies of compounds after 72 h of treatment in SUDHL5 cells; 3-5 biological replicates, mean ± s.e.m. **(E)** IC_50_ values (nM) of antiproliferation after 7 days of compound treatment in DLBCL and leukemia cells plotted against *BCL6* expression in transcripts/million (TPM)^95^; 3-4 biological replicates, mean ± s.d.; *R* of BCL6 TPM vs AUC computed by Pearson’s correlation and *P-*value computed by two-sided Student’s t-test.

To test the KAT-TCIP library, we used a BCL6-controlled green fluorescent protein (GFP) reporter gene in the DLBCL cell line KARPAS422 (K422)^39^. One compound, which we named **TCIP3** (Fig. 1B), produced the highest GFP signal, 10-times greater than DMSO treatment (Fig. 1C and Supplemental Fig. 1C). We observed a hook effect in GFP activation, a characteristic of bivalent molecules in which high concentrations of a compound saturate binding to either protein partner and prevent productive ternary complex formation^52,53^. To distinguish the effects of ternary complex formation from p300/CBP^BD^ or BCL6^BTB^ inhibition while controlling for drug size and cellular permeability, we synthesized negative controls containing the **TCIP3** linker but that lack the ability to bind either BCL6^BTB^ (**NEG1**) or p300/CBP^BD^ (**NEG2**) (Fig. 1B). When compared with **TCIP3**, **NEG1** and **NEG2** remained similarly membrane-permeable and potent binders of either p300/CBP^BD^ or BCL6^BTB^, respectively, as measured by cellular probe-displacement assays (nanoBRET^54^) (Supplemental Fig. 1D, E). Neither **NEG1** nor **NEG2** activated GFP expression, supporting ternary complex-dependent activity (Fig. 1C).

Given the ability of p300/CBP KAT-TCIPs to activate the BCL6 reporter gene, we investigated their effects on the viability of DLBCL. In a 72-hour cell viability assay, **TCIP3** inhibited the proliferation of SUDHL5 DLBCL cells with an average IC_50_ of 0.8 nM (Fig. 1D). **TCIP3** was over 1,000 times more cytotoxic than either **NEG1** or **NEG2**, and nearly all GNE-781-based KAT-TCIPs were 10 – 38,000-times more potent than the co-inhibition of p300/CBP and BCL6 (Fig. 1D, Supplemental Fig. 2A). In these cell proliferation/viability assays, **TCIP3** was 238-times more potent than the p300/CBP KAT catalytic-site inhibitor A-485^15^ and 12 times more potent than a p300/CBP PROTAC dCBP-1^30^ (Fig. 1D). Thus, **TCIP3** produces a potent anti-proliferative effect in DLBCL cells.

**TCIP3** exhibited substantially reduced cytotoxicity in B and T cells isolated from primary tonsillar lymphocytes of two separate donors and in primary human fibroblasts, as compared to its effects on SUDHL5 cells, and was less toxic than GNE-781, dCBP-1, and A-485 (Supplemental Fig. 2B, 2C). Because **TCIP3** exhibited cancer-specific cytotoxicity and was less toxic than p300/CBP-targeting agents in untransformed cells, we assessed the generality of these observations by testing the sensitivity of seven different lymphoma and leukemia lines. These included six lymphoma lines with high, medium, and low levels of BCL6 expression and one leukemia cell line (K562) with negligible BCL6 (Fig. 1E). There was a statistically significant correlation between BCL6 levels and area under the curve (AUC) values upon **TCIP3** treatment (Pearson’s *R* = -0.89, *P* = 0.0074). Co-treatment of the same cell lines with GNE-781 and BI-3812 produced no such correlation. Over 7 days of treatment, **TCIP3** inhibited the viability of high-BCL6 lines SUDHL5, K422, DB, and DAUDI at comparable potencies (1.2 nM, 4.0 nM, 9.8 nM, and 6.9 nM, respectively), approximately 200 to 1000 times greater than the co-treatment of the inhibitors GNE-781 and BI-3812 (Fig. 1E, Supplemental Fig. 2D, 2E). In cells with low BCL6 levels, **TCIP3** performed equivalently to the co-treatment of GNE-781 and BI-3812 (Fig. 1E, Supplemental Fig. 2D, 2E).

Each BCL6-high lymphoma cell line we tested originated from a germinal center B cell (GCB)-derived lymphoma in which dysregulation of BCL6 drives cancer progression by repressing pro-apoptotic and growth arrest genes^41^. p300/CBP are constitutively expressed at similar levels and are essential genes in most cells^55^. If **TCIP3** suppresses malignant cell growth by using high BCL6 expression to sequester p300/CBP from its acetyltransferase and transcriptional substrates, increasing BCL6 levels would increase cellular sensitivity. To directly test this model, we overexpressed the BTB domain of BCL6 in BCL6-low K562 cells. This construct codes for the BTB domain, which binds **TCIP3,** but not the DNA-binding zinc finger domains of BCL6 that might localize to death genes on chromatin. This line is otherwise sensitive to p300/CBP acetyltransferase inhibition^55^. Overexpressing BCL6^BTB^ did not increase sensitivity to **TCIP3** (Supplemental Fig. 2F). Thus, **TCIP3** activity requires not only high levels of BCL6, but also BCL6 that is actively repressing antiproliferative and cell death gene expression. Together, our data indicate that chemically induced recruitment of p300/CBP to BCL6 by **TCIP3** produces a potent, gain-of-function effect distinct from the effects caused by protein inhibition, sequestration, or degradation.

### Selective ternary complex formation in cells is required for activity

We investigated the relationship between p300/CBP-BCL6 ternary complex formation and the activity of KAT-TCIPs. First, to characterize the ternary complex biochemically, we assessed the KAT-TCIP library using TR-FRET with labeled recombinant p300^BD^, CBP^BD^, and BCL6^BTB^ domains (Supplemental Fig. 3A). Almost all compounds in our KAT-TCIP library induced p300/CBP-BCL6 ternary complexes and had antiproliferative activity (Supplemental Fig. 3B). **TCIP3** showed one of the highest increases in p300^BD^-BCL6^BTB^ TR-FRET signal with an AUC 23-times higher than **NEG1** (16-times higher in CBP^BD^-BCL6^BTB^ TR-FRET) and 18-times higher than **NEG2** (25-times higher in CBP^BD^-BCL6^BTB^ TR-FRET) (Supplemental Fig. 3C, D).

To probe how the formation of the ternary complex affects cellular viability, we treated SUDHL5 cells with 1 nM of **TCIP3** and co-treated these cells with one of three different BCL6^BTB^ inhibitors: BI-3812^48^, GSK137^49^, and an analog of CCT373566 (CCT373566a, Supplemental File 1) with a matched exit vector to BI-3812^46^. We carried out a similar **TCIP3** co-treatment experiment with the p300/CBP^BD^ inhibitor, GNE-781^50^. Co-treatment with each of these inhibitors buffered the effects of **TCIP3** on cell viability (Fig. 2A, B). These results establish that the activity of **TCIP3** depends on its dual engagement with p300/CBP and BCL6.

**Figure 2.**
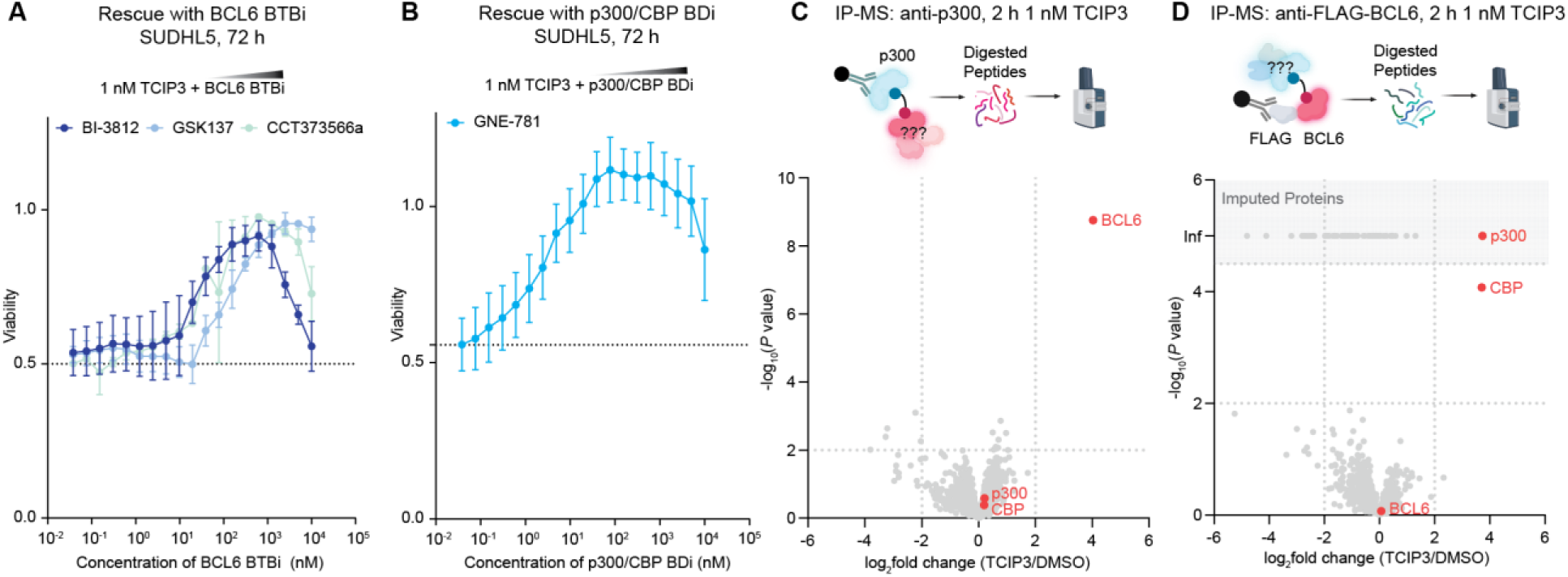
TCIP3 Kills Cells Via Chemically Induced Proximity (CIP) **(A)** Measurement of cell viability after competitive titration of constant 1 nM **TCIP3** with BCL6^BTB^ domain inhibitors (BI-3812^48^, GSK137^49^, or CCT373566a^46^), or **(B)** the p300/CBP bromodomain inhibitor GNE-781^50^; cells were treated simultaneously with **TCIP3** and the inhibitor or DMSO for 72 h; 3 biological replicates, mean ± s.e.m. **(B)** p300 immunoprecipitation-mass spectrometry (IP-MS) from SUDHL5 cells treated for 2 h with 1 nM **TCIP3**; plotted with cutoffs of log_2_(fold change)| ≥ 2 and *P* ≤ 0.01; 3 biological replicates. For **(C)** and **(D)**, *P*-values computed by a moderated t*-*test. **(C)** FLAG IP-MS from genomic knock-in FLAG-tagged *BCL6* SUDHL5 cells treated with 1 nM **TCIP3** for 2 h plotted with cutoffs of log_2_(fold change) ≥ 2 and *P* ≤ 0.01. Proteins that did not contain peptides for DMSO but contained peptides upon **TCIP3** treatment were imputed; 3 biological replicates.

The human genome encodes 62 structurally homologous bromodomains^56^ and 183 proteins with BTB domains^57^. To assess the selectivity of **TCIP3**, we conducted mass spectrometry of proteins immunoprecipitated (IP) with either p300 or BCL6 after 2 hours of 1 nM **TCIP3** addition to SUDHL5 cells (IP-MS) (Supplemental Table 1). Immunoprecipitation using an anti-p300 antibody produced exclusive enrichment of BCL6 (*P* = 1.8 x 10^-9^, log_2_(fold change) = 4.0) (Fig. 2C, Supplemental Fig. 3E, Supplemental Table 1). The sole proteins that showed statistically significant co-enrichment after immunoprecipitation with an anti-FLAG antibody from cells with FLAG inserted C-terminally to the *BCL6* gene (Methods) were CBP (*P* < 0.0001, log_2_(fold change) = 3.7) and p300 (no peptides detected in DMSO, imputed adj *P* < 0.05, log_2_(fold change) = 3.7) (Fig. 2D, Supplemental Fig. 3F, Supplemental Table 1). Despite BROMOscan data suggesting **TCIP3** may also engage BRPF1 at high compound concentrations (10 µM, Supplemental Fig. 3G), no other bromodomain-containing proteins were enriched (adj. *P* < 0.05, |log_2_(fold change)| > 2). To validate these results in a genetically unmodified cell, we immunoprecipitated BCL6 using an antibody raised against its native sequence and observed enrichment of both p300 and CBP (Supplemental Fig. 3H). Our proteomic data indicate that **TCIP3** is selective for both p300/CBP and BCL6 inside living cells.

### Rapid acetylation of BCL6 and BCL6-proximal chromatin

Biochemical specificity of **TCIP3** in forming a BCL6-p300/CBP ternary complex and the requirement of the complex for cytotoxicity implicates a proximity-dependent molecular mechanism of cell death. We examined the molecular effects of **TCIP3** in endogenous cells at the chromatin, RNA, and protein levels. We focused on acetylation of protein and chromatin substrates because p300/CBP has been described as acetylating a wide range of target substrates in its proximity^6^.

First, because p300/CBP-mediated acetylation of BCL6 at lysines 376, 377, and 379 in its unstructured repression domain 2 (RD2) was reported to derepress BCL6-target gene transcription by inhibiting the binding of repressive complexes to BCL6 without altering its DNA-binding capacity^58,59^, we assessed **TCIP3**-mediated BCL6 acetylation. Immunoprecipitation of all acetylated proteins from nuclear extracts with a pan-acetyl-lysine antibody showed that **TCIP3** induced a dose-dependent increase of acetylated BCL6 after 1 hour of addition beginning at 1 nM and increasing with compound dose, close to the antiproliferative IC_50_ value of 0.8 nM in these same cells (Fig. 3A).

**Figure 3.**
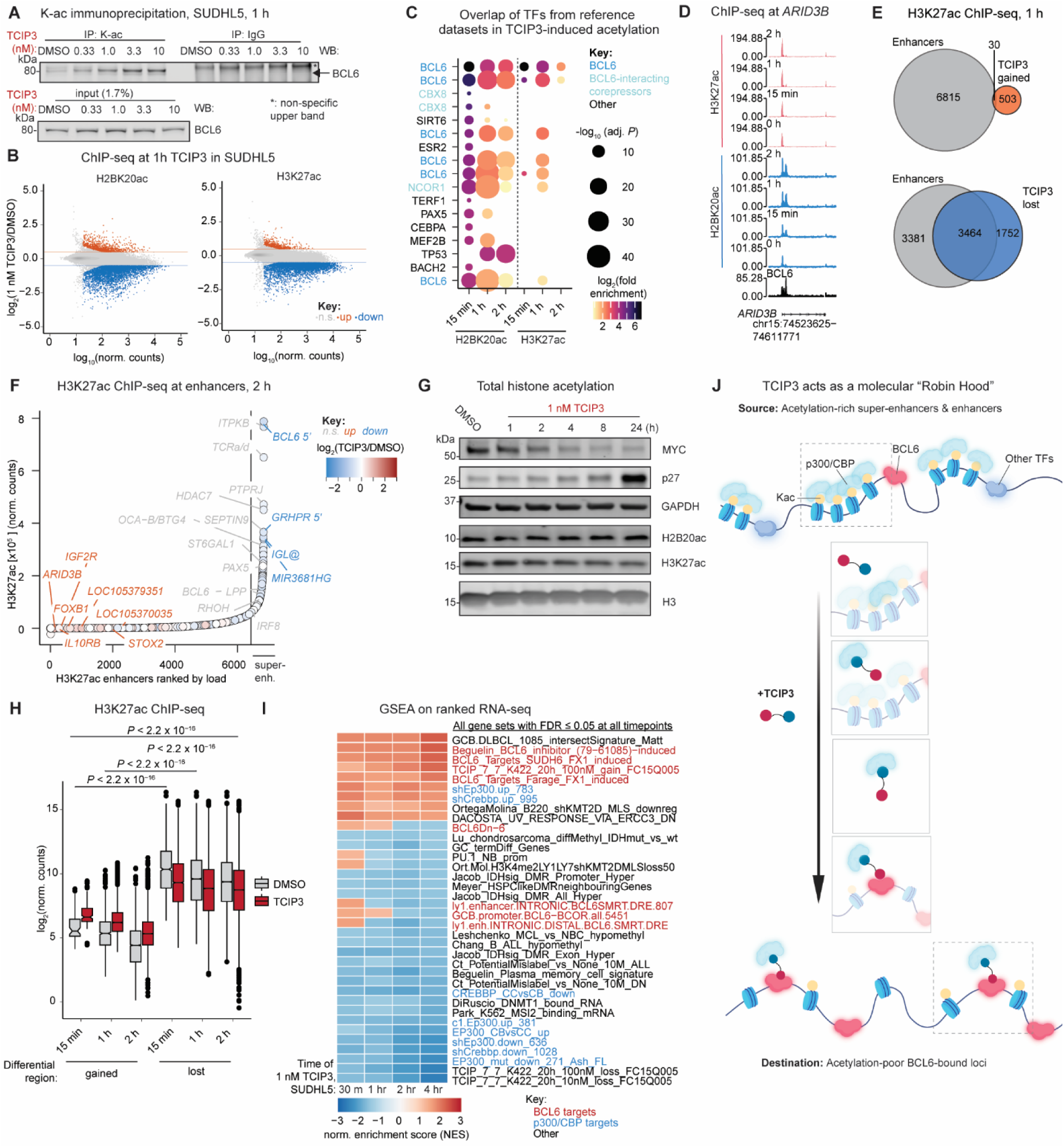
TCIP3 Redistributes p300/CBP Activity to BCL6 and Proximal Chromatin. **(A)** Acetylated lysine (K-ac)-IP and western blot (WB) for BCL6 after 1 h of **TCIP3** in SUDHL5 cells; representative of three biological replicates. **(B)** Changes in histone 3 lysine 27 and histone 2B lysine 20 acetylation (H3K27ac and H2BK20ac) as measured by chromatin immunoprecipitation sequencing (ChIP-seq) after 1 h of 1 nM **TCIP3** in SUDHL5 cells; significant: adj. *P* ≤ 0.05 and up, log_2_(fold change) ≥ 0.5; down, log_2_(fold change) ≤ -0.5; 2 biological replicates, *P*-values computed by two-sided Wald test and adjusted by multiple comparisons by Benjamini-Hochberg. **(C)** Enrichment of predicted transcription factor (TF) binding in gained H3K27ac and H2BK20ac peaks calculated by overlap with public ChIP-seq datasets from blood-lineage cells; full enrichment data in Supplemental Table 2; *P*-values computed by two-sided Fisher’s exact test and adjusted for multiple comparisons by Benjamini-Hochberg. **(D)** Induction of H2BK20ac and H3K27ac with time at the promoter of the BCL6 target gene *ARID3B*; BCL6 track is CUT&RUN in untreated SUDHL5 cells, tracks merged from two biological replicates and sequence-depth normalized and, for histone acetylation ChIP-seq, also input-subtracted. **(E)** Overlap of gained and lost H3K27ac peaks after 1 h of 1 nM **TCIP3** with annotated enhancers and super-enhancers in SUDHL5 cells. **(F)** Changes in H3K27ac at annotated enhancers and super-enhancers after 2 h of 1 nM **TCIP3**; significant: adj. *P* ≤ 0.05 and up, log_2_(fold change) ≥ 0.5; down, log_2_(fold change) ≤ -0.5; 2 biological replicates, *P*-values computed by two-sided Wald test and adjusted by multiple comparisons by Benjamini-Hochberg. **(G)** Western blot of SUDHL5 cells treated with 1 nM **TCIP3** for the indicated time periods; representative of 3 biological replicates. **(H)** Comparison of H3K27ac loading at differential regions at 15 min, 1 h, and 2 h of 1 nM **TCIP3**; differential regions defined as in **(A)**, *P*-values adjusted by Tukey’s test after type II analysis of variance (ANOVA). **(I)** Gene set enrichment analysis of ranked log_2_(fold change) in gene expression measured by RNA-sequencing (RNA-seq) after 1 nM **TCIP3** in SUDHL5 cells; only all gene sets adj. *P* ≤ 0.05 at all timepoints displayed, positive normalized enrichment scores (NES) indicate gene sets enriched in **TCIP3**-induced genes while negative NES scores indicate sets enriched in decreased genes, *P*-values computed by permutation and adjusted for multiple comparisons by Benjamini-Hochberg. **(J)** Model of how **TCIP3** redistributes p300/CBP from super-enhancers to BCL6.

Next, we hypothesized that **TCIP3** can induce p300/CBP to acetylate histone tail lysines near chromosomal BCL6 binding sites. Of the potential histone lysines that undergo acetylation, histone H3-K27 and histone H2B-K20 are of particular interest (H3K27ac and H2BK20ac, respectively) since they co-localize with and are generally considered to mark active enhancers and promoters^60,61^. To examine the effects of **TCIP3** on histone acetylation, we conducted chromatin immunoprecipitation sequencing (ChIP-seq) for H2BK20ac and H3K27ac after addition of 1 nM **TCIP3** to SUDHL5 cells for 15 min, 1 hour, and 2 hours. 69,719 and 66,995 peaks across all timepoints were reconstructed for H2BK20ac and H3K27ac, respectively. Most of the variance between conditions (∼90%) was attributable to compound treatment (Supplemental Fig. 4A). Differential peak analysis detected large gains and losses of acetylation: after 1 hour, 936 H2BK20ac peaks were gained and 4,346 were lost, while 533 H3K27ac peaks were gained and 5,216 were lost (Fig. 3B).

Peaks that gained acetylation were enriched in published regions of BCL6 binding in human B-cell and blood cancer cell lines (Fig. 3C and Supplemental Table 2). Enrichment for BCL6-binding sites was more pronounced in regions that gained H2BK20ac than in regions that gained H3K27ac (Fig. 3C), consistent with the recently reported propensity of p300/CBP to catalyze H2BK20ac^60^. Loci that gained acetylation included pro-apoptotic BCL6 targets such as *ARID3B*^62^ (Fig. 3D). We also identified 7,379 BCL6 binding sites in untreated SUDHL5 cells by CUT&RUN^63^ (Supplemental Fig. 4B). Comparison with this set of BCL6-bound sites showed that 21% (H3K27ac) to 25% (H2BK20ac) of gained acetylation regions overlapped (Supplemental Fig. 4B). The rapidly induced acetylation detected indicates redirection of p300/CBP acetyltransferase activity to BCL6-proximal DNA by **TCIP3**.

### Redistribution of p300/CBP activity from oncogenic regulatory regions

In contrast to gains in histone acetylation, losses occurred at annotated enhancers and super-enhancers in SUDHL5 cells (Fig. 3E and Supplemental Fig. 4C). Losses of H3K27ac and H2BK20ac were modest in magnitude across these regions but statistically significant (adj. *P* ≤ 0.05) after 15 min, 1, and 2 hours of **TCIP3** treatment (Supplemental Fig. 4D). The greatest losses in H3K27ac were concentrated in super-enhancers, broad regions of elevated histone acetylation that regulate the expression of cell-identity and proliferation genes^10,11,64^ (Fig. 3F). These losses were observed at several super-enhancers proximal to master regulators of the germinal center B cell and oncogenic drivers (Fig. 3F), including the *BCL6* gene itself, which contains an intronic enhancer as well as a requisite super-enhancer 150 kilobases upstream^65^ (Supplemental Fig. 4E). Total H3K27ac and H2BK20ac levels did not decrease after 1 nM **TCIP3** treatment at these immediate timepoints (Fig. 3G), indicating that TCIP3 does not reduce global histone acetylation. This dose and timepoint was nevertheless sufficient to decrease c-MYC and increase the BCL6-target p27 (Fig. 3G).

We were intrigued by the fact that most of the regions that lost acetylation due to **TCIP3** (*i.e.*, enhancers and super-enhancers) were already among the highest-acetylated loci on chromatin before treatment (Fig. 3F, H and Supplemental Fig. 4F). We reasoned that since these regions are considered rich in p300/CBP^11^, **TCIP3** should produce decreases in p300/CBP-target gene expression concurrent with increases in BCL6-regulated genes. The redirection of transcriptional activity should also occur on comparable timescales to the dynamic changes in acetylation observed. RNA-sequencing (RNA-seq) after 30 min, 1, 2 and 4 hours of 1 nM **TCIP3** addition in SUDHL5 cells revealed large numbers of genes induced (1,510) and decreased (2,126) (adj. *P* ≤ 0.05, |log_2_(fold change)| ≥ 0.5) (Supplemental Fig. 4G). Consistent with our hypothesis, gene set enrichment analysis showed that BCL6-bound genes were up, and p300/CBP-regulated genes were down (Fig. 3I). Most but not all genes near enhancers and super-enhancers that lost H3K27ac decreased in expression (Supplemental Fig. 4H). Overall changes in H3K27ac and gene expression showed a weak but statistically significant positive correlation (Supplemental Fig. 4I), indicating a modest direct relationship between histone acetylation and transcription of nearby genes. This is consistent with published data^60,66^.

We conclude that increases in BCL6 target genes result from the combination of the acetylation and inactivation of BCL6 protein (Fig. 3A) and p300/CBP recruitment to BCL6 target genes (Fig. 3B). Meanwhile, redirection of p300/CBP produces decreases of gene expression at its normal targets (Fig. 3F). The effective molecular mechanism of **TCIP3** calls to mind the character of Robin Hood: redistributing p300/CBP from “rich” sites to acetylate and activate “poor” genes, directed by BCL6 (Fig. 3J).

### Redistribution partly phenocopies proteomic effects of p300/CBP degradation

Given that **TCIP3** redistributed p300/CBP activity, we compared its effects on the cell to those produced by p300/CBP degraders. We first assessed changes in protein expression upon a 10 nM **TCIP3** treatment or a 250 nM dCBP-1 treatment at 6 hours (Supplemental Fig. 5A) and 24 hours (Fig. 4A, Supplemental Fig. 5B). **TCIP3** increased the abundance of both p300 and CBP (p300: adj. *P* ≤ 0.01, log_2_(fold change) = 0.37; CBP: adj. *P* ≤ 0.05, log_2_(fold change) = 0.58) (Fig. 4A). In contrast, and as expected^30^, 250 nM dCBP-1 treatment induced statistically significant losses in p300/CBP (p300: adj. *P* ≤ 0.05, log_2_(fold change) = -2.7; CBP: adj. *P* ≤ 0.01, log_2_(fold change) = -1.8) (Supplemental Fig. 5A, B). Matched concentrations of **NEG1** and **NEG2** displayed no significant changes relative to DMSO at either timepoint (Supplemental Fig. 5A, B).

**Figure 4.**
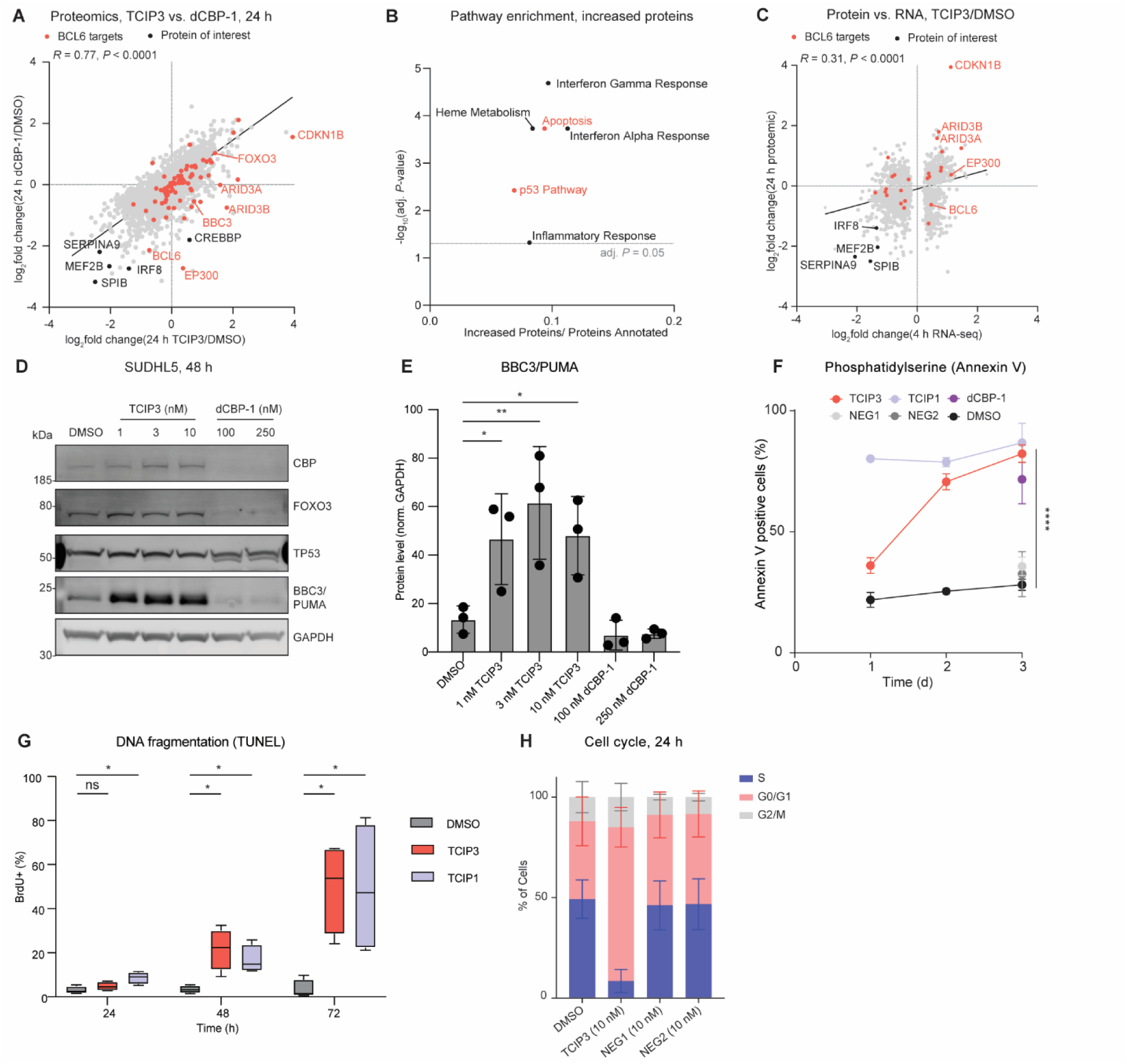
Activation of Apoptotic Signaling by TCIP3. **(A)** Comparison of whole-proteome profiling of SUDHL5 cells treated with 10 nM **TCIP3** or 250 nM of the p300/CBP degrader dCBP-1^30^ for 24 h; proteins labeled change statistically significantly (adj. *P* ≤ 0.05); 3 biological replicates; *P*-values computed using a moderated t-test and adjusted by Benjamini-Hochberg; *R* computed by Pearson’s correlation and *P-*value computed by two-sided Student’s t-test. **(B)** Signaling pathways (MSigDB Hallmark 2020) enriched in significantly increased proteins (adj. *P* < 0.05, log_2_(foldchange) > 1) after 24 h treatment of 10 nM **TCIP3** in SUDHL5. **(C)** Correlation of whole-proteome profiling of SUDHL5 cells treated with 10 nM **TCIP3** for 24 h to bulk transcriptomics (RNA-seq) performed on SUDHL5 cells treated with 1 nM TCIP3 for 4 h; only genes whose transcripts that change significantly (adj. *P* ≤ 0.05) were analyzed; 3 biological replicates; *P-*value for RNA-seq computed by two-sided Wald test and adjusted by Benjamini-Hochberg; *R* computed by Pearson’s correlation and *P-*value for Pearson’s correlation computed by two-sided Student’s t-test. **(D)** Western blot of pro-apoptotic proteins in SUDHL5 cells treated with indicated compounds and doses after 48 h; representative of 3 biological replicates. **(E)** Quantification of BBC3/PUMA protein levels from **(D);** *P-*values computed by Fisher’s LSD test after ANOVA; **: *P* < 0.01, *: *P* < 0.05; only the comparisons to DMSO were computed. 3 biological replicates, mean ± s.d. **(F)** Quantification of Annexin V-positive SUDHL5 cells treated with indicated compounds and doses at 24, 48, or 72 h; *P-*values adjusted by Tukey’s test after ANOVA. The following comparisons were significantly different from each other at 72 h (****: adj. *P* < 0.0001): 10 nM **TCIP3** vs DMSO, 10 nM **NEG1**, and 10 nM **NEG2** 72 h; 10 nM TCIP1^39^ vs DMSO, 10 nM NEG1, and 10 nM NEG2; 250 nM dCBP1 vs DMSO, 10 nm NEG1 (adj. *P =* 0.0023), and 10 nM NEG2 (adj. *P* = 0.0008). The following comparisons were significant at 48 h (****: adj. *P* < 0.0001): 10 nM **TCIP3** vs DMSO, 10 nM TCIP1 vs DMSO. The following comparisons were significant at 24 h: 10 nM **TCIP3** vs DMSO (****: adj. *P* < 0.0001), 10 nM TCIP1 vs DMSO (****: adj. *P* < 0.0001), 10 nM **TCIP3** vs 10 nM TCIP1 (*: adj. *P* = 0.0187). There were no significant differences between any other comparisons (adj. *P* ≥ 0.05). 3-12 biological replicates, mean ± s.e.m. **(G)** Concurrent analysis of apoptosis and cell cycle effects of TCIPs by terminal deoxynucleotidyl transferase dUTP nick end labeling (TUNEL) and total DNA content co-staining of SUDHL5 cells treated with DMSO, 10 nM TCIP1, or 10 nM **TCIP3** for 24, 48, or 72 h; mean ± s.e.m; 4 biological replicates; *P*-values computed by Fisher’s LSD test after analysis of variance (ANOVA); *:. *P* < 0.05; ns: not significant, *P* > 0.05; only the comparisons to DMSO at each timepoint were computed. **(H)** Quantification of percentage of fixed SUDHL5 cells in G1, S Phase, or G2/M phase after 24 h treatment with indicated compounds; 3 biological replicates, mean ± s.e.m; *P*-values adjusted by Tukey’s test after analysis of variance (ANOVA) on cells in either S, G0/G1, or G2/M phase. For G2/M cells, there were no significant differences (adj. *P* ≥ 0.05); for S and G0/G1 phase cells, **TCIP3** was significantly different from all other treatments (adj. *P* < 0.05). No other comparisons were significant.

Global proteomic changes induced by 10 nM **TCIP3** correlated (Pearson’s *R* = 0.77, *P* < 0.0001) with those observed upon 250 nM dCBP-1 treatment (Fig. 4A). This was striking not only because we treated cells with 25-times less **TCIP3** than dCBP-1, but also because p300/CBP protein levels increased in abundance due to **TCIP3** (Fig. 4A). Both compounds induced significant decreases (adj. *P* ≤ 0.05, log_2_(fold change) < -2) of germinal center B-cell-specific TFs including MEF2B, IRF8, SPIB (PU.1-related)^3,41,67^, and modestly of BCL6 itself (Fig. 4A, Supplemental Fig. 5C). This is consistent with the decreases in transcription observed at the genes encoding these TFs and the decreases in acetylation observed at their super-enhancers (Supplemental Fig. 4H and Fig. 3F). Many proteins which decreased upon p300/CBP degradation by dCBP-1 increased in abundance after **TCIP3** treatment. These included key BCL6-regulated and/or p53-target genes that play critical roles in apoptosis such as BBC3, ARID3A, and ARID3B (Fig. 4A and Supplemental Fig. 5A, B). Significantly increased proteins (adj. *P* <0.05, log_2_(fold change) > 1.0) were enriched with high confidence (adj. *P* < 0.05) for p53-target and apoptosis proteins by 10 nM **TCIP3** but not 250 nM dCBP1 treatment (Fig. 4B and Supplemental Fig. 5D). Proteomic changes induced by **TCIP3** also correlated with transcriptomic effects (Pearson’s *R* = 0.31, *P* < 0.0001); two examples of transcripts and proteins that increased are the BCL6-targets ARID3B and p27/*CDKN1B* (Fig. 4C, Fig. 3G, and Supplemental Fig. 5C). Our comparative proteomic analysis suggests that **TCIP3** is not only more potent at decommissioning p300/CBP-regulated signaling but also uniquely capable of activating apoptotic protein signaling.

### Activation of PUMA and induction of apoptosis

Global proteomics experiments revealed that PUMA/*BBC3* increased after 24 hours of treatment with **TCIP3** but not dCBP-1 (log_2_(**TCIP3** / DMSO fold change) = 0.72; adj. *P* = 0.0001) (Fig. 4A). PUMA expression continued to increase more than 6-fold after **TCIP3**-treatment for 48 hours (Fig. 4D, E). PUMA is a Bcl-2 homology 3 (BH3)-only containing pro-apoptotic protein sufficient for apoptosis caused by p53 and other stimuli^68–70^. In lymphocytes, PUMA is a target of p53, BCL6, and the transcription factor forkhead box O3 (FOXO3), the latter itself a BCL6- and p53-target gene^71^. p53 and FOXO3 protein levels did not change significantly by 48 hours (Supplemental Fig. 6A). Significant increases in PUMA levels prompted us to quantify the characteristics and kinetics of apoptotic induction by **TCIP3** by analyzing each stage of the apoptotic signaling cascade. Staining with Annexin V for the externalization of phosphatidylserine, one of the first events in a cell undergoing apoptosis, after 24, 48, and 72 hours of 10 nM **TCIP3** treatment showed that significant increases in Annexin V-positive cells began after 48 hours of **TCIP3** treatment (Fig. 4F), concurrent with the dramatic increases in PUMA at this timepoint (Fig. 4D, E). 82% of cells stained positive for Annexin V at 72 hours (Fig. 4F). This was 2.9 times higher than DMSO, 2.3 times higher than **NEG1**, and 2.5 times higher than **NEG2** treatment (Fig. 4F, Supplemental Fig. 6B), indicating that apoptosis depended on chemically induced proximity. Levels of cleaved caspase-3, the terminal executioner protease in the apoptotic cascade, increased with parallel kinetics to Annexin V (Supplemental Fig. 6C). DNA fragmentation, as measured by terminal deoxynucleotidyl transferase BrdU-dUTP nick end labeling (TUNEL) (Fig. 4G), and loss of membrane integrity, as measured by Trypan blue staining, both began after 48 hours of compound treatment (Supplemental Fig. 6D). While dCBP-1 also induced apoptosis after 72 hours of treatment (Fig. 4F), PUMA was not activated (Fig. 4D,E).

To determine the period of **TCIP3** treatment that is sufficient to trigger apoptosis, we treated cells with 10 nM **TCIP3** for either 24, 48, or 72 hours, washed cells with phosphate-buffered saline, and replaced cells with compound-free media (Supplemental Fig. 6E). Treatment for the first 24 or 48 hours of a 72-hour experiment was sufficient to induce apoptosis at levels comparable to a 72-hour continuous treatment. The kinetics of apoptosis observed suggest that 24 hours of **TCIP3** exposure is sufficient to activate one or more irreversible apoptotic signals, including PUMA, that manifests as apoptosis over the next 24-48 hours.

### TCIP3 induces G1 arrest

In addition to elevated apoptotic proteins, transcriptomic and proteomic profiling suggested that **TCIP3** may induce cell cycle arrest. We observed elevated levels of cell-cycle inhibitors such as *CDKN1B*/p27 accompanied by decreased levels of key master germinal center B cell regulators (Fig. 4A), which regulate cell proliferation^72–74^. Accordingly, we analyzed cell cycle progression after treating SUDHL5 cells with 10 nM of **TCIP3**, **NEG1**, **NEG2**, or DMSO for 24 hours. For this, we used fluorescent labels to detect total and newly synthesized DNA during a 2-hour pulse of 5-ethynyl-2’-deoxyuridine (EdU). We observed substantial and significant (*P* < 0.01) enrichment in cells arrested in G0/G1 after treatment with 10 nM **TCIP3** relative to DMSO and a corresponding significant (*P* < 0.01) reduction in S phase cells (Fig. 4H, Supplemental Fig. 6F). Cell cycle arrest after 24 hours was dependent on the ternary complex; none of the negative controls significantly changed the ratio of cells found in any of the cell cycle phases. Arrest in G0/G1 is consistent with decreased abundance of G2/M and mitotic spindle proteins detected by proteomics at 24 hours (Supplemental Fig. 5D). The measurements of apoptosis and cell cycle analysis indicate that **TCIP3** induces concurrent arrest of cell cycle progression and commitment to apoptosis in the first 24 hours of treatment, although apoptotic death does not occur until an additional 24-48 hours.

### Comparative pharmacology with BCL6-targeting TCIPs

The kinetics of apoptosis induction and cell cycle arrest differed strikingly from previous studies on two other BCL6-targeting TCIPs, TCIP1^39^ and CDK-TCIP1^40^, which recruit elongation factors associated with RNA Polymerase II to BCL6. To assess how the recruitment of chromatin modifiers to BCL6 differentially perturbs anti-proliferative mechanisms, we directly compared the effects of TCIP1 and CDK-TCIP1 with **TCIP3** on cell death and transcriptional reprogramming. **TCIP3** treatment exhibited slower kinetics of apoptosis induction than TCIP1 (Fig. 4F, G), which recruits the bromodomain and extraterminal domain (BET)-containing family member BRD4 to pro-apoptotic BH3-only genes such as BIM/*BCL2L11* and *PMAIP1*^39^. Concurrent analysis of the cell cycle and apoptotic effects of TCIP1 and **TCIP3** by TUNEL and total DNA content co-staining showed that while the predominant cellular effect after 24 hours of **TCIP3** treatment is to arrest cells in G1, 24 hours of TCIP1 treatment produces DNA fragmentation (Fig. 4G and Supplemental Fig. 7A). Unbiased clustering of transcriptomic changes mediated by **TCIP3,** TCIP1, and CDK-TCIP1, demonstrated that **TCIP3** produces a substantially different gene expression program that clusters independently from TCIP1 and CDK-TCIP1 (Supplemental Fig. 7B). Only some genes, including a set of cell-cycle-mediators, changed similarly for all three classes of TCIPs. These included *c-MYC* and the BCL6-target *CDKN1B* (p27) (Supplemental Fig. 7B). Only **TCIP3** induced dramatic changes in chromatin acetylation of enhancers and super-enhancers (Supplemental Fig. 7C). These epigenomic effects likely account for the similar proteomic changes observed between **TCIP3** and the p300/CBP degrader dCBP-1. Different molecular mechanisms correlated with a slightly different spectrum of cancer cell lines that were sensitive to each TCIP (Supplemental Fig. 7D). Nevertheless, all TCIPs were highly toxic to BCL6-driven lymphomas. Other sensitivities remain to be explored.

### Rapid and potent *c-MYC* repression

*c-MYC* transcripts were rapidly and significantly depleted upon 10 nM **TCIP3** treatment (mRNA at 2 hours of 1 nM **TCIP3**, adj. *P* ≤ 10^-100^, log_2_(fold change) = -2.71) (Fig. 5A), and global proteomics showed a complete loss of c-MYC peptides after 24 hours (Fig. 5B). c-MYC is a critical oncogenic driver expressed in all germinal center-derived lymphomas that regulates cell proliferation required for formation and maintenance of germinal centers^75,76,77^, prompting us to examine the contribution of its loss to the antiproliferative mechanism. *c-MYC* mRNA in SUDHL5 cells treated with 1 nM of **TCIP3** decreased with a t_1/2_ of 33 minutes, and repression persisted over 24 hours (Fig. 5C). c-MYC protein decreased following the loss of *c-MYC* mRNA (t_1/2_ = 2.5 hours, Supplemental Fig. 8A), indicating that **TCIP3** interferes with transcription of the *c-MYC* gene. Protein repression at 4 hours (IC_50_ ∼ 0.8 nM, Fig. 5D, E) led to significant (adj. *P* < 10^-10^) loss of c-MYC-target gene expression at the same timepoint and dose (Supplemental Fig. 8B). Thus, **TCIP3** represses transcription of the *c-MYC* gene and the entire MYC-coordinated gene expression network in DLBCL cells.

**Figure 5.**
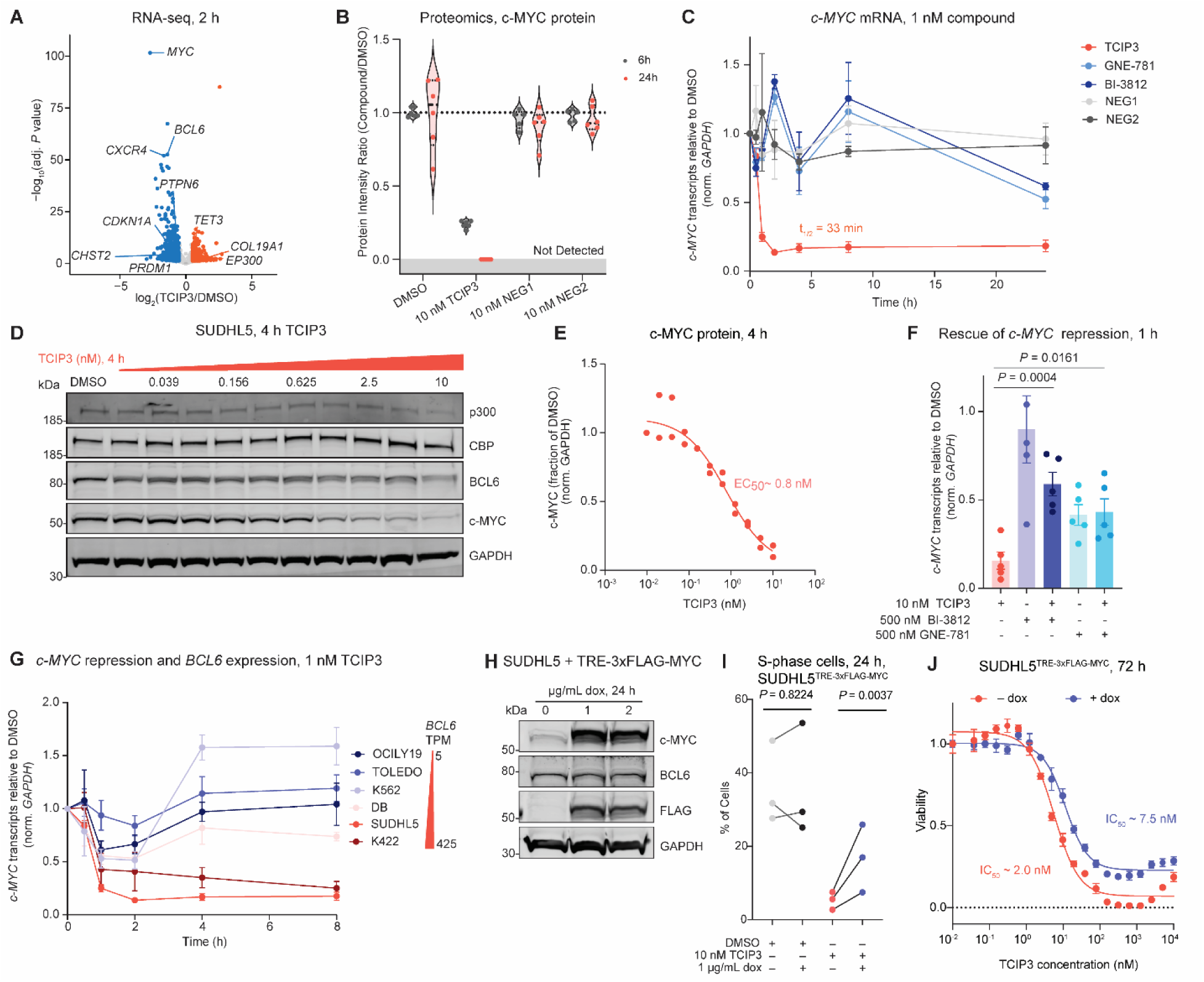
Rapid and Potent Reduction in c-MYC is Sufficient for Cell Cycle Arrest. **(A)** RNA-seq of SUDHL5 cells treated with 1 nM **TCIP3** for 2 h; colored are differential genes defined by adj. *P* ≤ 0.05 and |log_2_(fold change)| ≥ 0.5, *P*-values computed by two-sided Wald test and adjusted for multiple comparisons by Benjamini-Hochberg; 3 biological replicates. **(B)** All peptides of c-MYC detected by global proteomics after 10 nM **TCIP3**, **NEG1**, and **NEG2** treatment of SUDHL5 cells treated for 24 h; intensities are mean of 3 biological replicates. Lines represent median and interquartile range. **(C)** Time-course of *c-MYC* transcripts in SUDHL5 cells treated with 1 nM of compounds normalized to *GAPDH* and DMSO treatment as quantified through reverse transcription quantitative PCR (RT-qPCR); 3 biological replicates, mean ± s.e.m. **(D)** Western blot of c-MYC protein in SUDHL5 cells treated with **TCIP3** for 4 h; blot representative of 2 biological replicates. **(E)** Quantification of c-MYC protein normalized to GAPDH and DMSO levels from **(C)**. **(F)** Measurement of *c-MYC* mRNA in SUDHL5 cells (normalized to *GAPDH* and DMSO treatment) by RT-qPCR after competitive titration of constant 10 nM **TCIP3** with 500 nM of the BCL6^BTB^ domain inhibitor BI-3812^48^ or 500 nM of the p300/CBP bromodomain inhibitor GNE-781^50^; cells were treated simultaneously with **TCIP3** and the inhibitor or DMSO for 1 h; effects of co-treatment of DMSO and inhibitors shown for comparison; 3 biological replicates, mean ± s.e.m.; *P*-values adjusted by Tukey’s test after analysis of variance (ANOVA). Only comparisons of co-treatments to **TCIP3** were computed. **(G)** c*-MYC* transcripts in DLBCL and leukemia cells with varying *BCL6* expression^95^ (TPM: transcripts/million) treated with 1 nM of **TCIP3** normalized to *GAPDH* and DMSO treatment as quantified through RT-qPCR; 3 biological replicates, mean ± s.e.m. SUDHL5 data is from panel **B**. **(H)** Doxycycline-inducible overexpression of c-MYC in SUDHL5 cells. **(I)** S phase cells of SUDHL5^TRE-3xFLAG-MYC^ cells treated with or without 1 µg/mL doxycycline dissolved in ethanol or vehicle 24 h prior to 24 h treatment with 10 nM **TCIP3** or DMSO; 3 biological replicates, *P-*values computed by two-tailed ratio paired Students’ t-test. **(J)** Viability of SUDHL5^TRE-3xFLAG-MYC^ cells treated with 1 µg/mL doxycycline dissolved in ethanol or vehicle 24 h prior to 72 h treatment with **TCIP3**; mean ± s.e.m., 3 biological replicates.

To investigate the mechanism of *c-MYC* repression, we examined whether repression of *c-MYC* transcripts depended on induced proximity of p300/CBP and BCL6 by co-treating SUDHL5 cells with 10 nM **TCIP3** and either 500 nM of BI-3812 or GNE-781 to compete for binding with BCL6 or p300/CBP, respectively. Co-treatment with BI-3812 almost completely restored *c-MYC* levels to baseline, and co-treatment with 500 nM of GNE-781 restored *c-MYC* levels to those observed with GNE-781 treatment alone (Fig. 5F). *c-MYC* repression induced with 500 nM GNE-781 was still more than two times higher than repression in cells treated with 10 nM **TCIP3**. Neither GNE-781, dCBP-1, **NEG1, NEG2,** nor the BCL6^BTB^ inhibitor BI-3812 produced comparable *c-MYC* repression at matched concentrations at the transcript or proteomic level (Fig. 5C, B, Supplemental Fig. 8C). The p300/CBP-TCIP3-BCL6 ternary complex is therefore required for **TCIP3**-induced *c-MYC* repression.

*c-MYC* repression across six lymphoma cell lines correlated with BCL6 levels (Fig. 5G). BCL6-high expressing cells SUDHL5 and K422 displayed more pronounced *c-MYC* repression following a 1 nM **TCIP3** treatment than BCL6-low lymphoma cells OCILY19 and TOLEDO, or the leukemia line K562 (Fig. 5G). DB, a germinal center B-type lymphoma line with high *BCL6* levels (269 transcripts/million) and a recently identified *c-MYC* chromosomal rearrangement with the breakpoint at its promoter^78^, exhibited less repression (Fig. 5G). Indeed, repression was muted in BCL6-high Burkitt’s lymphoma cell lines (DAUDI and RAJI) with the characteristic chromosomal rearrangement of one or both alleles of the *c-MYC* promoter to the immunoglobulin heavy chain regulatory region^79,80^ (Supplemental Fig. 8D). These results imply that **TCIP3** may indirectly disrupt cis-regulation of *c-MYC* transcription. Consistent with this possibility, modest losses of histone acetylation at the *c-MYC* promoter and its known regulatory enhancers^78,81^ were observed only after *c-MYC* repression, likely as a result of lost transcription (Supplemental Fig. 8E, F). This stands in contrast to p300/CBP bromodomain inhibitors, which repress *c-MYC* transcription by displacing p300/CBP binding and activity from *c-MYC* enhancers and promoters^30,50^. Therefore, **TCIP3**-mediated repression of *c-MYC* is a mechanistically distinct consequence of chemically induced proximity of p300/CBP and BCL6.

### Restoration of *c-MYC* reverses cell cycle arrest but not death

To evaluate the importance of c-MYC repression in driving the cellular response to **TCIP3**, we overexpressed doxycycline(dox)-inducible 3x-FLAG-MYC in SUDHL5 cells (TRE-3x-FLAG-MYC) (Fig. 5H). Analysis of the cell cycle at 24 hours indicated that overexpression of c-MYC partially reversed the S-phase block caused by 10 nM **TCIP3** (Fig. 5I and Supplemental Fig. 8G). This correlated with a ∼3.8-fold proliferative benefit in cells expressing dox-induced MYC and treated with **TCIP3** (IC_50_ ∼ 7.5 nM), relative to **TCIP3**-treated cells without dox (IC_50_ ∼ 2.0 nM, Fig. 5J). These data suggest that c-MYC repression contributes to the G1 arrest but cannot reverse the cell killing effects of **TCIP3**, which likely involves direct transcriptional activation of pro-apoptotic factors (Figs. 3-4).

### TCIP3 is a molecular glue

The selectivity of **TCIP3** for its binding partners (Fig. 2C, D) and its use of chemical induced proximity to activate cell death led us to ask what features of the three-component assembly containing BCL6-**TCIP3**-p300 drive extraordinary potency and selectivity. To understand the determinants of specific ternary complex formation, we solved the crystal structure of a BCL6-**MNN-02-155**-p300 ternary complex. This KAT-TCIP features a 4-carbon alkyl linker and induces potent activation of the BCL6-target reporter gene and cell death with an IC_50_ comparable to **TCIP3** (Supplemental Fig. 1A, C). The asymmetric unit, resolved to a minimum Bragg spacing of 2.1 Å (Supplemental Table 3), contained a BCL6^BTB^ homodimer and two copies of a p300^BD^ molecule, each tethered to BCL6 by **MNN-02-155** (Fig. 6A, 6B, Supplemental Fig. 9A, B, C, Supplemental Video 1). **MNN-02-155** engages p300 and BCL6 without significant rearrangements of the parental monovalent protein-compound structures: GNE-781-CBP^50^ (RMSD = 0.35 Å, Supplemental Figure 9D) and BI-3802-BCL6^48^ (RMSD = 0.43 Å, Supplemental Figure 9E). A hydrogen bond connects the carbonyl of p300^G1085^ and the amide nitrogen in the **MNN-02-155** linker (Fig. 6C). Five residues engaged in fortuitous interactions constitute a p300-BCL6 neo-interface: p300^L1082^ contacts BCL6^S27^, while p300^Q1083^ contacts BCL6^R24^ and BCL6^R20^. The interactions are primarily hydrophobic in character (Fig. 6D).

**Figure 6.**
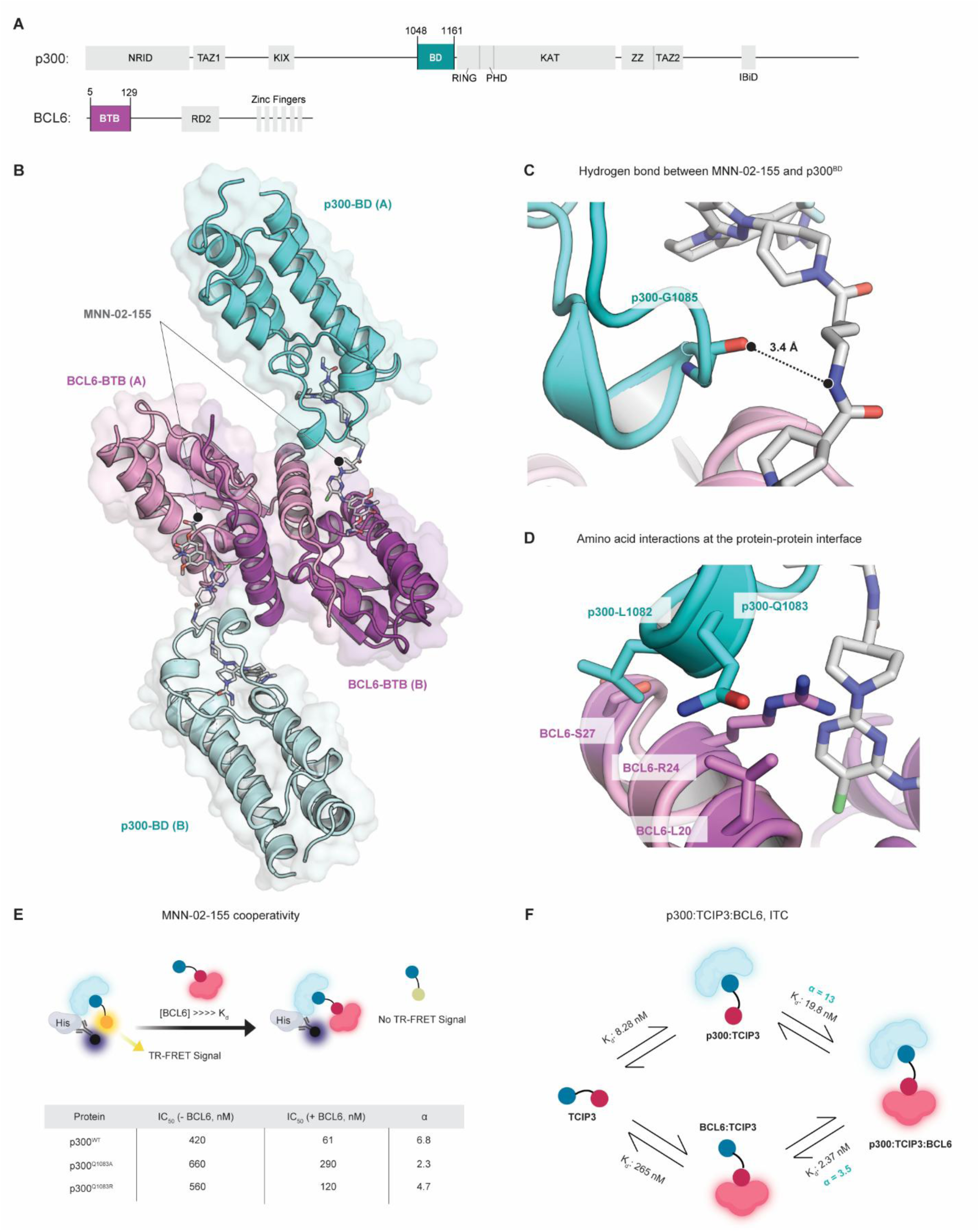
The Crystal Structure and Cooperativity of the Ternary Complex. **(A)** Domain structure of full-length p300/CBP and BCL6. The highlighted p300 bromodomain (BD) and BCL6 BTB domain (BTB) were crystallized in complex with **MNN-02-155**. **(B)** Co-crystal structure of the ternary complex formed by **MNN-02-155**, BCL6^BTB^, and p300^BD^. One dimer of BCL6 is bound to two molecules of MNN-02-155, each of which engages one protomer of p300^BD^. **(C)** Neo-hydrogen bond formed between **MNN-02-155** and the backbone of P300G^1085^. **(D)** Neo-protein-protein interactions formed at the interface of p300^BD^ and BCL6^BTB^ mediated by **MNN-02-155** binding. **(E)** Binary TR-FRET displacement assay assessing the binding of 6x-His-P300^BD^ to **MNN-02-155**. **MNN-02-155** was titrated into a biochemical complex of terbium-labeled 6x-His-p300^BD^ pre-incubated with **MNN-06-112**. Parallel experiments were performed by preincubating with BCL6 at concentrations exceeding its dissociation constant (>>> Kd). Binding affinities of p300^WT^, p300^Q1083A^, and p300^Q1083R^ to **MNN-02-155** were evaluated in the absence and presence of BCL6 and a cooperativity constant was calculated (α = K_d_ binary/K_d_ ternary). n = 3 independent experiments, mean. **(F)** K_d_ calculations for each binding event, based on n = 2-3 independent experiments, mean ± s.e.m. α was calculated for both p300 and BCL6 (α = K_d_ binary/K_d_ ternary).

The identification of chemically induced amino acid contacts between p300 and BCL6 suggested cooperativity in ternary complex formation^82^, a hallmark of molecular glues^23,26^. Cooperativity (α) is defined as the ratio of binary to ternary complex dissociation constants (K_d_s). We followed a reported procedure^83^ to measure α for p300, defined as the ratio between the K_d_ for p300-**MNN-02-155** binding versus the K_d_ for p300-[BCL6-**MNN-02-155**] binding. For p300^WT^, this value was α ∼ 6.8 (Fig. 6E), indicating strong ternary complex cooperativity. To understand the importance of the aforementioned contacts for KAT-TCIP binding and cooperativity, we measured α for p300Q^1083A^ or p300^Q1083R^ to reduce residue bulk or to introduce positive charge, respectively. Given the crystal structure, the latter mutation would clash with the interacting BCL6^R24^ amino acid side chain (Fig. 6D). p300^WT^ bound more favorably to **MNN-02-155** in the presence of BCL6 relative to p300^Q1083A^ and p300^Q1083R^ (Fig. 6E, Supplemental Fig. 10A). The presence of BCL6 did not affect the affinity of GNE-781 to the various p300 mutants (p300Q^1083A^ α ∼ 0.88; p300^Q1083R^ α ∼ 1.0; Supplemental Fig. 10B). Thus, fortuitous amino acid side chain contact between p300^Q1083^ and BCL6^BTB^ enhances complex formation, providing a biophysical explanation for potency and specificity.

We next used the cooperativity assay to profile other active GNE-781-derived KAT-TCIPs. **TCIP3** and **MNN-02-155** produced similar α values (α = 4.2 and α = 6.9, respectively; Supplemental Fig. 10C). We measured weaker cooperativity for compounds that were less cytotoxic such as **MNN-02-156** and **MNN-02-162** (Supplemental Fig. 10C). Correlation between α and cellular potency prompted us to investigate the average three-dimensional linker lengths of each GNE-781-based KAT-TCIP, as measured from predicted 3D conformations (Supplemental Fig. 10D). The computationally predicted linker length (Methods) of **MNN-02-155** (6.6 Å) deviated by less than 1 Å from the measured distance between p300^BD^ and BCL6^BTB^ in the co-crystal structure (6.1 Å), validating this approach (Supplemental Fig. 10D). Compounds with linker lengths from 4 to 12 Å demonstrated antiproliferative activity at IC_50_s < 10 nM, and compounds with rigidified linkers were consistently very potent (IC_50_ < 1 nM) (Supplemental Fig. 10D). Rigid linkers decrease rotational degrees of freedom and lower the entropic cost of ternary complex formation. Consequently, it is likely that they produce longer-lasting active ternary complexes inside the cell, assuming that membrane permeability is similar. Given the tolerability for a wide range of linker chemistry and the limited plasticity observed at the protein-protein interface in the co-crystal structure, we conclude that many different orientations of p300/CBP relative to BCL6 can support biological activity.

Finally, we used isothermal calorimetry (ITC) to quantify the binary and ternary complex binding affinities and thermodynamic parameters of p300^BD^, BCL6^BTB^, and **TCIP3** (Supplemental Fig. 10E). Calculations of α in both directions of ternary complex formation demonstrated that **TCIP3** exhibits significantly enhanced affinity for the ternary complex compared to binding with either p300^BD^ or BCL6^BTB^ alone (α ^p300 into BCL6:TCIP3^ = 3.5, α^BCL6 into p300: TCIP3^ =13, Fig. 6F). These values are comparable to those obtained with TR-FRET (Fig. 6E). Consistent with the expectation that the △G of ternary complex formation should be the same regardless of whether binding to p300 or BCL6 occurs first, there was only a 4% deviation in the overall free energy of complex formation between the p300 and BCL6 mediated binding mechanisms (Supplemental Figure 10F). Our biophysical and structural data confirms that **TCIP3**, by inducing the formation of a cooperative ternary complex containing new, stabilizing interactions between p300/CBP and BCL6, acts as a molecular glue.

To investigate how the relative orientation of p300^BD^ to BCL6^BTB^ may relate to biological activity, we superimposed the fragment of p300 seen in the p300-**MNN-02-155**-BCL6^BTB^ structure onto a crystal structure of the p300 catalytic core (PDB: 6GYR) containing the catalytic KAT domain^84,85^, the p300/CBP <u>a</u>uto-inhibitory loop (AIL)^86^, and the really interesting <u>n</u>ew <u>g</u>ene (RING) domain which regulates catalytic KAT activity^84^. **MNN-02-155** positions active p300 protomers in an orientation compatible with acetylation of BCL6 and proximal chromatin (Supplemental Fig. 10G, teal). Moreover, because BCL6 is an obligate homodimer ^87,88^, 2 molecules of p300 should be recruited to BCL6 binding sites over the genome. Superimposition revealed an opportunity for a second p300 protomer (Supplemental Fig. 10G, gray) to interact with each BCL6-bound p300 molecule. Activation by auto-acetylation occurs in trans across a p300 homodimer assembled on a dimeric binding partner^85^, indicating the potential for higher-ordered oligomers to form on chromatin, facilitating KAT trans-acetylation and reinforcing local chromatin acetylation. We propose a model in which **TCIP3** leverages cooperativity to form a stable and enzymatically active p300-BCL6 complex that goes on to initiate gene modulation on chromatin and cell death.

## DISCUSSION

The introduction of KAT-TCIPs expands the toolbox by which we can precisely reprogram endogenous gene expression networks. KATs are prominent therapeutic targets in lymphoma, but available small molecules inhibit their function globally and have the potential to cause multiple on-target toxicities. We alternatively converted a potent p300 inhibitor into a bivalent small molecule that leverages chemical induced proximity (CIP) to redirect p300/CBP acetyltransferase activity, triggering cell death in diffuse large B-cell lymphomas. The resulting KAT-TCIPs are small molecules that activate transcription of genes bound by the master transcriptional repressor BCL6, whose dysregulation drives DLBCL progression. The lead KAT-CIP, which we named **TCIP3**, activated BCL6-target gene expression and inhibited proliferation in DLBCL cells at sub-nanomolar IC_50_s without exhibiting toxicity in non-transformed tonsillar lymphocytes or fibroblasts.

**TCIP3** potency was contingent on the formation of a ternary complex between **TCIP3**, p300/CBP, and BCL6, which served as a driver of exquisite selectivity (Figs. 2 and 6). Small-molecule-induced cooperative ternary complexes confer beneficial pharmacological properties such as prolonged target residence time^23,26^, thereby enhancing the likelihood of sustained and potent therapeutic outcomes. Here, we used structural and biophysical analyses to demonstrate that **TCIP3** is a bivalent molecular glue that shares these advantageous characteristics. The crystal structure of a KAT-TCIP in complex with p300 and BCL6 exhibited potency enhancement by fortuitous amino acid contacts independent of plasticity at the protein-protein interface. This suggests that AI-driven docking strategies leveraging existing structures could accelerate the optimization of CIP-enabling compounds for therapeutic purposes.

KAT-TCIPs killed DLBCL cells by redirecting p300/CBP activity in a mechanism that depended on the oncogenic driver BCL6. The consequences of this on chromatin, transcription, and protein expression in the cell unexpectedly resembled changes induced by p300/CBP degradation. Three observations distinguish KAT-TCIPs from simple inhibitors or degraders. First, while the p300/CBP degrader dCBP-1 depletes both histone acetylation genome-wide and p300/CBP protein levels^30^, **TCIP3** selectively targeted DLBCL-specific super-enhancers and stabilized p300/CBP, indicating that it is not simply a super-inhibitor. Second, **TCIP3** activated apoptotic protein signaling. Elevated PUMA, a BH3-only protein sufficient for intrinsic apoptosis^68–70^, exemplifies this characteristic. Third, acute treatment produced rapid and durable repression of c-MYC in a manner dependent on chemically induced proximity to BCL6. All three molecular effects combined to elicit more potent cell killing than p300/CBP PROTACs, KAT inhibitors, or bromodomain inhibitors and BTB domain inhibitors of BCL6 (Fig. 1). The fact that c-MYC replacement could reverse cell cycle arrest but not death (Fig. 5) strongly suggests that anti-cancer activity is the product of multiple gain-of-function effects. Our results demonstrate the potential of redirecting epigenetic regulators via CIP to produce cancer cell killing.

The redistribution mechanism suggests that cells particularly vulnerable to disruption of p300/CBP that also have high BCL6 activity may be sensitive to **TCIP3**. In addition to DLBCL, follicular lymphoma, an incurable subtype of non-Hodgkin’s lymphoma, represents a potentially responsive disease. Patients exhibit both a high frequency of inactivating *CREBBP* mutations (65%) that lead to a loss of BCL6 antagonism and unchecked BCL6 activity^58,89,90,91^. Moreover, while BCL6-expressing lymphomas provide a compelling model to explore the therapeutic potential of the BCL6-targeting KAT-TCIPs described here, the widespread expression of p300/CBP across various cancers suggests the broader applicability of CIPs that redirect p300/CBP to pro-apoptotic TFs.

The combinatorial nature of transcription led us to hypothesize that recruiting acetyltransferases to BCL6 would reprogram gene networks distinct from those induced by the recruitment of other co-activators, despite localization to the same TF. We previously developed TCIP1 and CDK-TCIP1, molecules that redirect the positive transcription elongation factors BRD4 and CDK9, respectively, to activate BCL6-target genes^39,40^. **TCIP3** provoked substantially different gene expression changes and cell arrest/death phenotypes relative to these TCIPs (Fig. 4 and Supplemental Fig. 7). Differential activity is consistent with the complex multistep nature of transcriptional activation. While the elongation factors BRD4 and CDK9 work in concert to release promoter-proximal paused RNA Polymerase II, facilitating active transcription along the gene body^92^, p300/CBP promotes chromatin accessibility and TF binding, enabling the core transcriptional machinery to assemble^93,94^. Thus, we speculate that BRD4 and CDK9-based TCIPs may be more effective at inducing transcription of BCL6 sites where RNA Pol II is already in the vicinity of transcriptional start sites, while p300/CBP KAT-TCIPs may exhibit the most influential activity at BCL6 sites lacking RNA Pol II.

Consequently, transcriptional output and cell fate would be different. The modular pharmacology of TCIPs stands in contrast to targeted protein degraders, which result in target depletion-mediated effects regardless of the E3 ubiquitin ligase recruited. Our studies suggest TCIPs targeting the same transcription factor but recruiting different transcriptional/epigenetic modifiers could be used sequentially to overcome mechanism-based resistance.

## SUPPLEMENTAL FIGURES

**Supplemental Figure 1.**
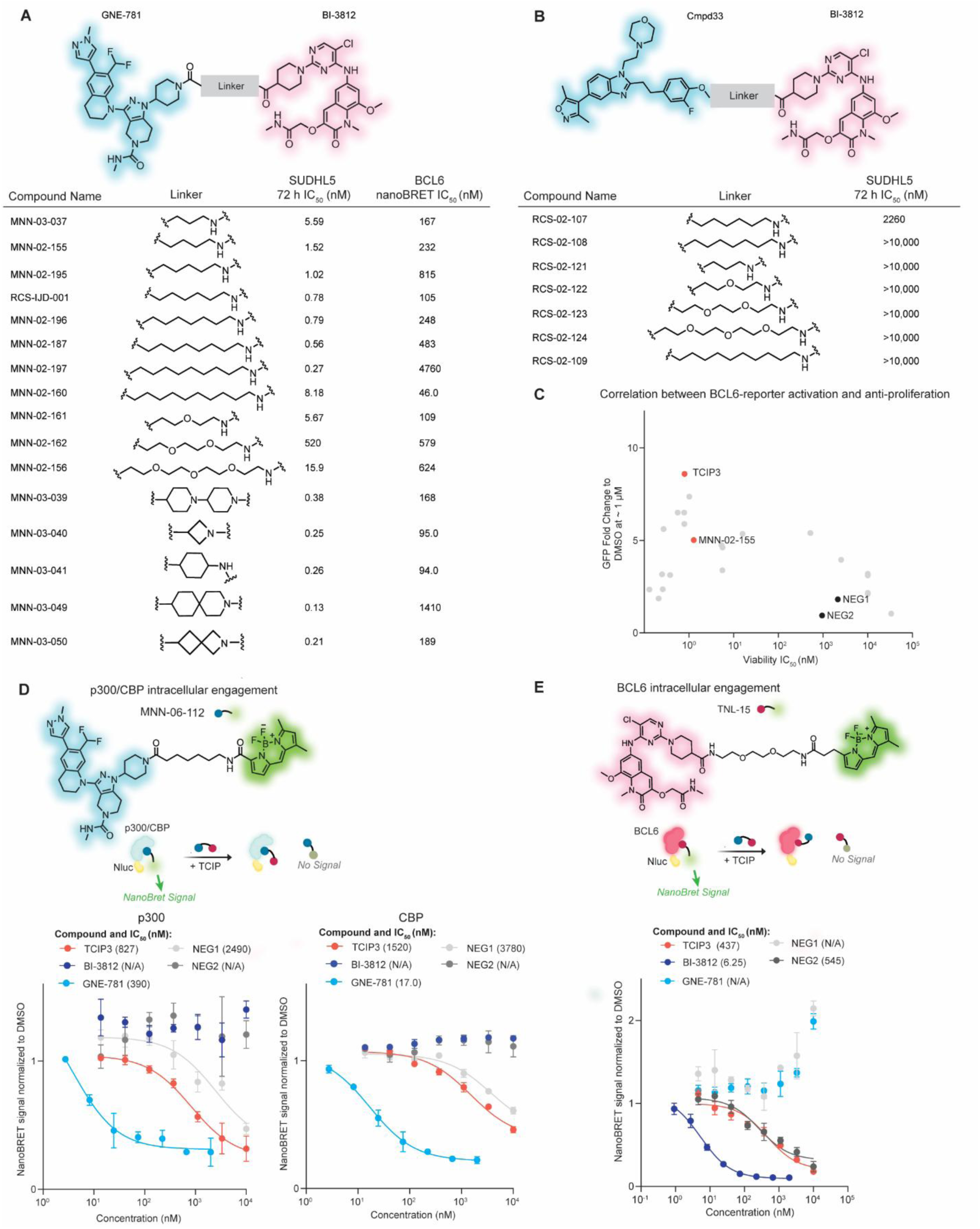
Structure-Activity Relationship of p300/CBP KAT-TCIP Library. **(A)** Design of GNE-781 and **(B)** Cmpd33-based KAT-TCIP libraries and corresponding IC_50_ values of cell viability after 72 h treatment in SUDHL5 and BCL6^BTB^ intracellular probe-displacement (nanoBRET: nano-bioluminescence resonance energy transfer) in HEK293T cells; for cell viability, mean of 1-4 biological replicates; for nanoBRET, mean of 3 technical replicates. **(B)** Reporter transactivation (fold change of BCL6-repressed GFP) after 24 h of treatment in K422 reporter cells versus IC_50_ values of cell viability after 72 h treatment in SUDHL5 for all KAT-TCIP compounds; for cell viability, mean of 1-4 biological replicates. **(C)** Assessment of compounds and corresponding IC_50_ values of p300 and CBP intracellular probe-displacement in 293T cells; the probe **MNN-06-112** is shown; Nluc: nano-luciferase. **(D)** Assessment of compounds and corresponding IC_50_ values in BCL6 nanoBRET in HEK293T cells; the probe **TNL-15** is shown. For **(D)**, **(E)**, mean ± s.e.m of 3 technical replicates.

**Supplemental Figure 2.**
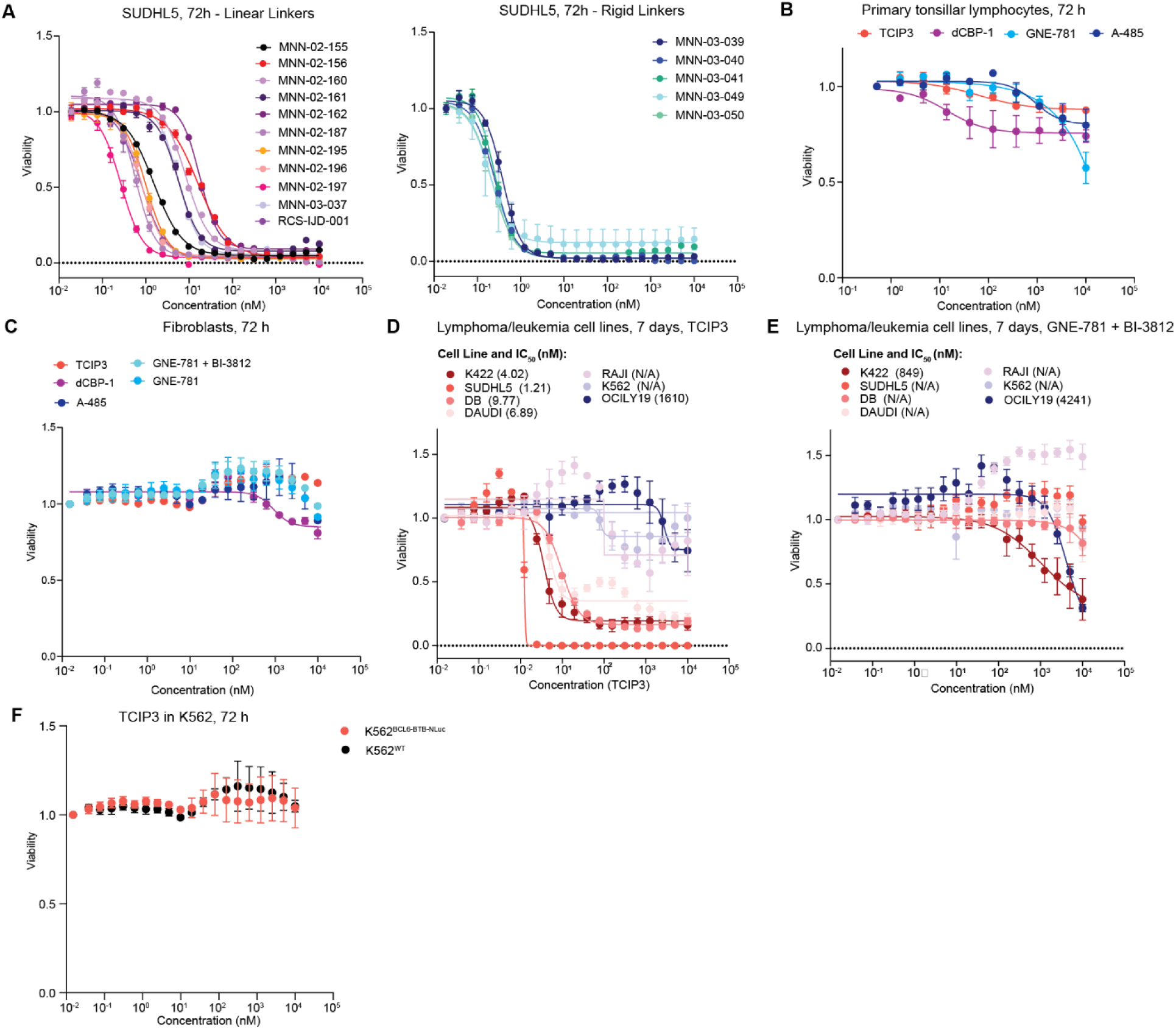
Assessment of KAT-TCIP Toxicity in Primary and DLBCL Cells. **(A)** Cell viability curves of GNE-781-based KAT-TCIPs containing either linear or rigid linkers after 72 h treatment in SUDHL5 cells; corresponds to summary data in Supplemental Fig. 1A; mean ± s.e.m. of 2-3 biological replicates or 3 technical replicates. **(B)** Viability effects of **TCIP3** and known p300/CBP targeting agents in primary tonsillar lymphocytes from two independent donors (Methods) after treatment for 72 h; mean ± s.e.m. **(C)** Viability effects of **TCIP3** and known p300/CBP targeting agents in human fibroblasts after treatment for 72 h; 2 biological replicates; mean ± s.e.m. **(D)** Viability effects of 7 days of **TCIP3** treatment in DLBCL and leukemia cell lines; 3-4 biological replicates; mean ± s.e.m. **(E)** Viability effects of 7 days of BI-3812 and GNE-781 co-treatment in DLBCL and leukemia cell lines; 3-4 biological replicates; mean ± s.e.m. **(F)** Viability effects after 72 h of **TCIP3** treatment in K562 cells overexpressing NLuc-BCL6^BTB^; 3 biological replicates; mean ± s.e.m.

**Supplemental Figure 3.**
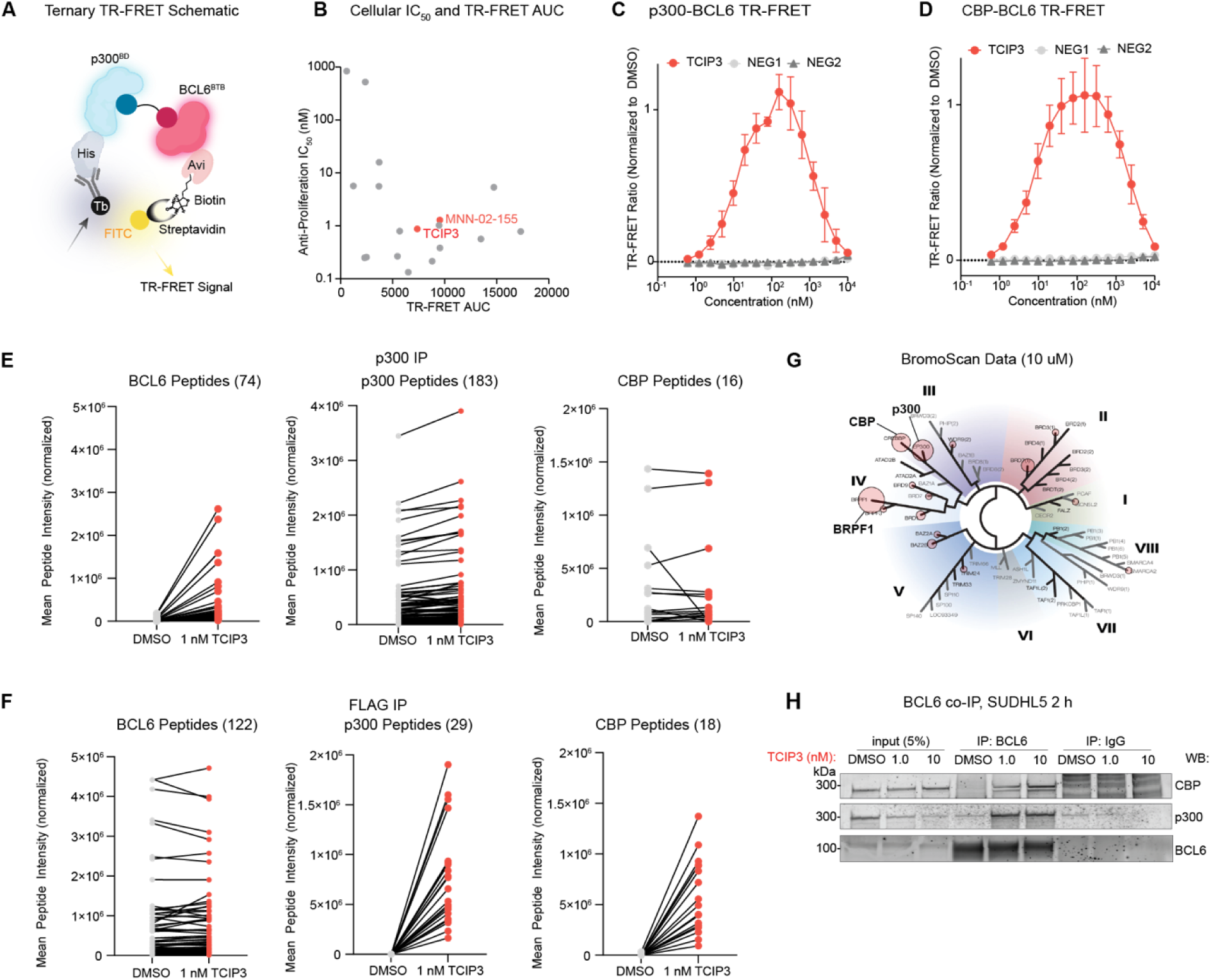
Biochemical and Cellular Engagement of BCL6 and p300/CBP. **(A)** Design of ternary TR-FRET assay using recombinant 6x-His-p300^BD^ or 6x-His-CBP^BD^ and biotinylated BCL6^BTB^-Avi to measure ternary complex formation. **(B)** Antiproliferation IC_50_ (nM) vs TR-FRET curve (AUC) for KAT-TCIPs synthesized from GNE-781; for TR-FRET, mean of 3 biological replicates each with 3 technical replicates; for cell viability IC_50_s, mean of 1-4 biological replicates. **(C)** p300-BCL6 or **(D)** CBP-BCL6 TR-FRET assay of **TCIP3, NEG1,** or **NEG2**; mean ± s.e.m. of 3 technical replicates. **(D)** Mean peptide intensities of BCL6, p300, or CBP matched between DMSO or **TCIP3** treatments in SUDHL5 cells treated for 2 h for p300 IP-MS shown in Figure 2C. **(E)** Mean peptide intensities of BCL6, p300, or CBP matched between DMSO or **TCIP3** treatments in FLAG-tagged *BCL6* SUDHL5 cells treated for 2 h for FLAG IP-MS shown in Figure 2D. For **E** and **F**, total number of unique, proteotypic peptides across all conditions provided in heading parentheses. **(G)** Selectivity of binding to 40 recombinant human bromodomains (BROMOscan) with **TCIP3** (10 µM). **(H)** BCL6 IP from SUDHL5 cells treated with the indicated doses of **TCIP3** for 2 h; representative of 3 biological replicates.

**Supplemental Figure 4.**
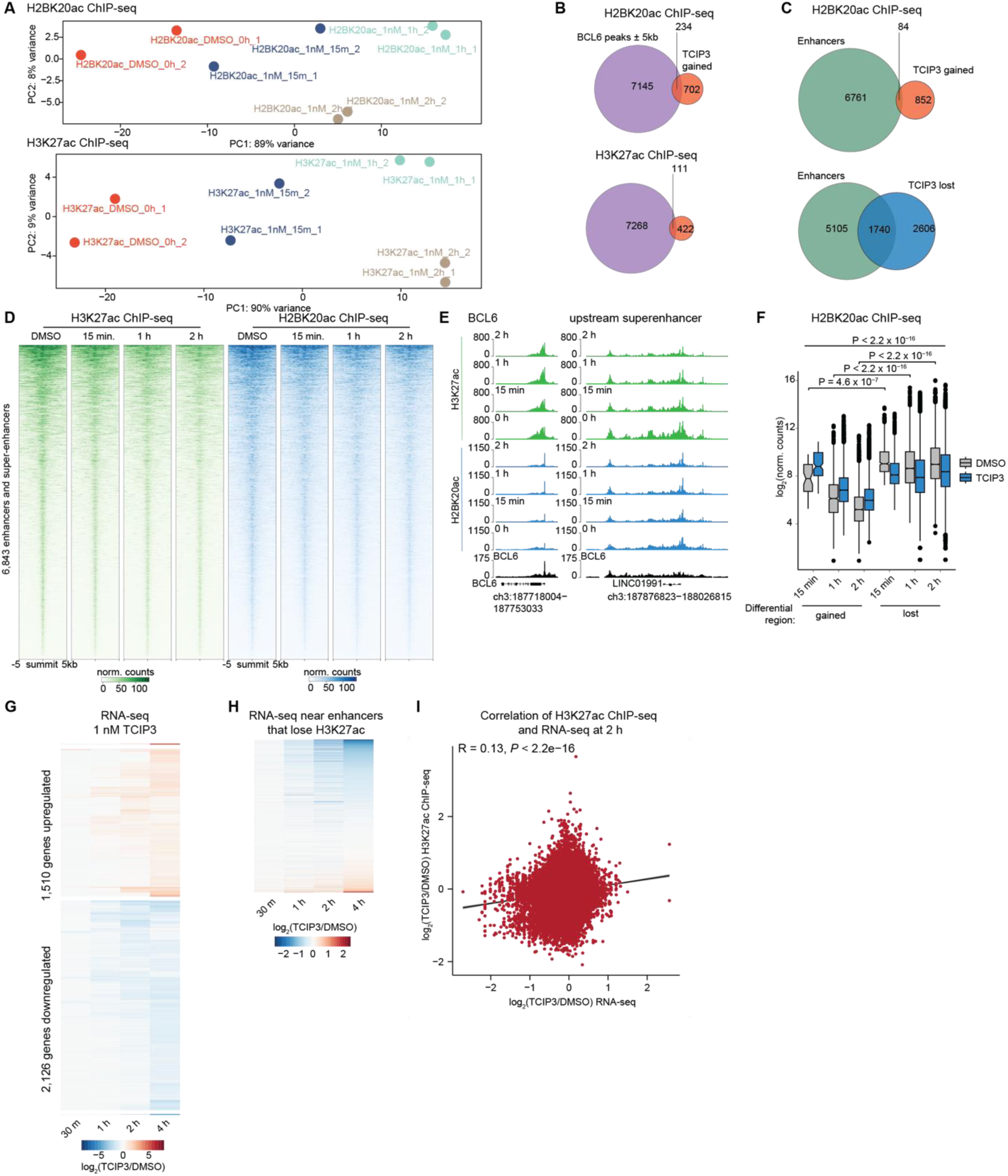
Reprogramming of Histone Lysine Acetylation and Gene Expression. **(A)** Principal component analysis of biological replicates for H3K27ac (2 per timepoint) and H2BK20ac (2 per timepoint) ChIP-seq experiments. **(B)** Overlap of gained H2BK20ac and H3K27ac peaks after 1 h of 1 nM TCIP3 with BCL6 summits ± 5 kilobases (kb) in SUDHL5 cells as measured by BCL6 CUT&RUN. **(C)** Overlap of gained and lost H2BK20ac peaks after 1 h of 1 nM **TCIP3** with annotated enhancers and super-enhancers in SUDHL5 cells.; differential regions defined as in Fig. 3A. **(D)** H3K27ac and H2BK20ac at all enhancers and super-enhancers for the indicated timepoints of **TCIP3** treatment; merged from 2 biological replicates and sequence-depth normalized and input-subtracted. **(E)** Induction of H2BK20ac and H3K27ac with time at the promoter of *BCL6* concomitant with loss at the *BCL6* upstream super-enhancer; BCL6 track is CUT&RUN in untreated SUDHL5 cells, tracks merged from two biological replicates and sequence-depth normalized and, for histone acetylation ChIP-seq, also input-subtracted. **(F)** Comparison of H2BK20ac loading at differential regions at 15 min, 1 h, and 2 h of 1 nM **TCIP3**; *P*-values adjusted by Tukey’s test after type II analysis of variance (ANOVA). **(G)** Time-dependent changes in gene expression; plotted are differential genes defined by adj. *P* ≤ 0.05 and |log_2_(fold change)| ≥ 0.5, *P*-values computed by two-sided Wald test and adjusted for multiple comparisons by Benjamini-Hochberg. **(H)** Changes in gene expression at genes near enhancers and super-enhancers that had statistically significant (adj. *P* ≤ 0.05 and log_2_(fold change) ≤ -0.5) losses in H3K27ac as measured by ChIP-seq Fig. 3E. **(I)** Correlation of changes at all H3K27ac peaks (66,995) after 2 h of 1 nM **TCIP3** treatment with changes in gene expression of nearest gene to peak; *R* computed by Pearson’s correlation and *P-*value computed by two-sided Student’s t-test.

**Supplemental Figure 5.**
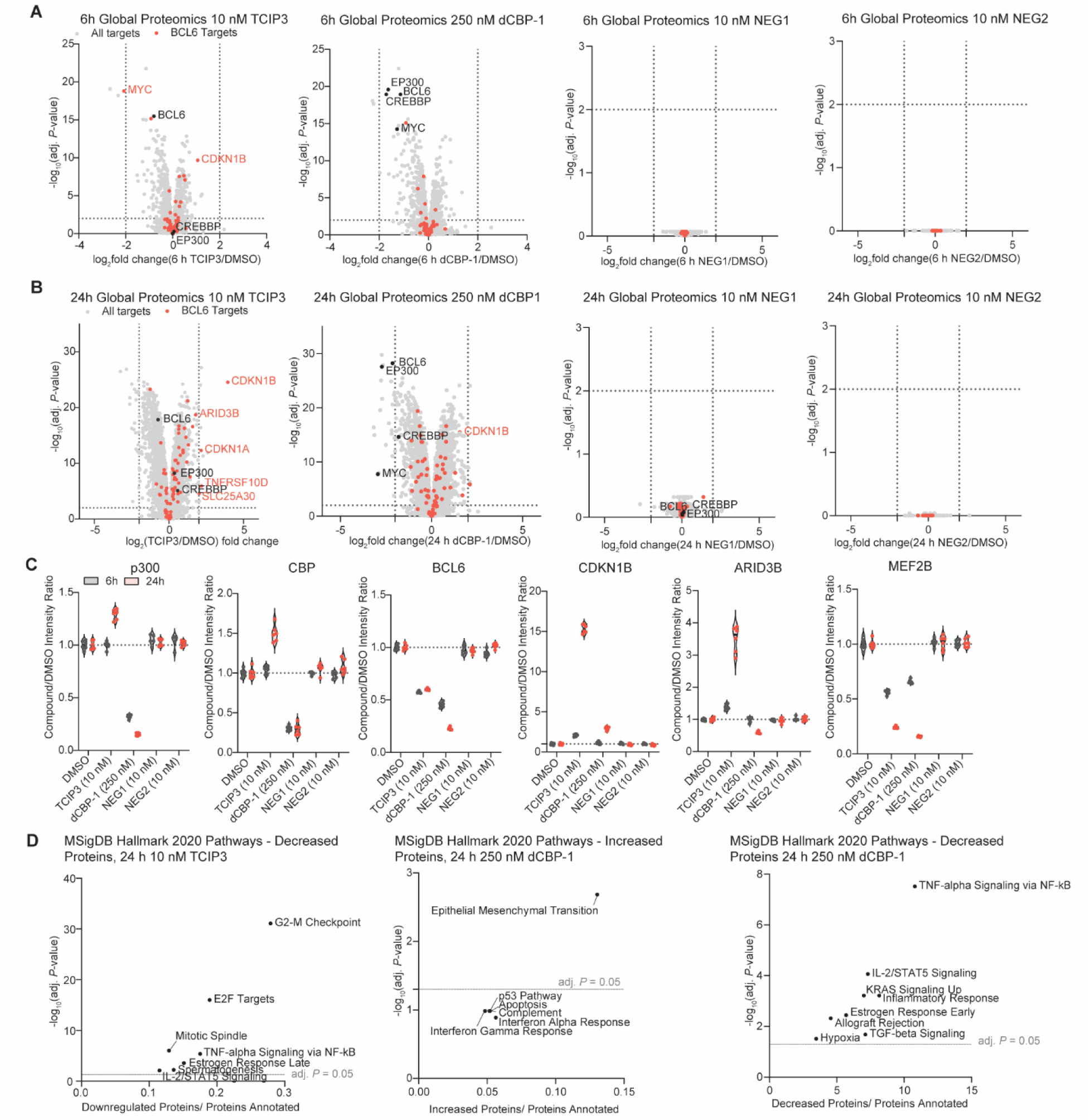
Proteomic Effects of TCIP3, dCBP1, NEG1, and NEG2. **(A)** Whole-proteome profiling of SUDHL5 cells treated with10 nM **TCIP3**, 250 nM dCBP1, 10 nM **NEG1**, or 10 nM **NEG2** for 24 h plotted with cutoffs of log_2_(fold change)| ≥ 2 and adj. *P* ≤ 0.05. **(B)** Whole-proteome profiling of SUDHL5 cells treated with 10 nM **TCIP3**, 250 nM dCBP1, 10 nM **NEG1**, or 10 nM **NEG2** for 6 h plotted with cutoffs of log_2_(fold change)| ≥ 2 and adj. *P* ≤ 0.05. For **(A)**, **(B)**: 3 biological replicates; *P*-values computed using a moderated t-test and adjusted by Benjamini-Hochberg. **(C)** Quantification of individual peptides of interest from global proteomics of all treatments and timepoints matched across treatments. Lines represent median and interquartile range. **(D)** Signaling pathways (MSigDB Hallmark 2020) enriched after 24 h treatment of indicated compounds in SUDHL5 in significantly decreased (adj. *P* < 0.05, log_2_(foldchange) < -1) or significantly increased (adj. *P* < 0.05, log_2_(foldchange) > 1) proteins.

**Supplemental Figure 6.**
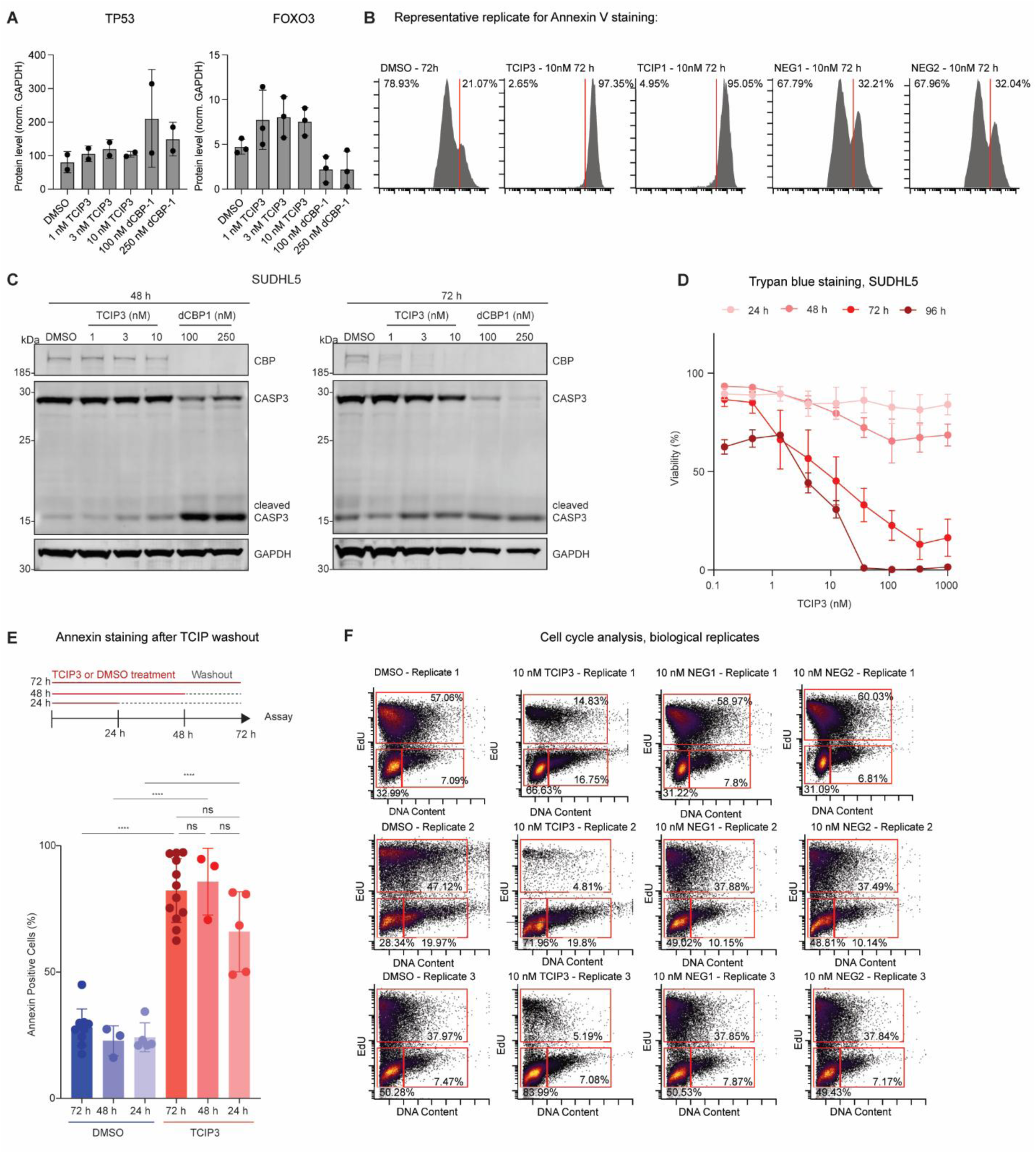
Characterization of Apoptosis and Cell Cycle Arrest. **(A)** Quantification of TP53 and FOXO3 protein levels from Fig. 4D; *P-*values computed by Fisher’s LSD test after ANOVA; only the comparisons to DMSO were computed; no comparisons were significant. **(B)** Representative (of 3-12 biological replicates) Annexin V staining replicate of SUDHL5 cells treated with **TCIP3, NEG1**, **NEG2**, or DMSO for 72 h. **(C)** Western blot of caspase 3 in SUDHL5 cells treated with indicated compounds and doses after 48 and 72 h; representative of 3 biological replicates. **(D)** Percent of cells alive after Trypan blue staining (Trypan blue negative); SUDHL5 cells treated with **TCIP3** for indicated timepoints and doses. 3 biological replicates; mean ± sem. **(E)** Quantification of Annexin V positive SUDHL5 cells treated with DMSO or 10 nM **TCIP3** for 24, 48, or 72 h and then washed with PBS and replaced with media for 48, 24, or 0 h, respectively; *P*-values adjusted by Tukey’s test after analysis of variance (ANOVA); ****: adj. *P* < 0.0001; all unlabeled comparisons were not significant. 72 h data corresponds to Fig. 4F. **(F)** Individual cell cycle analysis biological replicates corresponding to Fig. 4H.

**Supplemental Figure 7.**
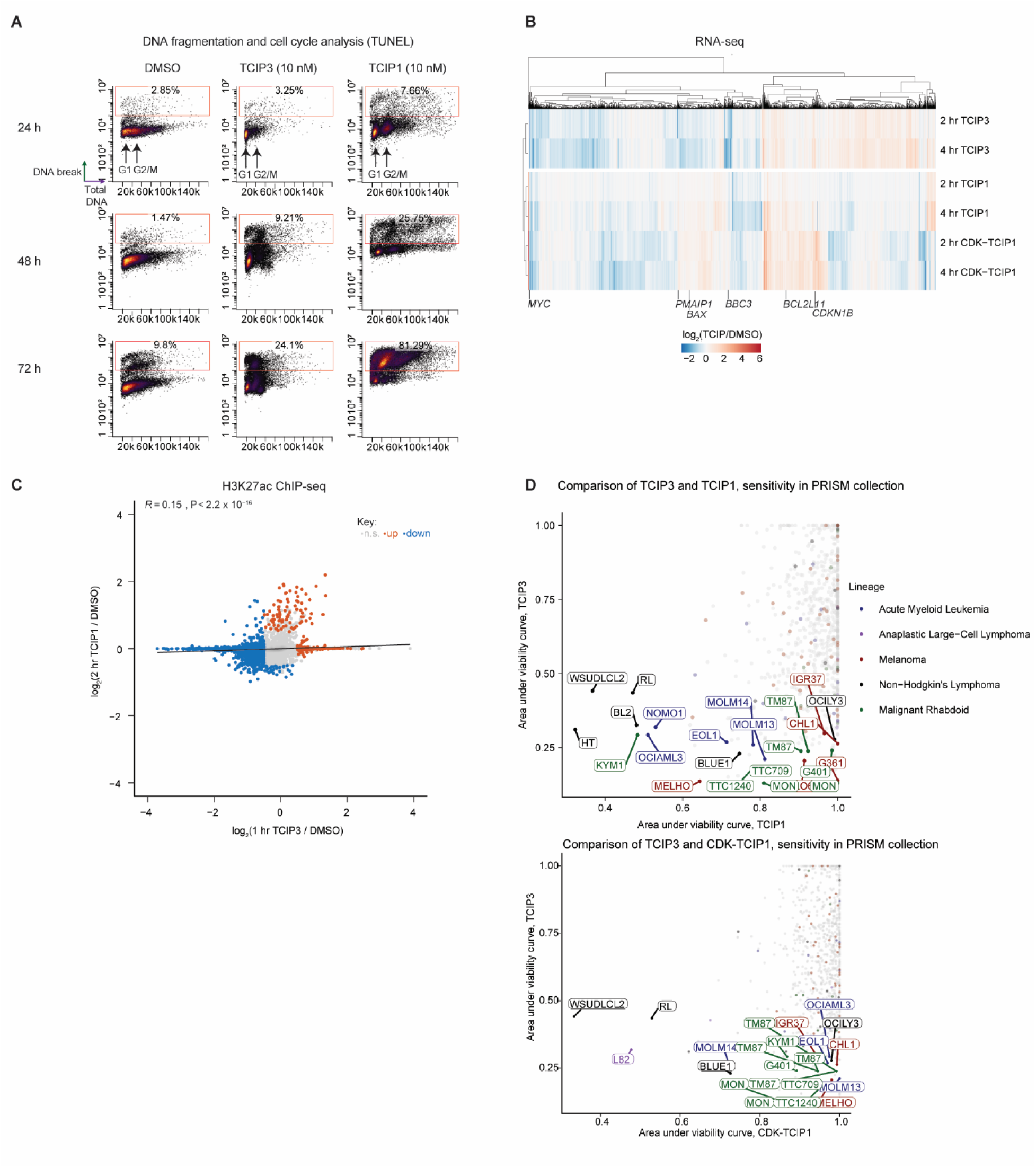
Comparison of BCL6-targeting TCIPs. **(A)** Concurrent analysis of apoptosis and cell cycle effects of TCIPs by terminal deoxynucleotidyl transferase dUTP nick end labeling (TUNEL) and total DNA content co-staining of SUDHL5 cells treated with DMSO, 10 nM TCIP1, or 10 nM **TCIP3** for 24, 48, or 72 h; corresponds to Fig. 4G; representative of 4 biological replicates; cell cycle phases labeled. **(B)** Unbiased clustering of differential gene expression caused by **TCIP3**, TCIP1^39^, and CDK-TCIP1^40^; differential genes were defined by adj. *P* ≤ 0.05 and |log_2_(fold change)| ≥ 0.5, *P*-values computed by two-sided Wald test and adjusted for multiple comparisons by Benjamini-Hochberg; 3-4 biological replicates. **(C)** Correlation of changes at all H3K27ac peaks (86,087) between TCIP1 and **TCIP3** treatment; colored are peaks that change significantly for either treatment, significance defined by adj. *P* ≤ 0.05 and |log_2_(fold change)| ≥ 0.5, *P*-values computed by two-sided Wald test and adjusted for multiple comparisons by Benjamini-Hochberg; 2 biological replicates; *R* computed by Pearson’s correlation and *P-*value computed by two-sided Student’s t-test. **(D)** Comparison of sensitivity of cell line to **TCIP3** with sensitivity to TCIP1^39^ and CDK-TCIP1^40^ in ∼900 cancer cell lines in PRISM^55,96^; labeled are top-ranked (most sensitive) lines colored by lineage.

**Supplemental Figure 8.**
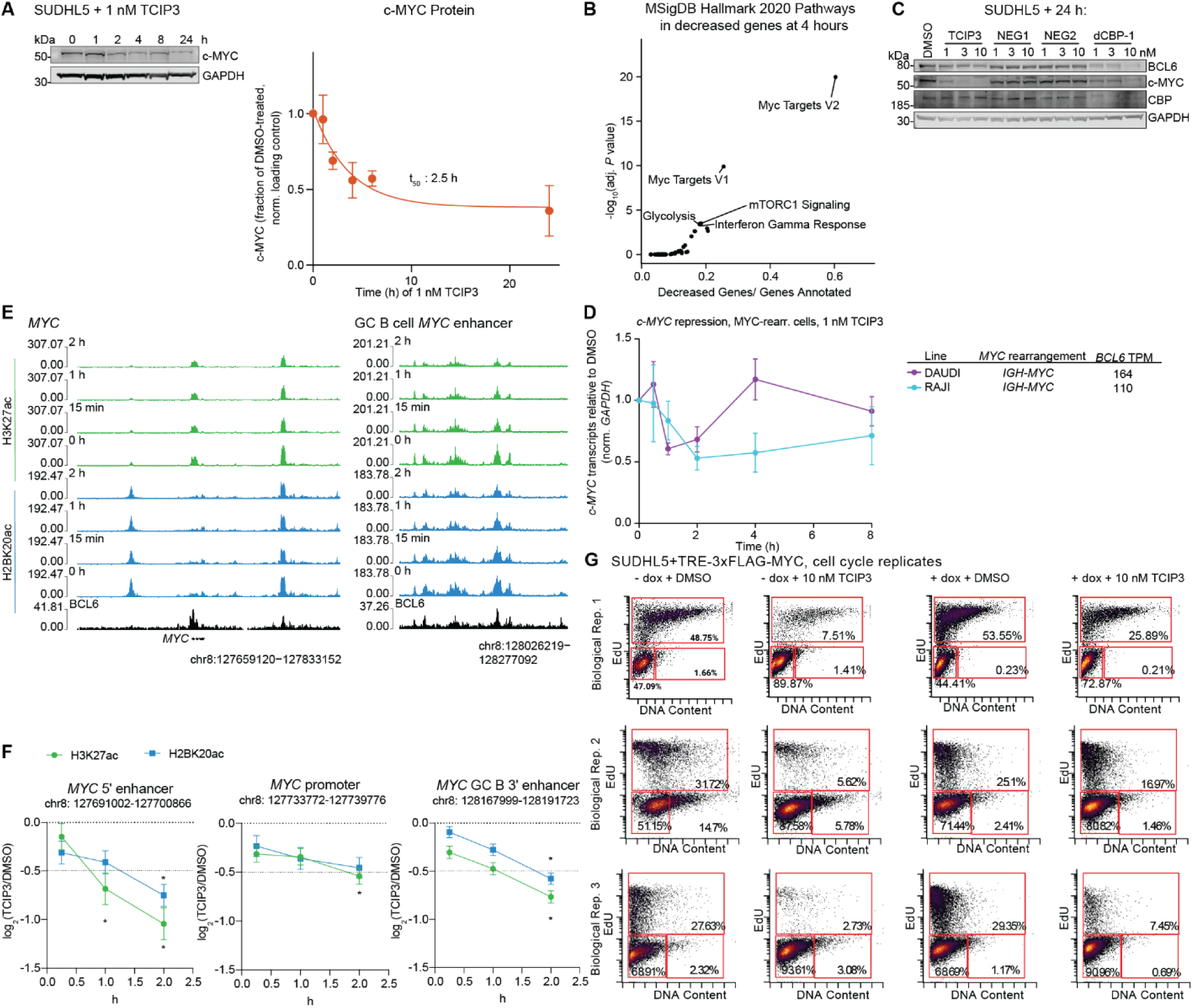
Characterization of c-MYC Repression. **(A)** c-MYC protein in SUDHL5 cells treated with 1 nM **TCIP3** for indicated timepoints; blot representative of 3 biological replicates, plotted is c-MYC normalized to loading controls and DMSO levels, fit to a one-phase exponential decay model, mean ± s.e.m. **(B)** Signaling pathways (MSigDB Hallmark 2020) significantly enriched among significantly decreased transcripts (adj. *P* ≤ 0.05 and log_2_(fold change) ≤ -0.5) after 4 h treatment of **TCIP3** (1 nM) in SUDHL5 cells. **(C)** Protein levels in SUDHL5 cells treated with indicated compounds for 24 h. **(D)** *c-MYC* transcripts in Burkitt’s lymphoma cells with chromosomal rearrangement of the *MYC* locus to the immunoglobulin heavy chain (IGH) locus^79,80^; *BCL6* expression from^95^; cells treated with 1 nM of **TCIP3** normalized to *GAPDH* and DMSO treatment as quantified through RT-qPCR; 3 biological replicates, mean ± s.e.m. **(E)** Changes in H2BK20ac and H3K27ac with time at the promoter of *c-MYC* and its upstream and downstream enhancer regions; BCL6 track is CUT&RUN in untreated SUDHL5 cells, tracks merged from two biological replicates and sequence-depth normalized and, for histone acetylation ChIP-seq, also input-subtracted. **(F)** Quantification of changes in H2BK20ac and H3K27ac differential peaks around the *c-MYC* locus; *: adj. *P* ≤ 0.05 and log_2_(fold change) ≤ -0.5; 2 biological replicates, mean ± s.e.m.; *P*-values computed by two-sided Wald test and adjusted by multiple comparisons by Benjamini-Hochberg. **(G)** Individual biological replicates of cell cycle analysis of SUDHL5^TRE-3xFLAG-MYC^ cells treated with or without 1 µg/mL doxycycline dissolved in ethanol or vehicle 24 h prior to 24 h treatment with 10 nM **TCIP3** or DMSO.

**Supplemental Figure 9.**
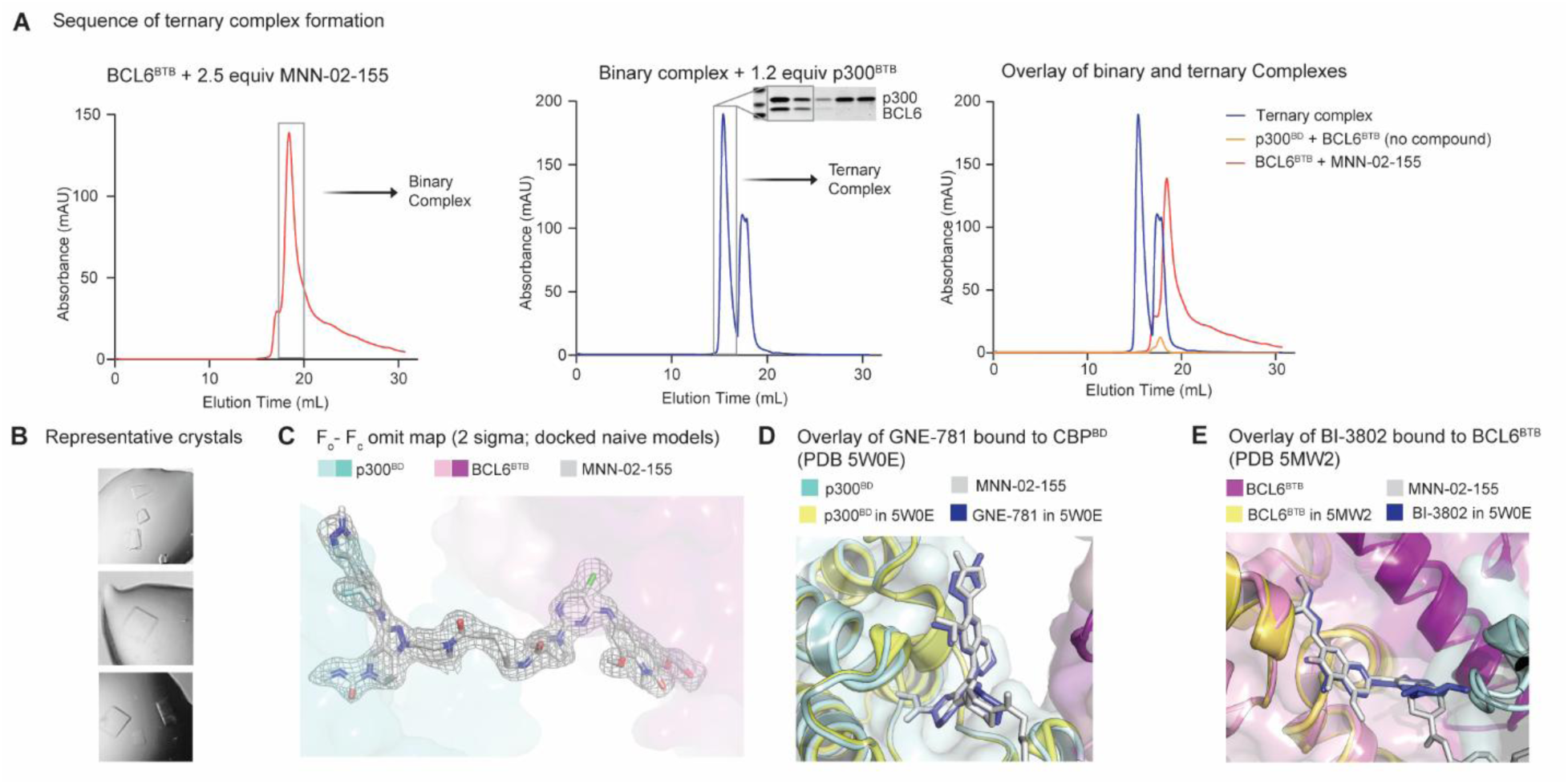
Structural Analyses of KAT-TCIPs. **(A)** Schematic of biochemical formation of a ternary complex between **MNN-02-155**, P300^BD^, and BCL6^BTB^ domains. Binary and ternary complexes were purified by size-exclusion and concentrated to desired concentrations. **(B)** Representative images of crystals containing the ternary complex. **(C)** F_o_-F_c_ map of **MNN-02-155** in the co-crystal structure with p300^BD^ and BCL6^BTB^ domains. **(D)** Overlay of the co-crystal structure with the published structure of GNE-781 bound to CBP (PDB: 5W0E). **(E)** Overlay of the co-crystal structure with the published structure of BI-3812 bound to BCL6 (PDB: 5MW2).

**Supplemental Figure 10.**
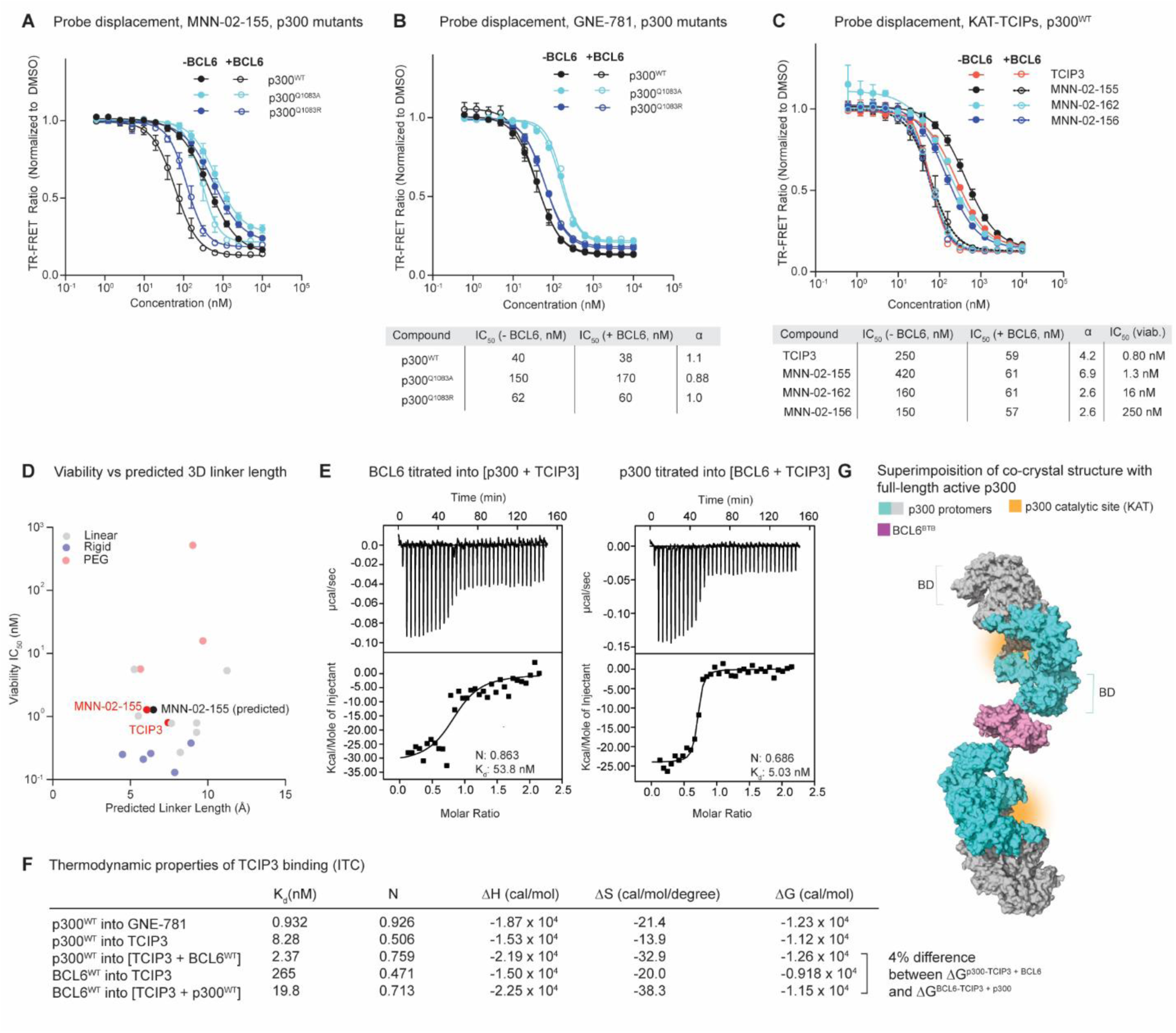
Biophysical Analyses of KAT-TCIPs. **(A)** Binary TR-FRET displacement curves of **MNN-02-155** corresponding to Fig. 7D. **(B)** Binary TR-FRET displacement assay and cooperativity analysis evaluating the inhibition of GNE-781 against p300^WT^, p300^1083A^, p300^1083R^ in the absence or presence of BCL6 (>>> K_d_). n = 2-3 biological replicates; mean ± s.e.m. **(C)** Binary TR-FRET displacement assay and cooperativity analysis evaluating the inhibition of **TCIP3, MNN-02-155, MNN-02-162, or MNN-02-156** against p300^WT^ in the absence or presence of BCL6 (>>> K_d_); cell viability IC_50_ after 72 h treatment in SUDHL5 (Supplemental Fig. 1) labeled in table. n = 2-3 biological replicates; mean ± s.e.m. **(D)** IC_50_s (nM) of KAT-TCIPs synthesized from GNE-781 treated for 72 h in SUDHL5 (from Supplemental Fig. 1A, 2A) plotted against predicted lowest-energy-conformation linker lengths (average of the 2-3 lowest energy conformations). **(E)** ITC traces of binding events to form a ternary complex, including BCL6^BTB^ (10 µM) titrated into p300^BD^ (20 µM) and **TCIP3** (1 µM), or p300^BD^ (20 µM) titrated into BCL6^BTB^ (40 µM) and **TCIP3** (2 µM), and respective N and K_d_ values; corresponds to Fig. 6E; representative of 3 biological replicates. **(F)** Thermodynamic parameters of binding measured from ITC. All measurements represent the mean of 2-3 biological replicates. **(G)** Superimposition of the co-crystal structure of **MNN-02-155** engaged with p300^BD^ and BCL6^BTB^ with the active p300 core, containing two p300 protomers, one teal and one gray (PDB: 6GYR).

**Supplemental Figure 11. Full Scans of Western Blots.**

**Supplemental Figure 12. Flow Gating Strategies.**

**Supplemental Table 1. Enriched proteins after p300 IP-MS in SUDHL5 cells, FLAG IP-MS in FLAG-tagged *BCL6* SUDHL5 cells, or after global proteome profiling after TCIP3, dCBP-1, NEG1, and NEG2 treatment in SUDHL5 cells.**

**Supplemental Table 2. Full results of enrichment of histone acetylation peaks in public transcription factor ChIP-seq datasets in blood-lineage cells (ChIP-atlas).**

Contains all significantly enriched TFs in H3K27ac and H2BK20ac differential peaks. Legend is on Sheet 1 of the Table.

**Supplemental Table 3: Crystallographic data collection and refinement statistics. Supplemental File 1: Chemical Synthesis and Characterization.**

**Supplemental Video 1: Co-Crystal Structure of MNN-02-155 in complex with p300^BD^ and BCL6^BTB^.**

## LIMITATIONS OF THE STUDY

KAT-TCIPs recruit the acetyltransferases p300/CBP to the master transcriptional repressor BCL6 to activate cell death in DLBCL cells. We primarily assessed our lead KAT-CIP molecule **TCIP3** in DLBCL cell lines with either high or low BCL6 expression, and the extent to which **TCIP3**’s pharmacology can be generalized across other DLBCL subtypes and other cancers remains to be explored. Additionally, we only incorporated two p300/CBP bromodomain-containing ligands and one BCL6-binding ligand into our KAT-TCIP designs. Other ligands may demonstrate different potencies and pharmacology. Although we investigated KAT-TCIP mediated acetylation of BCL6 and histone proteins, we did not comprehensively assess the possibility of other protein substrates. We did not investigate the *in vivo* efficacy of **TCIP3** in mouse models of DLBCL. **TCIP3** should be considered a tool molecule to manipulate cancer cell signaling and not a therapeutic candidate.

## ACKNOWLEDGEMENTS

We thank members of the Crabtree and Gray laboratories for constructive comments. We thank Dr. Anja Deutzmann and Dr. Jesse Engreitz for discussions about KAT-TCIP induced *c-MYC* repression, and Dr. Caitlin Mills for her suggestions regarding cell death studies. We thank members of the Weis laboratory for allowing us to use their ITC instrument. This work was supported by the Howard Hughes Medical Institute (G.R.C.); the National Institutes of Health (grants CA276167, CA163915, and MH126720-01 to G.R.C.; High-End Instrumentation Grant S10OD028697-01 to Stanford Chemistry, Engineering & Medicine for Human Health; R01CA201380 to MRG); the Mary Kay Foundation; and the Schweitzer Family Fund. Use of the Stanford Synchrotron Radiation Lightsource, SLAC National Accelerator Laboratory, is supported by the U.S. Department of Energy, Office of Science, Office of Basic Energy Sciences under Contract No. DE-AC02-76SF00515. The SSRL Structural Molecular Biology Program is supported by the DOE Office of Biological and Environmental Research, and by the National Institutes of Health, National Institute of General Medical Sciences (P30GM133894). The contents of this publication are solely the responsibility of the authors and do not necessarily represent the official views of NIGMS or NIH. M.N.N. was supported by the David L. Sze and Kathleen Donahue Interdisciplinary Fellowship. H.Y. was supported by a Leukemia and Lymphoma Society Special Fellow Award. N.S.G. was supported by funds from the Department of Chemical and Systems Biology and the Stanford Cancer Institute, both at Stanford University. M.R.G. was supported by a Leukemia & Lymphoma Society Scholar award, the MD Anderson B-cell Lymphoma Moonshot Program and the Schweitzer Family Fund. The Gray lab also receives or has received research funding from Novartis, Takeda, Astellas, Taiho, Jansen, Kinogen, Arbella, Deerfield, Springworks, Interline, and Sanofi.

## MATERIALS AND DATA AVAILABILITY

The mass spectrometry proteomics data have been deposited to the ProteomeXchange Consortium via the PRIDE^97^ partner repository with the dataset identifier PXD059919. Genomic sequencing data has been deposited to GSE287542 and GSE287543. The X-ray co-crystal structure of MNN-02-155 in complex with p300^BD^ and BCL6^BTB^ has been deposited to the Protein Data Bank (PDB: 9MZA). All other materials are available from the authors upon request.

## AUTHOR CONTRIBUTIONS

G.R.C., N.S.G., M.N.N., R.C.S., and S.G. conceived the project. S.G. and M.N.N. conducted cell biological, biochemical, and genomic studies and contributed equally to this work. M.N.N. and R.C.S. designed and conducted chemical syntheses. S.G. and S.A.N. conducted ChIP-seq studies with help from M.N.N. S.A.N. made the FLAG-BCL6 line and S.A.N. and B.G.D. performed IP-MS experiments. S.M.H, M.N.N., M.M., D.F., and S.G. conducted structural and biochemical studies. B.G.D. performed proteomic experiments. H.A., M.M, B.R., J.M.S., C.L., and H.M.J. performed experiments designed by G.R.C., S.G., N.S.G., M.N.N., B.G.D., and S.M.H. M.M. conducted NanoBRET studies. H.A performed Western Blots and generated lentiviruses. Y.W. assisted with studies of cell death. H.Y. and M.R.G. conducted BCL6 CUT&RUN studies and contributed gene set enrichment analyses relevant to DLBCL. T.L. synthesized the BI3812-BODIPY probe. A.K. and T.Z. contributed to TCIP biological application and chemical synthesis, respectively. L.C. and M.M.D. contributed to cytotoxicity testing in primary lymphocytes. G.R.C., M.N.N., S.G., S.M.H, and N.S.G. wrote the manuscript with input from all authors.

## METHODS

### Cell Culture

Lymphoma and leukemia cells were grown in RPMI-1640 medium (ATCC 30-2001) supplemented with 10% fetal bovine serum (FBS) and 1% 100X Penicillin-Streptomycin (Gibco, 15140122) in incubators at 37°C with 5% carbon dioxide. KARPAS422 cells were obtained from Sigma (06101702) and DB and SUDHL5 were obtained from the American Tissue Culture Collection (ATCC). The Daudi cell line, originally from ATCC, was generously provided by R. Levy’s laboratory at Stanford University. Raji cells, also from ATCC, were a gift from J. Cochran’s laboratory at Stanford University. TOLEDO and K562 cells were originally obtained from ATCC and were kindly shared by A. Alizadeh’s laboratory at Stanford University. Primary human tonsillar lymphocytes were from two separate donors (male, 6 years old; male, 44 years old) from the laboratory of M. M. Davis under IRB protocol numbers IRB-60741 (adult tonsils) and IRB-30837 (pediatric tonsils). BJ CRL-2522 human fibroblasts were obtained from ATCC and grown in DMEM media (ThermoFisher 11965118) supplemented with 10% fetal bovine serum (FBS) and 1% 100X Penicillin-Streptomycin (Gibco, 15140122) in incubators at 37°C with 5% carbon dioxide. Cells were routinely checked for mycoplasma and immediately checked upon suspicion. No cultures tested positive.

### Cell Viability Measurements

#### Compound treatment for 72 hours

Thirty thousand cells were seeded in 100 µl of media per well of a 96-well plate and treated with drug for the indicated times and doses. A resazurin-based indicator of cell health (PrestoBlue; P50200, Thermo Fisher) was added for 1.5 h at 37°C, at which point the fluorescence ratio at 560/590 nm was recorded (Tecan Spark). The background fluorescence was subtracted, and the signal was normalized to DMSO-treated cells. IC_50_ measurements on cell lines were calculated using at least three biological replicates (separate cell passages). Fit of dose–response curves to data and statistical analysis was performed using GraphPad PRISM using the four-parameter log(inhibitor) vs response function.

#### Compound treatment for 7 days

Fifteen thousand cells were seeded in 100 µl of media per well of a U-bottom 96-well plate and treated with digitally dispensed drugs (Tecan D300e) for the indicated times and doses. After 96 hours, plates were centrifuged for 750 rpm for 3 minutes, and old media was exchanged for 100 µl of fresh media. Drug treatment was then repeated at the indicated doses. 72 hours following media change, cells were transferred to 96 well flat-bottom opaque plates. Viability was measured by adding 25 µL Cell Titer-Glo reagent (Promega G7570) to each well and luminescence was measured (BMG Labtech Pherastar FS). The background luminescence was subtracted, and the signal was normalized to DMSO-treated cells. IC_50_ measurements on cell lines were calculated using at least three biological replicates by separate cell passages. Fit of dose–response curves to data and statistical analysis was performed using GraphPad PRISM using the four-parameter log(inhibitor) vs response function.

### Trypan Blue Cell Counting Assay

Thirty thousand cells were seeded into a 96-well plate and treated with digitally dispensed drugs (Tecan D300e) for the indicated times and doses. At either 24, 48, 72, or 96 hours, cells were transferred to a U-bottom well, centrifuged for 4 minutes at 500g, then aspirated. 10 µL phosphate-buffered saline (PBS) pH 7.4 and 10 µL Trypan Blue (Invitrogen T10282) were added to the well and mixed, at which point 10 µL of the mixture was transferred to a cell counting slide and the percentage of cells alive was recorded.

### PRISM Cell Proliferation Assay

The PRISM cell proliferation assay was carried out as previously described^98^. Briefly, up to 859 barcoded cell lines in pools of 20-25 were thawed and plated into 384-well plates (1250 cells/well for adherent cells, 2000 cells/well for suspension or mixed suspension/adherent pools). Cells were treated with an 8-point dose curve starting at 10 µM with threefold dilutions in triplicate and incubated for 120 hours, then lysed. Each cell’s barcode was read out by mRNA-based Luminex detection as described previously^55^ and input to a standardized R pipeline (https://github.com/broadinstitute/prism_data_processing) to generate viability estimates relative to vehicle treatment and fit dose-response curves. The area under the dose-response-curve (AUC), which is correlated with drug potency, was used as a metric of drug potency in a cell line, and correlated (Pearson’s) with dependency of the cell line to gene knockout^96^.

### Protein Expression and Purification

The bacterial expression vectors for 6xHis-p300^BD^ and 6xHis-CBP^BD^ used for isothermal calorimetry, crystallography, and TR-FRET assays were generous gifts from Nicola Burgess-Brown (p300: Addgene plasmid 746580; http://n2t.net/addgene:74658; RRID:Addgene_74658; CBP: Addgene plasmid 38977; http://n2t.net/addgene:38977; RRID:Addgene_38977). Mutations in p300 (Uniprot: Q09472) were introduced by site-directed mutagenesis (NEB E0554). The bacterial expression vector for BCL6^BTB^ used for isothermal calorimetry and crystallography (renamed pSG219C) was created as follows: codon-optimized coding sequences for human BCL6 (residues 5-129; Uniprot: P41182) and were synthesized and cloned into pET-48b(+) (Novagen). The open reading frame codes for N-terminal Trx and 6xHis tags as well as a 3C cleavage site. The bacterial expression vector for BCL6^BTB^ used for TR-FRET assays (renamed pSG233) included a C-terminal GS linker and AviTag (GLNDIFEAQKIEWHE) used for biotinylation. Both vectors coded for the following BCL6 mutations: C8Q, C67R, C84N^99^. These enhance stability but do not affect the affinity for BI3812 or for SMRT.

Protein expression was carried out for 18 hours in Rosetta(DE3) cells (Novagen 70954) at 18°C before pelleting cells by centrifugation. Cell pellets were resuspended in ∼2 mL per liter buffer D800 (20 mM HEPES, pH 7.5; 800 mM NaCl; 10 mM imidazole, pH 8.0; 2 mM beta-mercaptoethanol; 10 % glycerol (*v:v*)) supplemented with protease inhibitors (1 mM PMSF, 1 mM benzamidine, ∼20 µg/ml pepstatin, aprotinin, and leupeptin), and stored at -80 ***°***C.

To purify 6xHis-CBP^BD^ and 6xHis-p300^BD^, cell pellets were thawed in warm water. All subsequent steps were carried out at 4 ***°***C or on ice. Cells were lysed by sonication before centrifugation at 3,214 g for 1 hour. Clarified lysate was mixed with ∼0.5 mL/L of culture cobalt resin (GoldBio) for one hour. Beads were washed by low-speed centrifugation and subsequently by gravity flow with ∼25 column volumes of buffer D800 (followed by ∼10 column volumes of buffer B50 (D800 but with 50 mM NaCl). Protein was eluted with 50 mL C50 (B50 with 400 mM imidazole), and the eluate was applied to a 5 mL anion exchange column (Q HP, Cytiva) and eluted via salt gradient (8 CV; B50 to D800). Peak fractions were concentrated by ultrafiltration before application to a 24 ml gel filtration column (S200 increase, Cytiva) charged with GF150 buffer (20 mM Tris-HCl, pH 8.5, 150 mM NaCl, 1 mM tris(2-carboxyethyl)phosphine (TCEP)). Peak fractions were again concentrated by ultrafiltration, supplemented with 5% glycerol by volume (final), and aliquoted and frozen at -80 ***°***C. Individual aliquots were thawed and stored for no more than 24 hours at 4 ***°***C or on ice before use or disposal.

Purification of BCL6^BTB^ for crystallography and isothermal calorimetry followed the same procedure as 6xHis-CBP^BD^ and 6xHis-p300^BD^ with the following modifications: 6xHis-3C (homemade, produced with pET-NT*-HRV3CP, a kind gift from from Gottfried Otting (Addgene plasmid 162795; http://n2t.net/addgene:162795; RRID:Addgene_162795)) was added to the eluate from the anion exchange column for 18 hours at 4 ***°***C under slow rotation. Imidazole concentration was adjusted to 50 mM and the mixture was applied to a 5 mL nickel column (HisTrap FF Crude, Cytiva) and the flow-through captured. Flow-through was concentrated by ultrafiltration before application to a gel filtration column and peak fractions were concentrated and stored as written above.

Purification of biotinylated BCL6^BTB^-AviTag for TR-FRET assays followed the same procedure as BCL6^BTB^ with the following modification: following cleavage of the tag with 6xHis-3C, the mixture was charged with 14xHis-BirA (homemade; produced with pTP264, a kind gift from Dirk Görlich (Addgene plasmid 149334; http://n2t.net/addgene:149334; RRID:Addgene_149334)) with 0.01 mM D-biotin (Sigma 2031), 10mM MgCl_2_, and 10mM ATP and nutated for 1 hour 30 ***°***C. The sample was applied to a 5 mL nickel column (HisTrap FF Crude, Cytiva) and the flow-through captured. Flow-through was concentrated by ultrafiltration before application to a gel filtration column and peak fractions were concentrated and stored as written above. Biotinylation efficiency was confirmed to be almost 100% by incubation of a small sample with streptavidin beads and monitoring of the flowthrough by SDS-PAGE.

### TR-FRET Ternary Assay

10 µL reactions containing 10 nM 6x-His-p300^BD^ or 6x-His-CBP^BD^, 200 nM Biotinylated-Avi-BCL6^BTB^, 20 nM streptavidin-FITC (SA1001, Thermo), and 1:400 anti-6x-His-terbium (PerkinElmer 61HI2TLF) in buffer containing 20 mM HEPES pH 7.5, 150 mM NaCl, 0.1% BSA, 0.1% NP-40, and 1 mM TCEP were plated in low-volume 384 well plates. Drugs were digitally dispensed (Tecan D300e) into protein-containing wells, and the plate was allowed to incubate in the dark for 1 hour at room temperature. Emission at 490 nm (terbium) and 520 nm (FITC) was measured on a PHERAstar FS plate reader (BMG Labtech) upon excitation with 337 nm. The ratio of signal at 520 nm to 490 nm was calculated and normalized to DMSO-treated protein.

### TR-FRET Binary Assay

10 µL reactions containing 25 nM 6x-His-p300^BD^ or 6x-His-CBP^BD^, 50 nM MNN-06-112, 1:400 anti-6x-His-terbium (PerkinElmer 61HI2TLF), and with or without 10 µM BCL6^BTB^ in buffer containing 20 mM HEPES pH 7.5, 150 mM NaCl, 0.1% BSA, 0.1% NP-40, and 1 mM TCEP were plated in low-volume 384 well plates. Drugs were digitally dispensed (Tecan D300e) into protein-containing wells, and the plate was allowed to incubate in the dark for 1 hour at room temperature. Emission at 490 nm (terbium) and 520 nm (FITC) was measured on a PHERAstar FS plate reader (BMG Labtech) upon excitation with 337 nm. The ratio of signal at 520 nm to 490 nm was calculated and normalized to DMSO-treated protein.

### Crystallography

In GF150 buffer, 1 mg of BCL6^BTB^ protein was incubated for 1 hour on ice with 2.5 equivalents of MNN-02-155 that was diluted 10-fold from a 10 mM DMSO stock in GF150 prior to mixing. The mixture was then purified by size-exclusion chromatography as above. The centermost peaks were collected, concentrated by ultrafiltration, and then incubated with 1.2 equivalents of 6x-His-p300^BD^ on ice for 1 hour. This mixture was purified by size-exclusion chromatography, which resulted in two distinct peaks. Fractions from the early-eluting peak were collected and concentrated to approximately 4.5 mg/mL in a 3k Amicon Ultra-4 Centrifugal filter. The sample was used immediately for crystallization by the sitting drop vapor diffusion method. Rectangular plate-like crystals grew within 48 hours in multiple conditions. Crystals was cryoprotected in a solution containing the crystallization solution supplemented with 25% glycerol before cryo-cooling by dipping the crystal in liquid nitrogen. Data was collected at Stanford SSRL experimental beamline 9-2. The best diffraction dataset came from crystals grown in 0.15 M DL-Malic acid pH 7.0, 20 % PEG 3,350. Data were integrated and scaled in the P212121 space group using Aimless and data quality analyzed using Ctruncate^100,101^. We used data to a minimum Bragg spacing of 2.1 Å according to the CC1/2 cutoff suggested by Ctruncate. The L-test indicated twinning was not present in the crystal.

Initial phases were determined by molecular replacement using MOLREP [PMID 20057045] as implemented in CCP4^102^. We used as search models crystal structures of the BCL6 BTB domain (6CQ1) and the p300 bromodomain (5BT3) with the small molecule ligands removed^103^. The resolution cutoff for molecular placement was 3 Å, and we specified a multicopy search for hetero-multimers. This operation successfully placed individual copies of BCL6 and p300. Initial refinement using REFMAC^104^ and analysis of the overall Matthews Coefficient indicated the likely presence of two copies each for BCL6 and p300. Thus, these initial coordinates were used as an input for PHASER^105^, which placed two copies of the starting model in the unit cell. Refinement of this model using REFMAC converged rapidly and produced unambiguous extra density corresponding to the compound included in crystallography experiments. In later rounds of refinement, we added this compound, generating restraints using AceDRG^106^. We also added water molecules in later refinement rounds. Manual model adjustments were done in Coot^107^.

We note that, while the L-test indicated no twinning, the presence of multiple screw axes in the space group, as well as the existence of non-crystallographic symmetry within the biologically relevant protomer (2 BCL6:2 p300:2 TCIP) could potentially complicate data reduction and analysis. Specifically, we explored the possibility of translational non-crystallographic symmetry resulting in the averaging of non-equivalent protomers along a screw axis, which would result in higher-than-expected intensity statistics at low resolutions and would thus complicate L-tests for twinning. Unaccounted-for tNCS may also explain the relatively high Rfree value in the final refined model. However, the Phaser-MR solution was superior given the P212121 space group versus other orthorhombic possibilities (PHASER Refined LLG = 2749.7 versus 701.5 for the next best space group, P21221). Additionally, refinement produced features of the small molecule, which was not present in the molecular replacement models, indicative of high quality and unbiased maps. We note the presence of similar cases of potential unresolved tNCS in the literature with similar statistics. See^108^ and references therein. The final refined model has Rwork/Rfree values of .217/.281. There are no Ramachandran outliers, and the overall clashscore is 9.

### Isothermal Calorimetry

12 hours prior to performing ITC, frozen stocks of 6xHis-p300^BD^ and BCL6^BTB^ were dialyzed for two hours in ITC buffer at 4°C (50 mM HEPES pH 7.4, 100 µM TCEP, 150 mM NaCl), followed by dialysis in fresh ITC buffer overnight at 4°C to remove glycerol and equalize protein mixtures. Samples were centrifuged at 10,000g for 10 min to remove any precipitate. In all titrations, DMSO was added to protein mixtures to match the DMSO concentration of ligand dissolved in ITC buffer. For binary assays with TCIP3, dialyzed 6xHis-p300^BD^ or BCL6^BTB^ were titrated from the syringe into a cell containing TCIP3 dissolved in ITC buffer. For p300 titrations, 50 µM or 25 µM protein was titrated into 5 µM or 2.5 µM, respectively, of TCIP3. For BCL6 titrations, 70 µM or 40 µM protein was titrated into 7 µM or 5 µM, respectively, of TCIP3. For ternary titrations, 20x 6xHis-p300^BD^ was incubated with 1x TCIP3 in the cell to drive saturation of the binary complex, followed by a titration of 10x BCL6^BTB^ from the syringe, at 310 rpm stirring at 25°C. The following concentrations were used in each run: 15 µM BCL6^BTB^, 30 µM p300, and 1.5 µM TCIP3; 18 µM BCL6 ^BTB^, 36 µM 6xHis-p300^BD^, and 1.8 µM TCIP3; or 10 µM BCL6 ^BTB^, 20 µM 6xHis-p300^BD^, and 1 µM TCIP3. Alternatively, 20x BCL6^BTB^ was incubated with 1x TCIP3 in the cell to drive saturation of the binary complex, followed by a titration of 10x 6xHis-p300^BD^ from the syringe, at 310 rpm stirring at 25°C. The following concentrations were used in each run: 30 µM BCL6, 15 µM 6xHis-p300^BD^, and 1.5 µM TCIP3; 20 µM BCL6 ^BTB^, 10 µM 6xHis-p300^BD^, and 1 µM TCIP3; or 40 µM BCL6 ^BTB^, 20 µM 6xHis-p300^BD^, and 2 µM TCIP3. The first one or two injections and outliers from instrument noise were routinely excluded. Data were fit to a one-site model using MicroCal LLC Origin software. The following injection parameters were used for each run: (Total injection number: 35; Cell temp: 25 °C; Ref power: 10; Initial delay: 250 s; stirring speed: 310; Feedback mode: High; volume: 8 µL; Duration: 13.7 s; Filter period: 2; initial injection volume: 2 µL).

### nanoBRET

#### BCL6^BTB^ nanoBRET

The construct for NanoLuciferase(NanoLuc)-tagged BCL6^BTB^ was created as follows in a lentiviral construct: an N-terminal NanoLuc (subcloned from pNLSF-1, Promega N1351) was fused to the BTB domain of human BCL6 (Uniprot: P41182; aa1-129) with an internal GSG linker followed by a V5 tag. HEK293T cells were plated at a density of 6x10^5^ cells/mL in 2 mL of DMEM/well in a tissue culture treated 6-well plate and were allowed to incubate overnight at 37 °C. The next day, each well of cells was transfected with 2 ug of NanoLuc-BCL6^BTB^ (renamed pNSG218) using Lipofectamine 2000. The transfected cells were allowed to incubate overnight at 37 °C. The following day, the transfected cells were trypsizined, washed with PBS, counted with Trypan Blue, and brought to a final concentration of 1.25x10^5^ cells/mL in Fluorobrite DMEM (Gibco A1896701) supplemented with 10% FBS. 1 uM TNL-15 probe was added to the cells before plating 5000 cells per well in a volume of 40 uL per well in a 384-well plate. Control wells without the inclusion of TNL-15 probe were plated before its addition to the cells to be used as a negative control for data analysis. The plated cells were allowed to incubate overnight at 37 °C. The next day, the cells were treated with test compounds using a Tecan D300e Digital drug dispenser, normalized with DMSO, and allowed to incubate at 37 °C for 1 hour. After 1 h, 5 uL of NanoBRET NanoGlo Substrate plus Extracellular NanoLuc Inhibitor was added to each well (Promega N1661) and mixed on an orbital shaker at 200 g for 30 seconds. Data was then obtained using a PheraStar FS plate reader (BMG Labtech) measuring luminescence with 520-BP and 450-BP filters. Wells were normalized to treatment with DMSO.

#### p300^BD^ and CBP^BD^ nanoBRET

The constructs for NanoLuc-tagged p300^BD^ and CBP^BD^ were subcloned in pNLF-1 (Promega N1351) and created as follows: an N-terminal NanoLuc was fused to the bromodomains(BD) of human p300(Uniprot: Q09472; aa1040-1161) or human CBP (Uniprot: Q92793; aa1081-1197) with an internal GSSG linker. HEK293T cells were plated at a density of 6x10^5^ cells/mL in 2 mL of DMEM/well in a tissue culture treated 6-well plate and were allowed to incubate overnight at 37 °C. The next day, each well of cells was transfected with 2 ug of NanoLuc-p300^BD^ and NanoLuc-CBP^BD^ using Lipofectamine 2000. The transfected cells were allowed to incubate overnight at 37 °C. The following day, the transfected cells were trypsizined, washed with PBS, counted with Trypan Blue, and brought to a final concentration of 1.25x10^5^ cells/mL in Fluorobrite DMEM supplemented with 10% FBS. 100 nM of MNN-05-112 probe was added to the cells before plating 5000 cells per well in a volume of 40 uL per well in a 384-well plate. Control wells without the inclusion of MNN-05-112 probe were plated before its addition to the cells as a negative control for data analysis. The plated cells were allowed to incubate overnight at 37 °C. The next day, the cells were treated with test compounds using a Tecan D300e Digital drug dispenser, normalized with DMSO, and allowed to incubate at 37 °C for 1 hour. After 1 h, 5 uL of NanoBRET NanoGlo Substrate plus Extracellular NanoLuc Inhibitor was added to each well and mixed on an orbital shaker at 200 g for 30 seconds. Data was then obtained using a PheraStarFS plate reader (BMG Labtech) measuring luminescence with 520-BP and 450-BP filters. Wells were normalized to treatment with DMSO.

For all NanoBRET assays, a dose-response curve with 3 technical replicates was constructed for each compound, and corrected BRET ratios were calculated according to manufacturer assay protocol (Promega TM439). Data was fit using a standard four parameter log-logistic function using the R package drc or using GraphPad Prism.

### Annexin V staining

25,000 cells were plated in 200 µL of media in 96 well U-bottom plates and treated with compound. 72 hours later, the plate was spun at 500g for 4 minutes at 4°C, at which point cells were washed in 100 µL cold 2.5% FBS in PBS. Centrifugation and washing was repeated once more. Cells were then resuspended in 50 µL cold binding buffer (10 mM HEPES pH 7.4, 140 mM NaCl, 2.5 mM CaCl_2_) and 2.5 µL FITC-Annexin-V (Biolegend 640922). A no-stain control was also included to draw gates. The plate was allowed to incubate for 15 minutes at room temperature, at which point 100 µL of 2.5% FBS in PBS was added to each well. The plate was gently agitated through shaking, and samples were analyzed by flow cytometry on a BD Accuri. At least 50k events were collected per sample.

### Cell Cycle Analysis

2 million SUDHL5 cells were plated in 4 mL of media in 6-well plates and treated with 10 nM of drug. After 22 hours, cells were dosed with 10 µM 5-ethynyl-2’-deoxyuridine (EdU) and allowed to incubate for an additional 2 hours, at which point cells were harvested and collected by centrifuging at 300g for 4 minutes at 4°C. Cells were washed once with 1% BSA in PBS. 1.1 million cells in 1% BSA/PBS were then transferred to flow cytometry polypropylene tubes and centrifuged at 300g for 4 minutes at 4°C. Supernatant was removed, and cells were fixed for 15 minutes in the dark at room temperature with 100 µL of 4% PFA in PBS, in which gentle flicking was used to resuspend cells. Cells were then washed with 1 mL of 1% BSA/PBS and permeabilized with 100 µL of 0.1% Triton-X for 30 minutes at room temperature in the dark. Cells were washed with 3 mL 1% BSA/PBS, then incubated in the dark for 30 minutes with a cocktail of AlexaFluor-488-Azide (438 µL PBS, 10 µL 100 mM CuSO_4_, 2 µL 488-azide, and 50 µL of 1x EdU Reaction Buffer dissolved in water) (Invitrogen C10337). Cells were washed with 1 mL 1% BSA in PBS, then incubated with a reaction cocktail of 7-AAD/RNAseA for 30 minutes in the dark (500 µL 1% BSA/PBS, 2 µL 7-AAD (BD 559925), 5 µL RNAseA (Invitrogen 12091021)).

Cells were washed with 1 mL 1% BSA/PBS centrifuged a final time, then resuspended in 500 µL 1% BSA/PBS. Cells were gently agitated by shaking, then analyzed through flow cytometry on a BD Accuri. At least 100k events were collected per sample. Gates were drawn from single-stain and no-stain controls.

### TUNEL Analysis of DNA Fragmentation

1 million SUDHL5 cells were plated in 6 mL of media in 6-well plates and treated with compound for indicated timepoints and doses. 1.2 million total cells were counted, washed in PBS, fixed in 4% paraformaldehyde/PBS at a concentration of 10M/mL for 15 minutes at room temperature in the dark, washed in PBS, and stored in 70% ethanol at - 20°C until ready for processing (at least 24 hours). DNA breaks in fixed cells were labeled with bromolated deoxyuridine (Br-dUTP or BrdU) by incubation with deoxynucleotidyl transferase (TdT) for 60 min at 37°C, washed, and stained with FITC-labeled anti-BrdU antibodies for 30 min at room temperature in the dark (BD 556405). Cells were resuspended in 1% BSA/PBS with 1:100 RNaseA (Invitrogen 12091021) and co-stained with 7-AAD for 30 min at room temperature in the dark (BD 559925). The suspension was gently agitated by shaking and analyzed by flow cytometry within 1 hour (BD Accuri).

### BCL6 Reporter Assay

KARPAS422 cells were lentivirally transduced with a construct containing the reporter. Description of the reporter construct has been published previously^39^. After selection, cells were plated and treated with indicated amount of compound for 24 hours. Cells were washed in 2.5% FBS/PBS, stained with 1:250 v/v of 7-AAD (BD 559925) to distinguish live from dead cells, and harvested for flow cytometry on a BD Accuri. Given the polyclonal population after transduction, the area under the curve of the histogram representing FITC signal across all live cells was calculated as an integrative measure of total GFP signal. A GFP-positive gate two standard deviations from the mean was drawn from non-transduced cells and the area past a constant threshold was calculated and normalized to the signal from cells treated with DMSO.

### Lentivirus Production and Overexpression

#### c-MYC overexpression

An N-terminal fusion of 3xFLAG followed by a GGSGS linker fused to the coding sequence of human *c-MYC* (NM_002467.6)) was subcloned into pCW57-MCS1-2A-MCS2 (a gift from Adam Karpf (Addgene plasmid 71782; http://n2t.net/addgene:71782; RRID:Addgene_71782)).

#### NanoLuc-BCL6^BTB^ overexpression

This construct is the same used for BCL6 nanoBRET studies above. Lentivirus was produced from HEK293T cells via polyethylenimine transfection. Briefly, cells were transfected using 2^nd^-generation packaging plasmids and media replaced after 24 hours. Media containing virus 72 h after transfection was harvested, filtered, and used immediately or concentrated in PBS by ultracentrifugation (2 hours, 20,000 rpm, Beckman-Coulter Optima XE), flash-frozen, and stored at -80°C. Cells were lentivirally transduced with the overexpression construct by spinfection of virus (1000 g for 1 hour at 30°C) and selected using puromycin. Cells were maintained under antibiotic selection throughout experimental procedures.

### RNA Extraction, qPCR, and Sequencing Library Preparation

Cells were plated at 1M/mL and harvested in TRIsure (Bioline 38033). RNA was extracted using Direct-zol RNA MicroPrep columns (Zymo R2062) treated with DNAseI. cDNA was prepared (Meridian Bioscience BIO-65054) and used for qPCR (Meridian Bioscience BIO-94050) using an AppliedBiosystems QuantStudio 6Pro. Primer sequences were the following:

*c-MYC* fwd: CCTTCTCTCCGTCCTCGGAT;

*c-MYC* rev: CTTCTTGTTCCTCCTCAGAGTCG;

*GAPDH* fwd: GCCAGCCGAGCCACAT;

*GAPDH* rev: CTTTACCAGAGTTAAAAGCAGCCC.

For sequencing library preparation, rRNA was depleted (NEB E7400) and total RNA prepared into paired-end libraries (NEB E7765) and indexed (NEB E7335). Library size distributions were confirmed using an Agilent Bioanalyzer and High Sensitivity DNA reagents (Agilent 5067) and concentrations determined by qPCR. Equimolar pooled libraries were sequenced on an Illumina NovaSeq with 2 x 150 bp cycles.

### Acetyl-lysine and BCL6 Immunoprecipitation for Immunoblotting

Cells were plated at 1M/mL and treated with compound or DMSO at indicated timepoints and doses. Cells were harvested on ice, counted, normalized by count, and washed 1X in PBS containing compound or DMSO. For only BCL6 immunoprecipitation, compound or DMSO was maintained in nuclear preparation buffers at identical concentration to cell treatment throughout. Nuclei were prepared by incubation in Buffer A (25 mM HEPES pH 7.5, 25 mM KCl, 0.05 mM EDTA, 1mM MgCl_2_, 10% glycerol, 0.1% NP-40) supplemented with protease inhibitors (1 mM PMSF, ∼20 mg/ml pepstatin, aprotinin, and leupeptin) at 4 °C for 7 mins. Nuclear preparation was confirmed by Trypan blue staining and nuclei were pelleted by centrifugation for 5 min at 500g at 4 °C. Pelleted nuclei were resuspended in IP Buffer (25 mM HEPES pH 7.5, 150mM KCl, 0.05 mM EDTA, 1 mM MgCl_2_, 10% glycerol, 0.1% NP-40) supplemented with protease inhibitors and 1 µL/250 units benzonase (Sigma E1014). Chromatin was removed by incubation with rotation for 30 min and nuclei sheared using a 27-gauge needle exactly five times. Insoluble material was pelleted by centrifugation at 21,000g for 10 min at 4 °C and the supernatant containing soluble nuclear protein preserved. Extracts were normalized by total protein concentration (Bradford) and identical amounts of total protein and concentrations were used for immunoprecipitation. 1 µg acetylated-lysine antibodies (Cell Signaling 9441), anti-BCL6 antibodies (Cell Signaling D65C10), or normal rabbit IgG (Cell Signaling 2729) and paramagnetic beads conjugated to Protein G (Thermo 10003D) were added to samples and incubated with rotation at 4 °C for 18 hours. Samples were washed five times with 1 mL IP Buffer supplemented with protease inhibitors and eluted by denaturation in 1X NuPage LDS/RIPA sample buffer (Thermo NP0008) supplemented with beta-mercaptoethanol by incubation at 95 °C for 5 mins.

### Western Blots

Cells were plated at 1M/mL and treated with drug at indicated timepoints and doses. Cells were harvested on ice in RIPA buffer (50mM Tris-HCl pH 8, 150mM NaCl, 1% NP-40, 0.1% DOC, 1% SDS, protease inhibitor cocktail (∼20 mg/ml pepstatin, aprotinin, and leupeptin), 1mM DTT) and 1:200 benzonase (Sigma E1014) was added and incubated for 20 mins. After 10 min centrifugation at 14,000g and 4 °C, the supernatant was collected and protein concentration was measured by Bradford. SDS-PAGE analysis was carried out in either 4-12% or 12% Bis-Tris PAGE gels (Thermo NW04120 and NW04127). Antibodies used for immunoblots were: BCL6 (1:1000 *v:v*, Cell Signaling D65C10), p300 (1:250, Santa Cruz F-4 sc-48343), CBP (1:1000, Cell Signaling D6C5), c-MYC (1:1000, Cell Signaling D84C12), Caspase-3 (1:1000, Cell Signaling 9662), GAPDH (1:2000, Santa Cruz 6C5 sc-32233), BBC3/PUMA (1:1000, Cell Signaling E2P7G), FOXO3 (1:1000, Cell Signaling 75D8), H3K27ac (1:1000, abcam ab4729), H2BK20ac (1:1000, Cell Signaling D709W), H3 (1:10,000, Cell Signaling 1B1B2), H2B (1:10,000, Cell Signaling D2H6), p27/CDKN1B (1:1000, Cell Signaling D69C12), and TP53 (1:1000, Santa Cruz DO-1). ImageStudio (Licor) was used for blot imaging and quantification.

### Quantitative Global Proteome Profiling via LC-MS/MS

#### Cell Treatment

UDHL5 cells (6M cells in 6 mL RPMI-1640 media supplemented with 10% FBS and 1x pen-strep) were treated with 0.1% DMSO, 10 nM TCIP3, 10 nM Neg1, 10 nM Neg2, or 250 nM dCBP1 (1000x stocks in DMSO for final 0.1% DMSO concentration) in 3 biologically independent replicates. The cells were washed twice with TBS (50 mM Tris, pH 8.5, and 150 mM NaCl) and then stored at -80 °C until use.

#### Lysis & Digestion

Lysis was performed by first thermally denaturing the samples in residual wash buffer for 5 min at 95 °C. Fresh Lysis Buffer (8 M urea, 150 mM NaCl, and 100 mM HEPES, pH 8.0, in MS-grade water) was then added, and the lysates were homogenized using needle ultrasonication and then normalized to 30 µg of 2 mg/mL by diluting with Lysis Buffer via a Bradford protein concentration assay (Bio-Rad 5000006) in technical duplicates. The samples were then reduced and alkylated simultaneously by adding final concentrations of freshly prepared 10 mM tris(2-carboxyethyl)phosphine hydrochloride (TCEP; Sigma Aldrich C4706) and 40 mM chloroacetamide (CAM; Sigma Aldrich C0267) in 100 mM HEPES, pH 8.0, and shaking for 30-60 min. The samples were then diluted with 120 µL of 20 mM HEPES, pH 8.0, and CaCl_2_ (Sigma Aldrich C4901) for final concentrations of ∼1.3 M urea and 1 mM CaCl_2_. Digestion using 1:100 (w/w) protease to protein lysate with MS-grade Trypsin/Lys-C Protease Mix (Thermo A40009) was then performed by mixing overnight at 37 °C. The samples were then acidified by adding trifluoroacetic acid (TFA) until the sample reached pH ≤ 3, as confirmed by pH paper.

#### SPE Desalting

Samples were then desalted using SOLAµ SPE HRP Peptide Desalting columns (Thermo 60209-001) with a Positive Pressure Manifold and all MS-grade reagents. The desalting cartridges were activated with 200 µL of acetonitrile and then equilibrated with two washes of 200 µL Wash Buffer (2% ACN + 0.2% TFA in water). After loading samples into the cartridges, the samples were washed three times with 200 µL Wash Buffer and then eluted using 100 µL 50% acetonitrile with 0.1% formic acid (FA) in water. The desalted peptides were then immediately dried on a centrifugal vacuum concentrator (Thermo SPD120-115). The samples were then reconstituted in 0.1% FA in water and then analyzed by LC-MS/MS (see below).

### Generation of FLAG-BCL6 knock-in SUDHL5 cell lines

Construction of FLAG-BCL6 knock-in cell lines were adapted from previously described procedures (Savic et al. 2015, Meadows et al. 2020). Plasmid expressing wildtype Cas9 under the control of chicken β-actin (CBA) and a human U6 promoter-driven gRNAs to direct were obtained from Addgene (PX458_BCL6_iso1_1 and PX458_BCL6_iso1_2 were a gift from Eric Mendenhall & Richard M. Myers (plasmids #104046 and #104047; http://n2t.net/addgene:104046 and http://n2t.net/addgene:104047; RRID:Addgene_104046 and 104047, respectively). gRNAs were designed to direct Cas9 nuclease activity near the stop codon of *BCL6*. The donor plasmid contained a 3x FLAG epitope tag, P2A linker, and a Neomycin resistance gene, flanked by regions homologous to the C-terminus of the *BCL6* gene.

SUDHL5 cells were electroporated with donor and gRNA plasmids in a cuvette using the Amaxa™ Cell Line Nucleofector™ Kit V (Lonza, VVCA-1003) and Amaxa™ Nucleofector™ II transfection machine. 2 million cells were resuspended in Nucleofector solution containing supplement with 10 µg pooled plasmid (5 µg donor plasmid and 2.5 µg of each gRNA). This was done in biological triplicate (i.e. three separate electroporations performed), with cells from each electroporation maintained independently thereafter. Immediately after electroporation, cells were transferred to culture flask containing pre-warmed culture media. 24 hours post-transfection, cells were selected with G418 (Geneticin) at a concentration of 800 µg/mL. Selection was performed for 7 days. Cells were maintained under selection as a polyclonal pool for the generation of cell stocks. Subsequently, single clones were isolated by limiting dilution. To validate homologous recombination, genomic DNA from FLAG-BCL6 SUDHL5 cells was isolated using DNeasy Blood and Tissue kit (Qiagen, 69504). The C-terminus of the BCL6 gene was amplified using Q5 Hot Start High Fidelity 2X Master Mix (NEB M0494) with primers that bind internal to the 3x Flag tag and external to the homologous arms. The PCR product was purified using QIAquick PCR Purification Kit (Qiagen) and submitted for Sanger sequencing.

### FLAG and p300 Immunoprecipitation

50 million FLAG-BCL6 knock-in SUDHL5 cells were treated compound or DMSO (0.1%) at indicated doses for 2 hours. Cells were lysed in NE10 buffer (20 mM HEPES (pH 7.5), 10 mM KCl, 1 mM MgCl2, 0.1% Triton X-100 (v/v), protease inhibitors (Roche), 15 mM ß-mercaptoethanol), dounced 15 times and pelleted 5 min at 500 g. Nuclei were washed in NE10 buffer and then digested with 250 units benzonase (Millipore) for 30 min rotating at 25°C. Nuclei were resuspended in NE150 buffer (NE10 supplemented with 150mM NaCl) and incubated for 20 min. Lysates were pelleted at 16,000 g for 20 min at 4°C and supernatants were immunoprecipitated by incubating with 2.5 µg FLAG M2 antibody (F1804, Millipore-Sigma) or 2.5 µg p300 (A300-358A, Bethyl Laboratories) antibody with Dynabeads Protein G (Thermo Fisher) overnight at 4°C. The IP fraction was recovered by magnetic separation followed by three washes with NE10 buffer containing 150mM– 300mM NaCl but without Triton-X. For mass spectrometry analysis, the IP was then eluted from the beads with Ammonium Hydroxide, pH 11-12; 3% v/v.

### Immunoprecipitation-Mass Spectrometry (IP-MS)

Immunoprecipitation was performed as described above for FLAG-BCL6 and p300 in three biologically independent replicates. Enriched proteins were eluted three times with 150 µL of 3% v/v ammonium hydroxide (pH 11-12) for 15 min with gentle agitation. The collected eluate was then dried using a centrifugal vacuum concentrator (Thermo SPD120-115). The samples were reconstituted in 50 µL of reduction-alkylation buffer containing 8 M urea, 10 mM TCEP, 40 mM chloroacetamide, 4.4 mM CaCl_2_, and 100 mM HEPES, pH 8.0 and then incubated for at least 30 min while shaking. The samples were then diluted with 160 µL of 20 mM HEPES, pH 8.0, and then digested overnight at 37 °C with 500 ng of MS-grade Trypsin/Lys-C Protease Mix (50 ng/µL). The samples were then acidified by adding trifluoroacetic acid (TFA) until the sample reached pH ≤ 3, as confirmed by pH paper.

### LC-MS/MS diaPASEF Acquisition & Statistical Analysis

#### diaPASEF LC-MS/MS Analysis

The reconstituted desalted peptides were then analyzed using a nanoElute 2 UHPLC (Bruker Daltonics, Bremen, Germany) coupled to a timsTOF HT (Bruker Daltonics, Bremen, Germany) via a CaptiveSpray nano-electrospray source. The peptides were separated in the UHPLC using an Aurora Ultimate nanoflow UHPLC column with CSI fitting (25 cm x 75 µm ID, 1.7 µm C18; IonOptics AUR3-25075C18-CSI) over a 70 min gradient shown in the table below at a flow rate of 400 nL/min with column temperature maintained at 50 °C using Mobile Phase A (MPA; 3% acetonitrile + 0.1% FA in water) and Mobile Phase B (MPB; 0.1% FA in ACN).

| Time (min) | Composition % (v/v)<br>MPB |
| --- | --- |
| 0 | 6 |
| 40 | 17 |
| 55 | 25 |
| 64 | 34 |
| 65 | 85 |
| 70 | 92 |

The TIMS elution voltages were calibrated linearly with three points (Agilent ESI-L Tuning Mix Ions; 622, 922, 1,222 m/z) to determine the reduced ion mobility coefficients (1/K_0_). diaPASEF was performed using the MS settings 100 m/z for Scan Begin and 1700 m/z for Scan End in positive mode, the TIMS settings 0.70 V⋅s/cm^2^ for 1/K_0_ start, 1.30 V⋅s/cm^2^ for 1/K_0_ end, ramp time of 120.0 ms, 100% duty cycle, ramp rate of 7.93 Hz, and the capillary voltage set to 1600 V. diaPASEF windows from mass range 226.8 Da to 1226.8 Da and mobility range 0.70 1/K_0_ to 1.30 1/K_0_ were designed to provide 25 Da windows covering doubly and triply charged peptides as confirmed by DDA-PASEF scans, whereas singly charged peptides were excluded from the acquisition due to their position in the m/z-ion mobility plane.

#### Raw data processing

The raw diaPASEF files were processed using library-free analysis in FragPipe 22.0. DIA spectrum deconvolution was performed using diaTracer 1.1.5 with the following default settings: (i) “Delta Apex IM” to 0.01, (ii) “Delta Apex RT” to 3, (iii) “RF max” to 500, (iv) “Corr threshold” to 0.3, and (v) mass defect filter enabled with offset set to 0.1. The reviewed Homo sapiens protein sequence database was obtained from UniProt (07/13/2024; 20,468 entities) with decoys and common contaminants. In the MS Fragger 4.1 database search, the following settings were used: (i) initial precursor and fragment mass tolerances of 10 ppm and 20 ppm, respectively, (ii) enabled spectrum deisotoping, mass calibration, and parameter optimization, and (iii) isotope error set to “0/1/2”. For protein digestion, “stricttrypsin” for fully tryptic peptides was enabled with up to 1 missed cleavage, peptide length from 7 to 50, and peptide mass range from 500 to 5,000 Da. For modifications, methionine oxidation (2 max occurrences) and N-terminal acetylation (1 max occurrence) were set as variable modifications (maximum up to 3), while cysteine carbamidomethylation was set as a fixed modification. For validation, MSBooster (DIA-NN model) and Percolator were used for RT and MS/MS spectra prediction and PSM rescoring, while ProteinProphet (--maxppmdiff 2000000) was used for protein inference with FDR filtering (--picked --prot 0.01). The spectral library was then generated by EasyPQP 0.1.49 using default settings. Peptides were then quantified using DIA-NN 1.9.1 with 0.1% FDR and QuantUMS high accuracy settings. The DIA-NN parquet report containing peptide quantification and scoring was then analyzed in R.

#### Statistical Analysis

DIA-NN quantified peptides were then further filtered including (i) common contaminants and reverse sequences, (ii) 1% FDR filtering at the global.q.value (precursor Q value across all samples) and pg.q.value (protein group Q value in single injection), (iii) quantification of a given peptide in at least two replicates in one condition, and (iv) removal of singly charged peptides and non-proteotypic (unique) peptides. Protein intensities were then re-calculated using the MaxLFQ method^109^ provided in the DIA-NN R package. Differential statistics was then performed using the DEqMS^110^ R package, which performs a LIMMA-moderated t-test with an adjustment for number of detected peptides per protein, to determine the p-value, fold change, and Benjamini-Hochberg adjusted p-value. For the mean peptide intensity plots, the arithmetic mean was calculated for unique peptide precursors across conditions imputing a value of 1 unit for replicates where a peptide was not detected. For protein-level imputation in IP-MS, proteins were selected for imputation if (i) a fold change using DEqMS differential statistics could not be calculated and (ii) the protein was not detected in DMSO condition at all but detected in at least all but one replicate in the compound-treated condition. Selected proteins had imputed 15^th^ percentile MaxLFQ protein intensities for the DMSO condition and the subsequent fold changes between DMSO and compound conditions were recalculated.

### RNA-seq Analysis

Raw reads were checked for quality using fastqc (https://www.bioinformatics.babraham.ac.uk/projects/fastqc/) and trimmed from adapters using cutadapt^111^ using parameters cutadapt -a AGATCGGAAGAGCACACGTCTGAACTCCAGTCA -b AGATCGGAAGAGCGTCGTGTAGGGAAAGAGTGT--nextseq-trim=20--minimum-length 1. Transcripts were quantified using kallisto^112^ against the human Gencode v33 indexed transcriptome and annotations. Transcript isoforms were collapsed to genes and differential gene analysis was performed using DESeq2^113^ using apeglm^114^ to shrink fold changes. Pathway analyses were performed using Enrichr^115^ on genes defined by significance cutoffs as detailed in figure legends. GSEA analysis was performed using fgsea^116^ on differential genes ranked by log_2_(fold change) with an in-house dataset consisting of LymphDB (L. Staudt, NIH, https://lymphochip.nih.gov/signaturedb/), MSigDB pathways, and an internal dataset generated at MD Anderson of lymphoma-specific signaling. Correlation analyses with ChIP-seq used only genes with normalized (using DESeq2^113^, relative log expression (RLE)) mean expression ≥ 32; this number was chosen by automatic independent filtering for outlier and low mean counts^113^. Unbiased clustering analyses with TCIP1^39^ and CDK-TCIP1^40^ were performed using pheatmap^117^ using parameters cutree_cols = 6, cutree_rows=2 including only significantly changed genes in any of the treatment conditions (|log_2_(fold change)| ≥ 0.5 and adj. *P* ≤ 0.05; *P*-values computed by two-sided Wald test and adjusted by Benjamini-Hochberg). Re-processed published datasets from the Sequence Read Archive (SRA) were the following: TCIP1 – SRX20228454, SRX20228455, SRX202284546, SRX20228457, SRX20228458, SRX20228459, SRX20228448, SRX20228449, SRX20228450; CDK-TCIP1 – SRX22117221, SRX22117222, SRX22117223, SRX22117224, SRX22117225, SRX22117226, SRX22117227, SRX22117228, SRX22117229).

### ChIP-seq Experiment and Library Preparation

25-30 million cells were treated with TCIP3 or DMSO for indicated timepoints. Cells were washed in PBS containing TCIP3 or DMSO and crosslinked for 11 min in CiA Fix Buffer (50 mM HEPES pH 8.0, 1 mM EDTA, 0.5 mM EGTA, 100 mM NaCl) with addition of formaldehyde to a final concentration of 1%. The crosslinking reaction was quenched by glycine added at 0.125 M final concentration. Crosslinked cells were centrifuged at 1,000 x g for 5 min. Nuclei were prepared by 10 min incubation of resuspended pellet in CiA NP-Rinse 1 buffer (50 mM HEPES pH 8.0, 140 mM NaCl, 1 mM EDTA, 10% glycerol, 0.5% IPEGAL CA-630, 0.25% Triton X100) followed by wash in CiA NP-Rinse 2 buffer (10 mM Tris pH 8.0, 1 mM EDTA, 0.5 mM EGTA, 200 mM NaCl). The pellet was resuspended in CiA Covaris Shearing Buffer (0.1% SDS,1 mM EDTA pH 8.0, 10 mM Tris HCl pH 8.0) with protease inhibitors and sonicated for 20 min with a Covaris E220 sonicator (Peak Power 140, Duty Factor 5.0, Cycles/Burst 200). The distribution of fragments was confirmed with D1000 Tapestation or by agarose gel electrophoresis. 20 µg of chromatin per ChIP was used with anti-H3K27ac antibodies (abcam ab4729) and anti-H2Bk20ac antibodies (Cell Signaling D709W), with 40 ng Drosophila chromatin (53083, ActiveMotif) spiked in. After overnight incubation at 4 °C in IP buffer (50 mM HEPES pH 7.5, 300mM NaCl, 1mM EDTA, 1% Triton X100, 0.1% sodium deoxycholic acid salt (DOC), 0.1% SDS), IPs were washed twice with IP buffer, once with DOC buffer (10 mM Tris pH 8, 0.25 M LiCl, 0.5% IPEGAL CA-630, 0.5% sodium deoxycholic acid salt (DOC), 1mM EDTA), and once with 10 mM Tris/1 mM EDTA buffer (TE) pH 8. IPs and inputs were reverse-crosslinked in TE/0.5% SDS/0.5 µg/µL proteinase K for 55 °C /3 hours then 65 °C /18 hours, then DNA was purified using a PCR cleanup spin column (Takara #74609). Paired-end sequencing libraries were constructed using an NEBNext Ultra II DNA kit (E7645S) and indexed (NEB E7335). Library size distributions were confirmed using an Agilent Bioanalyzer and High Sensitivity DNA reagents (Agilent 5067) and concentrations determined by qPCR. Equimolar pooled libraries were sequenced on an Illumina NovaSeq XPlus with 2 x 150 bp cycles.

### CUT&RUN Experiment and Library Preparation

CUT&RUN was performed as previously described^118,119^ with minor modifications. Briefly, cells were collected and washed by PBS. After final wash, nuclei were isolated using nuclei extraction buffer (20mM HEPES pH 7.9, 10 mM KCl, 0.1% Triton X-100, 20% glycerol, 1x cOmplete protease inhibitors (Roche 11836153001), 0.5mM spermidine) and then washed twice with wash buffer (20mM HEPES pH 7.5, 150mM NaCl, 0.5mM Spermidine, 1x cOmplete protease inhibitor). Five-hundred thousand nuclei were counted and incubated with activated Concanavalin A coated beads (EpiCypher 21-1411) and subsequently mixed with BCL6 antibodies (Cell Signaling D65C10) in antibody buffer (wash buffer + 0.01% digitonin + 2 mM EDTA) for overnight incubation on nutator in a cold room. IgG controls (normal rabbit IgG, Cell Signaling 2729) were set in parallel for antibody enrichment and specificity validation. Nuclei were then washed three times with digitonin buffer (wash buffer + 0.01% digitonin) and incubated with pAG-MNase (EpiCypher 15-1116) for 10 minutes at room temperature. After 10 minutes of incubation, nuclei were washed three times again with digitonin buffer and resuspended with prechilled low-salt, high-Ca2+ buffer (20 mM HEPES pH 7.5, 0.5mM spermidine, 1x cOmplete protease inhibitor, 10mM CaCl_2_) to initiate 1-hour digestion at 0°C using an ice block. The reaction was terminated by switching into stop buffer (340 mM NaCl, 20mM EDTA, 4 mM EGTA, 50 µg/mL RNaseA, 50 µg/mL glycogen) and incubating at 37°C for 10 minutes to release digested DNA from the nuclei. DNA was then purified with Ampure beads (Beckman A63882), subjected to two-sided size selection(0.5X to remove large fragment and 2X to recover desired nucleosomal DNA fragments). Sequencing libraries were generated with KAPA Hyper Prep Kits (Roche KK8502) using 14 cycles of PCR amplification. Libraries were validated on a Tapestation 4200 (Agilent G2991BA), quantified by Qubit High Sensitivity dsDNA Kit (Life Technologies Q32854), multiplexed and sequenced on a NovaSeq6000 using 2x 100 bp cycles.

### ChIP-seq and CUT&RUN Analysis

The data quality was checked using fastq (https://www.bioinformatics.babraham.ac.uk/projects/fastqc/). The raw reads were trimmed from adapters with cutadapt (parameters: -a AGATCGGAAGAGCACACGTCTGAACTCCAGTCA-AAGATCGGAAGAGCGTCGTGTAGGGAAAGAGTGT) and raw reads were aligned to hg38 human genome assembly using bowtie2 (parameters: --local--maxins 1000). Low quality reads, duplicated reads and reads with multiple alignments were removed using samtools^120^ and Picard (https://broadinstitute.github.io/picard/). macs2^121^ was used to map position of peaks with FDR cutoff of 0.05. Bedtools^122^ was used to find a consensus set of peaks by merging peaks across multiple conditions (bedtools merge), count number of reads in peaks (bedtools intersect -c) and generate genome coverage (bedtools genomecov -bga). deepTools^123^ was used to generate coverage densities across multiple experimental conditions (deeptools computeMatrix and deeptools plotProfile) and to generate bigwig files (deeptools bamCoverage), where reads mapping to ENCODE blacklist regions were excluded^124^. All browser tracks and metaprofiles shown were calculated with sequence-depth-normalized, replicate-merged (bigWigMerge^125^) and, for ChIP-seq, input-subtracted data (deeptools bigWigCompare). Tracks were plotted using trackplot^126^. Peak differential analysis and PCA analysis was performed using DESeq2^113^ using apeglm^114^ to shrink fold changes across relevant peaksets. Overlap analyses were performed using valr^127^. Enhancers were annotated using ROSE^11^ by stitching together H3K27ac peaks in untreated cells within 12.5 kb but excluding regions within 2 kb of a transcription start site unless within a larger H3K27ac domain. For analyses of differential changes at enhancers, only read counts across annotated enhancers were considered in the peakset input to DESeq2. Peaks and enhancers were assigned first to their nearest gene by simple linear distance and manually annotated as needed from literature.

### Predicting KAT-TCIP Linker Length

LigPrep (Shrodinger Maestro 13.8) was performed on all KAT-TCIPs. The following settings were used: Force Field OPLS4, Ionization (Generate possible states at target pH: 7.00 ± 2.00), Epik Classic, Desalt, Generate Tautomers, Retain Specific Chiralities, Generate at Most 32 poses per ligand. The distance between the carbons within the amide bonds of each linker was measured. The average of the two or three lowest energy conformations was recorded.

**Chemical Synthesis:** See **Supplemental File 1**, Chemical Synthesis and Characterization.

