## Supplemental File 1 for "A Bivalent Molecular Glue Linking Lysine Acetyltransferases to Oncogene-induced Cell Death"

$^{1}H$ NMR of RCS-02-124…………………………………………………………………………….40
$^{1}H$ NMR of CCT373566a……………………………………………………………………….....40
References………………………………………………………………………………………….…41

**General synthetic methods:**

Unless otherwise noted, all reagents were purchased from commercial suppliers and used without further purification. Reactions were monitored using a Waters Acquity UPLC/MS system (Waters PDA eλ Detector, QDa Detector, Sample manager - FL, Binary Solvent Manager) using Acquity UPLC® BEH C18 column (2.1 x 50 mm, 1.7 µm particle size): solvent gradient = 85% A at 0 min, 1% A at 1.7 min; solvent A = 0.1% formic acid in water; solvent B = 0.1% formic acid in Acetonitrile; flow rate: 0.6 mL/min. Purification of reaction products was carried out using a gradient of 10-90% methanol in water containing 0.05% trifluoroacetic acid (TFA) over 40 min (45 min run time) at a flow of 40 mL/min. Assayed compounds were isolated and tested as TFA salts and purities of assayed compounds were in all cases greater than 95%, as determined by reverse-phase UPLC analysis. NMR spectra were acquired on a 500 MHz Bruker Avance III spectrometer, operating at the denoted spectrometer frequency given in MHz for the specified nucleus. All experiments were acquired at 298.0 K with a calibrated Bruker Variable Temperature Controller unless otherwise noted. The chemical shifts are reported in parts per million (ppm) and coupling constants (*J*) are given in Hertz (Hz). $^{1}$H NMR spectra are reported with the solvent resonance as the reference unless noted otherwise ($CDCl_3$ at 7.26 ppm, $CD_3OD$ at 3.31 ppm, DMSO-$d_6$ at 2.50 ppm). Peaks are reported as (s = singlet, d = doublet, t = triplet, q = quartet, m = multiplet or unresolved, br = broad signal, coupling constant(s) in Hz, integration). TEA = triethylamine; DIPEA = *N*,*N*-Diisopropylethylamine; HATU = Hexafluorophosphate Azabenzotriazole Tetramethyl Uronium; PyBOP = (Benzotriazol-1-yloxy)tripyrrolidinophosphonium hexafluorophosphate; EtOAc = ethyl acetate; THF = Tetrahydrofuran; DMF = N,N-Dimethylformamide; DMSO = Dimethyl sulfoxide; MeCN = acetonitrile; and TFA = trifluoroacetic acid.

**Synthesis of Key Intermediates**:

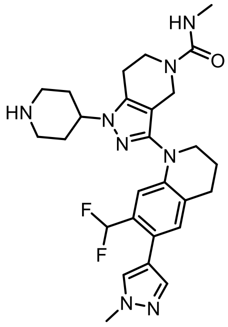

**GNE-781 - NH**

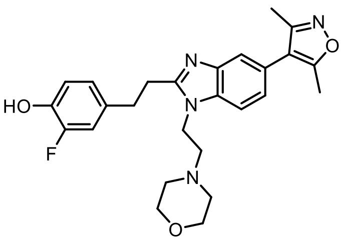

**Cmpd-33**

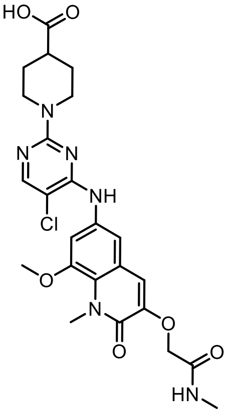

**BI-3812-COOH**

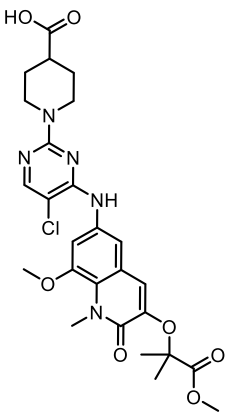

**BI-3812-Methyl Ester**

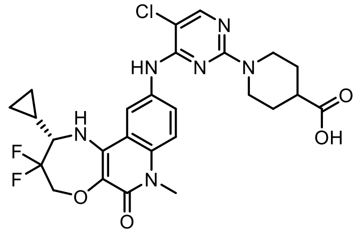

**CCT373566-COOH**

**BI-3812-COOH**[1], **BI-3812-Methyl-Ester**[1] and **CCT373566-COOH**[2] were synthesized according to previous literature procedures[1].

*Synthesis of Cmpd-33:*

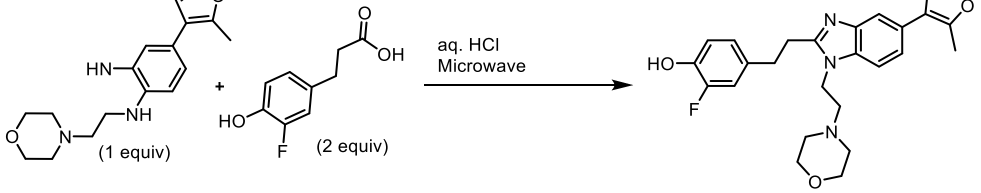

As previously prepared[3], a mixture of 4-(3,5-dimethylisoxazol-4-yl)-N1-(2-morpholinoethyl)benzene-1,2-diamine (0.23 g, 720 µmol, 1.0 equiv) and 3-(3-fluoro-4-hydroxyphenyl)propanoic acid (0.27 g, 1400 µmol, 2.0 equiv) was suspended in aqueous concentrated HCl (4.5 mL, 0.16 M) and heated in a microwave reactor for 15 min at 210 °C. The acid was quenched by careful addition of aq. sat. $NaHCO_3$ and the aqueous layer was extracted with EtOAc. The combined organic extracts were dried over $MgSO_4$ and concentrated under reduced pressure. Purification by preparative reverse-phase HPLC (0-60% MeOH in water) afforded 4-(2-(5-(3,5-dimethylisoxazol-4-yl)-1-(2-morpholinoethyl)-1H-benzo[d]imidazol-2-yl)ethyl)-2-fluorophenol as an oily solid (0.16 g, 34 µmol, 47%, MS m/z 465.40 $[M+H]^+$). **$^1$H NMR** (500 MHz, MeOD) δ = 8.00 – 7.96 (m, 1H), 7.72 (d, *J* = 1.5 Hz, 1H), 7.58 (dd, *J* = 8.6, 1.5 Hz, 1H), 7.02 (dd, *J* = 12.2, 1.8 Hz, 1H), 6.89 – 6.82 (m, 2H), 4.79 – 4.74 (m, 2H), 3.88 (d, *J* = 5.9 Hz, 4H), 3.53 (t, *J* = 7.6 Hz, 2H), 3.30 – 3.25 (m, 2H), 3.22 – 3.14 (m, 6H), 2.44 (s, 3H), 2.28 (s, 3H). **$^{19}$F NMR** (471 MHz, MeOD) δ = -77.3.

*Synthesis of GNE-781-NH:*

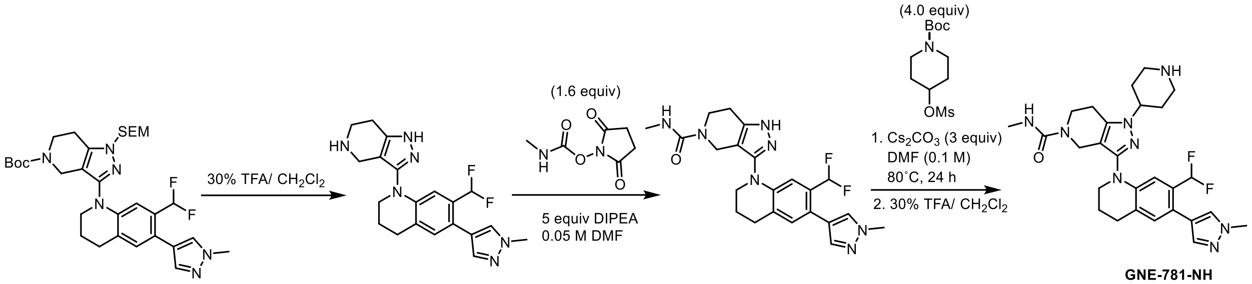

Tert-butyl 3-(7-(difluoromethyl)-6-(1-methyl-1H-pyrazol-4-yl)-3,4-dihydroquinolin-1(2H)-yl)-1-((2-(trimethylsilyl)ethoxy)methyl)-1,4,6,7-tetrahydro-5H-pyrazolo[4,3-c]pyridine-5-carboxylate[4] was dissolved in 30% TFA in $CH_2Cl_2$ (2.1 mL, 0.1 M) and stirred for 4 hours, at which point the reaction mixture was concentrated under reduced pressure. Residual TFA was removed by co-evaporation with DCM (3x), followed by high vacuum overnight to afford crude 7-(difluoromethyl)-6-(1-methyl-1H-pyrazol-4-yl)-1-(4,5,6,7-tetrahydro-1H-pyrazolo[4,3-c]pyridin-3-yl)-1,2,3,4-tetrahydroquinoline (MS m/z 385.23 $[M+H]^+$).

To a mixture of 7-(difluoromethyl)-6-(1-methyl-1H-pyrazol-4-yl)-1-(4,5,6,7-tetrahydro-1H-pyrazolo[4,3-c]pyridin-3-yl)-1,2,3,4-tetrahydroquinoline (40.0 mg, 10 µmol, 1.0 equiv) and TEA (73 µL, 52 µmol, 5 equiv) dissolved in DMF (2.1 mL, 0.05 M) at 0˚C was added 2,5-dioxopyrrolidin-1-yl methylcarbamate (29.0 mg, 170 µmol, 1.6 equiv). The reaction was allowed to stir for 4 hours, then was purified by reverse-phase HPLC (10-100% MeCN in water), concentrated under reduced pressure, and dried through lyophilization to afford 3-(7-(difluoromethyl)-6-(1-methyl-1H-pyrazol-4-yl)-3,4-dihydroquinolin-1(2H)-yl)-N-methyl-1,4,6,7-tetrahydro-5H-pyrazolo[4,3-c]pyridine-5-carboxamide (28 mg, 64 µmol, 62%, MS m/z 442.20 $[M+H]^+$).

A solution of 3-(7-(difluoromethyl)-6-(1-methyl-1H-pyrazol-4-yl)-3,4-dihydroquinolin-1(2H)-yl)-N-methyl-1,4,6,7-tetrahydro-5H-pyrazolo[4,3-c]pyridine-5-carboxamide (78.0 mg, 180 µmol, 1.0 eq), cesium carbonate (170 mg, 530 µmol, 3.0 equiv), tert-butyl 4-((methylsulfonyl)oxy)piperidine-1-carboxylate (200 mg, 710 µmol, 4.0 equiv) in DMF (1.8 mL, 0.1 M) was heated to 80˚C for 24 hours, after which point the reaction was purified by reverse-phase HPLC (10-100% MeCN in water). The second of two major peaks was collected and lyophilized to afford tert-butyl 4-(3-(7-(difluoromethyl)-6-(1-methyl-1H-pyrazol-4-yl)-3,4-dihydroquinolin-1(2H)-yl)-5-(methylcarbamoyl)-4,5,6,7-tetrahydro-1H-pyrazolo[4,3-c]pyridin-1-yl)piperidine-1-carboxylate (32 mg, 51 µmol, 29%, MS m/z 625.34 $[M+H]^+$). The resulting material was then stirred in 30% TFA v/v in $CH_2Cl_2$ (5.1 mL, 0.1 M) for 2 hours, at which point residual TFA was removed by co-evaporation with DCM (3x), followed by high vacuum overnight to afford crude 3-(7-(difluoromethyl)-6-(1-methyl-1H-pyrazol-4-yl)-3,4-dihydroquinolin-1(2H)-yl)-N-methyl-1-(piperidin-4-yl)-1,4,6,7-tetrahydro-5H-pyrazolo[4,3-c]pyridine-5-carboxamide (MS m/z 525.27 $[M+H]^{+)}$. **$^1$H NMR** (500 MHz, DMSO-$d_6$) δ = 7.75 (d, *J* = 0.8 Hz, 1H), 7.49 (d, *J* = 0.8 Hz, 1H), 7.09 (s, 1H), 6.79 (s, 1H), 6.78 (t, *J* = 55.0 Hz, 1H), 6.53 (t, *J* = 4.4 Hz, 1H), 4.12 – 4.03 (m, 1H), 4.01 (s, 2H), 3.86 (s, 3H), 3.62 – 3.54 (m, 4H), 3.02 (dt, *J* = 12.5, 3.2 Hz, 2H), 2.84 (t, *J* = 6.4 Hz, 2H), 2.71 (t, *J* = 5.7 Hz, 2H), 2.58 (td, *J* = 12.4, 2.8 Hz, 2H), 2.53 (d, *J* = 4.3 Hz, 3H), 2.02 – 1.93 (m, 2H), 1.89 – 1.82 (m, 2H), 1.78 (td, *J* = 11.1, 4.0 Hz, 2H). **$^{19}$F NMR** (471 MHz, MeOD) δ = -108.18.

### General Procedure A

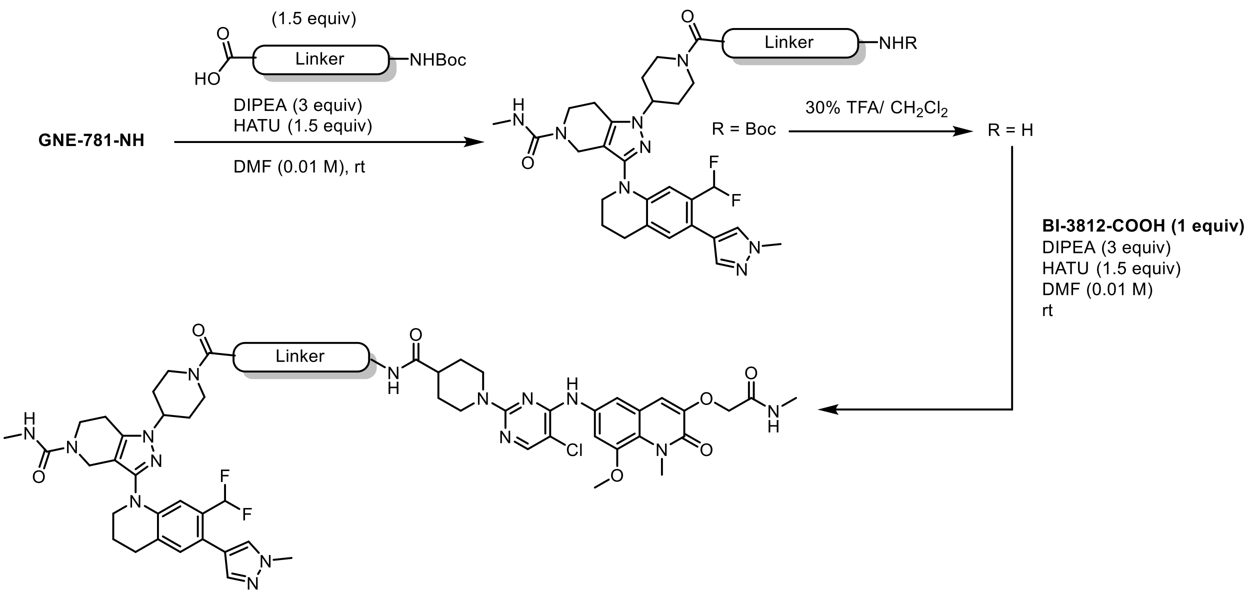

3-(7-(difluoromethyl)-6-(1-methyl-1H-pyrazol-4-yl)-3,4-dihydroquinolin-1(2H)-yl)-N-methyl-1-(piperidin-4-yl)-1,4,6,7-tetrahydro-5H-pyrazolo[4,3-c]pyridine-5-carboxamide (**GNE-781-NH$_2$**, 1.0 equiv) was added to a solution of linker (1.5 equiv) in DMF (0.1 M) and DIPEA (3.0 equiv), and the reaction was allowed to stir at room temperature until LC-MS indicated full conversion of limited starting material. The reaction mixture was purified by reverse phase chromatography (5-100% MeCN in water), concentrated under reduced pressure, then allowed to stir in 30% v/v TFA in $CH_2Cl_2$ (0.1 M) for 1 hour, or until LC-MS indicated full deprotection. The reaction was concentrated under reduced pressure, and residual TFA was removed by co-evaporation with DCM (3x), followed by high vacuum overnight. The residue was then dissolved in a minimal amount of DMF and added dropwise to a mixture of **BI-3812-COOH** (1 equiv) dissolved in DMF (0.01 M), HATU (1.5 equiv), and DIPEA (3 equiv) on ice. This reaction was allowed to stir until LC-MS indicated full consumption of limited starting material, at which point it was purified by reverse-phase HPLC (5-100% MeCN in water), concentrated under reduced pressure, and dried through lyophilization.

**Synthesis of TCIP3**:

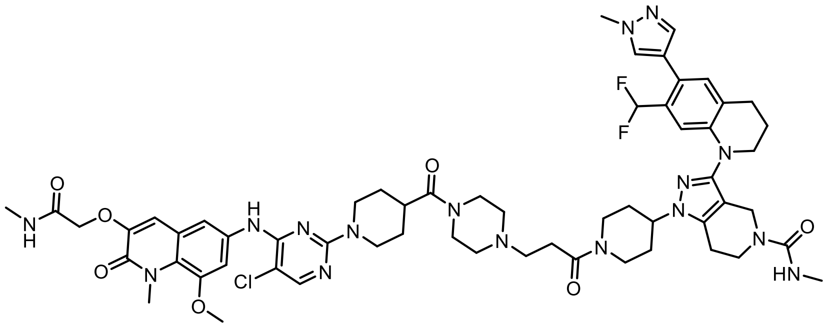

Compound MNN-03-038 was synthesized according to Procedure A using the linker 3-(4-(tert-butoxycarbonyl)piperazin-1-yl)propanoic acid to afford 1-(1-(3-(4-(1-(5-chloro-4-((8-methoxy-1-methyl-3-(2-(methylamino)-2-oxoethoxy)-2-oxo-1,2-dihydroquinolin-6-yl)amino)pyrimidin-2-yl)piperidine-4-carbonyl)piperazin-1-yl)propanoyl)piperidin-4-yl)-3-(7-(difluoromethyl)-6-(1-methyl-1H-pyrazol-4-yl)-3,4-dihydroquinolin-1(2H)-yl)-N-methyl-1,4,6,7-tetrahydro-5H-pyrazolo[4,3-c]pyridine-5-carboxamide (3.3 mg, 2.8 µmol, 30%, MS m/z 1177.74 $[M+H]^+$). **$^1$H NMR** (500 MHz, DMSO-$d_6$) δ = 9.57 (s, 1H), 8.94 (s, 1H), 8.09 (s, 1H), 7.98 (d, *J* = 4.7 Hz, 1H), 7.74 (s, 1H), 7.58 (d, *J* = 2.2 Hz, 1H), 7.50 (d, *J* = 2.3 Hz, 1H), 7.49 (d, *J* = 0.8 Hz, 1H), 7.09 (s, 1H), 6.99 (s, 1H), 6.77 (t, *J* = 55.2 Hz, 1H), 6.76 (s, 1H), 6.55 (s, 1H), 4.55 (s, 2H), 4.48 (m, *J* = 15.8 Hz, 4H), 4.41 – 4.32 (m, 1H), 4.23 (s, 1H), 4.02 (s, 2H), 3.95 (d, *J* = 13.1 Hz, 1H), 3.87 (s, 3H), 3.86 (s, 6H), 3.80-3.50 (7H, as assigned by HSQC), 3.36 (s, 2H), 3.26 – 3.17 (m, 1H), 3.10 (s, 1H), 3.04 – 2.94 (m, 5H), 2.91 (q, *J* = 6.8 Hz, 2H), 2.83 (q, *J* = 8.5 Hz, 3H), 2.76 (q, *J* = 8.1 Hz, 2H), 2.64 (d, *J* = 4.7 Hz, 3H), 2.54 (s, 3H), 1.94 (m, *J* = 13.2, 4.8 Hz, 5H), 1.84 – 1.63 (m, 3H), 1.50 (s, 2H). **$^{19}$F NMR** (471 MHz, DMSO-$d_6$) δ = -108.17.

**Synthesis of MNN-02-155**:

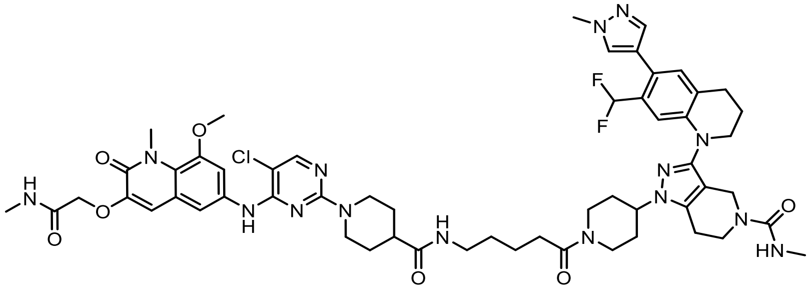

Compound MNN-02-155 was synthesized according to Procedure A using the linker 5-((tert-butoxycarbonyl)amino)pentanoic acid to afford 1-(1-(5-(1-(5-chloro-4-((8-methoxy-1-methyl-3-(2-(methylamino)-2-oxoethoxy)-2-oxo-1,2-dihydroquinolin-6-yl)amino)pyrimidin-2-yl)piperidine-4-carboxamido)pentanoyl)piperidin-4-yl)-3-(7-(difluoromethyl)-6-(1-methyl-1H-pyrazol-4-yl)-3,4-dihydroquinolin-1(2H)-yl)-N-methyl-1,4,6,7-tetrahydro-5H-pyrazolo[4,3-c]pyridine-5-carboxamide as a white solid (4.3 mg, 3.8 µmol, 40%, MS m/z 1136.52 $[M+H]^+$). **$^1$H NMR** (500 MHz, DMSO-$d_6$) δ = 9.03 (s, 1H), 8.10 (d, *J* = 6.0 Hz, 1H), 7.95 (q, *J* = 4.5 Hz, 1H), 7.79 (t, *J* = 5.6 Hz, 1H), 7.73 (s, 1H), 7.51 (s, 2H), 7.48 (s, 1H), 7.08 (s, 1H), 7.01 (s, 1H), 6.78 (s, 1H), 6.77 (t, *J* = 55.3 Hz, 1H), 6.54 (s, 1H), 4.55 (s, 2H), 4.49 – 4.43 (m, 3H), 4.31 (td, *J* = 10.6, 5.4 Hz, 1H), 4.01 (s, 2H), 3.95 (d, *J* = 13.7 Hz, 1H), 3.87 (d, *J* = 3.4 Hz, 3H), 3.85 (s, 3H), 3.85 (s, 3H), 3.60 (q, *J* = 6.1 Hz, 2H), 3.56 (t, *J* = 5.6 Hz, 2H), 3.21 – 3.09 (m, 1H), 3.03 (q, *J* = 7.0 Hz, 2H), 2.95 – 2.87 (m, 2H), 2.82 (t, *J* = 6.3 Hz, 2H), 2.76 – 2.71 (m, 2H), 2.68 (d, *J* = 12.5 Hz, 1H), 2.64 (d, *J* = 4.7 Hz, 3H), 2.54 (s, 3H), 2.42 – 2.35 (m, 1H), 2.33 (t, *J* = 7.3 Hz, 2H), 1.96 (q, *J* = 5.9 Hz, 2H), 1.91 – 1.82 (m, 3H), 1.70 (d, *J* = 13.1 Hz, 3H), 1.58-1.43 (m, 4H), 1.40 (t, *J* = 7.4 Hz, 2H). **$^{19}$F NMR** (471 MHz, DMSO-$d_6$) δ = -108.21.

**Synthesis of MNN-03-037**:

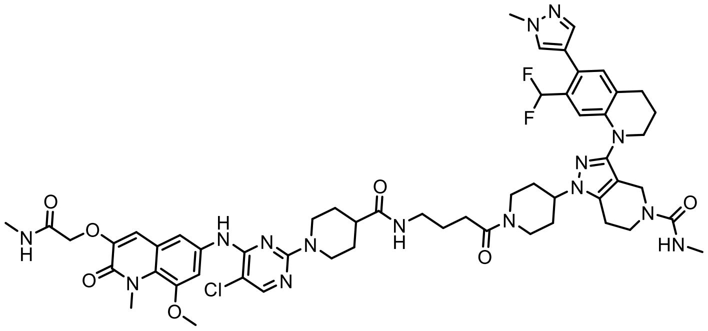

Compound MNN-03-037 was synthesized according to Procedure A using the linker 4-((tert-butoxycarbonyl)amino)butanoic acid to afford 1-(1-(4-(1-(5-chloro-4-((8-methoxy-1-methyl-3-(2-(methylamino)-2-oxoethoxy)-2-oxo-1,2-dihydroquinolin-6-yl)amino)pyrimidin-2-yl)piperidine-4-carboxamido)butanoyl)piperidin-4-yl)-3-(7-(difluoromethyl)-6-(1-methyl-1H-pyrazol-4-yl)-3,4-dihydroquinolin-1(2H)-yl)-N-methyl-1,4,6,7-tetrahydro-5H-pyrazolo[4,3-c]pyridine-5-carboxamide (1.95 mg, 1.7 µmol, 18%, MS m/z 1122.51 $[M+H]^+$). $^{1}$**H NMR** (500 MHz, DMSO-$d_6$) δ = 9.03 (s, 1H), 8.09 (d, *J* = 2.4 Hz, 1H), 7.97 – 7.91 (m, 1H), 7.81 (t, *J* = 5.7 Hz, 1H), 7.74 (d, *J* = 2.6 Hz, 1H), 7.51 (d, *J* = 2.7 Hz, 2H), 7.49 (s, 1H), 7.08 (s, 1H), 7.00 (s, 1H), 6.78 (s, 1H), 6.77 (t, *J* = 55.3 Hz, 1H), 6.54 (s, 1H), 4.55 (s, 2H), 4.46 (d, *J* = 12.9 Hz, 3H), 4.35 – 4.26 (m, 1H), 4.01 (s, 2H), 3.93 (d, *J* = 13.4 Hz, 1H), 3.87 (d, *J* = 3.1 Hz, 3H), 3.85 (s, 3H), 3.85 (s, 3H), 3.64 – 3.59 (m, 2H), 3.56 (t, *J* = 5.7 Hz, 2H), 3.14 (d, *J* = 12.3 Hz, 1H), 3.05 (q, *J* = 6.6 Hz, 2H), 2.96 – 2.87 (m, 2H), 2.81 (d, *J* = 6.5 Hz, 2H), 2.73 (d, *J* = 5.5 Hz, 2H), 2.69 (d, *J* = 12.7 Hz, 1H), 2.64 (d, *J* = 4.7 Hz, 3H), 2.54 (s, 3H), 2.42 – 2.36 (m, 1H), 2.33 (t, *J* = 7.5 Hz, 2H), 1.95 (p, *J* = 6.3 Hz, 2H), 1.87 (d, *J* = 17.1 Hz, 3H), 1.75 – 1.68 (m, 3H), 1.62 (p, *J* = 7.2 Hz, 2H), 1.50 (td, *J* = 12.2, 8.6 Hz, 2H). $^{19}$**F NMR** (471 MHz, DMSO-$d_6$) δ = -108.17.

**Synthesis of MNN-02-195**:

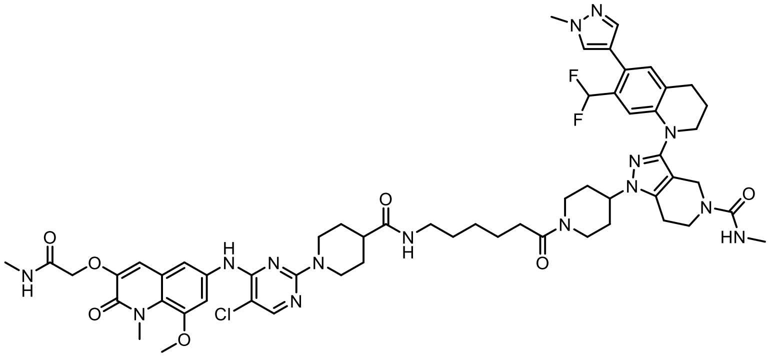

Compound MNN-02-195 was synthesized according to Procedure A using the linker 6-((tert-butoxycarbonyl)amino)hexanoic acid to afford 1-(1-(6-(1-(5-chloro-4-((8-methoxy-1-methyl-3-(2-(methylamino)-2-oxoethoxy)-2-oxo-1,2-dihydroquinolin-6-yl)amino)pyrimidin-2-yl)piperidine-4-carboxamido)hexanoyl)piperidin-4-yl)-3-(7-(difluoromethyl)-6-(1-methyl-1H-pyrazol-4-yl)-3,4-dihydroquinolin-1(2H)-yl)-N-methyl-1,4,6,7-tetrahydro-5H-pyrazolo[4,3-c]pyridine-5-carboxamide (3.5 mg, 3.1 µmol, 20%, MS m/z 1150.53 $[M+H]^+$). $^{1}$**H NMR** (500 MHz, DMSO-$d_6$) δ = 9.12 (s, 1H), 8.12 (d, *J* = 5.5 Hz, 1H), 7.95 (q, *J* = 4.5 Hz, 1H), 7.77 (t, *J* = 5.6 Hz, 1H), 7.73 (s, 1H), 7.51 (s, 2H), 7.48 (s, 1H), 7.08 (s, 1H), 7.01 (s, 1H), 6.78 (s, 1H), 6.77 (t, *J* = 55.2 Hz, 1H), 6.54 (s, 1H), 4.55 (s, 2H, assigned by HSQC), 4.44 (d, *J* = 13.2 Hz, 3H, assigned by HSQC), 4.36 – 4.26 (m, 1H, assigned by HSQC), 4.01 (s, 2H), 3.95 (d, *J* = 13.2 Hz, 1H), 3.86 (s, 3H), 3.85 (s, 3H), 3.85 (s, 3H), 3.60 (q, *J* = 5.8 Hz, 2H), 3.56 (t, *J* = 5.6 Hz, 2H), 3.15 (t, *J* = 12.0 Hz, 1H), 3.01 (q, *J* = 6.8 Hz, 2H), 2.93 (t, *J* = 12.5 Hz, 2H), 2.82 (t, *J* = 6.5 Hz, 2H), 2.73 (t, *J* = 5.9 Hz, 2H), 2.71 – 2.66

(m, 1H), 2.64 (d, $J$ = 4.6 Hz, 3H), 2.54 (s, 3H), 2.41 – 2.34 (m, 1H), 2.31 (t, $J$ = 7.5 Hz, 2H), 1.95 (t, $J$ = 5.8 Hz, 2H), 1.91 – 1.83 (m, 3H), 1.71 (dd, $J$ = 13.2, 3.7 Hz, 3H), 1.49 (dt, $J$ = 14.5, 5.6 Hz, 4H), 1.39 (p, $J$ = 7.1 Hz, 2H), 1.25 (qd, $J$ = 8.5, 5.8 Hz, 2H). **$^{19}$F NMR** (471 MHz, DMSO-$d_6$) δ = -108.19.

**Synthesis of RCS-IJD-001:**

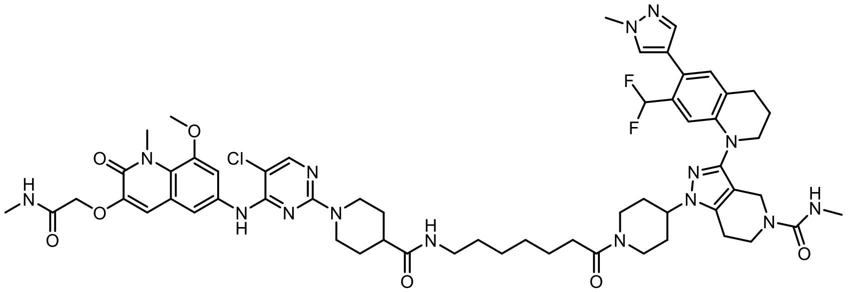

Compound RCS-SBM-IJD-001 was synthesized according to Procedures A using the linker 8-((tert-butoxycarbonyl)amino)octanoic acid to afford 1-(1-(7-(1-(5-chloro-4-((8-methoxy-1-methyl-3-(2-(methylamino)-2-oxoethoxy)-2-oxo-1,2-dihydroquinolin-6-yl)amino)pyrimidin-2-yl)piperidine-4-carboxamido)heptanoyl)piperidin-4-yl)-3-(7-(difluoromethyl)-6-(1-methyl-1H-pyrazol-4-yl)-3,4-dihydroquinolin-1(2H)-yl)-N-methyl-1,4,6,7-tetrahydro-5H-pyrazolo[4,3-c]pyridine-5-carboxamide (3.9 mg, 3.4 µmol, 18%, MS m/z 1165.48 $[M+H]^+$). **$^1$H NMR** (500 MHz, DMSO-$d_6$) δ = 9.16 (s, 1H), 8.12 (s, 1H), 7.95 (q, $J$ = 4.6 Hz, 1H), 7.76 (t, $J$ = 5.6 Hz, 1H), 7.73 (s, 1H), 7.50 (s, 2H), 7.48 (d, $J$ = 0.8 Hz, 1H), 7.08 (s, 1H), 7.01 (s, 1H), 6.79 (s, 1H), 6.77 (t, $J$ = 55.2 Hz, 1H), 6.53 (s, 1H), 4.55 (s, 2H, assigned by HSQC), 4.46 – 4.40 (3H, assigned by HSQC), 4.35 – 4.27 (m, 1H, assigned by HSQC), 4.01 (s, 2H), 3.95 (d, $J$ = 13.6 Hz, 1H), 3.86 (s, 3H), 3.86 (s, 3H), 3.85 (s, 3H), 3.60 (q, $J$ = 5.8 Hz, 2H), 3.56 (t, $J$ = 5.7 Hz, 2H), 3.15 (t, $J$ = 12.5 Hz, 1H), 3.00 (q, $J$ = 6.6 Hz, 2H), 2.97 – 2.89 (m, 2H), 2.82 (t, $J$ = 6.3 Hz, 2H), 2.73 (t, $J$ = 6.0 Hz, 2H), 2.68 (d, $J$ = 12.1 Hz, 1H), 2.64 (d, $J$ = 4.6 Hz, 3H), 2.54 (s, 3H), 2.42 – 2.34 (m, 1H), 2.31 (t, $J$ = 7.5 Hz, 2H), 1.96 (p, $J$ = 6.3 Hz, 2H), 1.88 (dd, $J$ = 18.7, 8.7 Hz, 3H), 1.74 – 1.68 (m, 3H), 1.56 – 1.42 (m, 4H), 1.36 (q, $J$ = 7.0 Hz, 2H), 1.25 (d, $J$ = 5.4 Hz, 4H). **$^{19}$F NMR** (471 MHz, DMSO-$d_6$) δ = -108.19.

**Synthesis of MNN-02-196:**

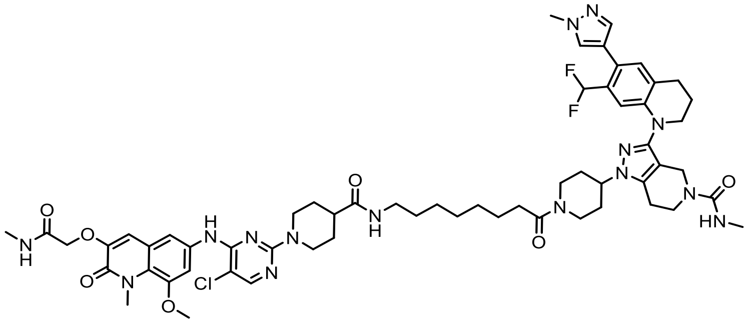

Compound MNN-02-196 was synthesized according to Procedure A using the linker 8-((tert-butoxycarbonyl)amino)octanoic acid to afford 1-(1-(8-(1-(5-chloro-4-((8-methoxy-1-methyl-3-(2-(methylamino)-2-oxoethoxy)-2-oxo-1,2-dihydroquinolin-6-yl)amino)pyrimidin-2-yl)piperidine-4-carboxamido)octanoyl)piperidin-4-yl)-3-(7-(difluoromethyl)-6-(1-methyl-1H-pyrazol-4-yl)-3,4-dihydroquinolin-1(2H)-yl)-N-methyl-1,4,6,7-tetrahydro-5H-pyrazolo[4,3-c]pyridine-5-carboxamide (1.4 mg, 1.2 µmol, 8%, MS m/z 1180.09 $[M+H]^+$). **$^1$H NMR** (500 MHz, DMSO) δ = 9.05 (s, 1H), 8.10 (s, 1H), 7.95 (q, $J$ = 4.7 Hz, 1H), 7.77 (t, $J$ = 5.6 Hz, 1H), 7.74 (s, 1H), 7.51 (s, 2H), 7.49 (s, 1H), 7.08 (s, 1H), 7.00 (s, 1H), 6.79 (s, 1H), 6.77 (t, $J$ = 55.2 Hz, 1H), 6.55 (s, 1H), 4.55 (s, 2H), 4.46 (d, $J$ = 12.6 Hz, 3H), 4.30 (td, $J$ = 10.5, 5.3 Hz, 1H), 4.01 (s, 2H), 3.95 (d, $J$ = 13.1 Hz, 1H),

3.86 (s, 3H), 3.85 (s, 3H), 3.85 (s, 3H), 3.60 (q, $J$ = 5.9 Hz, 2H), 3.56 (t, $J$ = 5.7 Hz, 2H), 3.14 (t, $J$ = 11.2 Hz, 1H), 3.00 (q, $J$ = 6.6 Hz, 2H), 2.91 (t, $J$ = 12.5 Hz, 2H), 2.82 (t, $J$ = 6.4 Hz, 2H), 2.78 – 2.71 (m, 2H), 2.69 (s, 1H), 2.64 (d, $J$ = 4.6 Hz, 3H), 2.53 (s, 3H), 2.41 – 2.34 (m, 1H), 2.30 (t, $J$ = 7.5 Hz, 2H), 1.95 (p, $J$ = 6.2 Hz, 2H), 1.92 – 1.79 (m, 3H), 1.70 (d, $J$ = 12.7 Hz, 3H), 1.49 (dq, $J$ = 13.2, 8.7 Hz, 4H), 1.35 (q, $J$ = 6.9 Hz, 2H), 1.24 (q, $J$ = 8.1 Hz, 6H). **$^{19}$F NMR** (471 MHz, DMSO-$d_6$) δ = -108.18.

**Synthesis of MNN-02-187**:

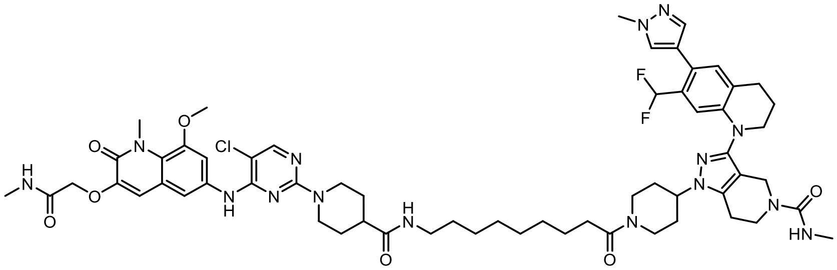

Compound MNN-02-187 was synthesized according to Procedure A using the linker 9-((tert-butoxycarbonyl)amino)nonanoic acid to afford (1-(1-(9-(1-(5-chloro-4-((8-methoxy-1-methyl-3-(2-(methylamino)-2-oxoethoxy)-2-oxo-1,2-dihydroquinolin-6-yl)amino)pyrimidin-2-yl)piperidine-4-carboxamido)nonanoyl)piperidin-4-yl)-3-(7-(difluoromethyl)-6-(1-methyl-1H-pyrazol-4-yl)-3,4-dihydroquinolin-1(2H)-yl)-N-methyl-1,4,6,7-tetrahydro-5H-pyrazolo[4,3-c]pyridine-5-carboxamide) (1.41 mg, 1.2 µmol, 8%, MS m/z 1193.80 $[M+H]^+$). **$^1$H NMR** (500 MHz, DMSO-$d_6$) δ = 8.95 (s, 1H), 8.08 (s, 1H), 7.96 (d, $J$ = 4.8 Hz, 1H), 7.77 (t, $J$ = 5.6 Hz, 1H), 7.74 (s, 1H), 7.52 (s, 2H), 7.49 (s, 1H), 7.08 (s, 1H), 7.00 (s, 1H), 6.79 (s, 1H), 6.77 (t, $J$ = 55.2 Hz, 1H), 6.55 (s, 1H), 4.55 (s, 2H), 4.47 (t, $J$ = 15.5 Hz, 3H), 4.30 (d, $J$ = 10.7 Hz, 1H), 4.01 (s, 2H), 3.94 (d, $J$ = 13.7 Hz, 1H), 3.86 (s, 3H), 3.86 (s, 3H), 3.85 (s, 3H), 3.60 (q, $J$ = 5.6 Hz, 2H), 3.56 (t, $J$ = 5.7 Hz, 2H), 3.15 (t, $J$ = 12.5 Hz, 1H), 2.99 (q, $J$ = 6.5 Hz, 2H), 2.89 (t, $J$ = 12.3 Hz, 2H), 2.81 (d, $J$ = 6.7 Hz, 2H), 2.73 (t, $J$ = 6.1 Hz, 2H), 2.68 (d, $J$ = 11.7 Hz, 1H), 2.64 (d, $J$ = 4.7 Hz, 3H), 2.56 – 2.51 (m, 3H), 2.40 – 2.33 (m, 1H), 2.30 (t, $J$ = 7.5 Hz, 2H), 1.95 (p, $J$ = 6.2 Hz, 2H), 1.87 (d, $J$ = 17.1 Hz, 3H), 1.72 – 1.65 (m, 3H), 1.54 – 1.40 (m, 4H), 1.39 – 1.32 (m, 2H), 1.23 (s, 8H). **$^{19}$F NMR** (471 MHz, DMSO-$d_6$) δ = -108.19.

**Synthesis of MNN-02-197**:

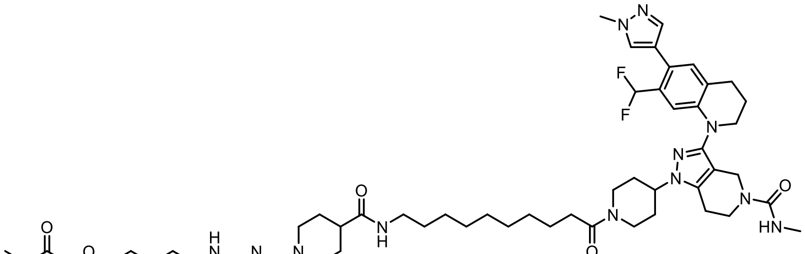

Compound MNN-02-197 was synthesized according to Procedure A using the linker 10-((tert-butoxycarbonyl)amino)decanoic acid to afford (1-(1-(10-(1-(5-chloro-4-((8-methoxy-1-methyl-3-(2-(methylamino)-2-oxoethoxy)-2-oxo-1,2-dihydroquinolin-6-yl)amino)pyrimidin-2-yl)piperidine-4-carboxamido)decanoyl)piperidin-4-yl)-3-(7-(difluoromethyl)-6-(1-methyl-1H-pyrazol-4-yl)-3,4-dihydroquinolin-1(2H)-yl)-N-methyl-1,4,6,7-tetrahydro-5H-pyrazolo[4,3-c]pyridine-5-carboxamide) (1.4 mg, 1.2 µmol, 8%, MS m/z 1207.76 $[M+H]^+$). **$^1$H NMR** (500 MHz, DMSO-$d_6$) δ = 9.10 (s, 1H), 8.11 (s, 1H), 7.95 (q, $J$ = 4.6 Hz, 1H), 7.75 (t, $J$ = 5.6 Hz, 1H), 7.74 (s, 1H), 7.51 (s, 2H), 7.48 (d, $J$ = 0.9 Hz, 1H), 7.08 (s, 1H), 7.01 (d, $J$ = 1.8 Hz, 1H), 6.79 (s, 1H), 6.77 (t, $J$ =

55.3 Hz, 1H), 6.54 (s, 1H), 4.55 (s, 2H), 4.45 (d, *J* = 12.9 Hz, 3H), 4.30 (m, *J* = 10.4, 5.3 Hz, 1H), 4.01 (s, 2H), 3.95 (d, *J* = 13.6 Hz, 1H), 3.87 (s, 3H), 3.86 (s, 3H), 3.85 (s, 3H), 3.64 – 3.58 (m, 2H), 3.56 (t, *J* = 5.7 Hz, 2H), 3.18 – 3.09 (m, 1H), 2.99 (q, *J* = 6.6 Hz, 2H), 2.97 – 2.89 (m, 2H), 2.82 (t, *J* = 6.4 Hz, 2H), 2.73 (t, *J* = 5.9 Hz, 2H), 2.68 (d, *J* = 12.0 Hz, 1H), 2.64 (d, *J* = 4.7 Hz, 3H), 2.54 (d, *J* = 1.2 Hz, 3H), 2.43 – 2.35 (m, 1H), 2.33 – 2.27 (m, 2H), 1.96 (p, *J* = 6.3 Hz, 2H), 1.91 – 1.80 (m, 3H), 1.71 (d, *J* = 13.3 Hz, 3H), 1.56 – 1.43 (m, 4H), 1.35 (t, *J* = 6.8 Hz, 2H), 1.29 – 1.20 (m, 10H). **$^{19}$F NMR** (471 MHz, DMSO-$d_6$) δ = -108.18.

**Synthesis of MNN-02-160**:

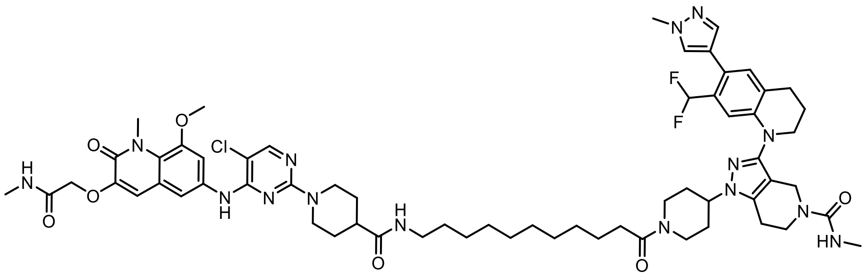

Compound MNN-02-160 was synthesized according to Procedure A to afford (1-(1-(11-(1-(5-chloro-4-((8-methoxy-1-methyl-3-(2-(methylamino)-2-oxoethoxy)-2-oxo-1,2-dihydroquinolin-6-yl)amino)pyrimidin-2-yl)piperidine-4-carboxamido)undecanoyl)piperidin-4-yl)-3-(7-(difluoromethyl)-6-(1-methyl-1H-pyrazol-4-yl)-3,4-dihydroquinolin-1(2H)-yl)-N-methyl-1,4,6,7-tetrahydro-5H-pyrazolo[4,3-c]pyridine-5-carboxamide) (0.89 mg, 0.73 µmol, 7%, MS m/z 1221.82 [M+H]$^+$). **$^1$H NMR** (500 MHz, DMSO-$d_6$) δ = 9.07 (s, 1H), 8.10 (s, 1H), 7.96 (q, *J* = 4.8 Hz, 1H), 7.77 (t, *J* = 5.6 Hz, 1H), 7.74 (d, *J* = 0.9 Hz, 1H), 7.52 (s, 2H), 7.48 (d, *J* = 0.8 Hz, 1H), 7.08 (s, 1H), 7.01 (s, 1H), 6.79 (s, 1H), 6.76 (t, *J* = 55.3 Hz, 1H), 6.55 (s, 1H), 4.55 (s, 2H), 4.47 (m, *J* = 13.0 Hz, 3H), 4.31 (tt, *J* = 10.6, 4.5 Hz, 1H), 4.01 (s, 2H), 3.98 – 3.91 (m, 1H), 3.87 (s, 3H), 3.86 (s, 3H), 3.85 (s, 3H), , 3.60 (q, *J* = 5.4 Hz, 2H), 3.56 (t, *J* = 5.7 Hz, 2H), 3.15 (t, *J* = 12.4 Hz, 1H), 2.99 (q, *J* = 6.6 Hz, 2H), 2.97 – 2.88 (m, 2H), 2.82 (t, *J* = 6.5 Hz, 2H), 2.73 (t, *J* = 6.0 Hz, 2H), 2.68 (d, *J* = 11.7 Hz, 1H), 2.64 (d, *J* = 4.6 Hz, 3H), 2.54 (d, *J* = 1.7 Hz, 3H), 2.42 – 2.34 (m, 1H), 2.33 – 2.27 (m, 2H), 1.96 (dt, *J* = 10.0, 6.3 Hz, 2H), 1.91 – 1.83 (m, 3H), 1.71 (d, *J* = 13.2 Hz, 3H), 1.49 (dq, *J* = 22.0, 8.1 Hz, 4H), 1.38 – 1.32 (m, 2H), 1.27 – 1.15 (m, 12H). **$^{19}$F NMR** (471 MHz, DMSO-$d_6$) δ = -108.16.

**Synthesis of MNN-02-161**:

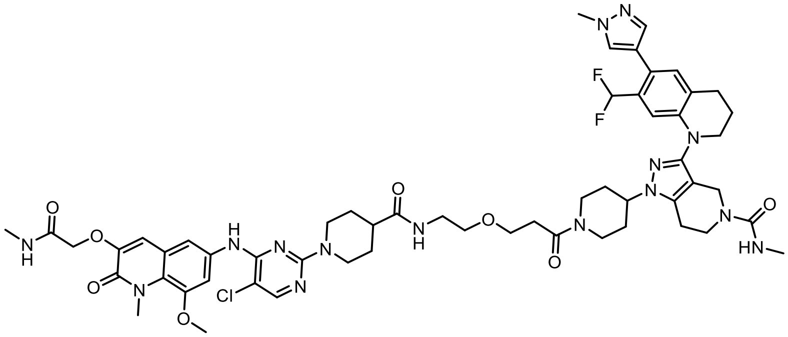

Compound MNN-02-161 was synthesized according to Procedure A using the linker 3-(2-((tert-butoxycarbonyl)amino)ethoxy)propanoic acid to afford 1-(1-(3-(2-(1-(5-chloro-4-((8-methoxy-1-methyl-3-(2-(methylamino)-2-oxoethoxy)-2-oxo-1,2-dihydroquinolin-6-yl)amino)pyrimidin-2-yl)piperidine-4-carboxamido)ethoxy)propanoyl)piperidin-4-yl)-3-(7-(difluoromethyl)-6-(1-methyl-1H-pyrazol-4-yl)-3,4-dihydroquinolin-1(2H)-yl)-N-methyl-1,4,6,7-tetrahydro-5H-pyrazolo[4,3-c]pyridine-5-carboxamide (2.5 mg, 2.2 µmol, 20%, MS m/z 1152.48 [M+H]$^+$). **$^1$H NMR** (500 MHz,

DMSO-$d_6$) δ = 8.83 (s, 1H), 8.05 (s, 1H), 7.96 (q, *J* = 4.6 Hz, 1H), 7.84 (t, *J* = 5.6 Hz, 1H), 7.73 (s, 1H), 7.53 (q, *J* = 2.3 Hz, 2H), 7.48 (s, 1H), 7.08 (s, 1H), 6.99 (s, 1H), 6.76 (t, *J* = 55.2 Hz, 1H), 6.76 (s, 1H, as determined by HSQC), 6.54 (d, *J* = 4.7 Hz, 1H), 4.54 (s, 2H), 4.47 (t, *J* = 18.1 Hz, 3H), 4.31 (d, *J* = 10.2 Hz, 1H), 3.98 (d, *J* = 18.1 Hz, 2H), 3.96 (s, 1H), 3.86 (s, 3H), 3.85 (s, 3H), 3.84 (s, 3H). 3.64 – 3.52 (m, 6H, determined by HSQC), 3.38 (t, *J* = 5.9 Hz, 2H), 3.17 (dt, *J* = 5.9, 2.8 Hz, 3H), 2.87 (s, 1H), 2.81 (d, 2H), 2.73 (t, *J* = 5.8 Hz, 2H), 2.72 – 2.68 (m, 1H), 2.64 (d, *J* = 4.7 Hz, 3H), 2.59 (dt, *J* = 12.3, 6.6 Hz, 1H), 2.54 (s, 3H), 2.44 – 2.36 (m, 1H), 1.95 (p, *J* = 6.3 Hz, 2H), 1.87 (d, *J* = 22.9 Hz, 3H), 1.69 (d, *J* = 9.7 Hz, 3H), 1.53 – 1.43 (m, 2H), 1.29 – 1.21 (m, 2H). **$^{19}$F NMR** (471 MHz, DMSO-$d_6$) δ = -108.16.

**Synthesis of MNN-02-162**:

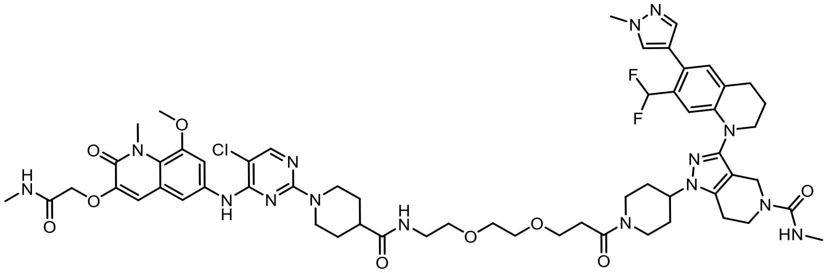

Compound MNN-02-162 was synthesized according to Procedure A using the linker 2,2-dimethyl-4-oxo-3,8,11-trioxa-5-azatetradecan-14-oic acid to afford 1-(1-(3-(2-(2-(1-(5-chloro-4-((8-methoxy-1-methyl-3-(2-(methylamino)-2-oxoethoxy)-2-oxo-1,2-dihydroquinolin-6-yl)amino)pyrimidin-2-yl)piperidine-4-carboxamido)ethoxy)ethoxy)propanoyl)piperidin-4-yl)-3-(7-(difluoromethyl)-6-(1-methyl-1H-pyrazol-4-yl)-3,4-dihydroquinolin-1(2H)-yl)-N-methyl-1,4,6,7-tetrahydro-5H-pyrazolo[4,3-c]pyridine-5-carboxamide (2.9 mg, 2.3 µmol, 20%, MS m/z 1196.40 $[M+H]^+$). **$^1$H NMR** (500 MHz, DMSO-$d_6$) δ = 8.85 (s, 1H), 8.06 (s, 1H), 7.95 (q, *J* = 4.6 Hz, 1H), 7.84 (t, *J* = 5.8 Hz, 1H), 7.73 (s, 1H), 7.53 (s, 2H), 7.48 (s, 1H), 7.08 (s, 1H), 7.00 (s, 1H), 6.77 (t, 1H), 6.77 (t, *J* = 55.2 Hz, 1H), 6.53 (d, *J* = 5.1 Hz, 1H), 4.54 (s, 2H), 4.47 (dd, *J* = 26.3, 12.9 Hz, 3H), 4.35 – 4.27 (m, 1H), 4.00 (d, *J* = 5.1 Hz, 2H), 3.96 (d, *J* = 5.9 Hz, 1H), 3.86 (s, 3H), 3.85 (s, 3H), 3.85 (s, 3H), 3.61 (td, *J* = 6.8, 2.2 Hz, 3H, determined by HSQC), 3.56 (q, *J* = 6.9 Hz, 3H, determined by HSQC). 3.48 (s, 4H), 3.37 (t, *J* = 5.9 Hz, 2H), 3.16 (q, *J* = 6.0 Hz, 3H), 2.88 (td, *J* = 13.0, 2.7 Hz, 2H), 2.82 (t, *J* = 6.4 Hz, 2H), 2.73 (q, *J* = 5.7 Hz, 2H), 2.64 (d, *J* = 4.7 Hz, 3H), 2.59 (q, *J* = 6.5 Hz, 1H), 2.54 (s, 3H), 2.40 (dh, *J* = 11.4, 3.7 Hz, 1H), 1.99 – 1.92 (m, 2H), 1.87 (d, *J* = 18.6 Hz, 3H), 1.77 – 1.66 (m, 3H), 1.48 (qd, *J* = 12.5, 4.1 Hz, 2H), 1.29 – 1.21 (m, 2H). **$^{19}$F NMR** (471 MHz, DMSO-$d_6$) δ = -108.17.

**Synthesis of MNN-02-156**:

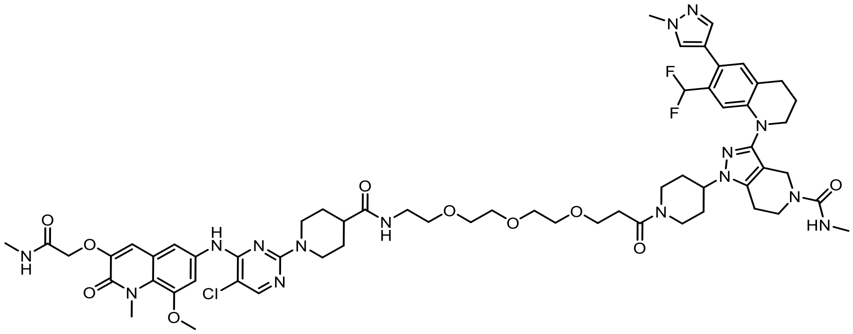

Compound MNN-02-156 was synthesized according to Procedure A using the linker 2,2-dimethyl-4-oxo-3,8,11,14-tetraoxa-5-azaheptadecan-17-oic acid to afford 1-(1-(1-(1-(5-chloro-4-((8-methoxy-1-methyl-3-(2-(methylamino)-2-oxoethoxy)-2-oxo-1,2-dihydroquinolin-6-yl)amino)pyrimidin-2-yl)piperidin-4-yl)-1-oxo-5,8,11-trioxa-2-azatetradecan-14-oyl)piperidin-4-yl)-

3-(7-(difluoromethyl)-6-(1-methyl-1H-pyrazol-4-yl)-3,4-dihydroquinolin-1(2H)-yl)-N-methyl-1,4,6,7-tetrahydro-5H-pyrazolo[4,3-c]pyridine-5-carboxamide. (2.0 mg, 1.6 µmol, 20%, MS m/z 1241.04 [M+H]$^+$). **$^1$H NMR** (500 MHz, DMSO-d$_6$) δ = 8.83 (s, 1H), 8.06 (d, *J* = 1.4 Hz, 1H), 7.96 (d, *J* = 4.9 Hz, 1H), 7.85 (q, *J* = 6.2 Hz, 1H), 7.74 (d, *J* = 1.7 Hz, 1H), 7.53 (s, 2H), 7.48 (s, 1H), 7.08 (s, 1H), 7.00 (s, 1H), 6.77 (t, *J* = 55.3 Hz, 1H), 6.77 (s, 1H, as determined by HSQC), 6.53 (d, *J* = 4.7 Hz, 1H), 4.54 (s, 2H), 4.47 (dd, *J* = 27.5, 12.8 Hz, 3H), 4.31 (s, 1H), 4.00 (s, 2H), 3.97 (s, 1H), 3.87 (s, 3H), 3.86 (s, 3H), 3.85 (s, 3H), 3.61 (dt, *J* = 6.6, 3.4 Hz, 3H, as determined by HSQC), 3.56 (dd, *J* = 7.0, 5.0 Hz, 3H, as determined by HSQC), 3.49 – 3.47 (m, 6H, as determined by HSQC), 3.47 – 3.44 (m, 2H, as determined by HSQC), 3.36 (t, *J* = 6.0 Hz, 2H), 3.17 (p, *J* = 5.8 Hz, 3H), 2.88 (td, *J* = 12.8, 2.7 Hz, 1H), 2.82 (t, *J* = 6.1 Hz, 2H), 2.76 – 2.72 (m, 2H), 2.70 (d, *J* = 12.8 Hz, 1H), 2.65 (d, *J* = 4.7 Hz, 3H), 2.62 – 2.56 (m, 1H), 2.53 (d, *J* = 3.3 Hz, 3H), 2.45 – 2.33 (m, 1H), 1.99 – 1.92 (m, 2H), 1.91 – 1.84 (m, 3H), 1.75 – 1.66 (m, 3H), 1.54 – 1.43 (m, 2H), 1.28 – 1.16 (m, 2H). **$^{19}$F NMR** (471 MHz, DMSO-d$_6$) δ = -108.16.

**Synthesis of MNN-03-050**:

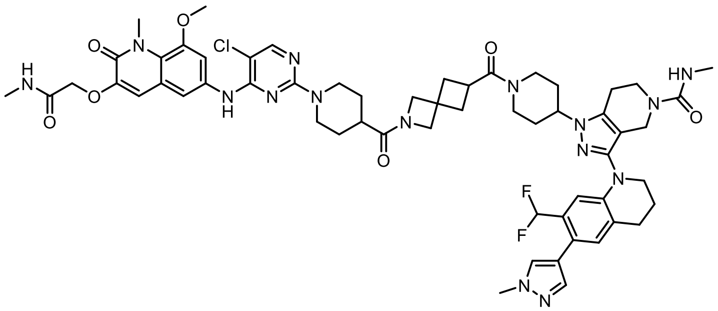

Compound MNN-03-050 was synthesized according to Procedure A using the linker 2-(tert-butoxycarbonyl)-2-azaspiro[3.3]heptane-6-carboxylic acid to afford 1-(1-(2-(1-(5-chloro-4-((8-methoxy-1-methyl-3-(2-(methylamino)-2-oxoethoxy)-2-oxo-1,2-dihydroquinolin-6-yl)amino)pyrimidin-2-yl)piperidine-4-carbonyl)-2-azaspiro[3.3]heptane-6-carbonyl)piperidin-4-yl)-3-(7-(difluoromethyl)-6-(1-methyl-1H-pyrazol-4-yl)-3,4-dihydroquinolin-1(2H)-yl)-N-methyl-1,4,6,7-tetrahydro-5H-pyrazolo[4,3-c]pyridine-5-carboxamide (4.7 mg, 4.1 µmol, 40%, MS m/z 1160.73 [M+H]$^+$). **$^1$H NMR** (500 MHz, DMSO-d$_6$) δ = 9.17 (s, 1H), 8.12 (s, 1H), 7.94 (q, *J* = 4.7 Hz, 1H), 7.74 (s, 1H), 7.52 (dd, *J* = 7.1, 2.3 Hz, 1H), 7.48 (d, *J* = 2.6 Hz, 2H), 7.09 (s, 1H), 7.00 (s, 1H), 6.81 (s, 1H), 6.77 (t, *J* = 55.1 Hz, 1H), 6.54 (s, 1H), 4.55 (s, 2H), 4.42 (d, *J* = 12.8 Hz, 3H), 4.32 (dq, *J* = 11.2, 5.3 Hz, 1H), 4.24 (s, 1H), 4.08 (s, 1H), 4.02 (s, 2H), 3.86 (d, *J* = 1.3 Hz, 3H), 3.86 (s, 3H), 3.85 (s, 3H), 3.80 (d, *J* = 13.9 Hz, 1H), 3.70 (s, 1H), 3.59 (d, *J* = 7.0 Hz, 2H), 3.56 (t, *J* = 5.6 Hz, 2H), 3.26 (td, *J* = 8.4, 2.7 Hz, 2H), 3.11 (t, 2H), 2.97 (s, 2H), 2.83 (t, *J* = 6.4 Hz, 2H), 2.73 (h, *J* = 6.2 Hz, 2H), 2.64 (d, *J* = 4.7 Hz, 3H), 2.54 (s, 3H), 2.34 (td, *J* = 8.1, 4.0 Hz, 4H), 2.25 (d, *J* = 8.2 Hz, 1H), 1.97 (p, *J* = 6.1 Hz, 2H), 1.87 (d, *J* = 13.2 Hz, 2H), 1.81 (s, 1H), 1.69 (d, *J* = 31.1 Hz, 3H), 1.44 (t, *J* = 12.0 Hz, 2H). **$^{19}$F NMR** (471 MHz, DMSO-d$_6$) δ = -108.19.

**Synthesis of MNN-03-041**:

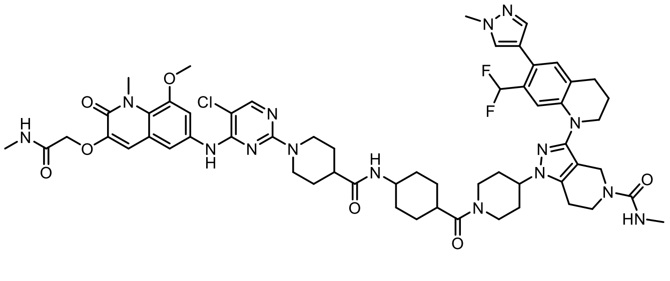

Compound MNN-03-041 was synthesized according to Procedure A using the linker 4-((tert-butoxycarbonyl)amino)cyclohexane-1-carboxylic acid to afford 1-(1-(4-(1-(5-chloro-4-((8-methoxy-1-methyl-3-(2-(methylamino)-2-oxoethoxy)-2-oxo-1,2-dihydroquinolin-6-yl)amino)pyrimidin-2-yl)piperidine-4-carboxamido)cyclohexane-1-carbonyl)piperidin-4-yl)-3-(7-(difluoromethyl)-6-(1-methyl-1H-pyrazol-4-yl)-3,4-dihydroquinolin-1(2H)-yl)-N-methyl-1,4,6,7-tetrahydro-5H-pyrazolo[4,3-c]pyridine-5-carboxamide (4.7 mg, 4.1 µmol, 40%, MS m/z 1162.38 $[M+H]^+$). **$^1$H NMR** δ = 8.96 (s, 1H), 8.09 (s, 1H), 7.94 (q, *J* = 4.8 Hz, 1H), 7.75 (d, *J* = 7.5 Hz, 1H), 7.73 (s, 1H), 7.52 (p, *J* = 2.6 Hz, 1H), 7.50 (d, *J* = 2.3 Hz, 1H), 7.48 (t, *J* = 1.4 Hz, 1H), 7.08 (s, 1H), 7.04 – 6.98 (m, 1H), 6.81 (s, 1H), 6.76 (t, *J* = 55.3 Hz, 1H), 6.54 (s, 1H), 4.55 (d, *J* = 2.3 Hz, 3H), 4.50 (d, *J* = 13.2 Hz, 2H), 4.32 (dq, *J* = 10.8, 5.4 Hz, 1H), 4.02 (s, 3H), 3.87 (d, *J* = 2.8 Hz, 3H), 3.86 (d, *J* = 1.7 Hz, 3H), 3.85 (s, 3H), 3.77 (s, 1H), 3.60 (t, *J* = 5.7 Hz, 2H), 3.56 (t, *J* = 5.6 Hz, 2H), 3.18 (t, *J* = 12.7 Hz, 1H), 2.98 (t, *J* = 12.4 Hz, 1H), 2.92 – 2.85 (m, 3H), 2.85 – 2.80 (m, 2H), 2.74 (t, *J* = 6.0 Hz, 2H), 2.69 (s, 1H), 2.68 – 2.59 (m, 3H), 2.54 (s, 3H), 1.96 (q, *J* = 5.9 Hz, 3H), 1.88 (s, 2H), 1.68 (d, *J* = 12.7 Hz, 7H), 1.48 (d, *J* = 29.4 Hz, 6H). **$^{19}$F NMR** (471 MHz, DMSO-$d_6$) δ = -108.22.

**Synthesis of MNN-03-040:**

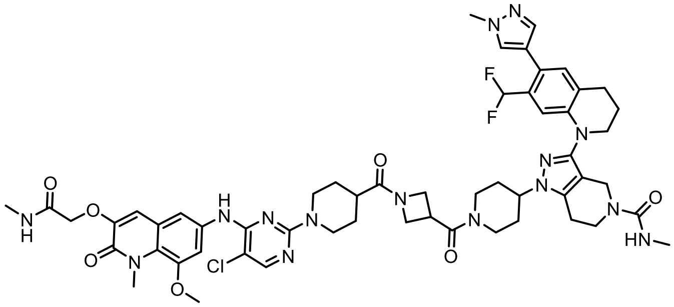

Compound MNN-03-040 was synthesized according to Procedure A using the linker 1-(tert-butoxycarbonyl)azetidine-3-carboxylic acid to afford 1-(1-(1-(1-(5-chloro-4-((8-methoxy-1-methyl-3-(2-(methylamino)-2-oxoethoxy)-2-oxo-1,2-dihydroquinolin-6-yl)amino)pyrimidin-2-yl)piperidine-4-carbonyl)azetidine-3-carbonyl)piperidin-4-yl)-3-(7-(difluoromethyl)-6-(1-methyl-1H-pyrazol-4-yl)-3,4-dihydroquinolin-1(2H)-yl)-N-methyl-1,4,6,7-tetrahydro-5H-pyrazolo[4,3-c]pyridine-5-carboxamide (0.82 mg, 0.73 µmol, 6%, MS m/z 1120.61 $[M+H]^+$). **$^1$H NMR** (500 MHz, DMSO-$d_6$) δ = 9.20 (s, 1H), 8.13 (d, *J* = 1.9 Hz, 1H), 7.94 (p, *J* = 4.6 Hz, 1H), 7.74 (d, *J* = 8.1 Hz, 1H), 7.51 (t, *J* = 2.7 Hz, 1H), 7.49 (dd, *J* = 3.7, 1.7 Hz, 2H), 7.08 (s, 1H), 7.00 (d, *J* = 5.1 Hz, 1H), 6.79 (s, 1H), 6.78 (t, *J* = 55.2, 3.1 Hz, 1H), 6.55 (s, 1H), 4.55 (d, *J* = 3.3 Hz, 2H), 4.49 – 4.36 (m, 4H), 4.36 – 4.23 (m, 2H), 4.01 (s, 2H), 3.98 (d, *J* = 3.7 Hz, 1H), 3.90 (dt, *J* = 9.1, 6.4 Hz, 1H), 3.86 (d, *J* = 3.3 Hz, 3H), 3.85 (d, *J* = 1.8 Hz, 3H), 3.85 (d, *J* = 2.9 Hz, 3H), 3.74 (p, *J* = 7.8 Hz, 1H), 3.66 (d, *J* = 13.3 Hz, 1H), 3.61 (q, *J* = 5.6 Hz, 2H), 3.56 (t, *J* = 5.6 Hz, 2H), 3.19 – 3.07 (m, 1H), 2.99 (t, *J* = 12.5 Hz, 2H), 2.82 (t, *J* = 6.7 Hz, 2H), 2.78 (s, 1H), 2.74 (t, *J* = 6.4 Hz, 2H), 2.64 (dd, *J* = 4.7, 2.1 Hz, 3H), 2.54 (d, *J* = 1.8 Hz, 3H), 1.96 (p, *J* = 6.3 Hz, 2H), 1.91 (d, *J* = 11.8 Hz, 3H), 1.81 – 1.72 (m, 2H), 1.68 (t, *J* = 15.0 Hz, 2H), 1.52 – 1.36 (m, 2H). **$^{19}$F NMR** (471 MHz, DMSO-$d_6$) δ = -108.21.

**Synthesis of MNN-03-049**:

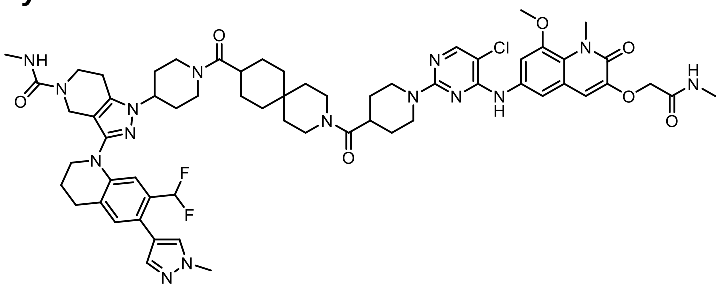

Compound MNN-03-049 was synthesized according to Procedure A using the linker 3-(tert-butoxycarbonyl)-3-azaspiro[5.5]undecane-9-carboxylic acid to afford 1-(1-(3-(1-(5-chloro-4-((8-methoxy-1-methyl-3-(2-(methylamino)-2-oxoethoxy)-2-oxo-1,2-dihydroquinolin-6-yl)amino)pyrimidin-2-yl)piperidine-4-carbonyl)-3-azaspiro[5.5]undecane-9-carbonyl)piperidin-4-yl)-3-(7-(difluoromethyl)-6-(1-methyl-1H-pyrazol-4-yl)-3,4-dihydroquinolin-1(2H)-yl)-N-methyl-1,4,6,7-tetrahydro-5H-pyrazolo[4,3-c]pyridine-5-carboxamide (2.7 mg, 2.2 µmol, 30%, MS m/z 1216.57 $[M+H]^+$). **$^1$H NMR** (500 MHz, DMSO-$d_6$) δ = 9.11 (s, 1H), 8.11 (s, 1H), 7.94 (q, *J* = 4.6 Hz, 1H), 7.74 (s, 1H), 7.52 (d, *J* = 2.3 Hz, 1H), 7.51 – 7.46 (m, 2H), 7.09 (s, 1H), 7.01 (s, 1H), 6.81 (s, 1H), 6.77 (t, *J* = 55.2 Hz, 1H), 6.54 (s, 1H), 4.55 (s, 2H), 4.45 (d, *J* = 12.7 Hz, 3H), 4.37 – 4.28 (m, 1H), 4.02 (s, 3H), 3.87 (s, 3H), 3.86 (s, 3H), 3.86 (s, 3H), 3.64 – 3.59 (m, 2H), 3.56 (t, *J* = 5.6 Hz, 2H), 3.47 (s, 2H), 3.41 (s, 2H), 3.18 (t, *J* = 12.7 Hz, 1H), 3.00 (s, 2H), 2.95 – 2.88 (m, 1H), 2.83 (t, *J* = 6.4 Hz, 2H), 2.74 (t, *J* = 5.7 Hz, 2H), 2.69 (s, 1H), 2.64 (d, *J* = 4.7 Hz, 3H), 2.60 (s, 1H), 2.54 (s, 3H), 1.96 (h, *J* = 6.8 Hz, 3H), 1.85 (d, *J* = 16.9 Hz, 2H), 1.73 – 1.63 (m, 5H), 1.50 (d, *J* = 16.4 Hz, 6H), 1.38 (s, 1H), 1.30 (s, 1H), 1.16 (dd, *J* = 24.9, 9.9 Hz, 4H). **$^{19}$F NMR** (471 MHz, DMSO-$d_6$) δ = -108.24.

**Synthesis of MNN-03-039**:

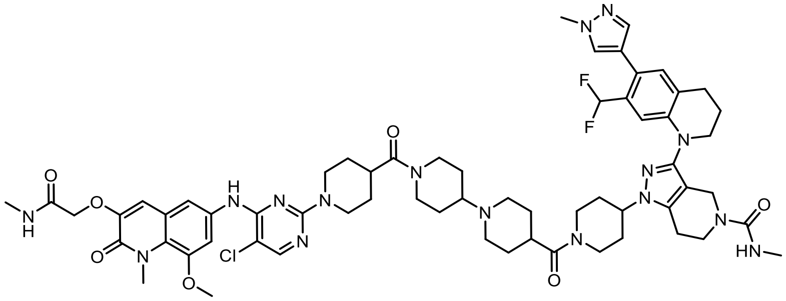

Compound MNN-03-039 was synthesized according to Procedure A using the linker 1'-(tert-butoxycarbonyl)-[1,4'-bipiperidine]-4-carboxylic acid to afford (1-(1-(1'-(1-(5-chloro-4-((8-methoxy-1-methyl-3-(2-(methylamino)-2-oxoethoxy)-2-oxo-1,2-dihydroquinolin-6-yl)amino)pyrimidin-2-yl)piperidine-4-carbonyl)-[1,4'-bipiperidine]-4-carbonyl)piperidin-4-yl)-3-(7-(difluoromethyl)-6-(1-methyl-1H-pyrazol-4-yl)-3,4-dihydroquinolin-1(2H)-yl)-N-methyl-1,4,6,7-tetrahydro-5H-pyrazolo[4,3-c]pyridine-5-carboxamide) (3.1 mg, 2.5 µmol, 40%, MS m/z 1232.68 $[M+H]^+$). **$^1$H NMR** (500 MHz, DMSO-$d_6$) δ = 9.24 (s, 1H), 8.99 (s, 1H), 8.10 (s, 1H), 7.97 (q, *J* = 4.7 Hz, 1H), 7.75 (s, 1H), 7.57 (d, *J* = 2.2 Hz, 1H), 7.50 (d, *J* = 2.3 Hz, 1H), 7.49 (d, *J* = 0.8 Hz, 1H), 7.09 (s, 1H), 7.00 (s, 1H), 6.81 (s, 1H), 6.78 (t, *J* = 55.2 Hz, 1H), 6.56 (s, 1H), 4.55 (s, 2H), 4.48 (s, 4H), 4.34 (d, *J* = 10.5 Hz, 1H), 4.24 – 4.05 (m, 4H, determined by HSQC), 4.03 (s, 2H), 3.87 (s, 3H), 3.86 (s, 3H), 3.85 (s, 3H), 3.61 (t, *J* = 5.6 Hz, 2H), 3.56 (t, *J* = 5.6 Hz, 2H), 3.44 (s, 3H), 3.27 – 3.16 (m, 1H), 3.01 (dd, *J* = 29.5, 16.6 Hz, 7H, determined by HSQC), 2.83 (d, *J* = 6.5 Hz, 2H), 2.75 (s, 3H), 2.64 (d, *J* = 4.6 Hz, 3H), 2.54 (s, 3H), 1.97 (q, *J* = 5.8 Hz, 5H), 1.87 (d, *J* = 12.8 Hz, 5H), 1.78 – 1.61 (m, 3H), 1.61 – 1.35 (m, 4H). **$^{19}$F NMR** (471 MHz, DMSO-$d_6$) δ = -108.18.

**Synthesis of NEG1:**

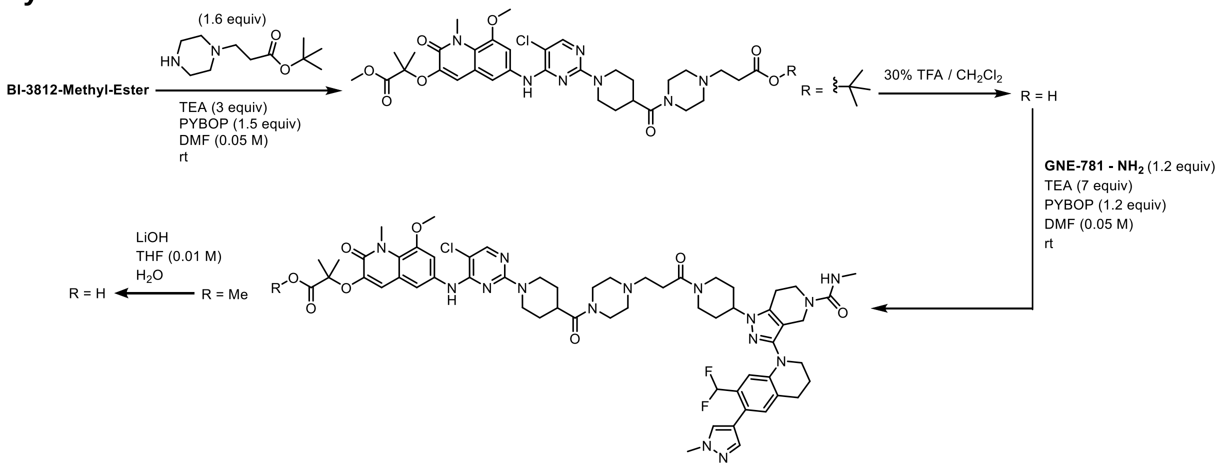

To a solution of 1-(5-chloro-4-((3-((1-ethoxy-2-methyl-1-oxopropan-2-yl)oxy)-8-methoxy-1-methyl-2-oxo-1,2-dihydroquinolin-6-yl)amino)pyrimidin-2-yl)piperidine-4-carboxylic acid (19 mg, 34 µmol, 1.0 eq) in DMF (670 µL, 0.05 M) was added tert-butyl 3-(piperazin-1-yl)propanoate (12 mg, 54 µmol, 1.6 equiv), TEA (23 µL,170 µmol, 5 equiv), and HATU (18 mg, 47 µmol, 1.4 equiv) on ice. The reaction was allowed to stir at room temperature until LC-MS indicated full conversion of limited starting material. The reaction mixture was purified by reverse phase chromatography (10-100% MeCN in water), and concentrated under reduced pressure to afford ethyl 2-((6-((2-(4-(4-(3-(tert-butoxy)-3-oxopropyl)piperazine-1-carbonyl)piperidin-1-yl)-5-chloropyrimidin-4-yl)amino)-8-methoxy-1-methyl-2-oxo-1,2-dihydroquinolin-3-yl)oxy)-2-methylpropanoate (20 mg, 2.5 µmol, 76%, MS m/z 770/32 $[M+H]^+$).

Ethyl 2-((6-((2-(4-(4-(3-(tert-butoxy)-3-oxopropyl)piperazine-1-carbonyl)piperidin-1-yl)-5-chloropyrimidin-4-yl)amino)-8-methoxy-1-methyl-2-oxo-1,2-dihydroquinolin-3-yl)oxy)-2-methylpropanoate was then allowed to stir in 30% v/v TFA in $CH_2Cl_2$ (0.25 mL, 0.1 M) for 2 hours. The reaction was concentrated under reduced pressure, and residual TFA was removed by co-evaporation with DCM (3x), followed by high vacuum overnight to afford a crude product. To a solution of 3-(4-(1-(5-chloro-4-((3-((1-ethoxy-2-methyl-1-oxopropan-2-yl)oxy)-8-methoxy-1-methyl-2-oxo-1,2-dihydroquinolin-6-yl)amino)pyrimidin-2-yl)piperidine-4-carbonyl)piperazin-1-yl)propanoic acid (17 mg, 24 µmol, 1.0 equiv) dissolved in DMF (480 µL, 0.05 M), TEA (23 µL, 170 µmol, 7.0 equiv), and PyBOP (15 mg, 29 µmol, 1.2 equiv) was added **GNE-781-NH$_2$** (3-(7-(difluoromethyl)-6-(1-methyl-1H-pyrazol-4-yl)-3,4-dihydroquinolin-1(2H)-yl)-N-methyl-1-(piperidin-4-yl)-1,4,6,7-tetrahydro-5H-pyrazolo[4,3-c]pyridine-5-carboxamide) (11 mg, 21 µmol, 0.9 equiv) dissolved in a minimal amount of DMF in a dropwise manner on ice. The reaction was allowed to stir until LC-MS indicate completion, at which point it was purified by reverse phase chromatography (5-100% MeCN in water), and concentrated under reduced pressure to afford ethyl 2-((6-((5-chloro-2-(4-(4-(3-(4-(3-(7-(difluoromethyl)-6-(1-methyl-1H-pyrazol-4-yl)-3,4-dihydroquinolin-1(2H)-yl)-5-(methylcarbamoyl)-4,5,6,7-tetrahydro-1H-pyrazolo[4,3-c]pyridin-1-yl)piperidin-1-yl)-3-oxopropyl)piperazine-1-carbonyl)piperidin-1-yl)pyrimidin-4-yl)amino)-8-methoxy-1-methyl-2-oxo-1,2-dihydroquinolin-3-yl)oxy)-2-methylpropanoate (7.9 mg, 24 µmol, 27%, MS m/z 1222.24 $[M+H]^+$).

To a solution of ethyl 2-((6-((5-chloro-2-(4-(4-(3-(4-(3-(7-(difluoromethyl)-6-(1-methyl-1H-pyrazol-4-yl)-3,4-dihydroquinolin-1(2H)-yl)-5-(methylcarbamoyl)-4,5,6,7-tetrahydro-1H-pyrazolo[4,3-c]pyridin-1-yl)piperidin-1-yl)-3-oxopropyl)piperazine-1-carbonyl)piperidin-1-yl)pyrimidin-4-yl)amino)-8-methoxy-1-methyl-2-oxo-1,2-dihydroquinolin-3-yl)oxy)-2-methylpropanoate (7.9 mg, 6.5 µmol, 1.0 equiv) dissolved in THF (230 µL, 0.01 M) and $H_2O$ (4 µL) was added LiOH (2.7 mg,

64 µmol, 10 equiv). The reaction was allowed to stir overnight, at which point the reaction was purified via reverse-phase HPLC (10-100% MeCN in water), concentrated under reduced pressure, and dried through lyophilization to afford a white solid (3.5 mg, 2.9 µmol, 45%, MS m/z 1193.10 $[M+H]^+$). **$^1$H NMR** (500 MHz, DMSO-$d_6$) δ 9.62 (s, 1H), 8.96 (s, 1H), 8.10 (s, 1H), 7.74 (s, 1H), 7.64 (d, *J* = 2.2 Hz, 1H), 7.53 (d, *J* = 2.3 Hz, 1H), 7.49 (s, 1H), 7.09 (s, 1H), 6.88 (s, 1H), 6.78 (t, *J* = 55.2 Hz, 1H), 6.76 (s, 1H), 6.55 (s, 1H), 4.49 (t, *J* = 16.1 Hz, 5H), 4.38 (t, *J* = 9.0 Hz, 1H), 4.24 (s, 2H), 4.02 (s, 2H), 3.95 (d, *J* = 13.3 Hz, 1H), 3.87 (s, 3H), 3.86 (s, 3H), 3.84 (s, 3H), 3.61 (dt, *J* = 10.9, 6.0 Hz, 2H), 3.56 (dd, *J* = 7.6, 4.8 Hz, 3H), 3.45 (s, 1H), 3.36 (s, 2H), 3.27 – 3.18 (m, 1H), 3.08 (s, 2H), 3.04 – 2.96 (m, 4H), 2.91 (d, *J* = 7.5 Hz, 2H), 2.83 (d, *J* = 6.2 Hz, 2H), 2.79 (dd, *J* = 12.6, 2.6 Hz, 1H), 2.75 (q, *J* = 4.3 Hz, 2H), 2.54 (s, 3H), 1.96 (td, *J* = 8.7, 4.5 Hz, 5H), 1.77 (dd, *J* = 11.6, 4.0 Hz, 2H), 1.71 (d, *J* = 12.5 Hz, 2H), 1.55 (s, 6H).

**Synthesis of NEG2:**

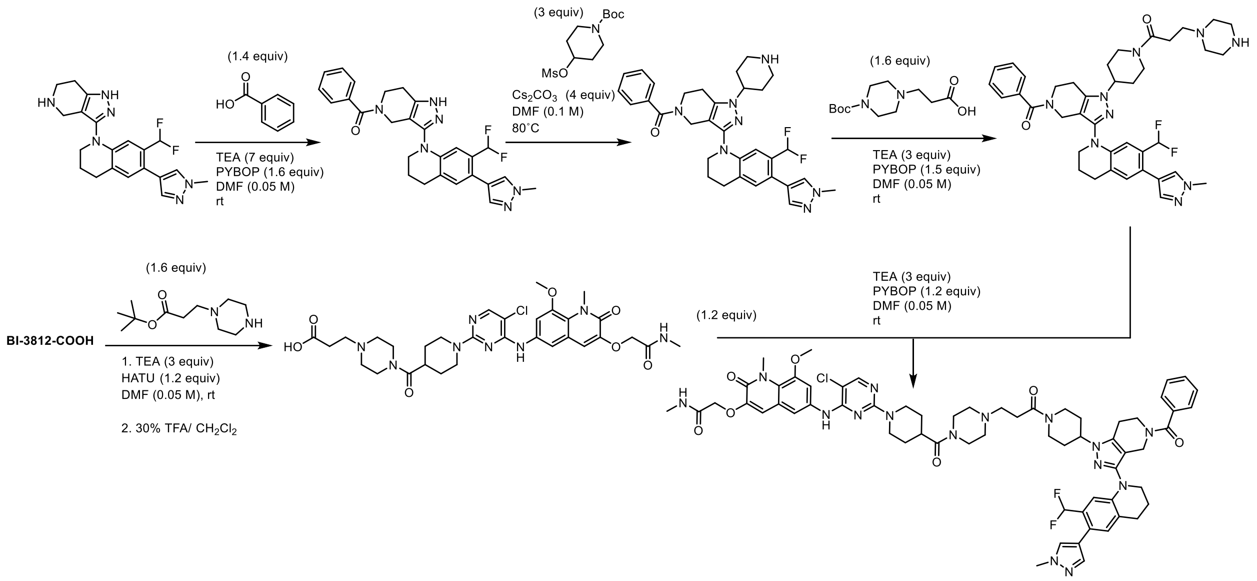

To a solution of benzoic acid (180 mg, 150 µmol, 1.4 equiv) in DMF (2.1 mL, 0.05 M) with TEA (102 µL, 728 µmol, 7 equiv) and Pybop (87 mg, 170 µmol, 1.6 equiv) was added 7-(difluoromethyl)-6-(1-methyl-1H-pyrazol-4-yl)-1-(4,5,6,7-tetrahydro-1H-pyrazolo[4,3-c]pyridin-3-yl)-1,2,3,4-tetrahydroquinoline (40.0 mg, 104 µmol, 1.0 equiv). The reaction was allowed to stir at room temperature until LC-MS indicated completion. The reaction was purified by reverse phase chromatography (10-100% MeCN in water), and concentrated under reduced pressure to afford (3-(7-(difluoromethyl)-6-(1-methyl-1H-pyrazol-4-yl)-3,4-dihydroquinolin-1(2H)-yl)-1,4,6,7-tetrahydro-5H-pyrazolo[4,3-c]pyridin-5-yl)(phenyl)methanone (40 mg, 81 µmol, 78%, MS m/z 489.16 $[M+H]^+$).

A solution of (3-(7-(difluoromethyl)-6-(1-methyl-1H-pyrazol-4-yl)-3,4-dihydroquinolin-1(2H)-yl)-1,4,6,7-tetrahydro-5H-pyrazolo[4,3-c]pyridin-5-yl)(phenyl)methanone (39 mg, 810 µmol, 1.0 equiv), tert-butyl 4-((methylsulfonyl)oxy)piperidine-1-carboxylate (90 mg, 320 µmol, 4.0 equiv), and cesium carbonate (79 mg, 240 µmol, 3.0 equiv) in DMF(810 µL, 0.1 M) was heated at 80˚C while stirring for 24 hours. The reaction was purified by reverse-phase HPLC (90% water: 100% MeCN), and fractions from the second peak were collected and concentrated under high vacuum (MS m/z 672.30 $[M+H]^+$). The residue was stirred in 30% v/v $TFA/CH_2Cl_2$ for 1 hour, then concentrated under reduced pressure, and residual TFA was removed by co-evaporation with DCM (3x), followed by high vacuum overnight to afford (3-(7-(difluoromethyl)-6-(1-methyl-1Hpyrazol-4-yl)-3,4-dihydroquinolin-1(2H)-yl)-1-(piperidin-4-yl)-1,4,6,7-tetrahydro-5H-pyrazolo[4,3-c]pyridin-5-yl)(phenyl)methanone (11 mg, 19 µmol, 24%, MS m/z 572.38 $[M+H]^+$).

To a solution of 1-(5-chloro-4-((8-methoxy-1-methyl-3-(2-(methylamino)-2-oxoethoxy)-2-oxo-1,2-dihydroquinolin-6-yl)amino)pyrimidin-2-yl)piperidine-4-carboxylic acid (BI-3812-COOH) (50 mg, 94 µmol, 1.0 equiv) in DMF (0.05 M, 1.9 mL) with TEA (39 µL, 280 µmol, 3.0 equiv) and PyBOP (59 mg, 130 µmol, 1.2 equiv) on ice was added tert-butyl 3-(piperazin-1-yl)propanoate (32 mg, 150 µmol, 1.6 equiv). The reaction was allowed to stir at room temperature until LC-MS indicated completion of the reaction, then it was purified by reverse phase chromatography (5-100% MeCN in water), and concentrated under reduced pressure to afford tert-butyl 3-(4-(1-(5-chloro-4-((8-methoxy-1-methyl-3-(2-(methylamino)-2-oxoethoxy)-2-oxo-1,2-dihydroquinolin-6-yl)amino)pyrimidin-2-yl)piperidine-4-carbonyl)piperazin-1-yl)propanoate (37 mg, 55 µmol, 58%, MS m/z 727.17 $[M+H]^+$). The product was then allowed to stir in 30% v/v TFA/$CH_2Cl_2$ (0.55 mL, 0.1 M) for 2 hours, concentrated under reduced pressure, and residual TFA was removed by co-evaporation with DCM (3x) followed by high vacuum overnight to afford 3-(4-(1-(5-chloro-4-((8-methoxy-1-methyl-3-(2-(methylamino)-2-oxoethoxy)-2-oxo-1,2-dihydroquinolin-6-yl)amino)pyrimidin-2-yl)piperidine-4-carbonyl)piperazin-1-yl)propanoic acid as a crude product (MS m/z 672.23 $[M+H]^+$).

To a solution of 3-(4-(1-(5-chloro-4-((8-methoxy-1-methyl-3-(2-(methylamino)-2-oxoethoxy)-2-oxo-1,2-dihydroquinolin-6-yl)amino)pyrimidin-2-yl)piperidine-4-carbonyl)piperazin-1-yl)propanoic acid (9.9 mg, 15 µmol, 1.2 equiv) ) in DMF (250 µL, 0.05 M) with TEA (39 µL, 280 µmol, 3.0 equiv) and PyBOP (59 mg, 130 µmol, 1.2 equiv) on ice was added (3-(7-(difluoromethyl)-6-(1-methyl-1H-pyrazol-4-yl)-3,4-dihydroquinolin-1(2H)-yl)-1-(piperidin-4-yl)-1,4,6,7-tetrahydro-5H-pyrazolo[4,3-c]pyridin-5-yl)(phenyl)methanone (7.0 mg, 12 µmol, 1.0 equiv). The reaction was allowed to stir for 3 hours, at which point it was purified by reverse-phase HPLC (10-100% MeCN in water) to afford 2-((6-((2-(4-(4-(3-(4-(5-benzoyl-3-(7-(difluoromethyl)-6-(1-methyl-1H-pyrazol-4-yl)-3,4-dihydroquinolin-1(2H)-yl)-4,5,6,7-tetrahydro-1H-pyrazolo[4,3-c]pyridin-1-yl)piperidin-1-yl)-3-oxopropyl)piperazine-1-carbonyl)piperidin-1-yl)-5-chloropyrimidin-4-yl)amino)-8-methoxy-1-methyl-2-oxo-1,2-dihydroquinolin-3-yl)oxy)-N-methylacetamide as a white solid (5.8 mg, 4.8 µmol, 39%, MS m/z 1224.63 $[M+H]^+$). **$^1$H NMR** (500 MHz, DMSO-$d_6$) δ = 9.72 (s, 1H), 9.12 (s, 1H), 8.13 (s, 1H), 7.96 (d, *J* = 4.9 Hz, 1H), 7.77 (s, 1H), 7.57 (t, *J* = 1.7 Hz, 1H), 7.51 (s, 1H), 7.49 (d, *J* = 2.3 Hz, 1H), 7.45 (s, 3H), 7.19 (s, 1H), 7.00 (s, 1H), 6.79 (t, *J* = 55.0 Hz, 1H), 6.36 (s, 1H), 4.56 (s, 2H), 4.47 (d, *J* = 13.6 Hz, 3H), 4.47 (s, 1H), 4.26 (s, 3H), 3.92 (s, 1H), 3.87 (s, 6H), 3.86 (s, 3H), 3.55 (s, 7H), 3.37 (s, 3H), 3.23 (s, 1H), 3.11 (s, 1H), 3.07 – 2.84 (m, 7H), 2.80 (d, *J* = 15.6 Hz, 2H), 2.71 (d, *J* = 21.4 Hz, 1H), 2.65 (d, *J* = 4.6 Hz, 3H), 2.16 – 1.96 (m, 4H), 1.95 – 1.79 (m, 3H), 1.71 (d, *J* = 12.5 Hz, 3H), 1.53 (s, 2H). **$^{19}$F NMR** (471 MHz, DMSO-$d_6$) δ = -108.02.

**Synthesis of MNN-06-112:**

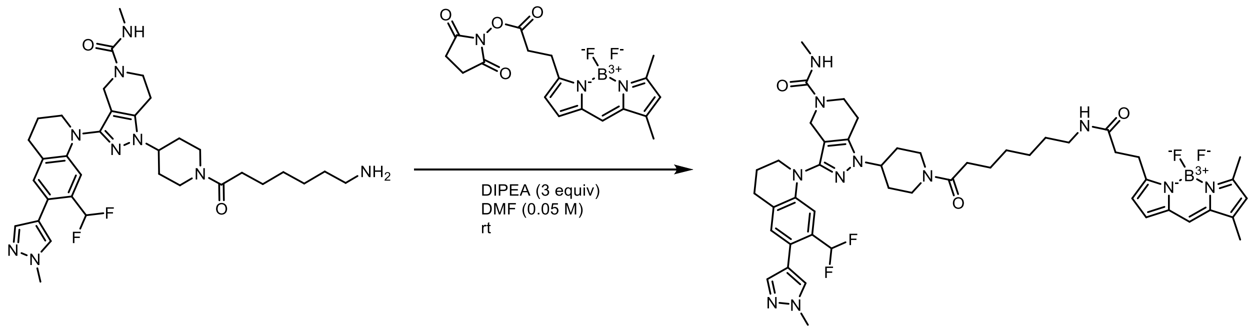

3-(7-(difluoromethyl)-6-(1-methyl-1H-pyrazol-4-yl)-3,4-dihydroquinolin-1(2H)-yl)-N-methyl-1-(piperidin-4-yl)-1,4,6,7-tetrahydro-5H-pyrazolo[4,3-c]pyridine-5-carboxamide (6.0 mg, 10 µmol, 1 equiv) was reacted with 7-((tert-butoxycarbonyl)amino)heptanoic acid (6.0 mg, 20 µmol, 2 equiv) as described in Procedure A to afford tert-butyl (7-(4-(3-(7-(difluoromethyl)-6-(1-methyl-1H-pyrazol-4-yl)-3,4-dihydroquinolin-1(2H)-yl)-5-(methylcarbamoyl)-4,5,6,7-tetrahydro-1H-pyrazolo[4,3-c]pyridin-1-yl)piperidin-1-yl)-7-oxoheptyl)carbamate (4.5 mg, 6.0 µmol, 50%), which was then stirred in 30% TFA/$CH_2Cl_2$ v/v (0.060 mL, 0.1 M) for 2 hours to afford a crude mixture. To a solution of 2,5-dioxopyrrolidin-1-yl 3-(5,5-difluoro-7,9-dimethyl-5H-5l4,6l4-dipyrrolo[1,2-c:2',1'-f][1,3,2]diazaborinin-3-yl)propanoate (8.0 mg, 12 µmol, 1.0 equiv) in DMF (250 µL, 0.05 M) and DIPEA (6.6 µL, 37 µmol, 3.0 equiv) on ice was added 2,5-dioxopyrrolidin-1-yl 3-(5,5-difluoro-7,9-dimethyl-5H-5l4,6l4-dipyrrolo[1,2-c:2',1'-f][1,3,2]diazaborinin-3-yl)propanoate (4.8 mg, 12 µmol, 1.0 equiv). The reaction was allowed to stir for 8 hours and was then purified by reverse-phase HPLC (10-100% MeCN in water) to afford 1-(1-(7-(3-(5,5-difluoro-7,9-dimethyl-5H-5l4,6l4-dipyrrolo[1,2-c:2',1'-f][1,3,2]diazaborinin-3-yl)propanamido)heptanoyl)piperidin-4-yl)-3-(7-(difluoromethyl)-6-(1-methyl-1H-pyrazol-4-yl)-3,4-dihydroquinolin-1(2H)-yl)-N-methyl-1,4,6,7-tetrahydro-5H-pyrazolo[4,3-c]pyridine-5-carboxamideand isolated as an orange solid (2.4 mg, 2.6 µmol, 22%, MS m/z 926.36 $[M+H]^+$). **$^1$H NMR** (500 MHz, DMSO-$d_6$) δ = 7.86 (t, *J* = 5.6 Hz, 1H), 7.74 (s, 1H), 7.68 (s, 1H), 7.49 (s, 1H), 7.10 – 7.05 (m, 2H), 6.79 (s, 1H), 6.77 (t, *J* = 55.3 Hz, 1H), 6.54 (s, 1H), 6.33 (d, *J* = 4.0 Hz, 1H), 6.29 (s, 1H), 4.46 (d, *J* = 13.0 Hz, 1H), 4.31 (tt, *J* = 10.2, 4.4 Hz, 1H), 4.01 (s, 3H), 3.95 (d, *J* = 13.6 Hz, 1H), 3.86 (s, 3H), 3.60 (q, *J* = 5.5 Hz, 2H), 3.56 (t, *J* = 5.7 Hz, 2H), 3.20 – 3.11 (m, 1H), 3.05 (dt, *J* = 16.5, 7.2 Hz, 4H), 2.83 (t, *J* = 6.5 Hz, 2H), 2.73 (t, *J* = 5.7 Hz, 2H), 2.69 (d, *J* = 12.3 Hz, 1H), 2.54 (d, *J* = 1.9 Hz, 3H), 2.46 (s, 3H), 2.46 (d, *J* = 15.4 Hz, 2H), 2.32 (t, *J* = 7.5 Hz, 2H), 2.25 (s, 3H), 1.96 (p, *J* = 6.1 Hz, 2H), 1.88 (t, *J* = 11.2 Hz, 3H), 1.72 (dt, *J* = 14.1, 10.0 Hz, 1H), 1.48 (p, *J* = 7.3 Hz, 2H), 1.38 (p, *J* = 7.0 Hz, 2H), 1.26 (dq, *J* = 10.4, 6.6 Hz, 5H). **$^{19}$F NMR** (471 MHz, DMSO-$d_6$) δ -108.11, -143.24 (dd, *J* = 66.9, 33.0 Hz).

**Synthesis of TNL15:**

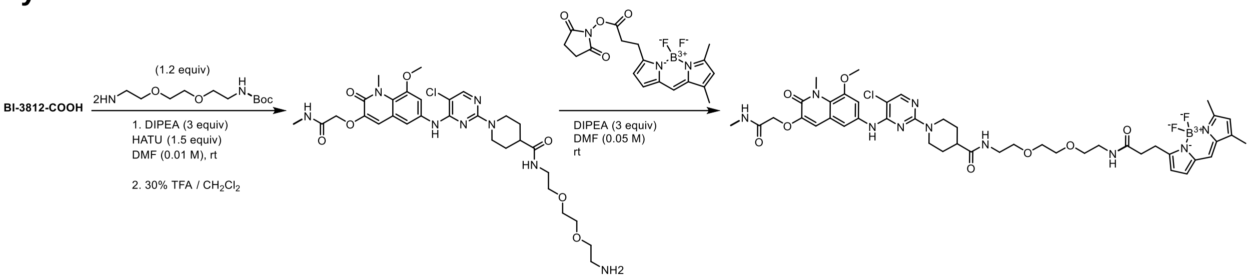

A solution of 1-(5-chloro-4-((8-methoxy-1-methyl-3-(2-(methylamino)-2-oxoethoxy)-2-oxo-1,2-dihydroquinolin-6-yl)amino)pyrimidin-2-yl)piperidine-4-carboxylic acid (**BI-3812-COOH**) (10.0 mg, 19 µmol, 1.0 equiv), HATU (11 mg, 28 µmol, 1.5 equiv), tert-butyl (2-(2-(2-aminoethoxy)ethoxy)ethyl)carbamate (5.6 mg, 23 µmol, 1.2 equiv), and DIPEA (9.8 µL, 57 µmol, 3 equiv) was stirred at room temperature until LC-MS indicate completion of the reaction. The reaction was purified by reverse phase chromatography (5-100% MeCN in water), concentrated under reduced pressure, stirred with 30% v/v TFA in $CH_2Cl_2$ (0.57 mL, 0.1 M) for 2 hours, concentrated under reduced pressure, and residual TFA was removed by co-evaporation with DCM (3x) followed by high vacuum overnight to afford N-(2-(2-(2-aminoethoxy)ethoxy)ethyl)-1-(5-chloro-4-((8-methoxy-1-methyl-3-(2-(methylamino)-2-oxoethoxy)-2-oxo-1,2-dihydroquinolin-6-yl)amino)pyrimidin-2-yl)piperidine-4-carboxamide as a crude product. To a solution of N-(2-(2-(2-aminoethoxy)ethoxy)ethyl)-1-(5-chloro-4-((8-methoxy-1-methyl-3-(2-(methylamino)-2-oxoethoxy)-2-oxo-1,2-dihydroquinolin-6-yl)amino)pyrimidin-2-yl)piperidine-4-carboxamide (10.0 mg, 15 µmol, 1.0 equiv), DMF (300 µL, 0.05 M), and DIPEA (7.9 µL, 45 µmol, 3 equiv), was added 2,5-dioxopyrrolidin-1-yl 3-(5,5-difluoro-7,9-dimethyl-5H-5l4,6l4-dipyrrolo[1,2-c:2',1'-f][1,3,2]diazaborinin-3-yl)propanoate (5.9 mg, 15 µmol, 1.0 equiv). The reaction was allowed to stir for 2 hours, then purified by reverse-phase HPLC (10-100% MeCN in water) to afford 1-(5-chloro-4-((8-methoxy-1-methyl-3-(2-(methylamino)-2-oxoethoxy)-2-oxo-1,2-dihydroquinolin-6-yl)amino)pyrimidin-2-yl)-N-(2-(2-(2-(3-(5,5-difluoro-7,9-dimethyl-5H-5l4,6l4-dipyrrolo[1,2-c:2',1'-f][1,3,2]diazaborinin-3-yl)propanamido)ethoxy)ethoxy) ethyl) piperidine-4-carboxamide (5.1 mg, 5.4 µmol, 36%, MS m/z 935.4 $[M+H]^+$). **$^1$H NMR** (500 MHz, DMSO-$d_6$) δ = 9.04 (s, 1H), 8.10 (s, 1H), 8.01 (t, *J* = 5.6 Hz, 1H), 7.95 (q, *J* = 4.7 Hz, 1H), 7.89 (t, *J* = 5.7 Hz, 1H), 7.67 (s, 1H), 7.54 – 7.49 (m, 2H), 7.06 (d, *J* = 4.1 Hz, 1H), 7.00 (s, 1H), 6.34 (d, *J* = 4.0 Hz, 1H), 6.28 (d, *J* = 1.1 Hz, 1H), 4.55 (s, 2H), 4.46 (d, *J* = 13.0 Hz, 2H), 3.86 (s, 3H), 3.85 (s, 3H), 3.50 (s, 4H), 3.40 (q, *J* = 6.0 Hz, 4H), 3.20 (dq, *J* = 13.8, 6.8 Hz, 4H), 3.06 (t, *J* = 7.8 Hz, 2H), 2.96 – 2.86 (m, 2H), 2.64 (d, *J* = 4.6 Hz, 3H), 2.48 (d, *J* = 7.7 Hz, 2H), 2.45 (s, 3H), 2.41 (dt, *J* = 11.5, 3.9 Hz, 1H), 2.24 (s, 3H), 1.70 (d, *J* = 12.8 Hz, 2H), 1.50 (qd, *J* = 12.4, 4.0 Hz, 2H). **$^{19}$F NMR** (471 MHz, DMSO-$d_6$) δ = -143.24 (dd, *J* = 66.9, 32.8 Hz).

**General Procedure B:**

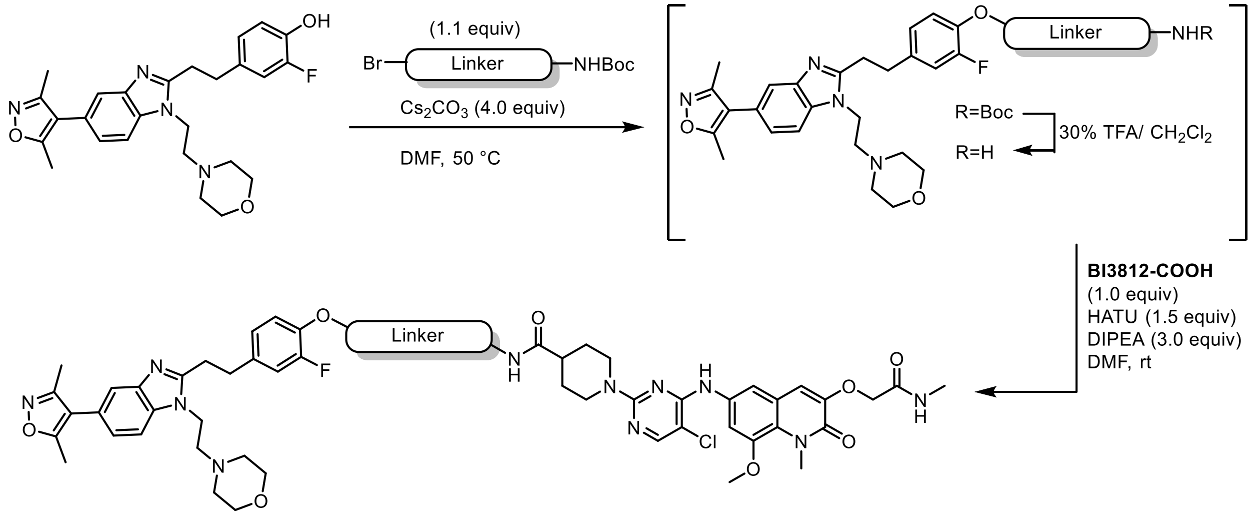

To a solution of 4-(2-(5-(3,5-dimethylisoxazol-4-yl)-1-(2-morpholinoethyl)-1H-benzo[d]imidazol-2-yl)ethyl)-2-fluorophenol (1.0 equiv) and linker (1.1 equiv) in DMF (0.2 M) was added $Cs_2CO_3$ (4.0 equiv) and the resulting suspension was vigorously stirred at 50 °C until LC-MS and/or TLC analysis indicated full conversion of limiting starting material. The reaction mixture was diluted with water and EtOAc. After separation of the organic layer, the aqueous phase was extracted with EtOAc (3x) and the combined organic extracts were washed with brine, dried over $Na_2SO_4$, and concentrated under reduced pressure. The resulting intermediate was taken up in 30% v/v TFA in $CH_2Cl_2$ and the resulting solution was stirred at ambient temperature until LC-MS showed quantitative formation of free amine intermediate. All volatiles were removed under reduced pressure. The intermediate was used without further purification, dissolved in DMF (0.2 M), and added to a solution of **BI3812-COOH** (1.0 equiv), HATU (1.5 equiv), and DIPEA (3.0 equiv) in DMF. The reaction mixture was purified by reverse-phase HPLC to afford the product upon lyophilization.

Synthesis of **RCS-02-107:**

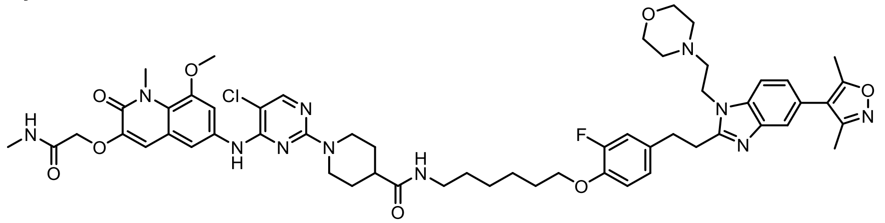

The corresponding compound was prepared following General Procedure B using *tert*-butyl (6-bromohexyl)carbamate as the linker. Purification by preparative reverse-phase HPLC (15-90% MeOH in water) afforded 1-(5-chloro-4-((8-methoxy-1-methyl-3-(2-(methylamino)-2-oxoethoxy)-2-oxo-1,2-dihydroquinolin-6-yl)amino)pyrimidin-2-yl)-N-(6-(4-(2-(5-(3,5-dimethylisoxazol-4-yl)-1-(2-morpholinoethyl)-1H-benzo[d]imidazol-2-yl)ethyl)-2-fluorophenoxy)hexyl)piperidine-4-carboxamide (1.9 mg, 1.8 μmol, 10%, MS m/z 1076.61 $[M+H]^+$). $^{1}$**H NMR** (500 MHz, DMSO-$d_6$) δ = 8.94 (s, 1H), 8.09 (s, 1H), 7.96 (d, *J* = 5.0 Hz, 1H), 7.86 (d, *J* = 8.4 Hz, 1H), 7.79 (t, *J* = 5.7 Hz, 1H), 7.71 (d, *J* = 1.5 Hz, 1H), 7.53 (q, *J* = 2.3 Hz, 2H), 7.42 – 7.38 (m, 1H), 7.25 – 7.19 (m, 1H), 7.09 – 7.05 (m, 2H), 7.00 (s, 1H), 4.69 (d, *J* = 8.0 Hz, 2H), 4.54 (s, 2H), 4.49 (d, *J* = 13.1 Hz, 2H), 3.99 (t, *J* = 6.5 Hz, 2H), 3.87 (s, 3H), 3.85 (s, 3H), 3.43 (s, 2H), 3.32 (t, *J* = 8.0 Hz, 2H), 3.14 (t, *J* = 8.0 Hz, 2H), 3.02 (q, *J* = 6.6 Hz, 2H), 2.90 (t, *J* = 12.3 Hz, 2H), 2.64 (d, *J* = 4.6 Hz, 3H), 2.41 (s, 3H), 2.39 – 2.34 (m, 1H), 2.23 (s, 3H), 1.69 (q, *J* = 9.1 Hz, 4H), 1.55 – 1.45 (m, 2H), 1.38 (dd, *J* = 12.7, 6.5 Hz, 4H), 1.33 – 1.27 (m, 2H). $^{19}$**F NMR** (471 MHz, DMSO-$d_6$) δ = -74.4.

Synthesis of **RCS-02-108:**

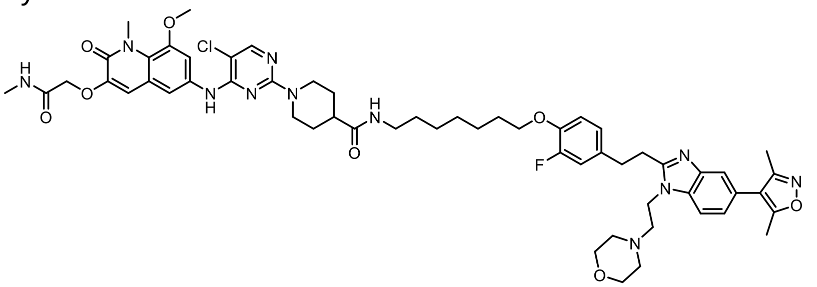

The corresponding compound was prepared following General Procedure B using *tert*-butyl (7-bromoheptyl)carbamate as the linker. Purification by preparative reverse-phase HPLC (15-90% MeOH in water) afforded 1-(5-chloro-4-((8-methoxy-1-methyl-3-(2-(methylamino)-2-oxoethoxy)-2-oxo-1,2-dihydroquinolin-6-yl)amino)pyrimidin-2-yl)-N-(7-(4-(2-(5-(3,5-dimethylisoxazol-4-yl)-1-(2-morpholinoethyl)-1H-benzo[d]imidazol-2-yl)ethyl)-2-fluorophenoxy)heptyl)piperidine-4-carboxamide (1.7 mg, 1.6 µmol, 9%, MS m/z 1090.67 [M+H]$^{+}$). $^{1}$**H NMR** (500 MHz, DMSO-$d_6$) δ = 8.90 (s, 1H), 8.08 (s, 1H), 7.96 (d, *J* = 5.1 Hz, 1H), 7.82 (d, *J* = 8.4 Hz, 1H), 7.78 (t, *J* = 5.6 Hz, 1H), 7.69 (s, 1H), 7.53 (d, *J* = 1.5 Hz, 2H), 7.38 (d, *J* = 8.4 Hz, 1H), 7.24 – 7.18 (m, 1H), 7.07 (d, *J* = 6.6 Hz, 2H), 7.00 (s, 1H), 4.67 (t, *J* = 7.9 Hz, 2H), 4.54 (s, 2H), 4.50 (d, *J* = 13.0 Hz, 2H), 3.99 (t, *J* = 6.5 Hz, 2H), 3.87 (s, 3H), 3.84 (s, 3H), 3.45 – 3.38 (m, 2H), 3.31 (t, *J* = 8.0 Hz, 2H), 3.14 (t, *J* = 8.0 Hz, 2H), 3.02 (q, *J* = 6.6 Hz, 2H), 2.92 – 2.86 (m, 2H), 2.64 (d, *J* = 4.7 Hz, 3H), 2.41 (s, 3H), 2.38 – 2.34 (m, 1H), 2.23 (s, 3H), 1.69 (t, *J* = 7.8 Hz, 4H), 1.50 (p, *J* = 10.1 Hz, 2H), 1.37 (q, *J* = 7.2 Hz, 4H), 1.33 – 1.22 (m, 6H). $^{19}$**F NMR** (471 MHz, DMSO-$d_6$) δ = -74.2.

Synthesis of **RCS-02-109:**

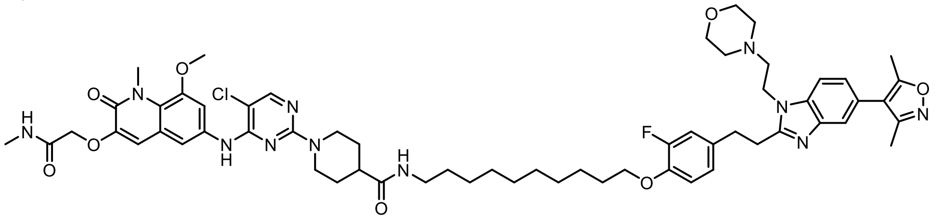

The corresponding compound was prepared following General Procedure B using *tert*-butyl (10-bromodecyl)carbamate as the linker. Purification by preparative reverse-phase HPLC (15-90% MeOH in water) afforded 1-(5-chloro-4-((8-methoxy-1-methyl-3-(2-(methylamino)-2-oxoethoxy)-2-oxo-1,2-dihydroquinolin-6-yl)amino)pyrimidin-2-yl)-N-(10-(4-(2-(5-(3,5-dimethylisoxazol-4-yl)-1-(2-morpholinoethyl)-1H-benzo[d]imidazol-2-yl)ethyl)-2-fluorophenoxy)decyl)piperidine-4-carboxamide (9.0 mg, 8.0 µmol, 50%, MS m/z 1132.93 [M+H]$^{+}$). $^{1}$**H NMR** (500 MHz, DMSO-$d_6$) δ = 8.97 (s, 1H), 8.12 (s, 1H), 8.03 – 7.98 (m, 1H), 7.91 (d, *J* = 8.4 Hz, 1H), 7.81 (q, *J* = 5.8 Hz, 1H), 7.75 (d, *J* = 1.6 Hz, 1H), 7.57 (q, *J* = 2.3 Hz, 2H), 7.47 – 7.42 (m, 1H), 7.27 (d, *J* = 12.6 Hz, 1H), 7.12 (d, *J* = 5.3 Hz, 2H), 7.04 (s, 1H), 4.75 (t, *J* = 7.9 Hz, 2H), 4.59 (s, 2H), 4.53 (d, *J* = 13.1 Hz, 2H), 4.06 – 3.96 (m, 2H), 3.91 (s, 3H), 3.89 (s, 3H), 3.47 (s, 2H), 3.37 (t, *J* = 8.0 Hz, 2H), 3.23 – 3.14 (m, 2H), 3.05 (q, *J* = 6.5 Hz, 2H), 2.95 (t, *J* = 12.4 Hz, 2H), 2.69 (d, *J* = 4.6 Hz, 3H), 2.45 (s, 3H), 2.43 – 2.40 (m, 1H), 2.28 (s, 3H), 1.74 (q, *J* = 7.0 Hz, 4H), 1.54 (td, *J* = 14.2, 6.8 Hz, 2H), 1.46 – 1.38 (m, 4H), 1.36 – 1.25 (m, 10H). $^{19}$**F NMR** (471 MHz, DMSO-$d_6$) δ = -74.3.

Synthesis of **RCS-02-121:**

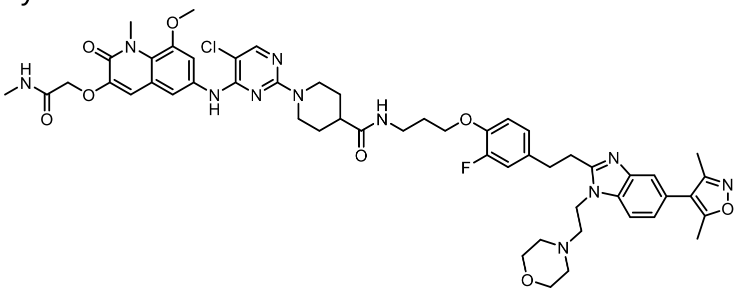

The corresponding compound was prepared following General Procedure B using *tert*-butyl (3-bromopropyl)carbamate as the linker. Purification by preparative reverse-phase HPLC (15-90% MeOH in water) afforded 1-(5-chloro-4-((8-methoxy-1-methyl-3-(2-(methylamino)-2-oxoethoxy)-2-oxo-1,2-dihydroquinolin-6-yl)amino)pyrimidin-2-yl)-N-(3-(4-(2-(5-(3,5-dimethylisoxazol-4-yl)-1-(2-morpholinoethyl)-1H-benzo[d]imidazol-2-yl)ethyl)-2-fluorophenoxy)propyl)piperidine-4-carboxamide (3.5 mg, 3.4 µmol, 20%, MS m/z 1034.50 $[M+H]^+$). **$^1$H NMR** (500 MHz, DMSO-$d_6$) δ = 8.95 (s, 1H), 8.09 (s, 1H), 7.96 (d, *J* = 4.7 Hz, 1H), 7.92 (d, *J* = 5.6 Hz, 1H), 7.87 (d, *J* = 8.5 Hz, 1H), 7.71 (d, *J* = 1.6 Hz, 1H), 7.57 – 7.48 (m, 2H), 7.41 (dd, *J* = 8.3, 1.6 Hz, 1H), 7.27 – 7.20 (m, 1H), 7.11 – 7.04 (m, 2H), 6.99 (s, 1H), 4.71 (t, *J* = 8.0 Hz, 2H), 4.54 (s, 2H), 4.49 (d, *J* = 13.1 Hz, 2H), 4.02 (t, *J* = 6.2 Hz, 2H), 3.86 (s, 3H), 3.85 (s, 3H), 3.44 (s, 2H), 3.36 – 3.30 (m, 2H), 3.19 (q, *J* = 6.6 Hz, 2H), 3.14 (t, *J* = 8.0 Hz, 2H), 2.91 (t, *J* = 12.7 Hz, 2H), 2.64 (d, *J* = 4.6 Hz, 3H), 2.41 (s, 3H), 2.40 – 2.35 (m, 1H), 2.23 (s, 3H), 1.84 (p, *J* = 6.6 Hz, 2H), 1.72 (d, *J* = 12.8 Hz, 2H), 1.55 – 1.44 (m, 2H). **$^{19}$F NMR** (471 MHz, DMSO-$d_6$) δ = -74.4.

Synthesis of **RCS-02-122:**

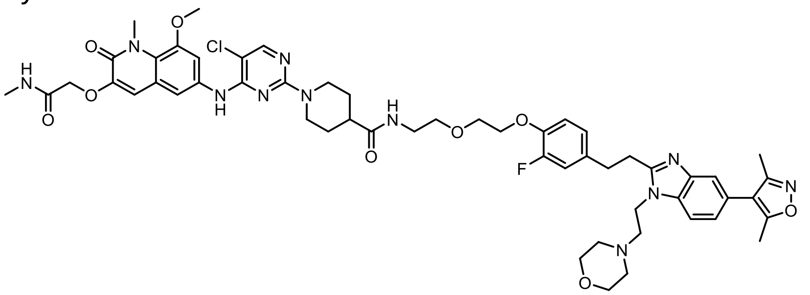

The corresponding compound was prepared following General Procedure B using *tert*-butyl (2-(2-bromoethoxy)ethyl)carbamate as the linker. Purification by preparative reverse-phase HPLC (15-90% MeOH in water) afforded 1-(5-chloro-4-((8-methoxy-1-methyl-3-(2-(methylamino)-2-oxoethoxy)-2-oxo-1,2-dihydroquinolin-6-yl)amino)pyrimidin-2-yl)-N-(2-(2-(4-(2-(5-(3,5-dimethylisoxazol-4-yl)-1-(2-morpholinoethyl)-1H-benzo[d]imidazol-2-yl)ethyl)-2-fluorophenoxy)ethoxy)ethyl)piperidine-4-carboxamide (5.3 mg, 5.0 µmol, 29%, MS m/z 1064.51 $[M+H]^+$). **$^1$H NMR** (500 MHz, DMSO-$d_6$) δ = 8.96 (s, 1H), 8.09 (s, 1H), 7.95 (d, *J* = 4.7 Hz, 1H), 7.91 – 7.85 (m, 2H), 7.71 (d, *J* = 1.5 Hz, 1H), 7.52 (q, *J* = 2.3 Hz, 2H), 7.45 – 7.39 (m, 1H), 7.22 (dd, *J* = 12.6, 1.9 Hz, 1H), 7.14 – 7.04 (m, 2H), 7.00 (s, 1H), 4.70 (d, *J* = 8.2 Hz, 2H), 4.54 (s, 2H), 4.48 (d, *J* = 13.1 Hz, 2H), 4.15 – 4.10 (m, 2H), 3.86 (s, 3H), 3.85 (s, 3H), 3.75 – 3.70 (m, 2H), 3.47 (t, *J* = 6.0 Hz, 2H), 3.33 (t, *J* = 8.0 Hz, 2H), 3.21 (q, *J* = 5.9 Hz, 2H), 3.14 (t, *J* = 8.0 Hz, 2H), 2.91 (t, *J* = 12.3 Hz, 2H), 2.64 (d, *J* = 4.7 Hz, 3H), 2.44-2.39 (m, 1H, assigned by HSQC), 2.41 (s, 3H), 2.23 (s, 3H), 1.70 (d, *J* = 12.6 Hz, 2H), 1.54 – 1.43 (m, 2H). **$^{19}$F NMR** (471 MHz, DMSO-$d_6$) δ = -74.5.

Synthesis of **RCS-02-123:**

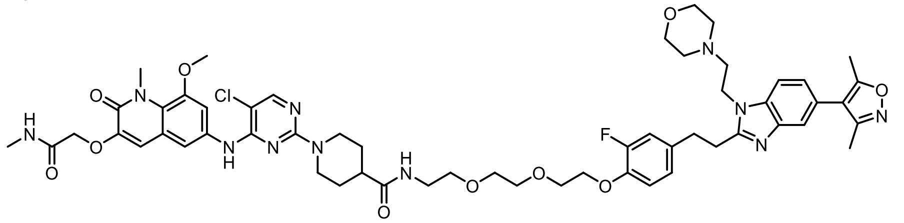

The corresponding compound was prepared following General Procedure B using *tert*-butyl (2-(2-(2-bromoethoxy)ethoxy)ethyl)carbamate as the linker. Purification by preparative reverse-phase HPLC (15-90% MeOH in water) afforded 1-(5-chloro-4-((8-methoxy-1-methyl-3-(2-(methylamino)-2-oxoethoxy)-2-oxo-1,2-dihydroquinolin-6-yl)amino)pyrimidin-2-yl)-N-(2-(2-(2-(4-(2-(5-(3,5-dimethylisoxazol-4-yl)-1-(2-morpholinoethyl)-1H-benzo[d]imidazol-2-yl)ethyl)-2-fluorophenoxy)ethoxy)ethoxy)ethyl)piperidine-4-carboxamide (10 mg, 9.0 µmol, 52%, MS m/z 1108.52 $[M+H]^+$). **$^1$H NMR** (500 MHz, DMSO-$d_6$) δ = 9.12 (s, 1H), 8.12 (s, 1H), 7.96 (d, *J* = 8.1 Hz, 2H), 7.88 (t, *J* = 5.5 Hz, 1H), 7.76 (d, *J* = 1.5 Hz, 1H), 7.53 – 7.51 (m, 2H), 7.49 (d, *J* = 8.5 Hz, 1H), 7.28 – 7.20 (m, 1H), 7.15 – 7.06 (m, 2H), 7.02 (s, 1H), 4.78 (s, 2H), 4.55 (s, 2H), 4.45 (d, *J* = 13.0 Hz, 2H), 4.13 (t, *J* = 4.6 Hz, 2H), 3.87 (s, 3H), 3.85 (s, 3H), 3.76 – 3.73 (m, 2H), 3.59 (dd, *J* = 6.0, 3.5 Hz, 2H), 3.54 – 3.47 (m, 4H), 3.40 (q, *J* = 7.2 Hz, 4H), 3.23 – 3.13 (m, 4H), 2.94 (t, *J* = 12.6 Hz, 2H), 2.65 (d, *J* = 4.7 Hz, 3H), 2.44 – 2.41 (m, 1H, assigned by HSQC), 2.42 (s, 3H), 2.25 (s, 3H), 1.72 (d, *J* = 12.7 Hz, 2H), 1.57 – 1.47 (m, 2H). **$^{19}$F NMR** (471 MHz, DMSO-$d_6$) δ = -74.8.

Synthesis of **RCS-02-124:**

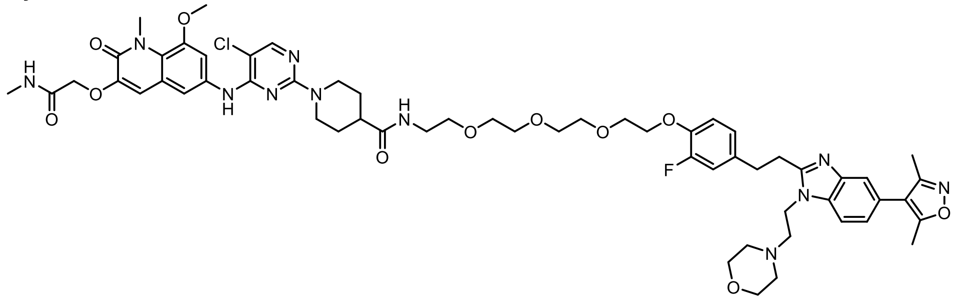

The corresponding compound was prepared following General Procedure B using *tert*-butyl (2-(2-(2-(2-bromoethoxy)ethoxy)ethoxy)ethyl)carbamate as the linker. Purification by preparative reverse-phase HPLC (15-90% MeOH in water) afforded 1-(5-chloro-4-((8-methoxy-1-methyl-3-(2-(methylamino)-2-oxoethoxy)-2-oxo-1,2-dihydroquinolin-6-yl)amino)pyrimidin-2-yl)-N-(2-(2-(2-(2-(4-(2-(5-(3,5-dimethylisoxazol-4-yl)-1-(2-morpholinoethyl)-1H-benzo[d]imidazol-2-yl)ethyl)-2-fluorophenoxy)ethoxy)ethoxy)ethoxy)ethyl)piperidine-4-carboxamide (5.8 mg, 5.0 µmol, 29%, MS m/z 1152.65 $[M+H]^+$). **$^1$H NMR** (500 MHz, DMSO-$d_6$) δ = 8.97 (s, 1H), 8.08 (s, 1H), 7.96 (d, *J* = 4.8 Hz, 1H), 7.91 – 7.84 (m, 2H), 7.71 (d, *J* = 1.5 Hz, 1H), 7.52 (t, *J* = 1.8 Hz, 2H), 7.42 (dd, *J* = 8.3, 1.6 Hz, 1H), 7.23 (dd, *J* = 11.9, 2.4 Hz, 1H), 7.13 – 7.05 (m, 2H), 7.00 (s, 1H), 4.71 (t, *J* = 7.9 Hz, 2H), 4.54 (s, 2H), 4.47 (d, *J* = 13.1 Hz, 2H), 4.14 – 4.10 (m, 2H), 3.87 (s, 3H), 3.85 (s, 3H), 3.74 – 3.71 (m, 2H), 3.57 (dd, *J* = 6.1, 3.6 Hz, 2H), 3.54 – 3.47 (m, 8H), 3.39 (t, *J* = 6.0 Hz, 2H), 3.34 (t, *J* = 8.0 Hz, 2H), 3.21 – 3.12 (m, 4H), 2.91 (t, *J* = 12.4 Hz, 2H), 2.64 (d, *J* = 4.6 Hz, 3H), 2.43 – 2.38 (m, 1H, assigned by HSQC), 2.41 (s, 3H), 2.23 (s, 3H), 1.70 (d, *J* = 12.8 Hz, 2H), 1.54 – 1.43 (m, 2H). **$^{19}$F NMR** (471 MHz, DMSO-$d_6$) δ = -74.5.

**Synthesis of CCT373566a:**

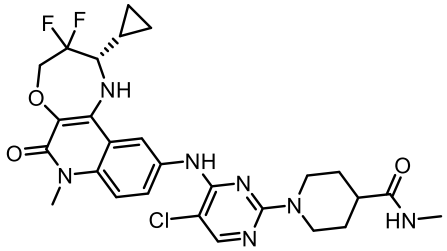

To a solution of **CCT373566-COOH** (10.0 mg, 18 µmol, 1.0 equiv) in DMF (0.360 mL, 0.05 M), TEA (0.017 mL, 120 µmol, 7.0 equiv), and HATU (8.1 mg, 21 µmol, 1.2 equiv) was added methanamine (0.72 mg, 23 µmol, 1.3 equiv) at 0 ˚C. The reaction was allowed to stir for two hours, at which point it was purified by preparative reverse-phase HPLC (15-90% MeOH in water) to afford (S)-1-(5-chloro-4-((2-cyclopropyl-3,3-difluoro-7-methyl-6-oxo-1,2,3,4,6,7-hexahydro-[1,4]oxazepino[2,3-c]quinolin-10-yl)amino)pyrimidin-2-yl)-N-methylpiperidine-4-carboxamide (5.6 mg, 180 µmol, 55%, MS m/z 574.03 $[M+H]^+$). **$^1$H NMR** (500 MHz, DMSO-$d_6$) δ = 9.23 (s, 1H), 8.18 (d, *J* = 2.3 Hz, 1H), 8.12 (s, 1H), 7.75 – 7.68 (m, 2H), 7.47 (s, 1H), 7.45 (s, 1H), 6.21 (s, 1H), 4.50 – 4.40 (m, 2H, as determined by HSQC), 4.40 – 4.32 (m, 3H, as determined by HSQX), 3.57 (s, 3H), 3.23 (dd, *J* = 11.0, 6.1 Hz, 1H), 2.86 (t, *J* = 12.6 Hz, 2H), 2.55 (d, *J* = 4.6 Hz, 3H), 2.34 (ddd, *J* = 15.5, 11.2, 3.7 Hz, 1H), 1.66 (d, *J* = 12.9 Hz, 2H), 1.54 – 1.41 (m, 2H), 1.32 (q, *J* = 4.4 Hz, 1H), 0.75 – 0.66 (m, 1H), 0.55 – 0.47 (m, 2H), 0.34 (q, *J* = 5.9 Hz, 1H). **$^{19}$F NMR** (471 MHz, DMSO-$d_6$) δ = -100.23.

**$^{1}$H NMR of GNE-781-NH:**

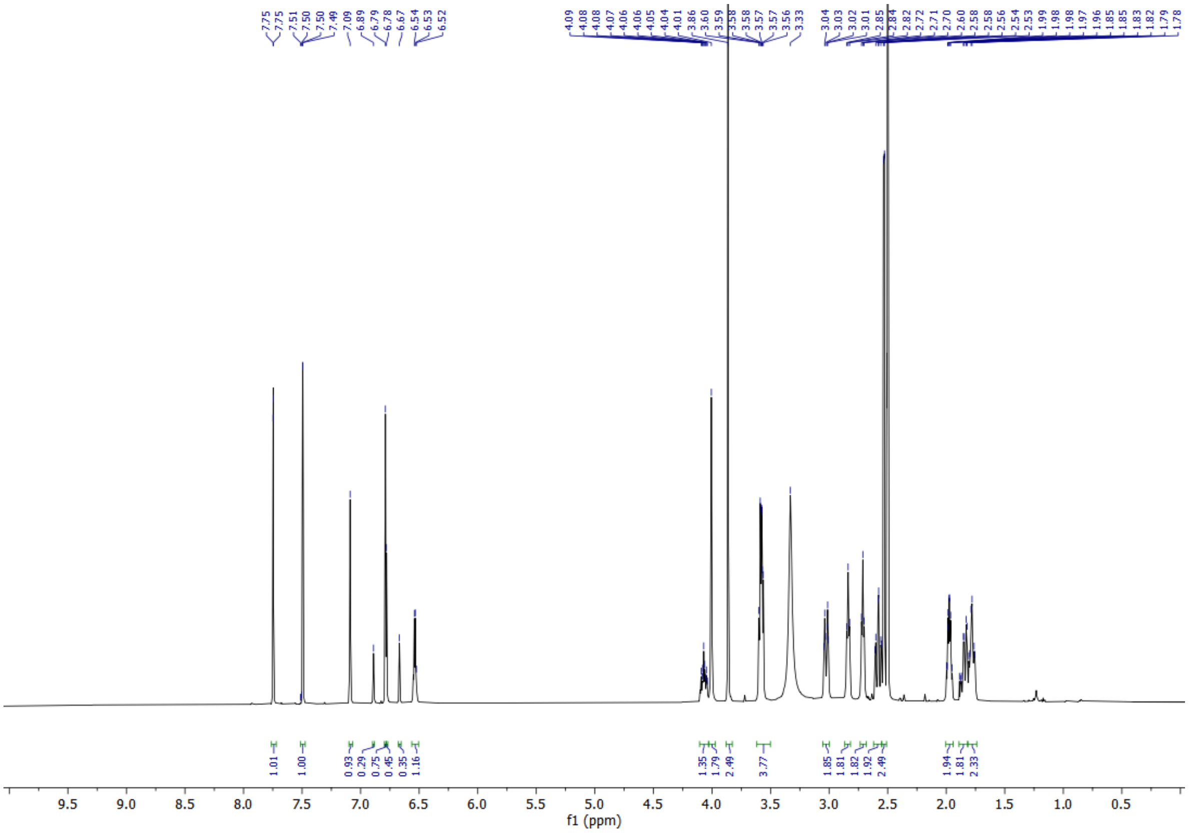

**$^{1}$H NMR of TCIP3:**

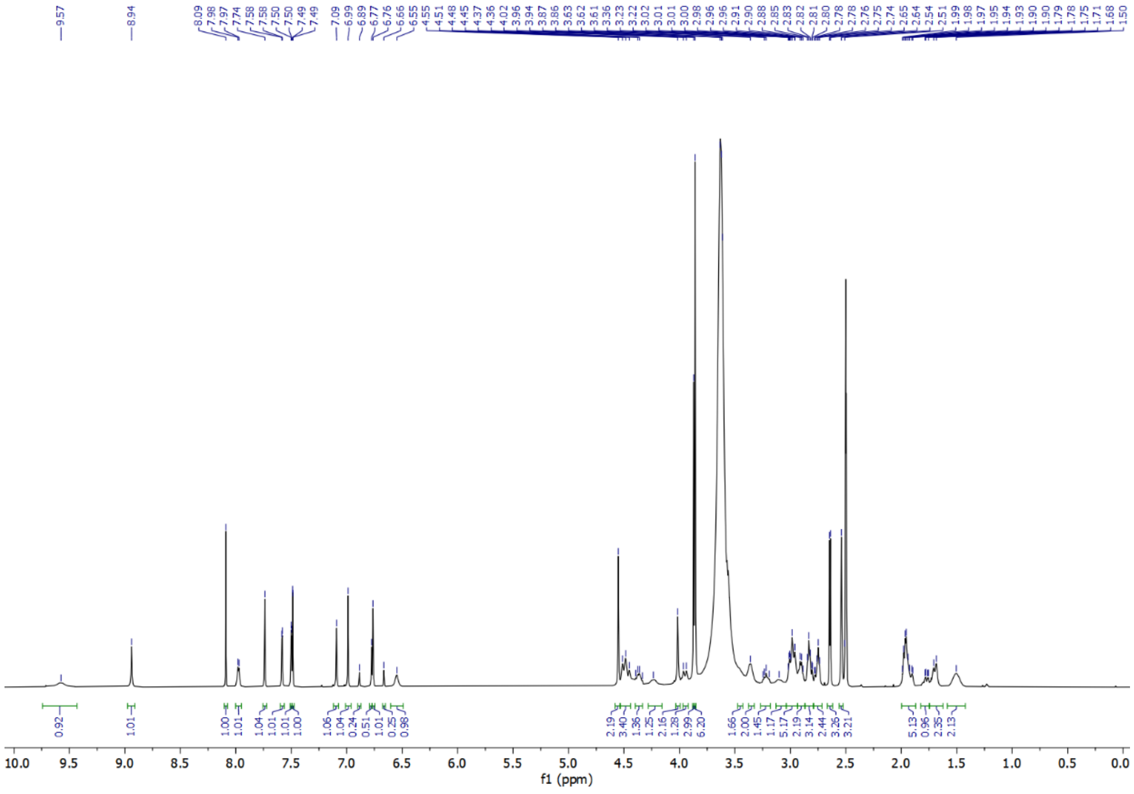

**$^1$H NMR of MNN-02-155:**

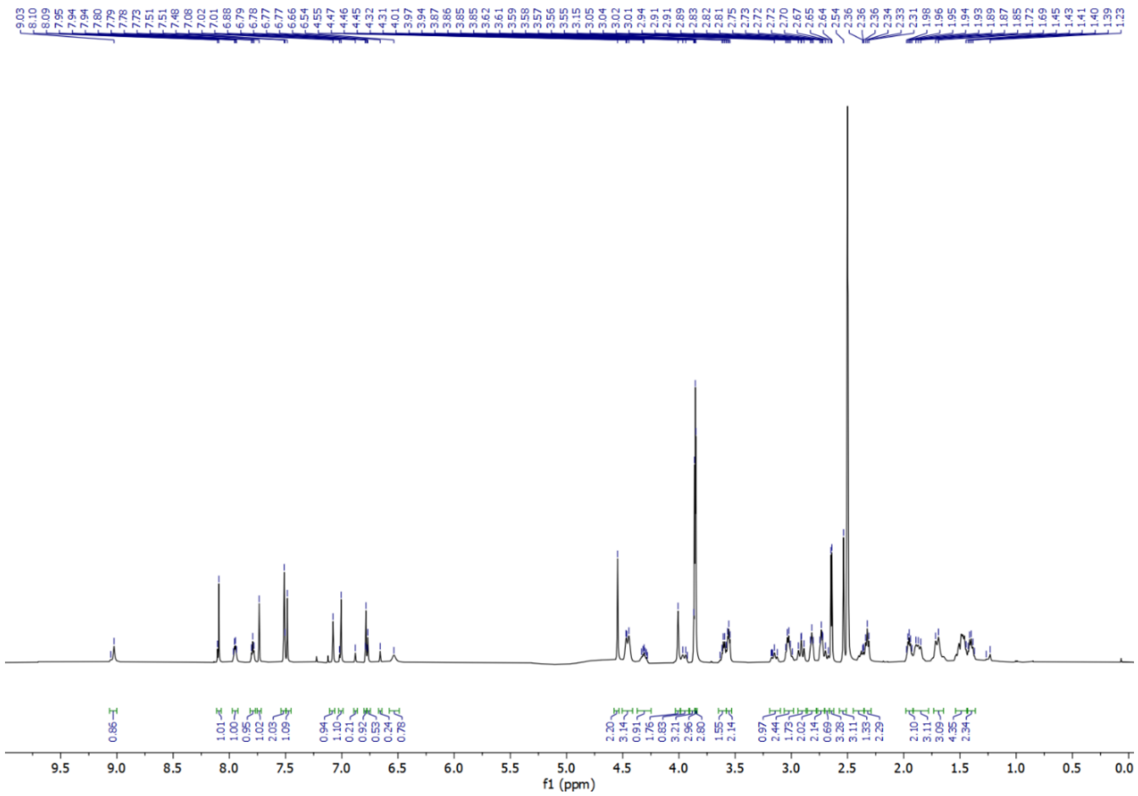

**$^1$H NMR of MNN-03-037:**

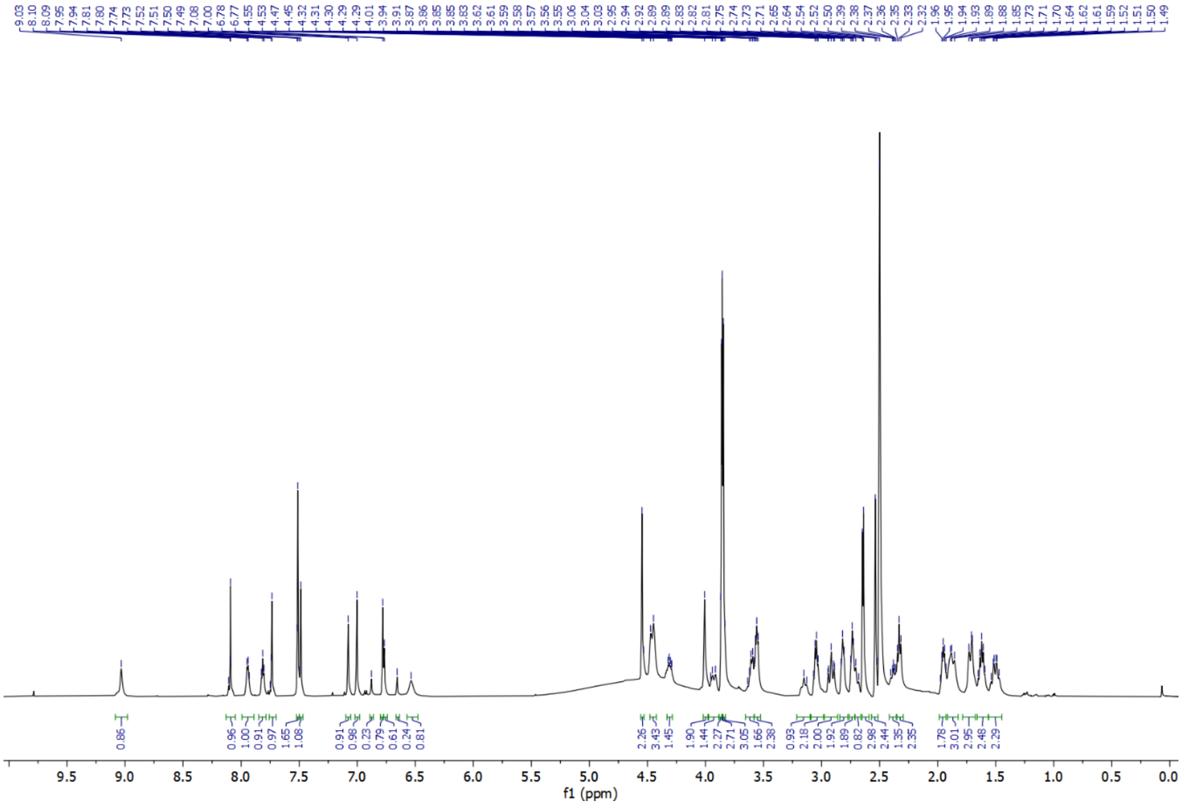

**$^1$H NMR of MNN-02-195:**

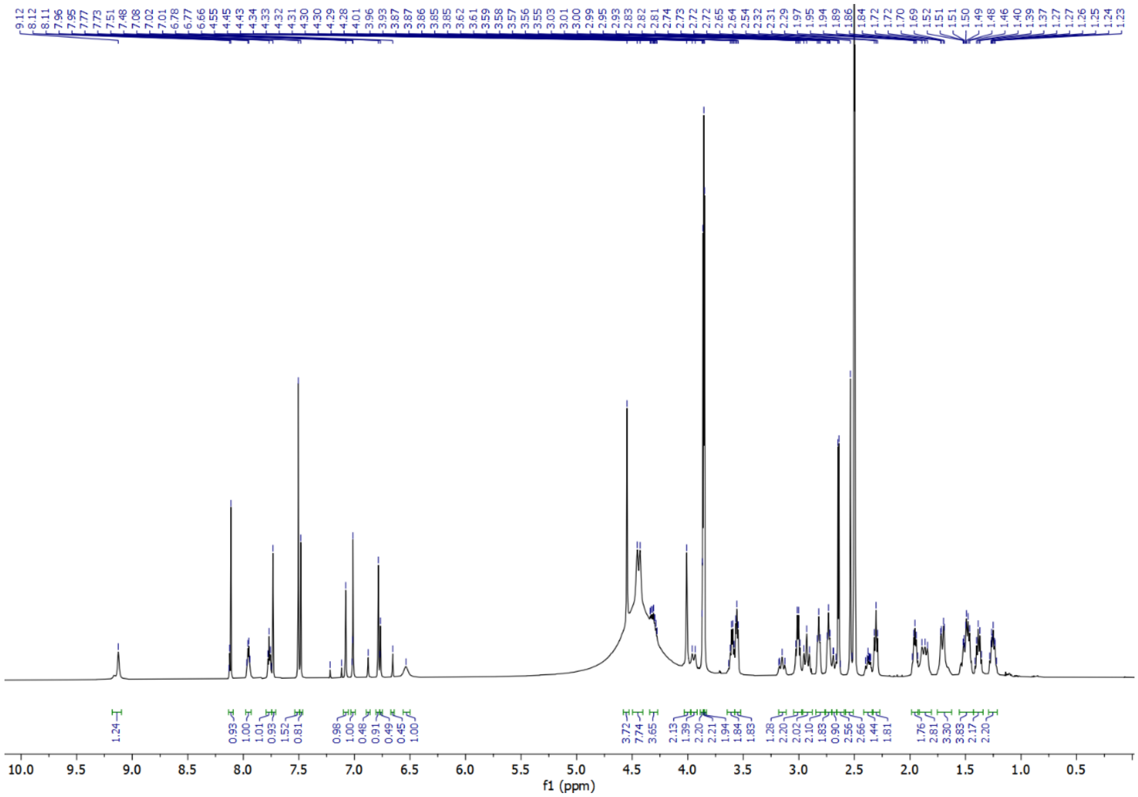

**$^1$H NMR of RCS-IJD-001:**

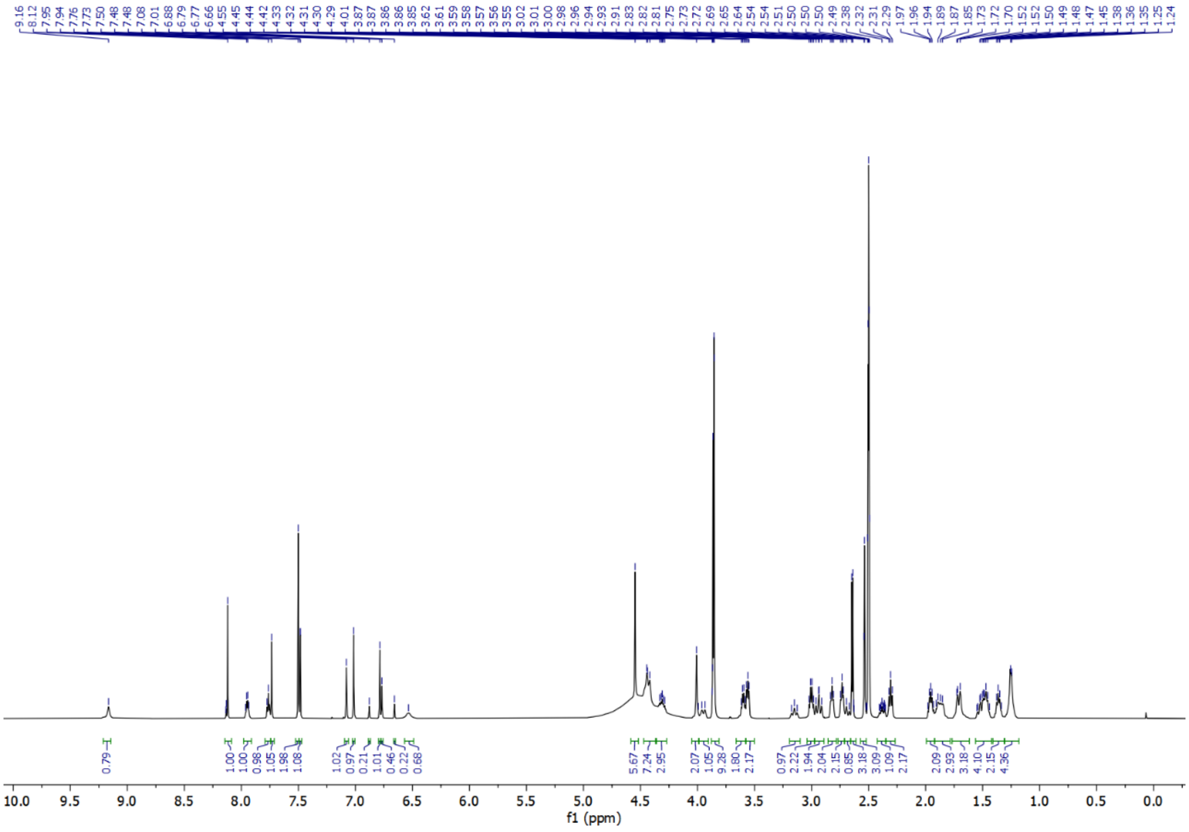

**$^{1}$H NMR of MNN-02-196:**

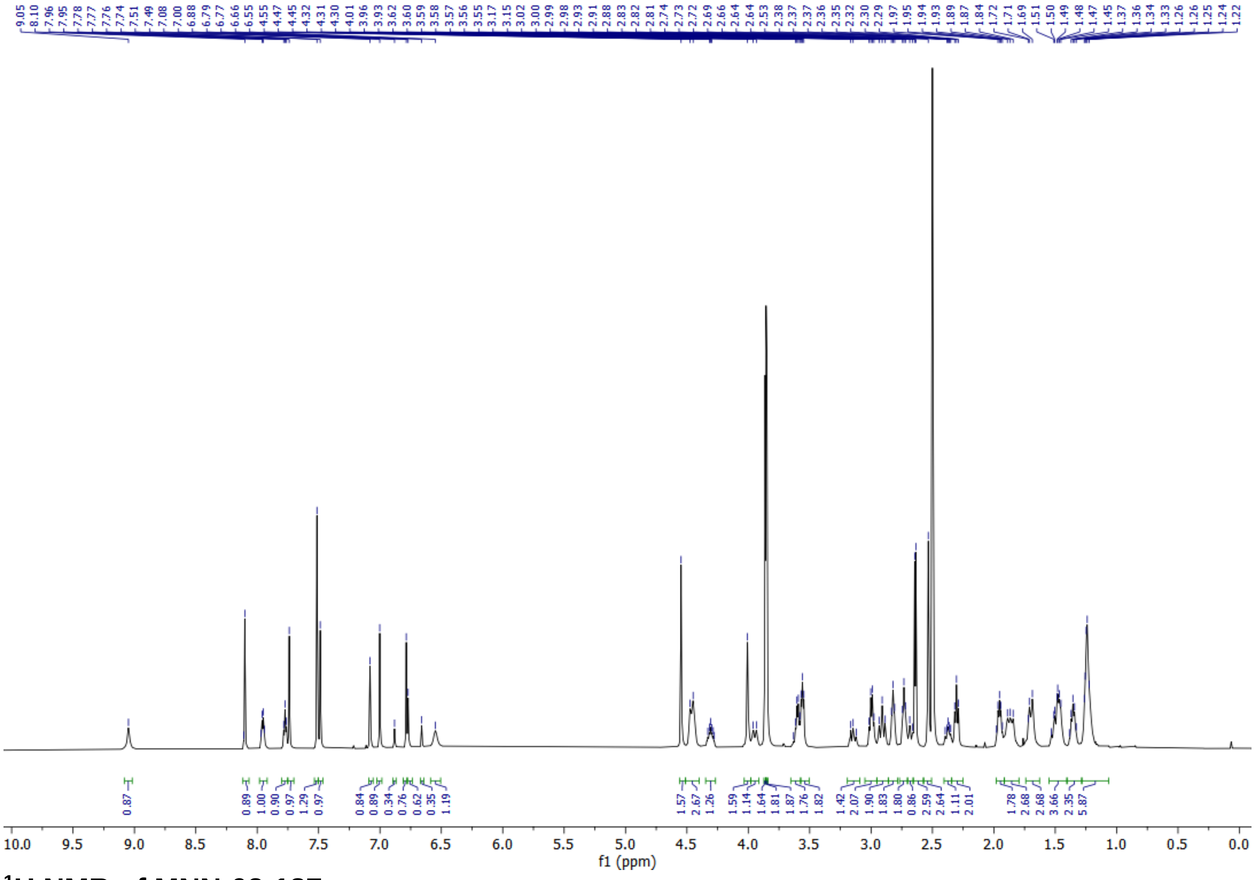

**$^{1}$H NMR of MNN-02-187:**

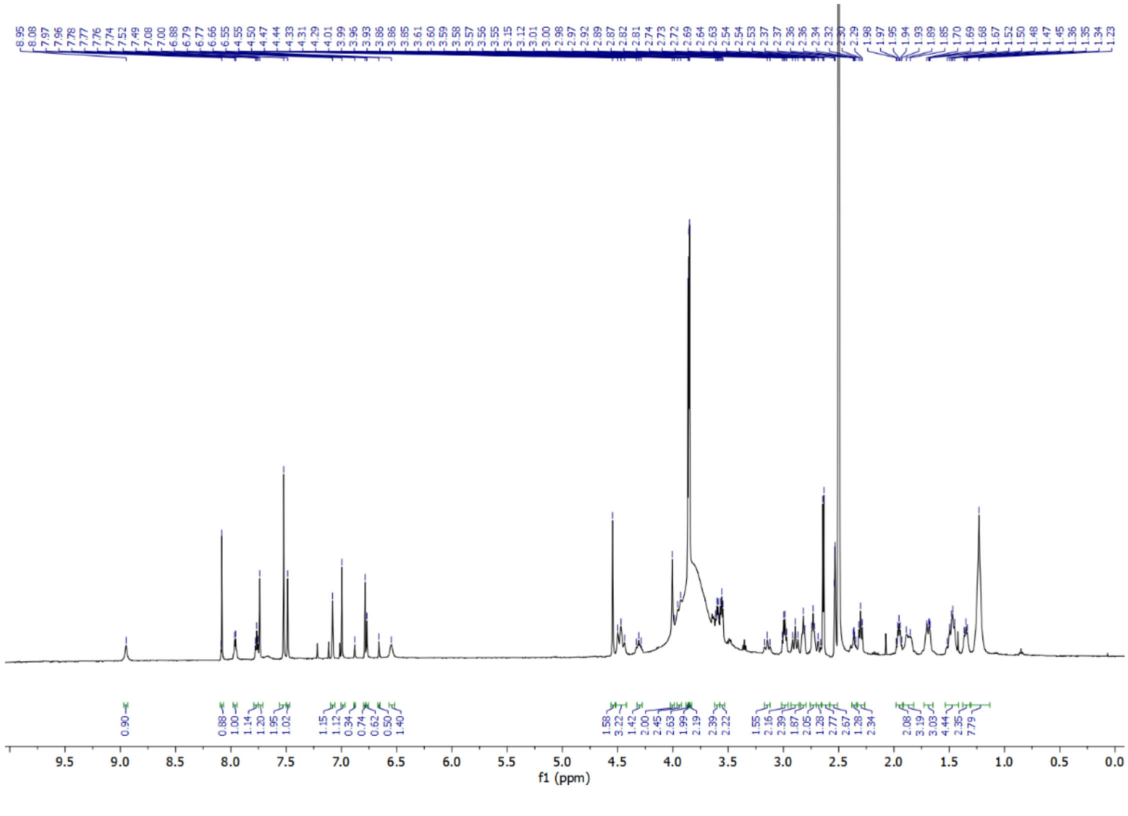

**$^1$H NMR of MNN-02-197:**

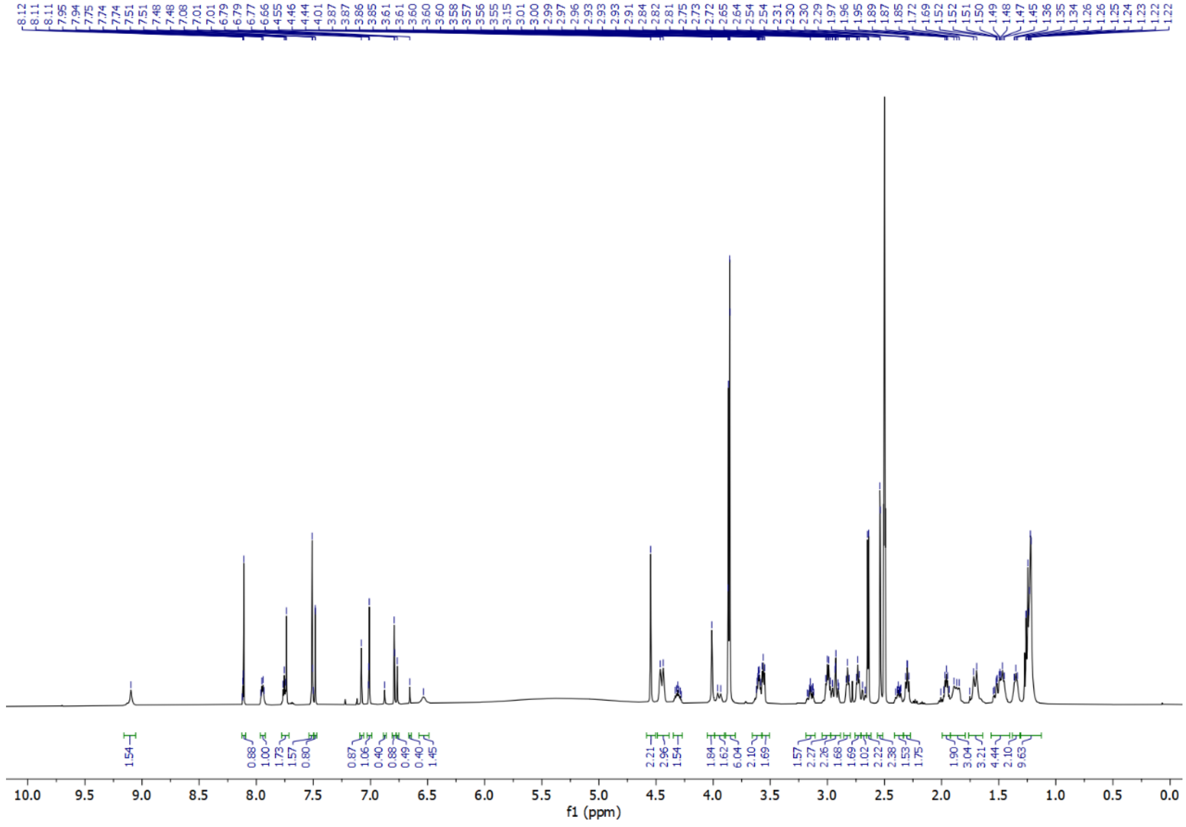

**$^1$H NMR of MNN-02-160:**

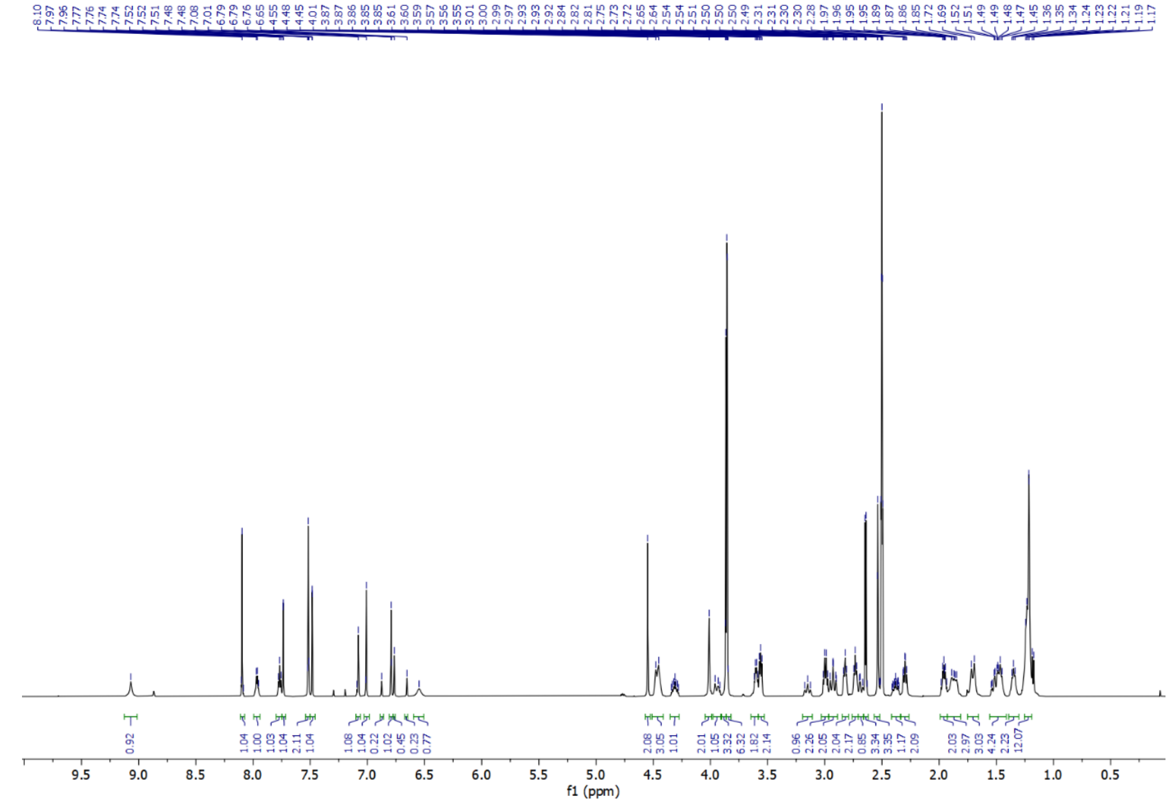

**[1]H NMR of MNN-02-161:**

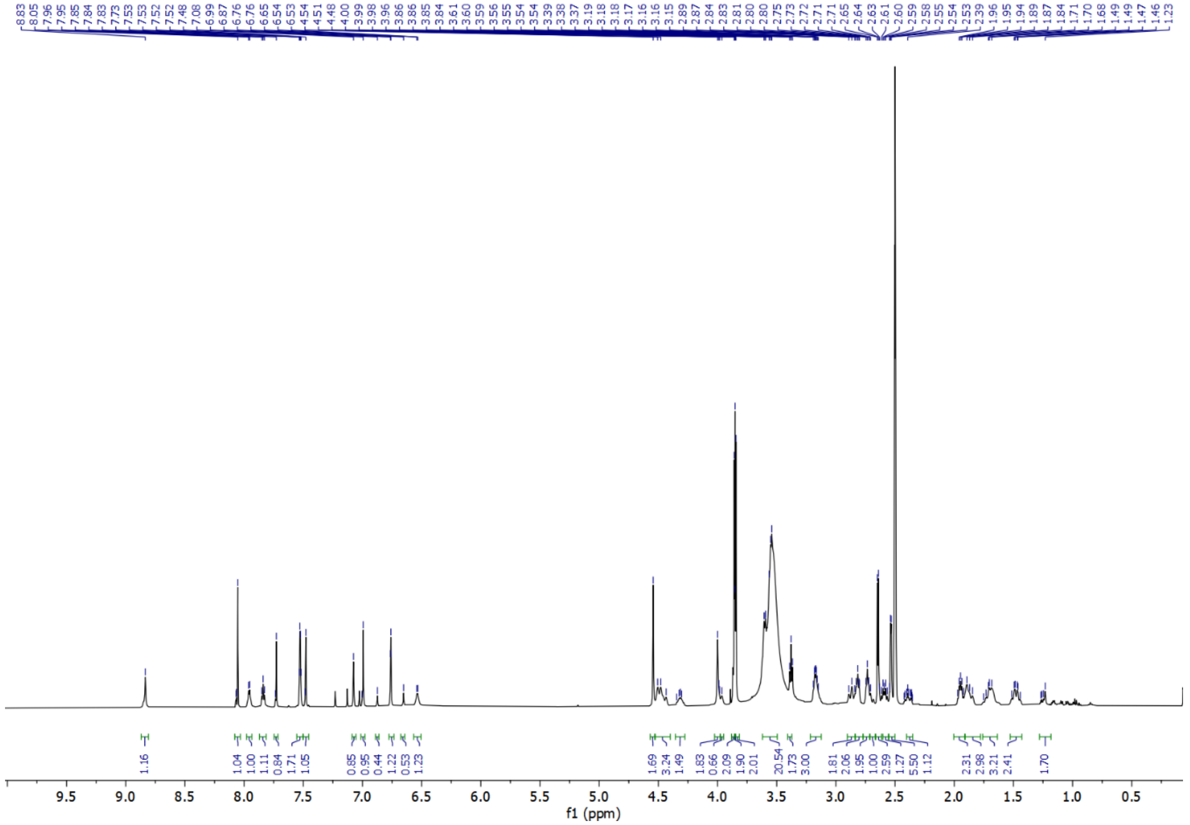

**[1]H NMR of MNN-02-162:**

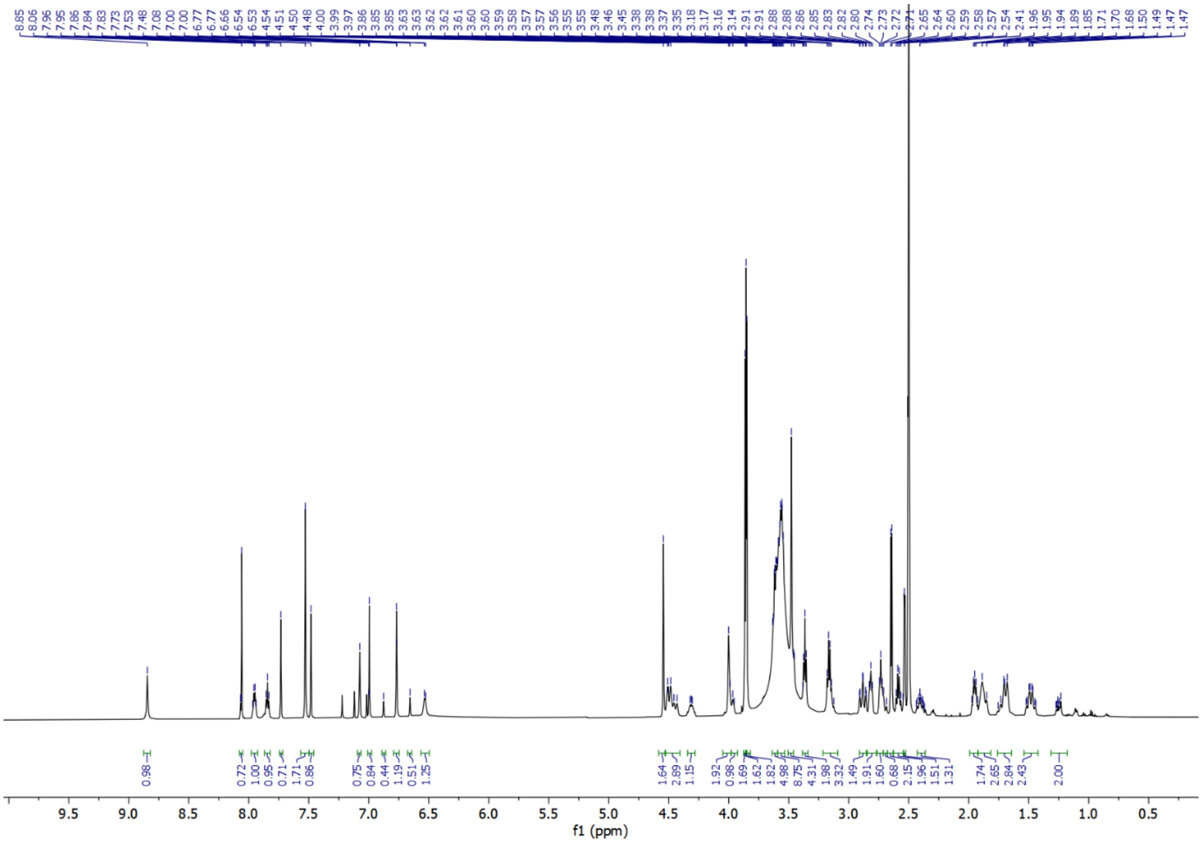

**$^{1}$H NMR of MNN-02-156:**

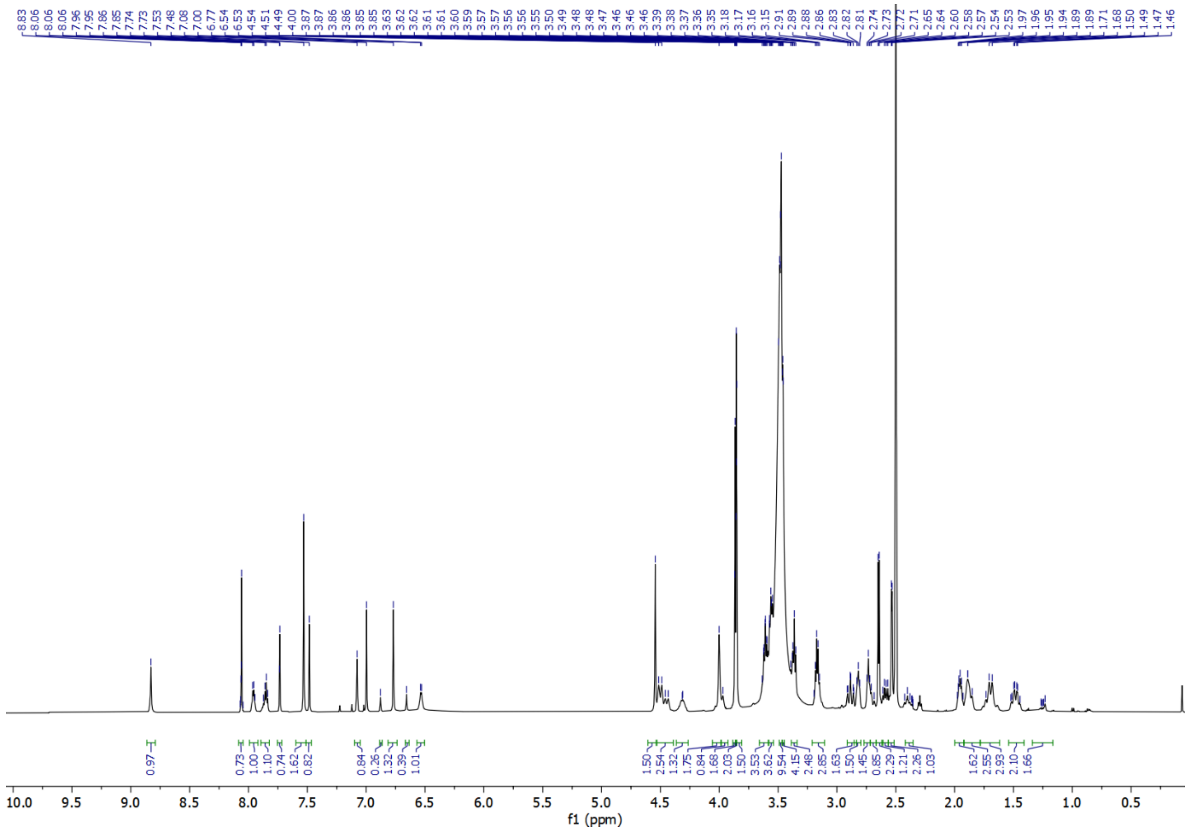

**$^{1}$H NMR of MNN-03-050:**

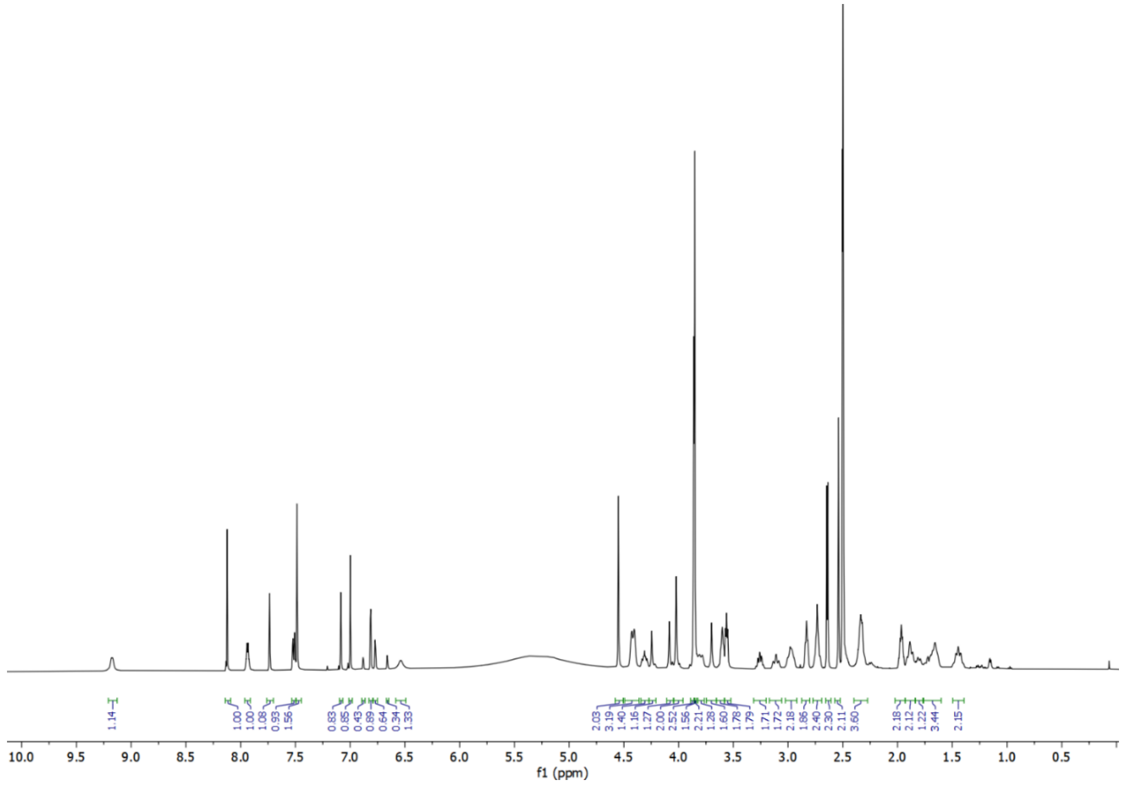

[1]H NMR of MNN-03-041:

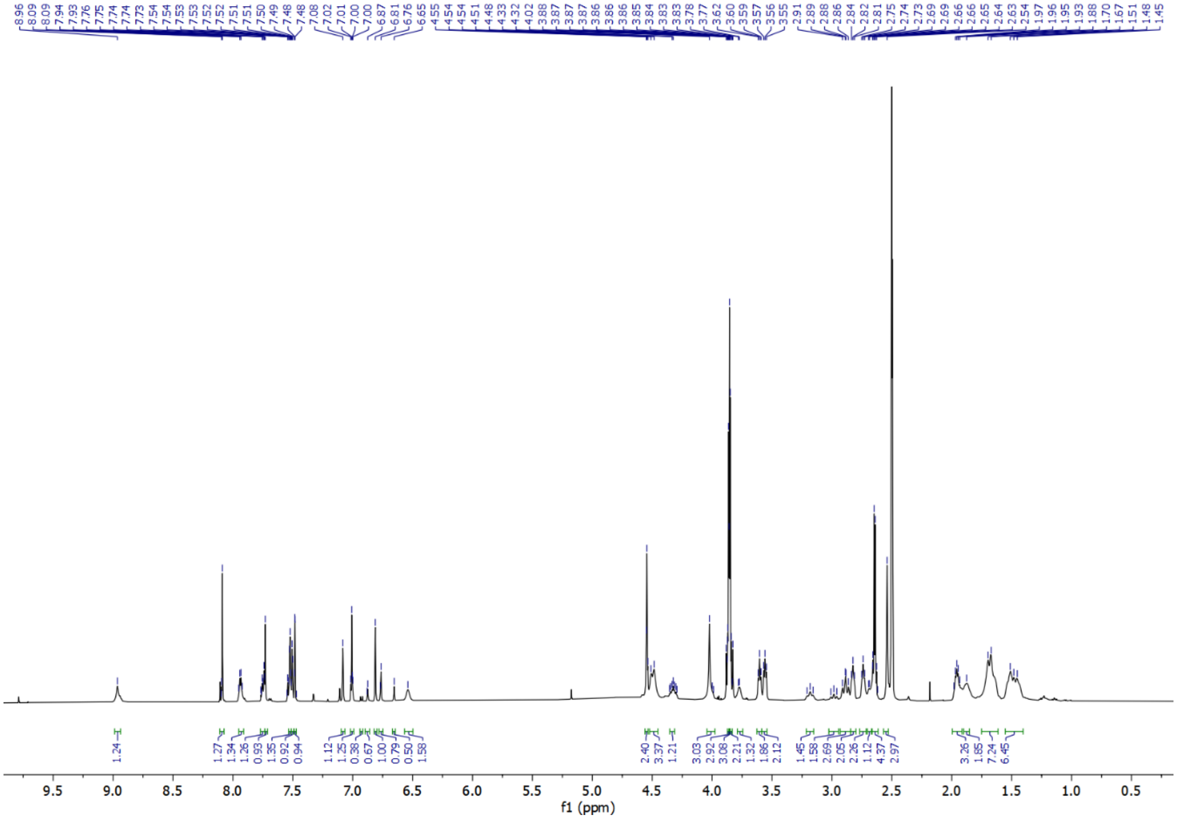

[1]H NMR of MNN-03-040:

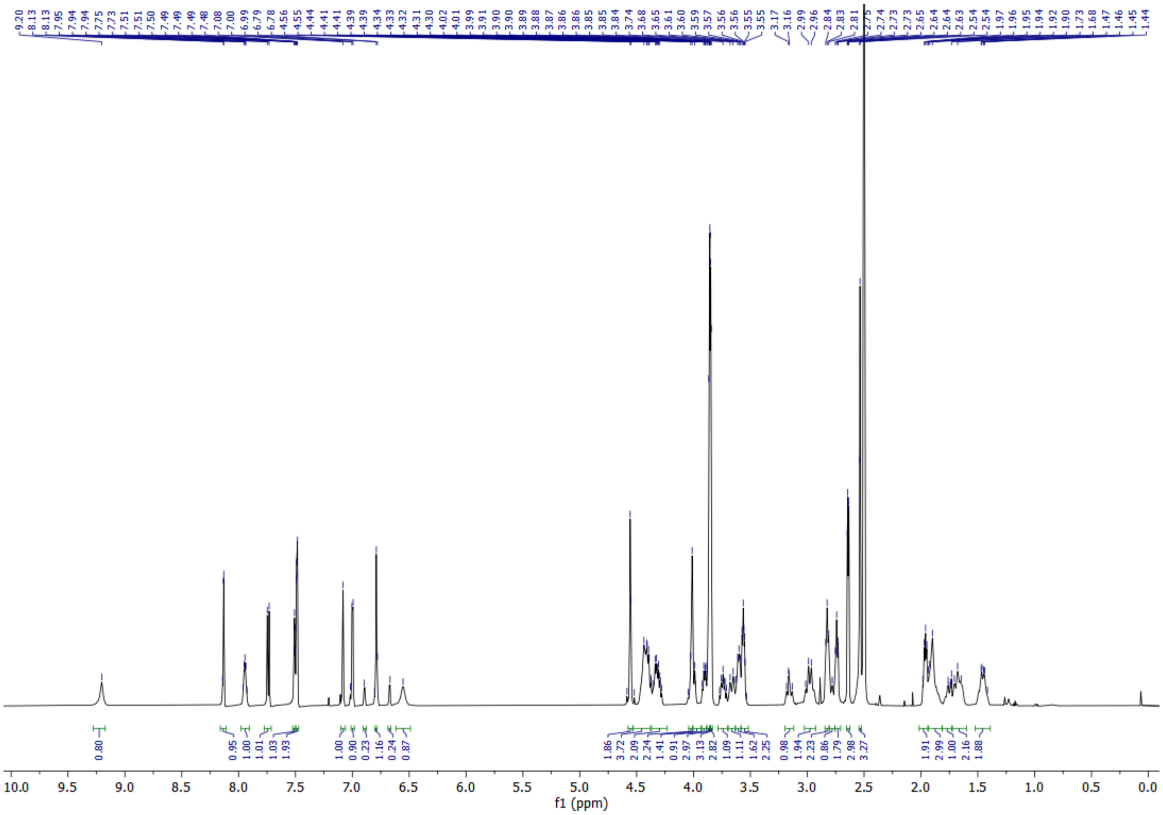

**[1]H NMR of MNN-03-049:**

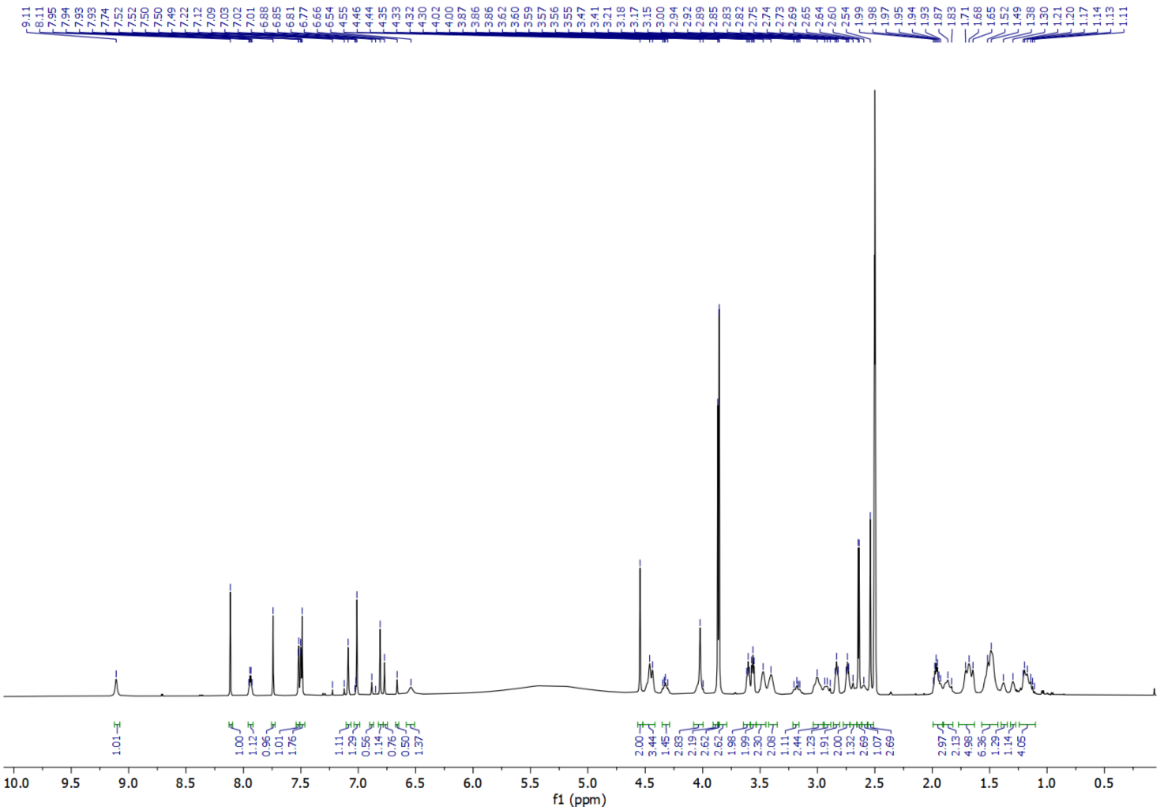

**[1]H NMR of MNN-03-039:**

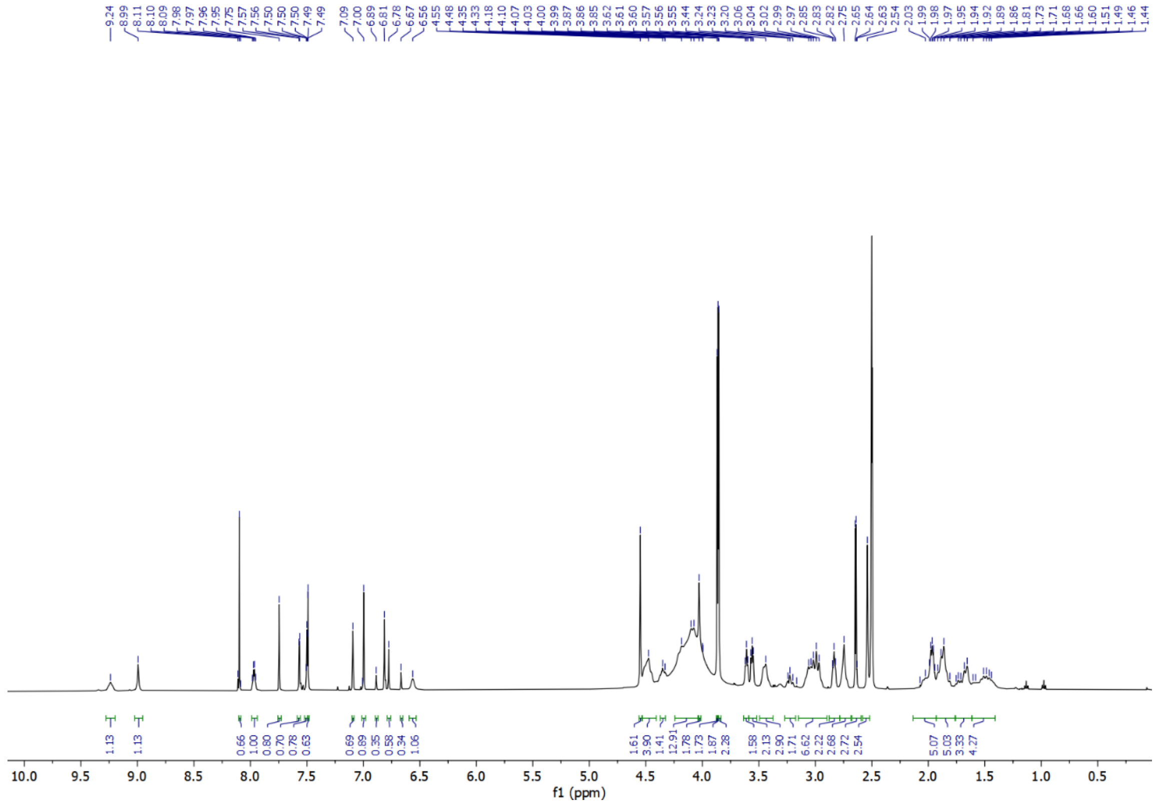

**$^1$H NMR of NEG1:**

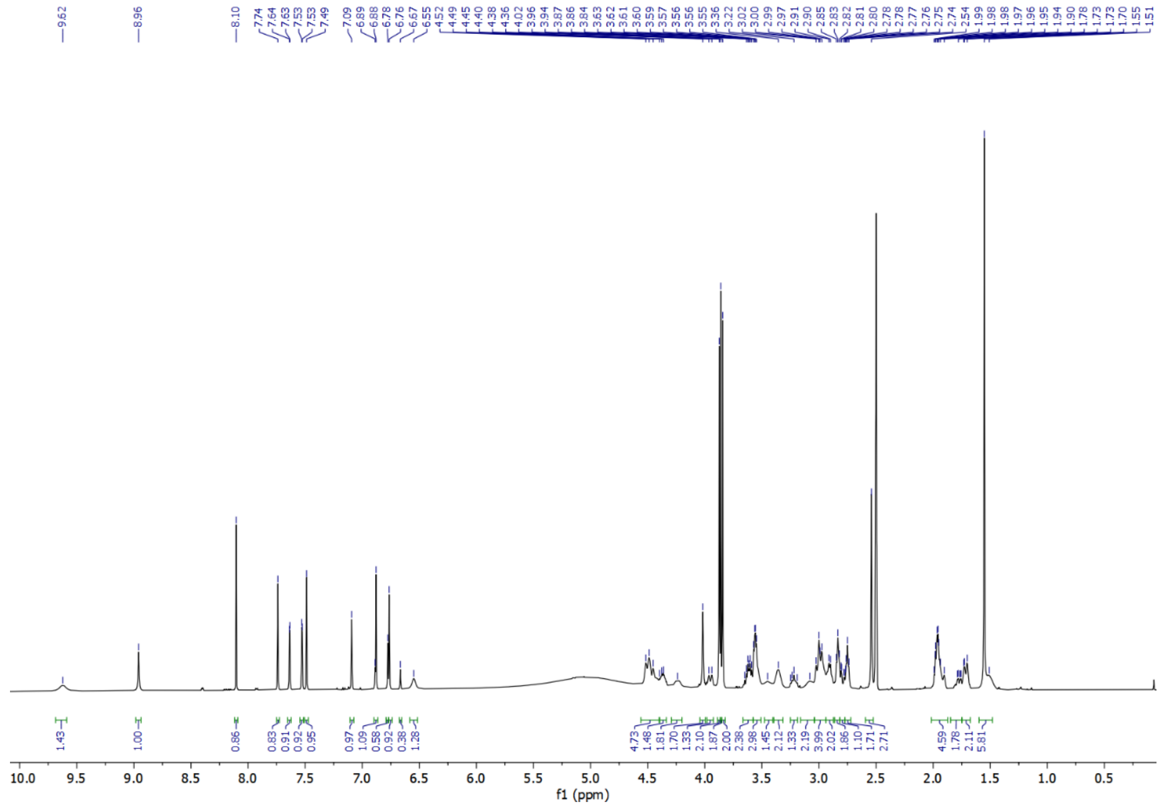

**$^1$H NMR of NEG2:**

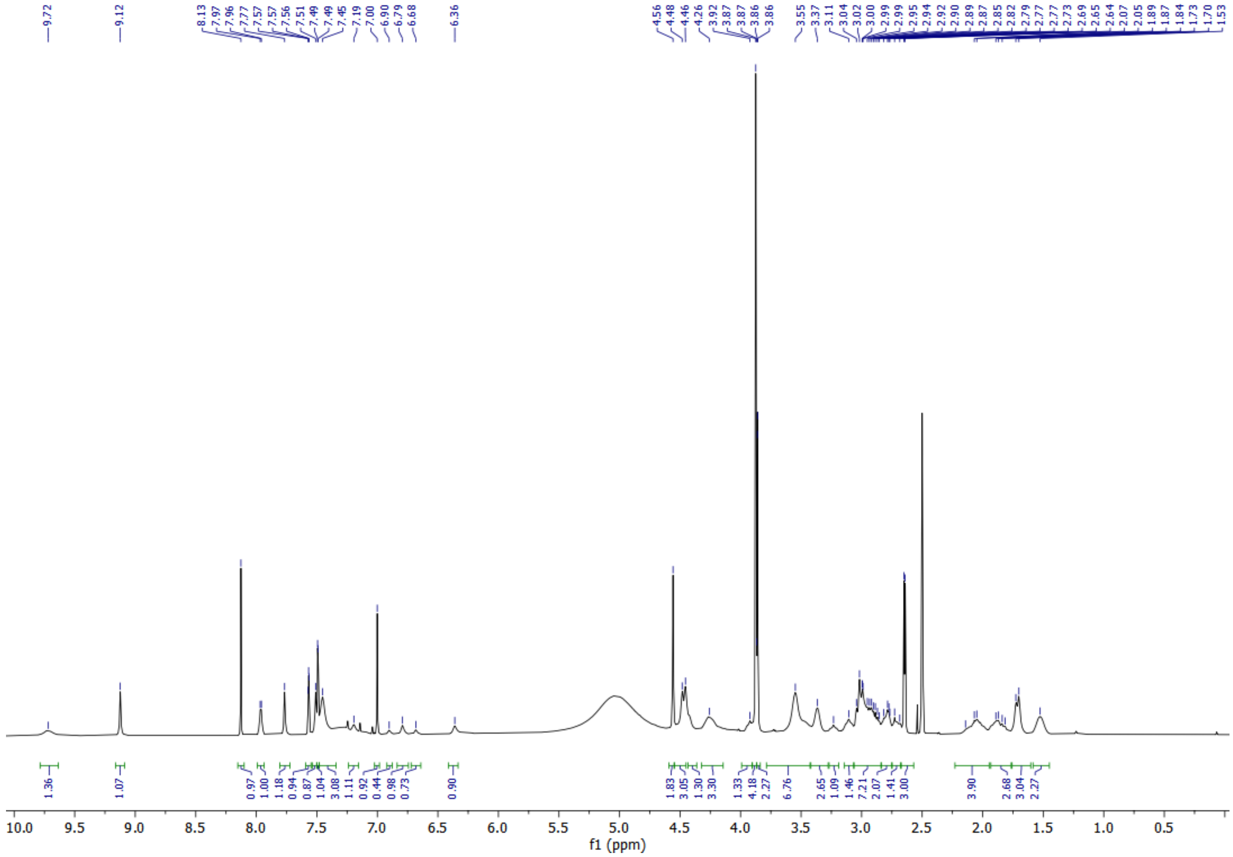

**[1]H NMR of MNN-06-112:**

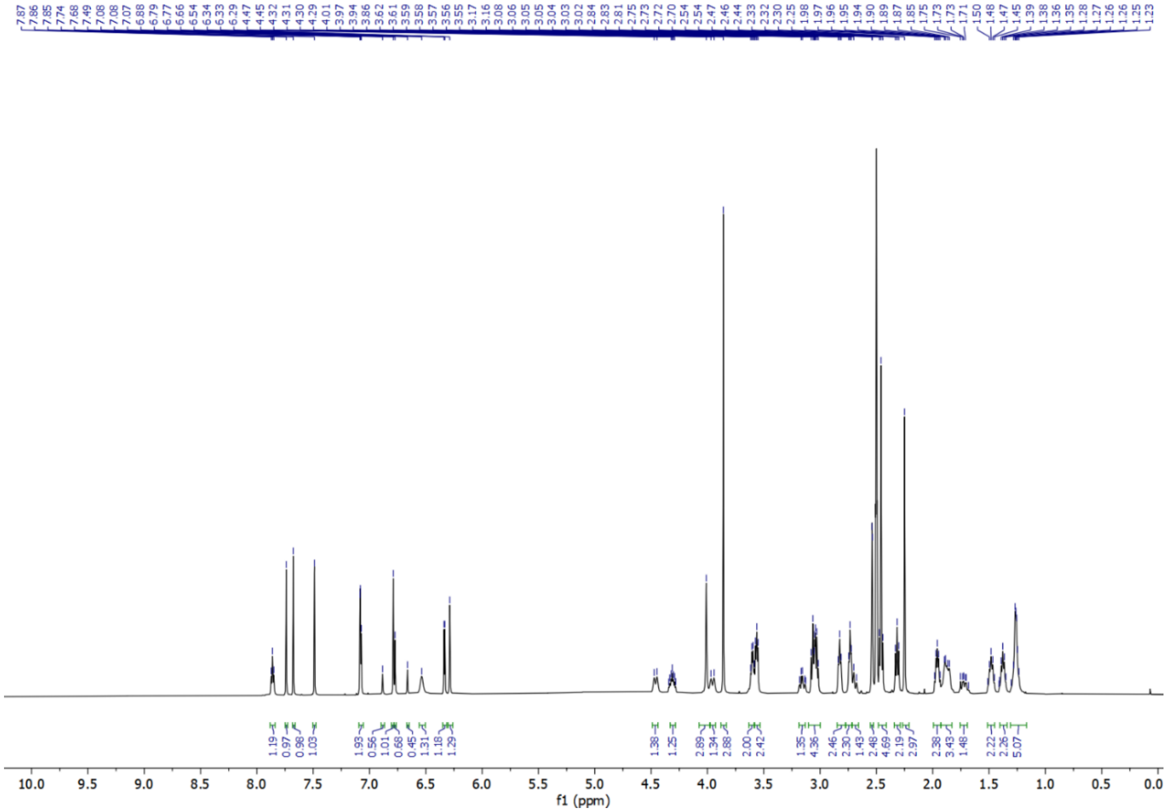

**[1]H NMR of TNL15:**

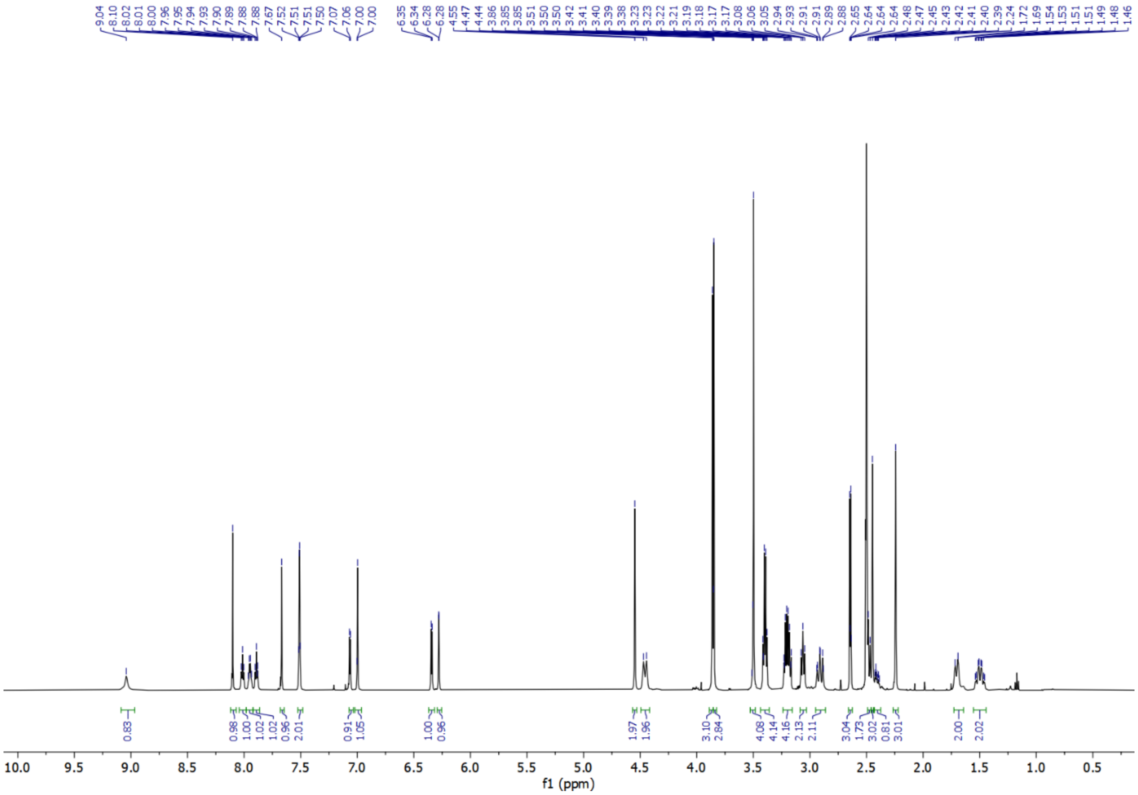

### $^{1}H$ NMR of RCS-02-107:

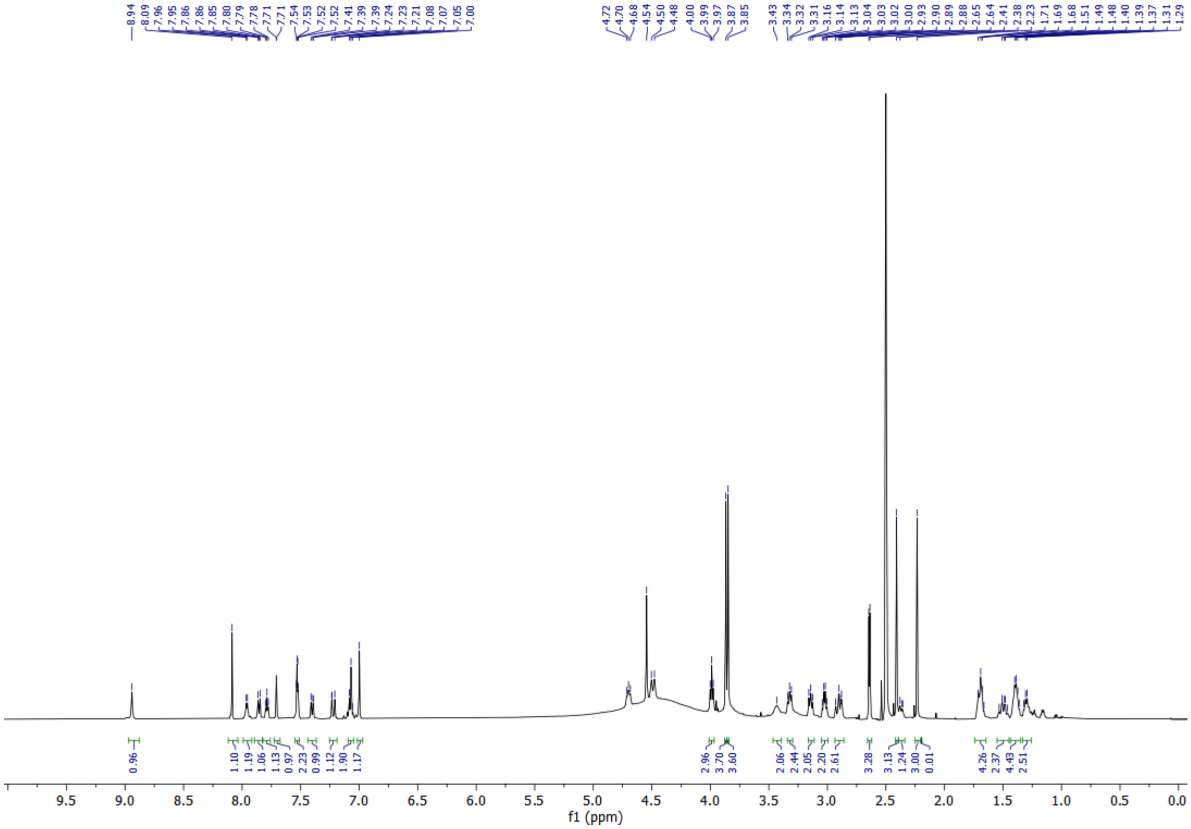

### $^{1}H$ NMR of RCS-02-108:

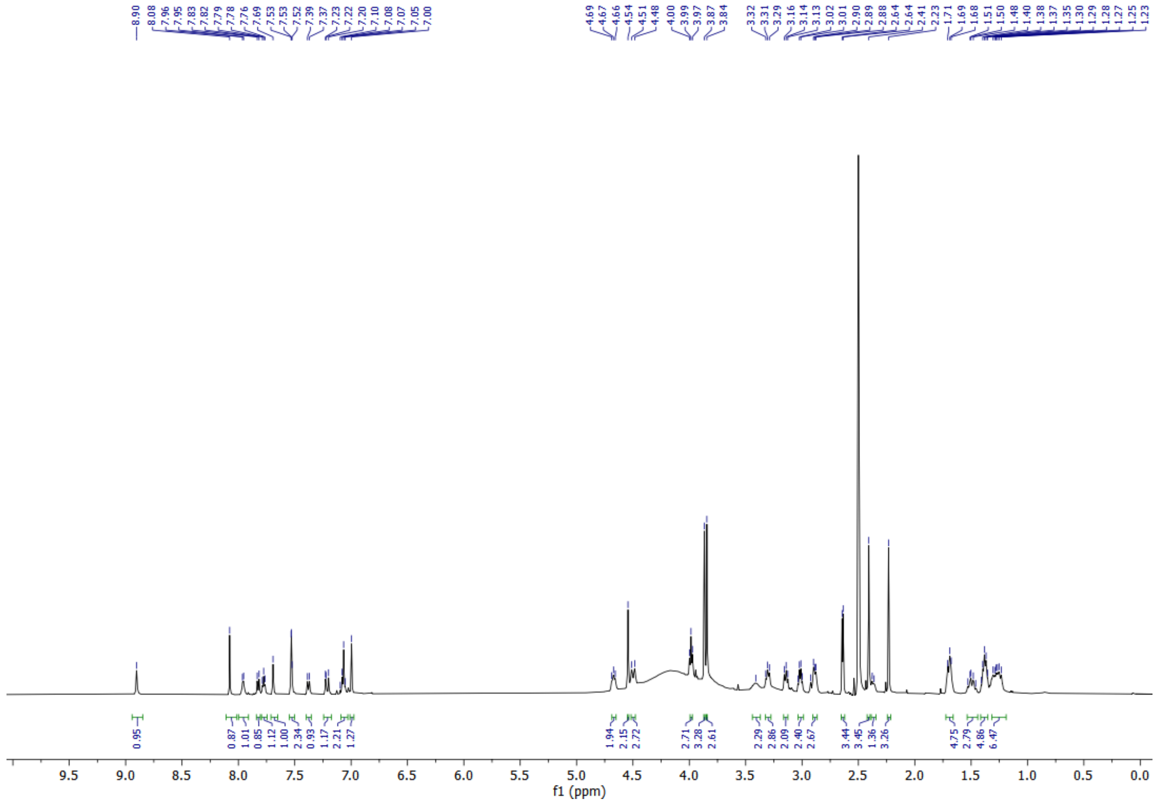

### [1]H NMR of RCS-02-109:

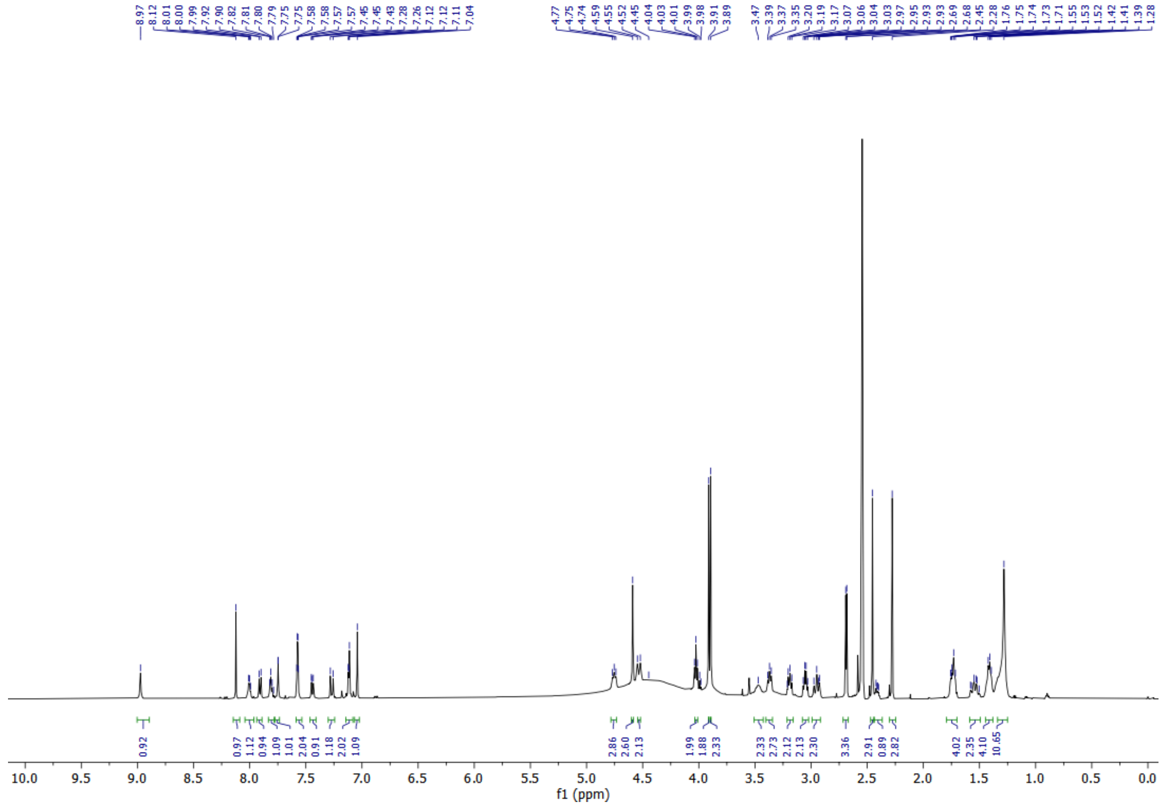

### [1]H NMR of RCS-02-121:

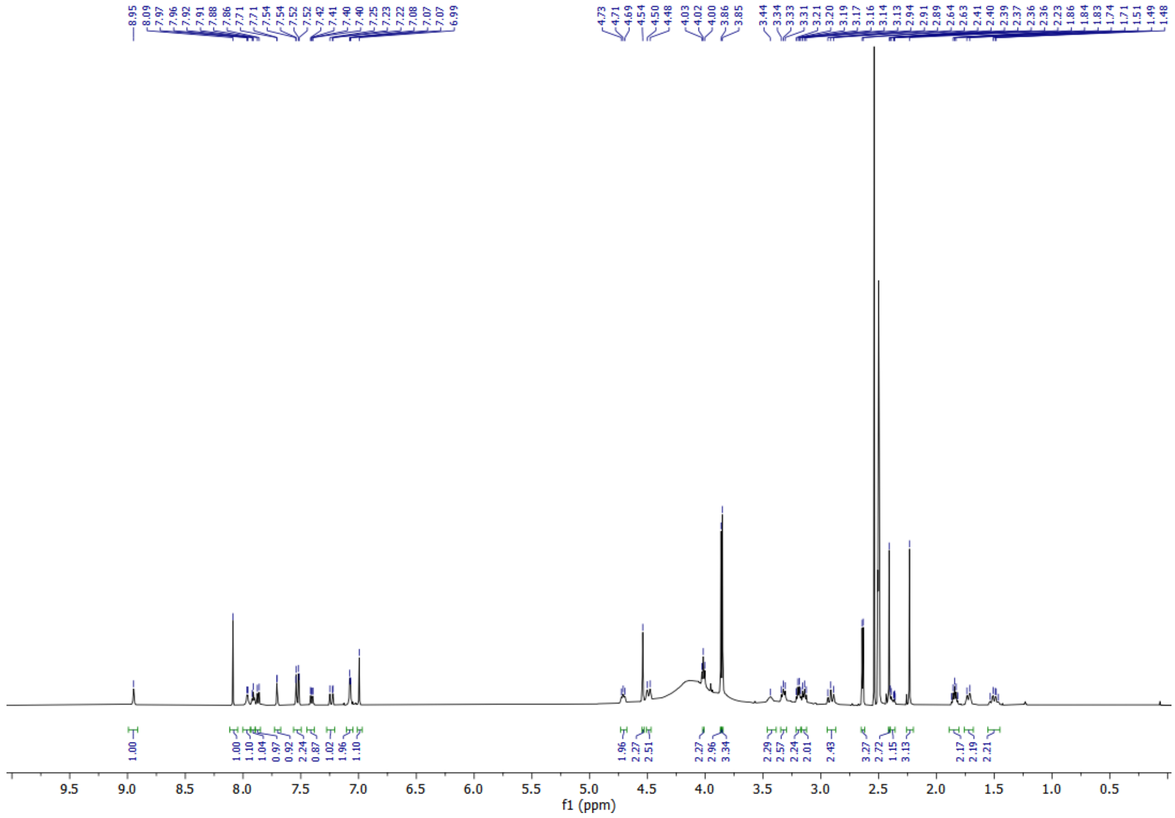

**[1]H NMR of RCS-02-122:**

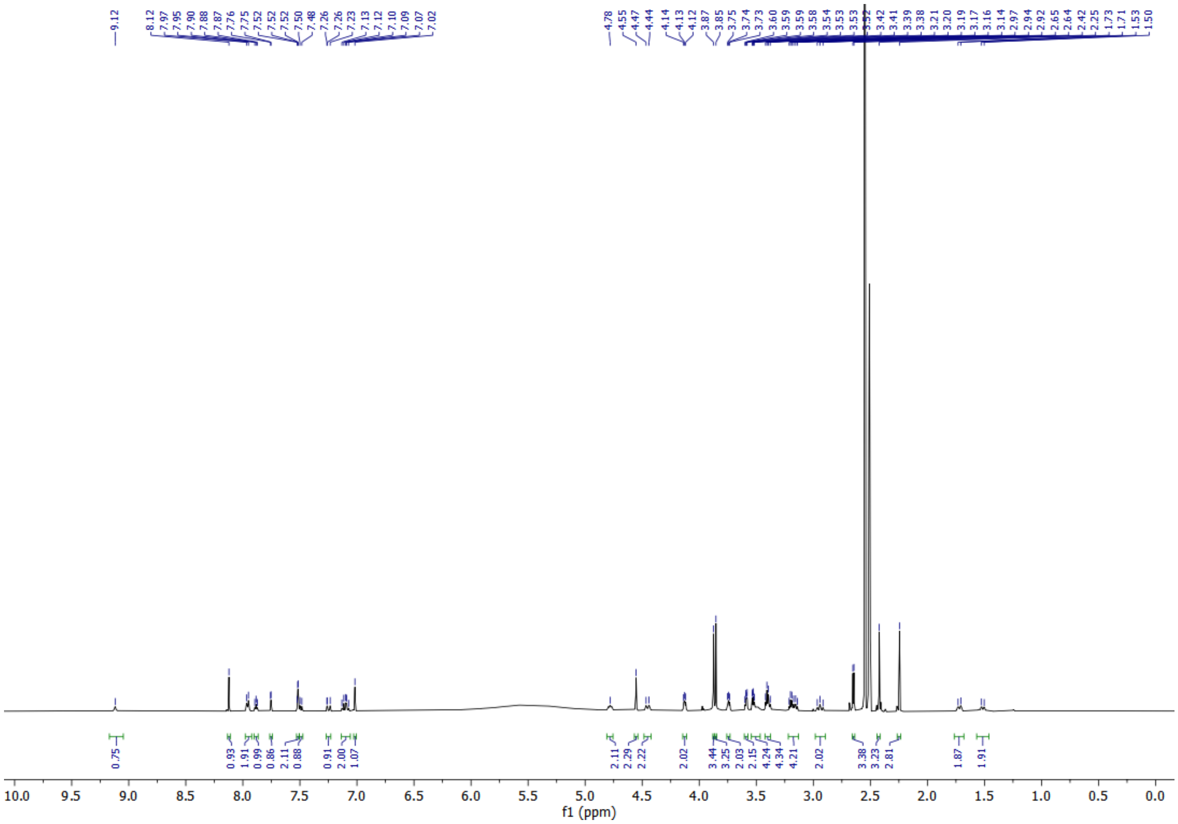

**[1]H NMR of RCS-02-123:**

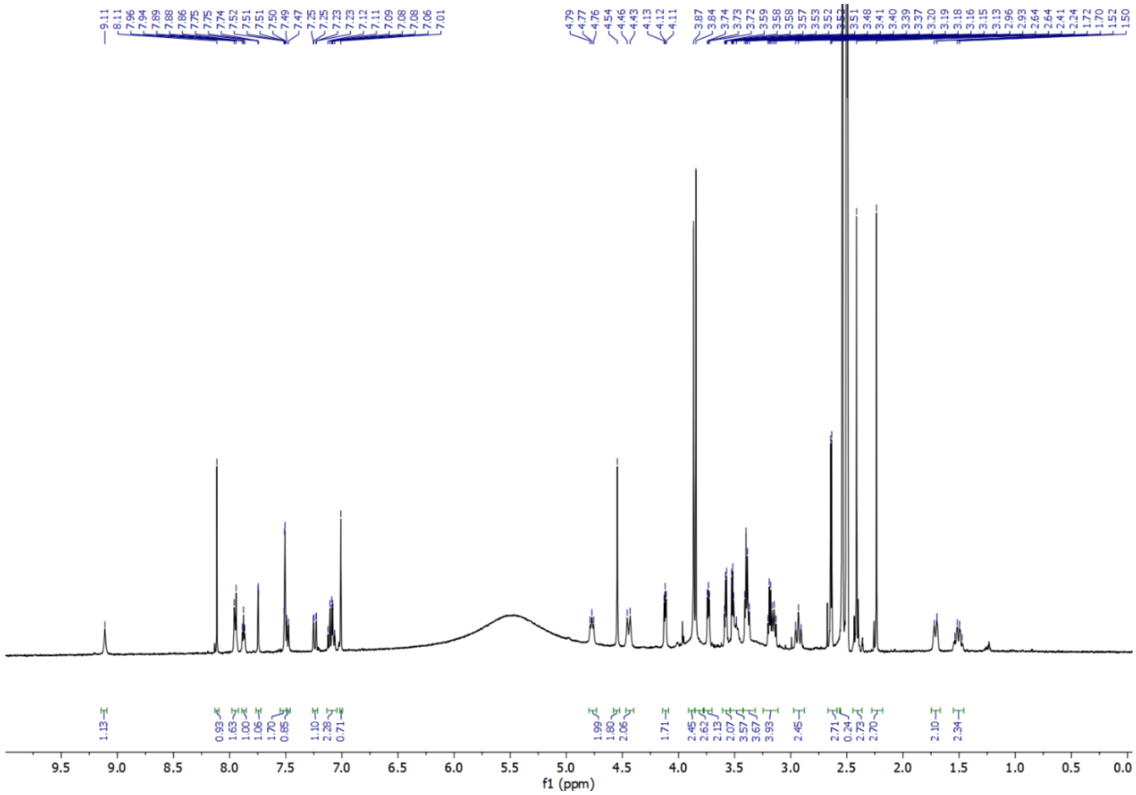

**$^1$H NMR of RCS-02-124:**

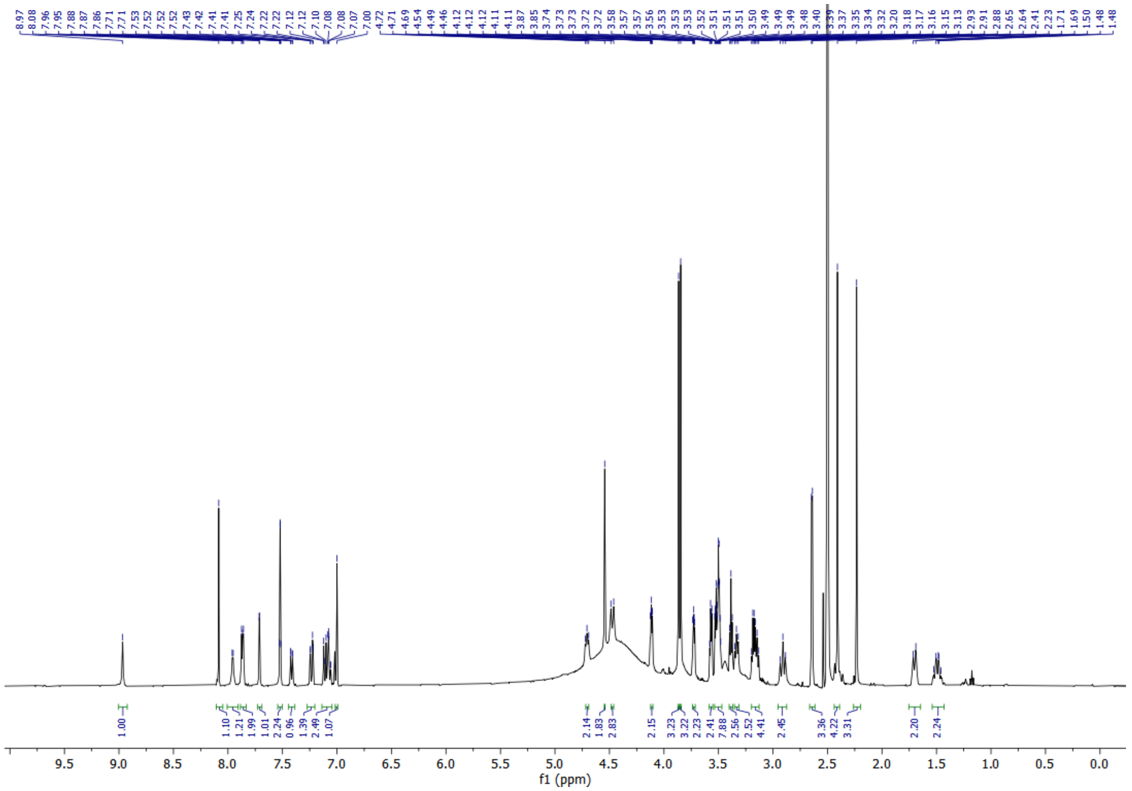

**$^1$H NMR of CCT373566a:**

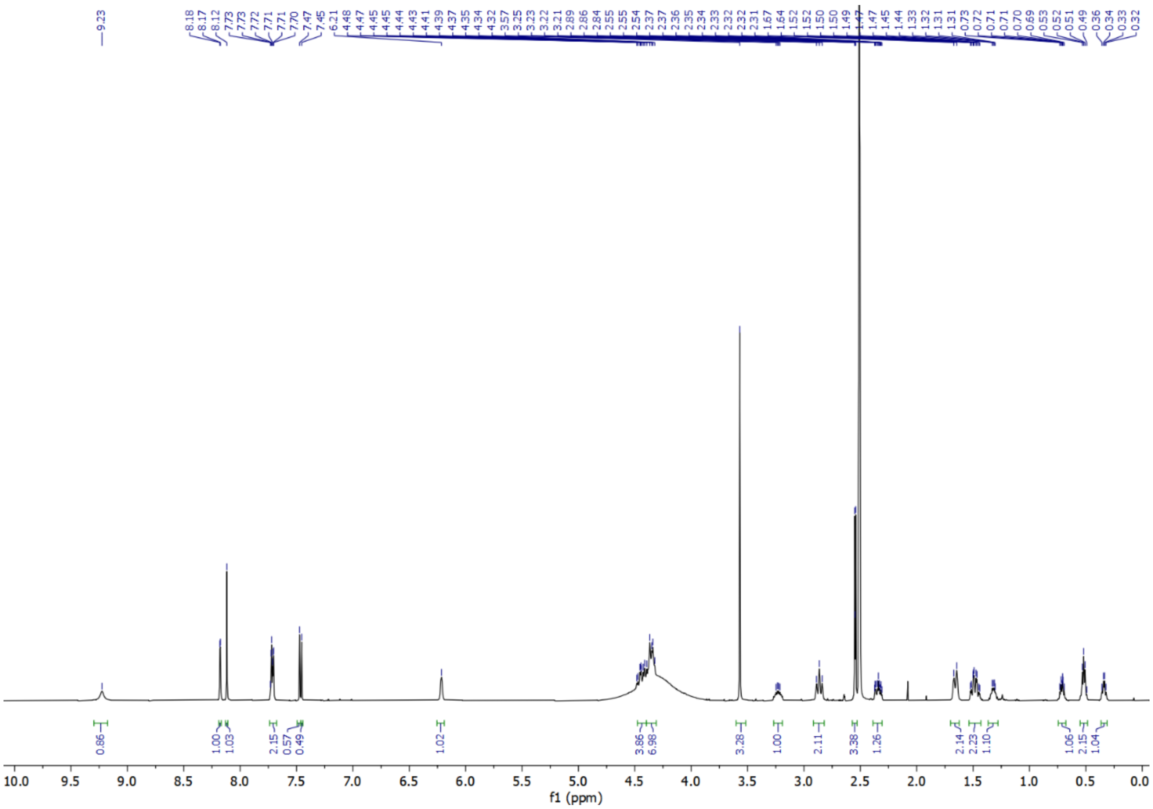
