## Supplemental Figure 11 for "A Bivalent Molecular Glue Linking Lysine Acetyltransferases to Oncogene-induced Cell Death"

**A**

Full scans for Supplemental Fig. 4F

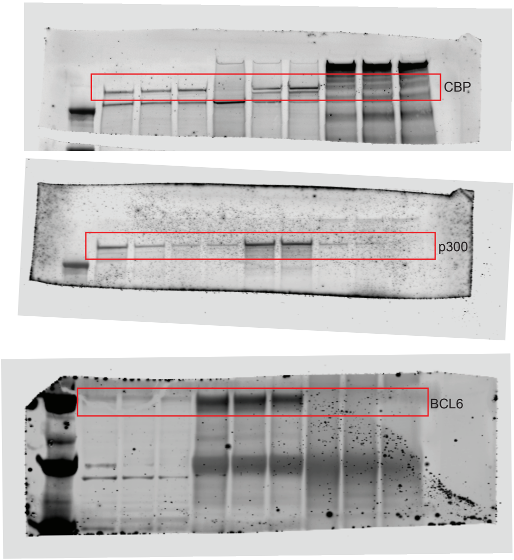


**B**

Full scans for Fig. 3A

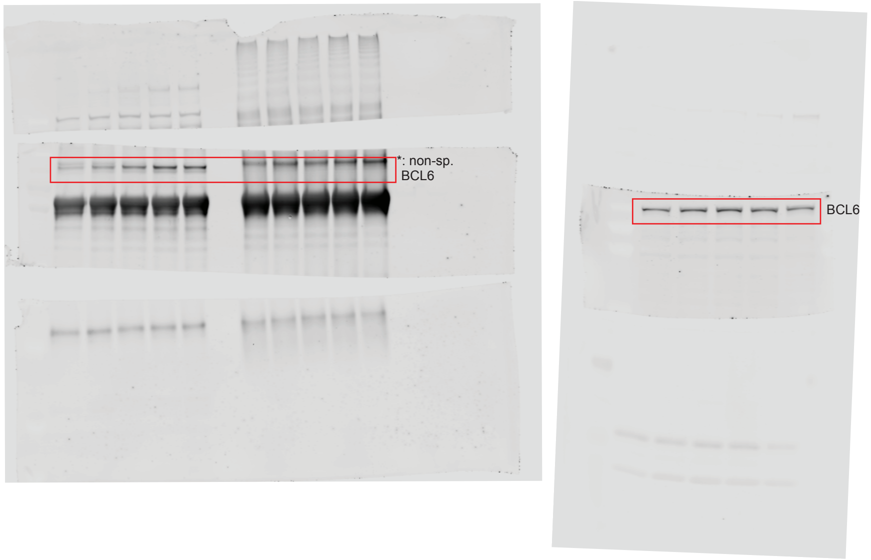


**C**

Full scans for Fig. 3G

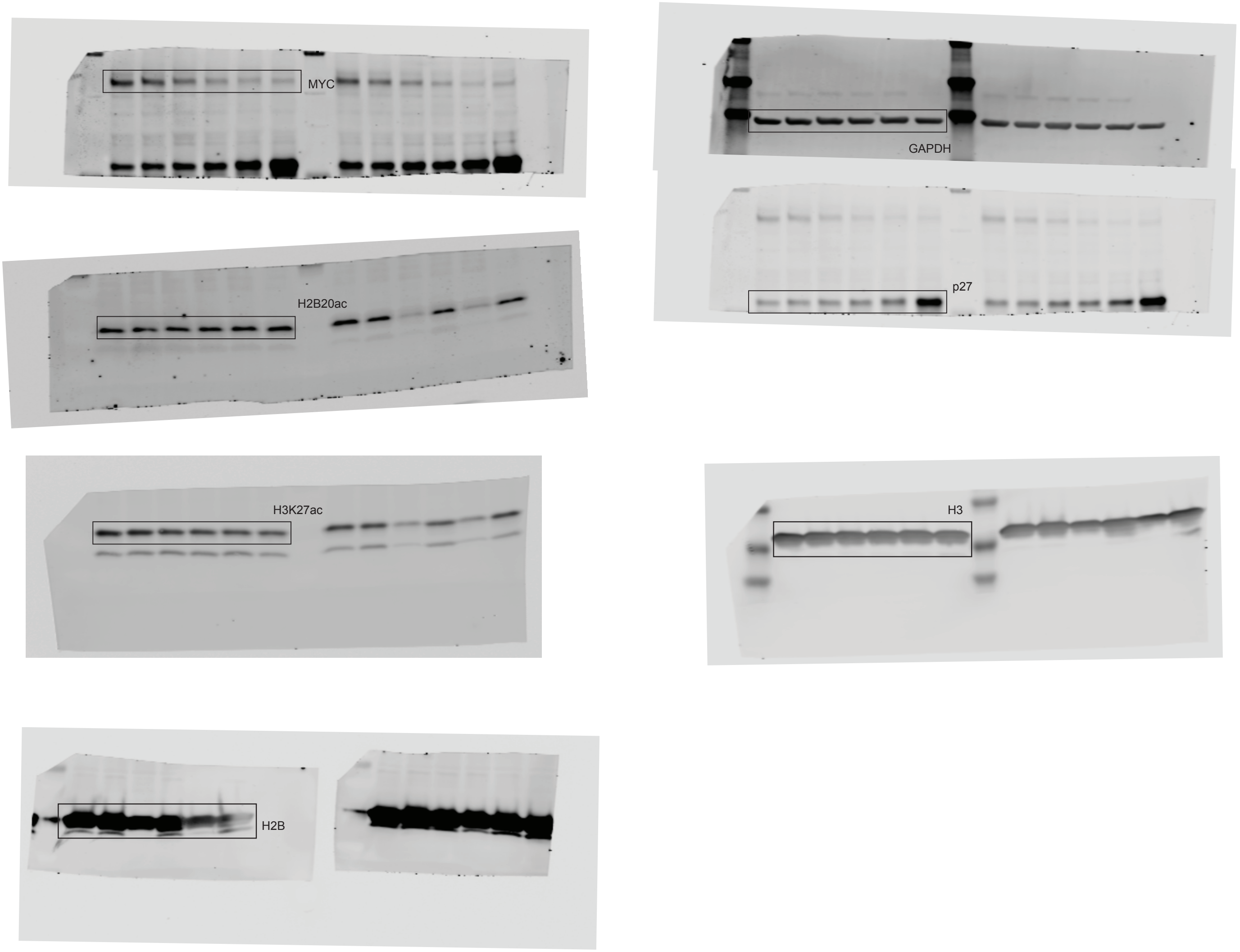

D

Full scans for Fig. 4D

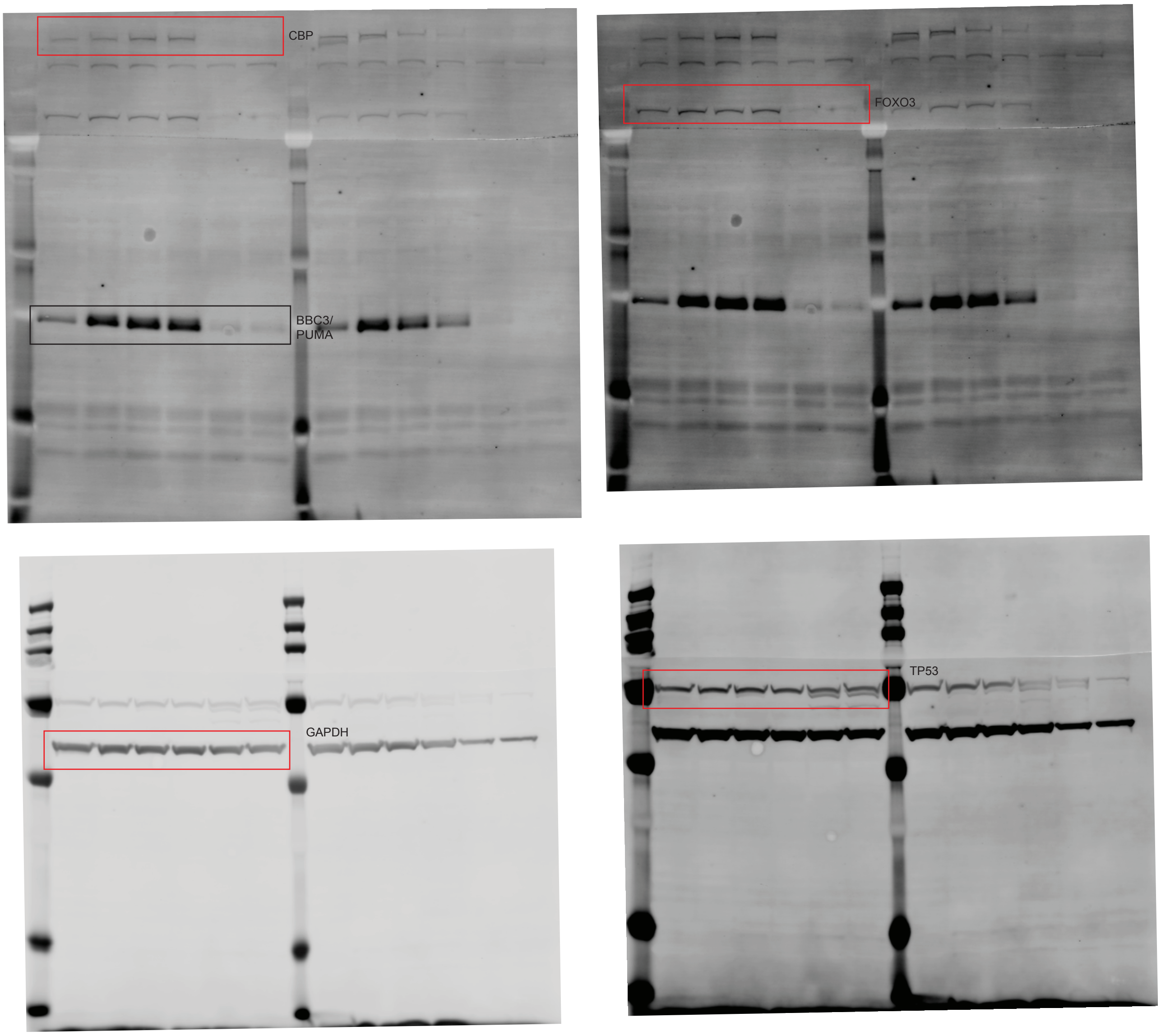


E

Full scans for Supplemental Fig. 7C

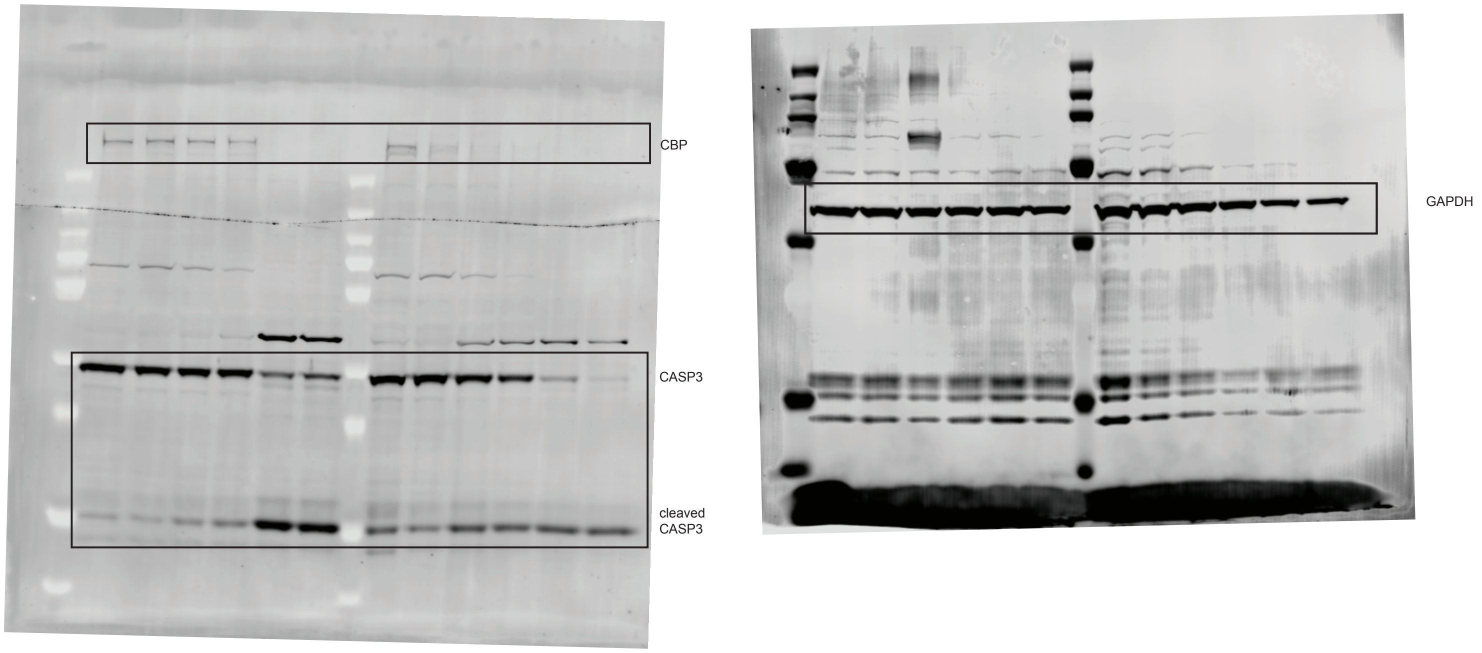

**F**

Full scans for Fig. 5C

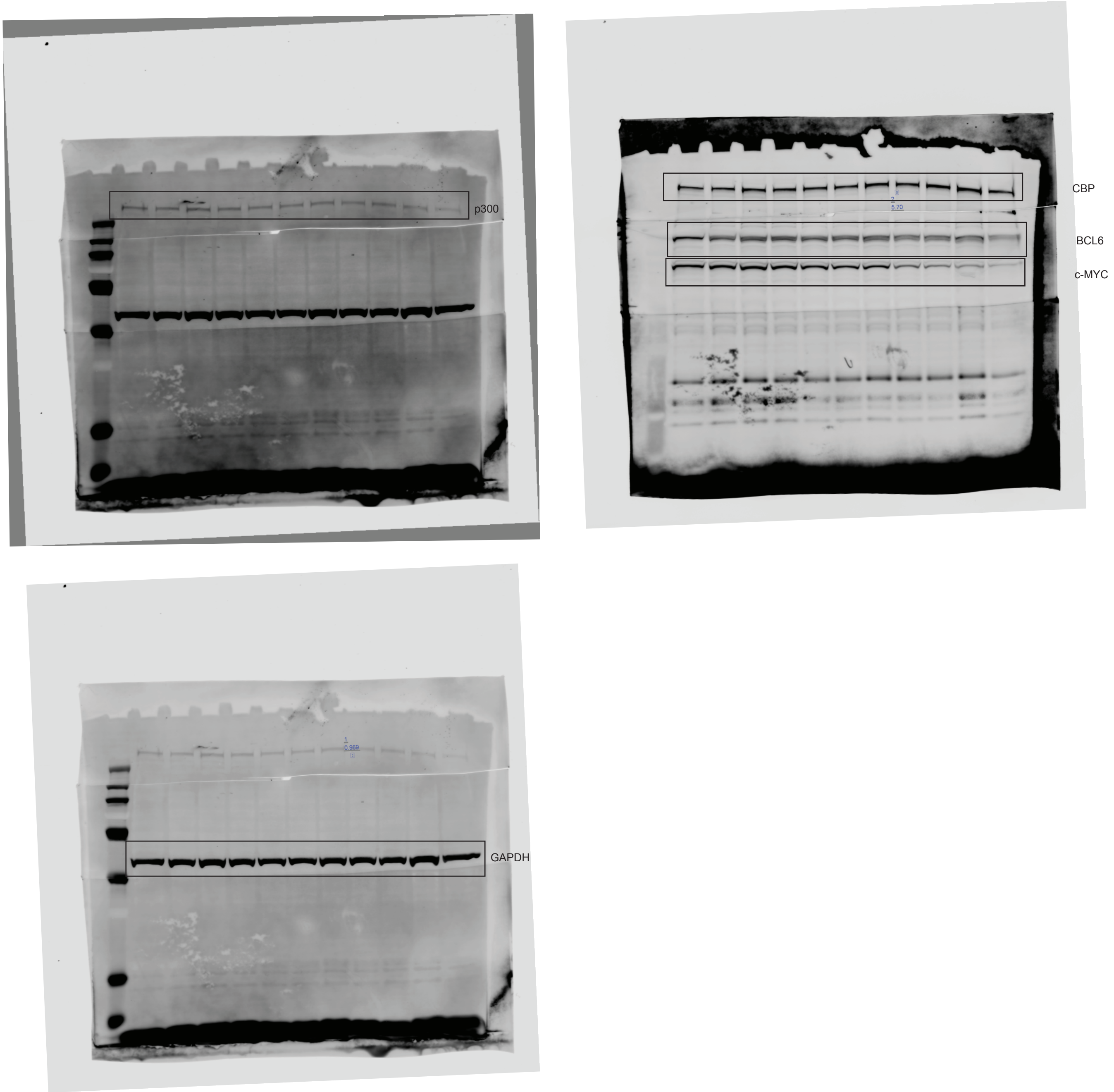


**G**

Full scans for Fig. 5I

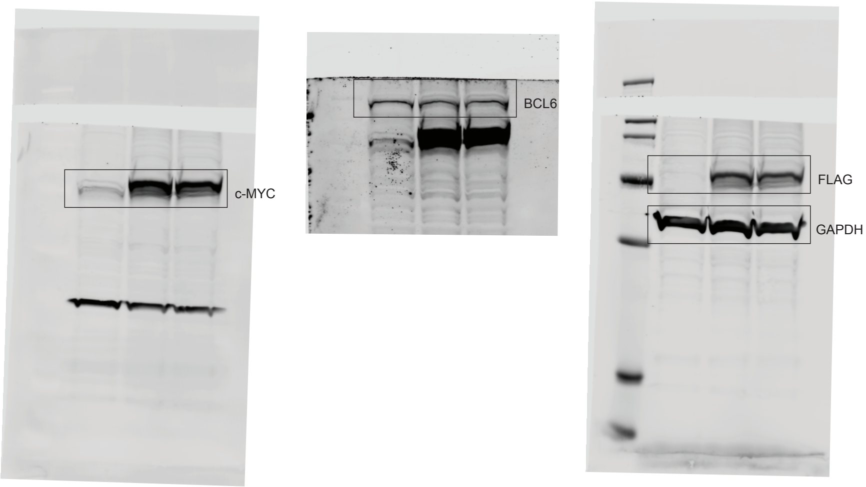


**H**

Full scans for Supplemental Fig. 9A

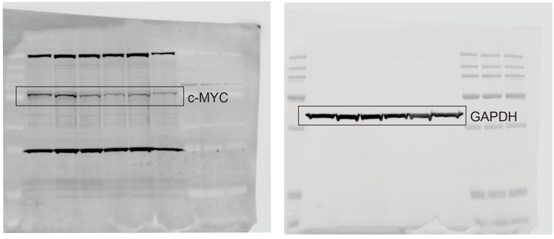

**I**

Full scans for Supplemental Fig. 9B

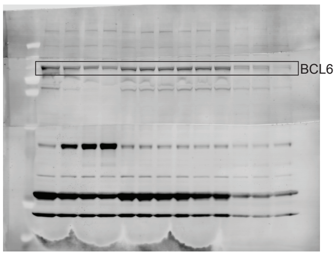


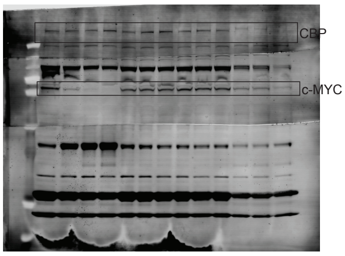


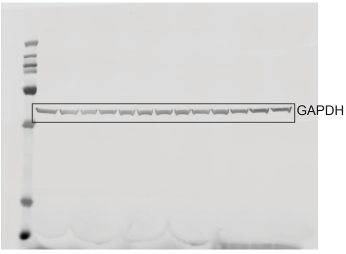
