## Supplementary figures and images for "A Bivalent Molecular Glue Linking Lysine Acetyltransferases to Oncogene-induced Cell Death"

### Supplemental Figure 12

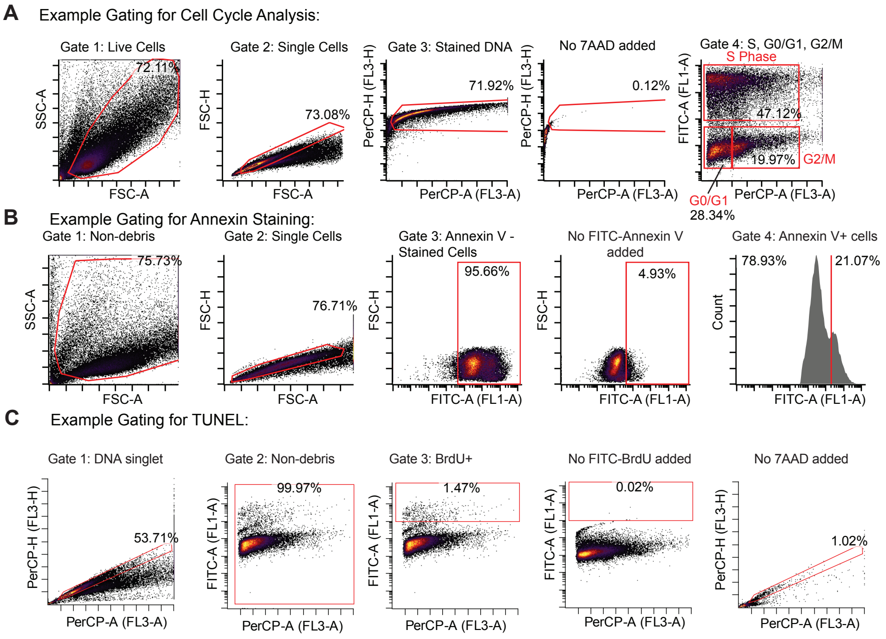
