## Supplemental Table 3 for "A Bivalent Molecular Glue Linking Lysine Acetyltransferases to Oncogene-induced Cell Death"

**Table 1. Data collection and refinement statistics (molecular replacement)**

| | $BCL6^{BTB}$-TCIP3-$p300^{BD}$ |
|---|---|
| **Data collection** | |
| Space group | P 21 21 21 |
| Cell dimensions | |
| *a*, *b*, *c* (Å) | 76.39, 94.79, 97.62 |
| α, β, γ (°) | 90.00, 90.00, 90.00 |
| Resolution (Å) | 60.16-2.01 (2.06-2.01) |
| $R_{merge}$ | .122(1.764) |
| $I / \sigma I$ | 6.7(0.8) |
| Completeness (%) | 99.2(98.6) |
| Redundancy | 5.4(5.5) |
| **Refinement** | |
| Resolution (Å) | 68.00-2.10 (2.18-2.10) |
| No. reflections | 39619 |
| $R_{work}$ / $R_{free}$ | 0.218/0.277(0.41/0.38) |
| No. atoms | |
| Protein | 3906 |
| Ligand | 162 |
| Water | 427 |
| *B*-factors | |
| Protein | 53.25 |
| Ligand | 45.87 |
| Water | 53.81 |
| R.m.s. deviations | |
| Bond lengths (Å) | 0.0074 |
| Bond angles (°) | 1.7090 |

*Values in parentheses are for highest-resolution shell.
